# Chromosomal Duplications of MurZ (MurA2) or MurA (MurA1), Amino Acid Substitutions in MurZ (MurA2), and Absence of KhpAB Obviate the Requirement for Protein Phosphorylation in *Streptococcus pneumoniae* D39

**DOI:** 10.1101/2023.03.26.534294

**Authors:** Ho-Ching Tiffany Tsui, Merrin Joseph, Jiaqi J. Zheng, Amilcar J. Perez, Irfan Manzoor, Britta E. Rued, John D. Richardson, Pavel Branny, Linda Doubravová, Orietta Massidda, Malcolm E. Winkler

## Abstract

GpsB links peptidoglycan synthases to other proteins that determine the shape of the respiratory pathogen *Streptococcus pneumoniae* (pneumococcus; *Spn*) and other low-GC Gram-positive bacteria. GpsB is also required for phosphorylation of proteins by the essential StkP(*Spn*) Ser/Thr protein kinase. Here we report three classes of frequently arising chromosomal duplications (≈21-176 genes) containing *murZ* (MurZ-family homolog of MurA) or *murA* that suppress Δ*gpsB* or Δ*stkP*. These duplications arose from three different repeated sequences and demonstrate the facility of pneumococcus to modulate gene dosage of numerous genes. Overproduction of MurZ or MurA alone or overexpression of MurZ caused by Δ*khpAB* mutations suppressed Δ*gpsB* or Δ*stkP* phenotypes to varying extents. Δ*gpsB* and Δ*stkP* were also suppressed by MurZ amino-acid changes distant from the active site, including one in commonly studied laboratory strains, and by truncation or deletion of the homolog of IreB(ReoM). Unlike in other Gram-positive bacteria, MurZ is predominant to MurA in pneumococcal cells. However, Δ*gpsB* and Δ*stkP* were not suppressed by Δ*clpCP*, which did not alter MurZ or MurA amounts. These results support a model in which regulation of MurZ and MurA activity, likely by IreB(*Spn*), is the only essential requirement for protein phosphorylation in exponentially growing D39 pneumococcal cells.

## 1 INTRODUCTION

Bacterial survival depends on the regulation of the synthesis and assembly of the peptidoglycan (PG) cell wall (Rohs & Bernhardt, 2021, Egan *et al*., 2020, Kumar *et al*., 2022). PG determines cell shape and morphology and protects against osmotic stress(Booth & Lewis, 2019, Egan *et al*., 2020, Garde *et al*., 2021). The proteins that carry out the numerous steps of PG synthesis are major targets for clinically relevant antibiotics, for which widespread resistance has developed (Booth & Lewis, 2019, Egan *et al*., 2020, Bush & Bradford, 2016). In Gram-positive bacteria, such as *Streptococcus pneumoniae* (pneumococcus; *Spn*), the PG cell wall also provides a scaffolding for attachment of capsule, wall teichoic acids, and extracellular proteins and virulence factors (Booth & Lewis, 2019, Briggs *et al*., 2021, Kumar *et al*., 2022). *S. pneumoniae* is a commensal bacterium of the human nasopharynx and a major opportunistic respiratory-tract pathogen that kills millions of people annually worldwide, including following influenza and COVID-19 infections (Sender *et al*., 2021, Cox *et al*., 2020, Weiser *et al*., 2018). *S. pneumoniae* is continuing to acquire antibiotic resistance to a broad range of antibiotics and is now classified as a “superbug” by the CDC and WHO (WHO, 2017, CDC *et al*., 2019).

The GpsB protein is a major regulator of PG synthesis in low-GC Gram-positive bacteria (Claessen *et al*., 2008, Rismondo *et al*., 2016, Cleverley *et al*., 2019, Fleurie *et al*., 2014, Rued *et al*., 2017). In *Bacillus subtilis* (*Bsu*), Δ*gpsB* results in growth and morphological abnormality in high salt media and synthetic lethality with Δ*ezrA* or Δ*ftsA* (Claessen *et al*., 2008), while in *Listeria monocytogenes* (*Lmo*), Δ*gpsB* causes marked growth and division defects at 37°C and is lethal at 42°C (Rismondo *et al*., 2016). Δ*gpsB* mutants of *Enterococcus faecalis* (*Efa*) also show growth defects at 45°C, but grow normally at 37°C (Minton *et al*., 2022). In contrast, in derivatives of serotype-2 *S. pneumoniae* D39 progenitor strains, *gpsB* is essential at 37°C, and GpsB depletion leads to drastic cell enlargement and elongation, incomplete closure of septal division rings, and eventual cell lysis (Land *et al*., 2013, Rued *et al*., 2017). Depletion of GpsB in *S. aureus* (*Sau*), however, arrests cell division without coincident cell enlargement and ultimately causes aberrant membrane accumulation (Eswara *et al*., 2018).

Combined studies indicate that GpsB plays species-specific roles in regulating PG synthesis (Cleverley *et al*., 2019, Hammond *et al*., 2019). Based on genetic and biochemical studies, one role shared by GpsB in different bacteria is as an adaptor that docks PG synthases to other cell-wall enzymes and scaffold proteins to form complexes for division and septal and lateral PG synthesis (Rued *et al*., 2017, Cleverley *et al*., 2019, Halbedel & Lewis, 2019, Sacco *et al*., 2022). Binding between GpsB homologs and Class A PBPs, including PBP1(*Bsu*), PBPA1(*Lmo*), and aPBP2a(*Spn*) and Class C PBP4(*Sau*) occurs by a conserved mechanism, wherein Arg residues in amino-terminal, cytoplasmic microdomains of the PBPs bind to a specific site in the amino-terminal domain of GpsB (Cleverley *et al*., 2019, Sacco *et al*., 2022). Species-specific binding to other subsets of PG synthesis and cell division proteins occurs at other surfaces in GpsB homologs (Cleverley *et al*., 2019). For example, besides interacting with aPBP2a, GpsB(*Spn*) is in complexes with EzrA, MreC, StkP, and possibly bPBP2x, bPBP2b, and aPBP1a, but not with FtsZ and FtsA (Rued *et al*., 2017, Cleverley *et al*., 2019). Unlike other GpsB homologs, GpsB(*Sau*) binds to a non-conserved C-terminal tail of FtsZ, which affects FtsZ polymerization (Sacco *et al*., 2022). GpsB(*Sau*) also interacts with teichoic acid biogenesis proteins through binding motifs that are not widely conserved in GpsB from other bacteria (Eswara *et al*., 2018, Hammond *et al*., 2022). The significance of GpsB in maintaining cell wall integrity during antibiotic stress in *S. pneumoniae* was underscored by a genome-wide association study of clinical isolates that revealed significant correlation of β-lactam resistance and the presence of *gpsB* variants (Mobegi *et al*., 2017).

An additional important regulatory function of GpsB is the maintenance of protein phosphorylation mediated by conserved homologues of serine/threonine kinases, StkP(*Spn*), PrkC(*Bsu*), and IreK(*Efa*) (Rued *et al*., 2017, Pompeo *et al*., 2015, Fleurie *et al*., 2014, Minton *et al*., 2022). In *S. pneumoniae*, phosphorylation of StkP and other StkP substrates is significantly reduced in Δ*gpsB* mutants of laboratory strains Rx1, R6, or R800 or upon depletion of GpsB in D39-derived strains (Rued *et al*., 2017, Fleurie *et al*., 2014). The link between GpsB function and protein phosphorylation was further supported in D39-derived strains by the finding that Δ*gpsB* is suppressed by mutations that inactivate the cognate PhpP Ser/Thr protein phosphatase, such as *phpP*(G229D), which restore protein phosphorylation (Rued *et al*., 2017). Notably, *phpP*(G229D), restores the growth and cell morphology of Δ*gpsB* mutants to nearly those of WT cells (Rued *et al*., 2017), indicating that GpsB mediates StkP phosphorylation of one or more proteins required for exponential growth of *S. pneumoniae*.

StkP(*Spn*) belongs to the subfamily of eukaryotic-type Ser/Thr kinases (ESTKs) and together with cognate PP2C-type phosphatase PhpP(*Spn*), constitutes a signaling system (Echenique *et al*., 2004, Novakova *et al*., 2005). Based on phenotypes of Δ*stkP* mutants in different genetic backgrounds, StkP has been implicated in the regulation of cell growth and cell division (Beilharz *et al*., 2012, Giefing *et al*., 2010, Hirschfeld *et al*., 2019, Fleurie *et al*., 2012), competence (Echenique *et al*., 2004, Saskova *et al*., 2007, Rued *et al*., 2017), stress resistance (Saskova *et al*., 2007)), acidic stress-induced lysis (Pinas *et al*., 2018), capsule synthesis and virulence (Echenique *et al*., 2004, Kant *et al*., 2023), pilus expression and adherence (Herbert *et al*., 2015), and β-lactam susceptibility (Dias *et al*., 2009). However, the essentiality of both *gpsB*(*Spn*) and *stkP*(*Spn*) has been controversial. Based on numerous studies of common laboratory strains R6 (and its derivative R800) and Rx1, *gpsB* and *stkP* have generally been classified as non-essential (Rued *et al*., 2017, Fleurie *et al*., 2014), despite variations in growth properties and cell morphologies consistent with the presence of suppressor mutations (Beilharz *et al*., 2012, Fleurie *et al*., 2012, Massidda *et al*., 2013, Rued *et al*., 2017, Ulrych *et al*., 2021, Vollmer *et al*., 2019). In contrast, *gpsB* and *stkP* are essential in D39-derived strains (Land *et al*., 2013, Rued *et al*., 2017), from which the laboratory strains were originally derived (Cuppone *et al*., 2021, Lanie *et al*., 2007, Santoro *et al*., 2019). Depletion and transformation experiments clearly indicate that *gpsB* is essential in D39 strains and that Δ*gpsB* mutants accumulate suppressor mutations (Land *et al*., 2013, Rued *et al*., 2017). In contrast, Δ*stkP* mutants are unstable and rapidly acquire suppressor mutations that cause faster growth (Beilharz *et al*., 2012, Rued *et al*., 2017, Ulrych *et al*., 2021). Moreover, the primary cell morphology changes caused by StkP depletion remain unknown, as do mutations in the common laboratory strains that bypass the essentiality of *gpsB* and *stkP*.

Multiple proteins phosphorylated by StkP(*Spn*) have been identified in studies comparing global phosphoproteomes of Δ*stkP* mutants with that of their isogenic encapsulated D39 (*cps*^+^) or unencapsulated D39 (Δ*cps*) parent strains (Hirschfeld *et al*., 2019, Sun *et al*., 2010, Ulrych *et al*., 2021). Several proteins associated with division and PG synthesis are phosphorylated in pneumococcal cells, including DivIVA (Novakova *et al*., 2010, Fleurie *et al*., 2012), MapZ (LocZ) (Fleurie *et al*., 2014, Holeckova *et al*., 2014), KhpB (Jag/EloR) (Zheng *et al*., 2017, Ulrych *et al*., 2016, Stamsas *et al*., 2017), MacP (Fenton *et al*., 2018), FtsZ (Ulrych *et al*., 2021), GpsB (Hirschfeld *et al*., 2019, Ulrych *et al*., 2021), MpgA (formerly MltG(*Spn*) (Hirschfeld *et al*., 2019, Taguchi *et al*., 2021, Ulrych *et al*., 2021), and IreB (Ulrych *et al*., 2021). In addition, the pattern of protein phosphorylation changes between exponentially and antibiotic stressed cells (Ulrych *et al*., 2021). Nevertheless, the roles of phosphorylation of individual proteins in growing D39 cells remains problematic, because phosphoablative and phosphomimetic mutants of cell division and PG synthesis proteins, such as DivIVA, MapZ, and KhpB, have not consistently shown aberrant phenotypes in exponentially growing cultures (Holeckova *et al*., 2014, Massidda *et al*., 2013, Zheng *et al*., 2017, Manuse *et al*., 2016, Fleurie *et al*., 2012, Grangeasse, 2016). It has not yet been determined which StkP-phosphorylation proteins are required for normal exponential growth of D39 strains.

Besides *phpP* null mutations, Δ*gpsB*(*Spn*) was suppressed by two large chromosomal duplications that also contained deletions (Rued *et al*., 2017). Notably, these duplications contain *murZ* (Rued *et al*., 2017, Wamp *et al*., 2020), which encodes one of two homologs of the UDP-N-acetylglucosamine 1-carboxyvinyltransferase that converts PEP and UDP-GlcNAc to Pi and UDP-N-acetyl-3-O-(1-carboxyvinyl)-alpha-D-glucosamine in the first committed step in the synthesis of the PG precursor Lipid II (Brown *et al*., 1995, Zhou *et al*., 2022). Like other low-GC Gram-positive bacteria, *S. pneumoniae* encodes two distinct homologs of this enzyme (Fig. 1) (Blake *et al*., 2009, Chan *et al*., 2022, Du *et al*., 2000, Kedar *et al*., 2008, Kock *et al*., 2004, Mascari *et al*., 2022, Vesic & Kristich, 2012). The two homologs in *S. pneumoniae* strains were annotated as MurZ (MurA2) (Spd_0967) and MurA (MurA1) (*Spn*)(Spd_1764) (Hoskins *et al*., 2001) (Fig. 1). The MurA-family homolog, which is the sole enzyme present in Gram-negative bacteria (Brown *et al*., 1995, Du *et al*., 2000, Hummels *et al*., 2023, Zhou *et al*., 2022), often plays a predominant enzymatic role in Gram-positive bacteria and is essential in *B. subtilis*, *B. anthracis*, and *L. monocytogenes* (Kock *et al*., 2004, Kedar *et al*., 2008, Rismondo *et al*., 2017), and required for normal growth of *E. faecalis* and *S. aureus* (Vesic & Kristich, 2012, Blake *et al*., 2009, Mascari *et al*., 2022). MurZ(*Spn*) and MurA(*Spn*) have a synthetic lethal relationship, where one homolog functions in the absence of the other, but both homologs cannot be deleted in the same strain(Du *et al*., 2000). Absence of MurAA(*Efa*) and MurAB(*Efa*) or MurA(*Sau*) and MurZ(*Sau*) is also synthetically lethal, where MurA-family MurAA(*Efa*) or MurA(*Sau*) is catalytically dominant in cells (Blake *et al*., 2009, Vesic & Kristich, 2012, Mascari *et al*., 2022). In contrast, previous biochemical studies demonstrated that MurZ(*Spn*) purified from strain R6 has a considerably higher (≈3.5-fold) catalytic efficiency (*k_cat_*/*K_m_*) for UDP-GlcNAc than MurA(*Spn*) (Du *et al*., 2000). Consistent with these kinetic results, a Δ*murZ*(*Spn*) mutant substantially reduced the circumferential velocity of the bPBP2x:FtsW septal PG synthase, without changing the rate of FtsZ treadmilling (Perez *et al*., 2019). However, the relative contributions of MurZ and MurA to pneumococcal growth and physiology remain unknown.

**Figure 1.**
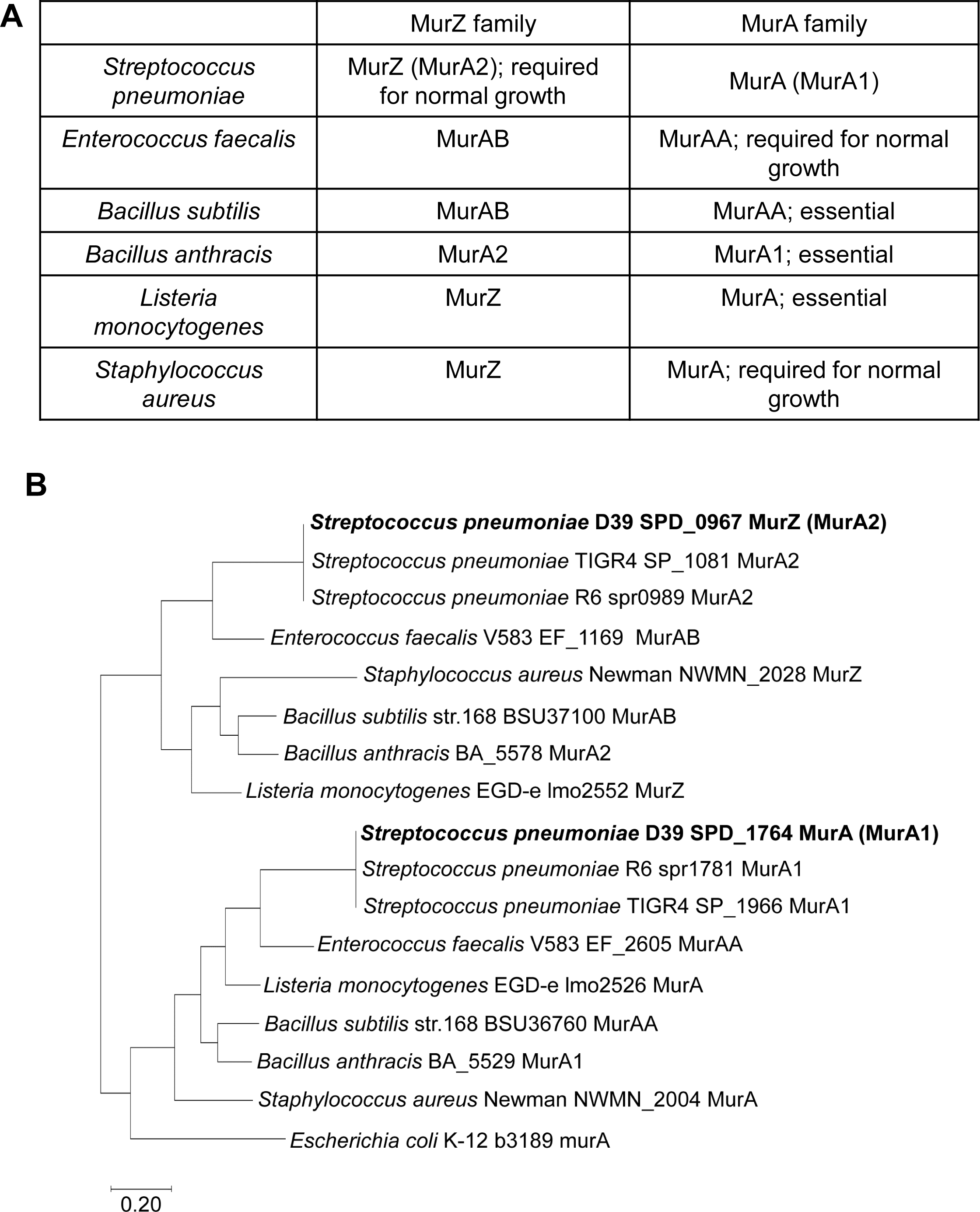
Two evolutionary branches of the MurA-family and MurZ-family homologs of *S. pneumoniae* and other Gram-positive bacteria. (A) Nomenclature and function of MurA and MurZ homologs from six Gram-positive bacteria *S. pneumoniae* (*Spn*) (Du *et al*.), *E. faecalis* (*Efa*) (Vesic & Kristich, 2012), *B. subtilis* (*Bsu*) (Kock *et al*., 2004), *B. anthracis* (*Ban*) (Kedar *et al*., 2008), *L. monocytogenes* (*Lmo*) (Rismondo *et al*., 2017), and *S. aureus* (*Sau*) (Blake *et al*., 2009). (B) Partial evolutionary tree of the MurZ-family and MurA-family homologs from five Gram-positive bacteria *Spn*, *Efa*, *Sau, Bsu* and *Lmo*, and the single MurA-homolog in Gram-negative bacterium *E. coli*. MurZ(*Spn*) (Spd_0967)(*Spn*) is phylogenetically closely related to MurAB(*Efa*), MurAB(*Bsu*), MurZ(*Sau*), and MurZ(*Lmo*), while MurA(*Spn*) (Spd_1764) is phylogenetically closely related to MurAA(*Efa*), MurAA(*Bsu*), MurA(*Sau*), and MurA(*Lmo*). Note that in the original annotation of the *S. pneumoniae* D39 genome, the MurZ(*Spn*) homolog was called “MurA1” and the MurA(*Spn*) homolog was called “MurA2” (Lanie *et al*., 2007, Slager *et al*., 2018). For consistency with the field, the revised nomenclature in the table is used.

Concurrent with our previous study (Rued *et al*., 2017) and the work reported here on suppression of Δ*gpsB* in *S. pneumoniae* D39 strains, suppressors of Δ*gpsB* were isolated in *L*. *monocytogenes* (Rismondo *et al*., 2017, Wamp *et al*., 2020). Remarkably, these studies by Rismondo, Wamp and colleagues showed that Δ*gpsB*(*Lmo*) is suppressed by mutations in the *murZ*(*Lmo*) homolog, *reoY*(*Lmo*), which encodes a protein of unknown function in *Bacillus* and *Enterococcus* species, *clpC*(*Lmo*), which encodes an ATPase subunit of the ClpP protease, *reoM*(*Lmo*), which encodes a small protein that is phosphorylated by the PrkA(*Lmo*) kinase, and *prpC*(*Lmo*), which encodes the cognate phosphatase to PrkA(*Lmo*) (Rismondo *et al*., 2016, Wamp *et al*., 2020, Wamp *et al*., 2022). In parallel work, Vesić and Kristich linked MurAA(Efa) function to protein phosphorylation by demonstrating that overexpression of MurAA(*Efa*) restored cephalosporin resistance to a mutant lacking the IreK(*Efa*) kinase (Vesic & Kristich, 2012).

These and other supporting data have led to a model whereby regulation of MurA(*Lmo*) stability is mediated by the phosphorylation level of ReoM(*Lmo*) (Wamp *et al*., 2022, Wamp *et al*., 2020). According to this model, unphosphorylated ReoM(*Lmo*) may act as an adaptor, along with ReoY(*Lmo*) and MurZ(*Lmo*), to direct MurA(*Lmo*) degradation by the ClpCP(*Lmo*) protease. Phosphorylation of ReoM(*Lmo*) by PrkA(*Lmo*) in response to PG signals and stress are postulated to increase MurA(*Lmo*) amount and increase PG precursor synthesis for PG synthases in response to beta-lactam antibiotics. In support of this model, overexpression of MurA(*Lmo*), but not MurZ(*Lmo*), suppressed Δ*gpsB*(*Lmo*), and amino acid changes in MurA(*Lmo*) were identified that uncouple ReoM(*Lmo*)-mediated degradation by ClpCP(*Lmo*) (Wamp *et al*., 2022). Moreover, *reoM*(*Lmo*), *reoY*(*Lmo*), and *clpC*(*Lmo*) mutations suppress the conditional lethality of Δ*gpsB* as well as the lethality of Δ*prkA* in one genetic background of *L. monocytogenes* (Wamp *et al*., 2022, Wamp *et al*., 2020). Notably, Kelliherr and colleagues confirmed this general model by isolating suppressors in this set of genes that decrease sensitivity of Δ*prkA*(*Lmo*) to β-lactam antibiotics and relieve infection-linked phenotypes (Kelliher *et al*., 2021). However, a link between general protein phosphorylation and GpsB function has not been reported in *L. monocytogenes*, and it was speculated that lack of GpsB(*Lmo*) leads to misregulation of Class A PBP function that somehow signals to the PrkA(*Lmo*) kinase (Wamp *et al*., 2020).

In this paper, we expand our previous study of Δ*gpsB* suppression in *S. pneumoniae* D39. We report that most Δ*gpsB*(*Spn*) and Δ*stkP* suppressors are duplications of regions containing *murZ*(*Spn*) or *murA*(*Spn*). We show that these duplications range from ≈20 gene to >150 genes and are anchored by different repeat sequences flanking *murZ*(*Spn*) or *murA*(*Spn*), attesting to remarkable genetic plasticity in the pneumococcal chromosome (Slager *et al*., 2018). Consistent with the isolation of these duplication suppressors, we show that overexpression of MurZ(*Spn*) or MurA(*Spn*) suppressed Δ*gpsB*(*Spn*) or Δ*stkP* lethality. In addition, lack of the pneumococcal KhpAB RNA-binding protein resulted in overexpression of MurZ(*Spn*), which accounts for suppression of Δ*gpsB*(*Spn*) by Δ*khpA*(*Spn*) or *khpB*(*Spn*). Yet, determinations of growth, morphology, and sensitivity to fosfomycin indicated that MurZ(*Spn*) is predominant to MurA(*Spn*), although their cellular amounts are approximately equal.

In addition, we isolated mutations containing amino-acid changes in a region of MurZ(*Spn*) away from its catalytic site that suppressed Δ*gpsB*(*Spn*), without restoring protein phosphorylation, or Δ*stkP*. Other amino acid changes in this region of MurZ(*Spn*) acted as suppressors, including one present in laboratory strains R6 and Rx1. An isolated stop-codon mutation near the end of *ireB*(*Spn*) and a constructed markerless Δ*ireB*(*Spn*) deletion also suppressed Δ*gpsB*(*Spn*) or Δ*stkP*. However, genetic suppression and western blotting experiments indicated that MurZ(*Spn*) and MurA(*Spn*) are not degraded by the ClpCP(*Spn*) protease. Tn-seq and depletion experiments further showed that StkP is essential in D39 strains and that the primary morphology phenotype caused by lack of StkP is a defect in division septation, resulting in longer, but not wider, cells. Altogether, these findings support the conclusion that GpsB(*Spn*) and StkP are essential in exponentially growing *S. pneumoniae* D39 cells, because phosphorylation is required for the regulation of MurZ(*Spn*) and MurA(*Spn*) activity, but not their amounts.

## 2 RESULTS

### 2.1 Chromosome duplications containing *murZ* or *murA* are present in Δ*gpsB* or Δ*stkP* suppressor strains of *S. pneumoniae* D39

Previously, we reported five spontaneous missense mutations in *phpP* (Thr/Ser protein phosphatase) (Table 1, lines 2 and 5-8) and two mutants containing large chromosomal duplications/deletions (Table 1, lines 3-4) that suppress the essentiality of Δ*gpsB* in unencapsulated *S. pneumoniae* D39 (Rued *et al*., 2017). However, we did not determine the basis for Δ*gpsB* suppression or how the duplications/deletions formed in these mutants. To this end, we screened 20 additional Δ*gpsB* spontaneous suppressors from independent transformations by sequencing for *phpP* mutations or by PCR for the Δ(*spd_1029’-spd_1037’*)-region deletion present in the *sup gpsB-2* and sup *gpsB-3* duplication/deletion mutants (Rued *et al*., 2017). Fifteen of 20 suppressors contained Δ(*spd_1032’-spd_1036’*)-region deletions, indicative of adjacent duplications (Table 1, line 13). Whole-genome sequencing of the remaining 5 suppressors indicated that *sup gpsB-8* contains an ≈163 kb (149 genes) duplication of Ω[*spd_0889’-spd_1037’*] (Fig. 2A, S1B, and S2B; Table 1, line 9), *sup gpsB-9* and *sup gpsB-10* contain an ≈18 kb (21 genes) duplication or quadruplication, respectively, of Ω[*spd_0966’* to *spd_0986’*] (Fig. 2A and S1C; Table 1, lines 10-11), *sup gpsB-11* contains a *murZ*(D280Y) missense mutation as well as two other mutations (Table 1, line 12 and footnote), and *sup gpsB-27* contains a nonsense mutation *ireB*(Q84(STOP)), truncating IreB by 4 amino acids, as well as a (7→6) slippage mutation in an intergenic region (Table 1, line 14 and footnote). Genetic separation showed that *murZ*(D280Y) or *ireB*(Q84(STOP)) was necessary and sufficient for Δ*gpsB* suppression (Table 2, lines 6 and 12). Consistent with involvement of MurZ in Δ*gpsB* suppression, the duplicated regions of *sup gpsB*-2-3 and *sup gpsB*-*8*-*10* contain *murZ (spd_0967*) (Fig. 2A, 3, and S1B-S1C).

**Figure 2.**
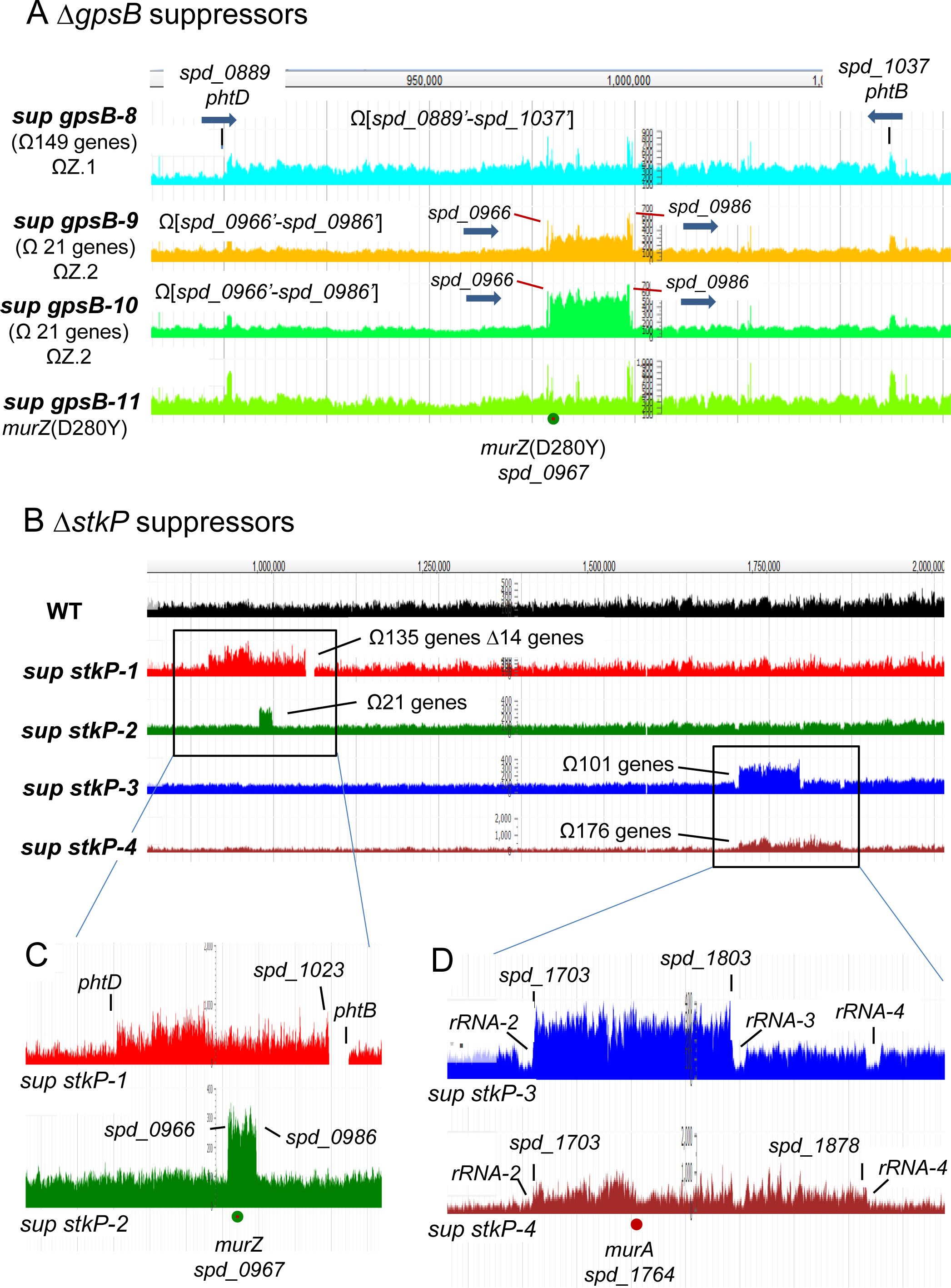
Chromosomal duplications containing *murZ* or *murA* are present in Δ*gpsB* or Δ*stkP* suppressor strains of *S. pneumoniae* D39. (A) Snapshot of genome browser output of Δ*gpsB sup* strains from genome coordinates 870 to 1100 kb. Three new Δ*gpsB* suppressor strains contain chromosomal duplication or quadruplication of multiple genes, all of which include *murZ. Sup gpsB-8* contains a ≈163 kb duplication of chromosomal region from *spd_0889’* to *spd_1037’,* while *sup gpsB-9* and *sup gpsB-10* contain a duplication or quadruplication, respectively, of the chromosomal region from *spd_0966’* to *spd_0986’*. *sup gpsB-10* has a *murZ*(D280Y) mutation and no chromosomal duplication. Black lines point to the flanking regions of the duplication found in *sup gpsB-8*, which are 1324-bp inverted repeats present in *phtD* (*spd_0889*) and *phtB* (*spd_1037*), encoding 2 of the 3 pneumococcal histidine triad proteins. The red lines point to the flanking regions (*spd_0966* and *spd_0986*) of duplication or quadruplication found in *sup gpsB-9,* and *-10*, respectively. *spd_0966* and *spd_0986* are pseudogenes containing IS1167 degenerate transposase sequences. Thick blue arrows show the gene orientations of *phtB*, *phtD*, *spd_0966*, and *spd_0986*. (B) Snapshot of genome browser output of Δ*stkP sup* strains from genome coordinates 750 to 2,000 kb. (C) *Sup stkP-1* contains a duplication/deletion between *phtD* and *phtB,* and *sup stkP-2* contains a duplication between *spd_0966* and *spd_0986.* (D) Large duplications found in *sup stkP-3* and *-4* are flanked by tRNA + rRNA clusters rRNA/rRNA3 and rRNA/rRNA4 respectively. *Sup stkP-3* showed a decrease in sequence reads of the four *rRNA-1-4* operons (*rRNA-1, rRNA-2, rRNA-3*, *and rRNA-4*) compared to the surrounding region. It is possible that either *rRNA-2* or *rRNA-3,* or both *rRNA-2* and *rRNA-3,* are deleted in this strain, but because of the sequence identity of the *rRNA* operons, deletion of one or two operons manifest as a decrease of reads for all four operons.

**Figure 3.**
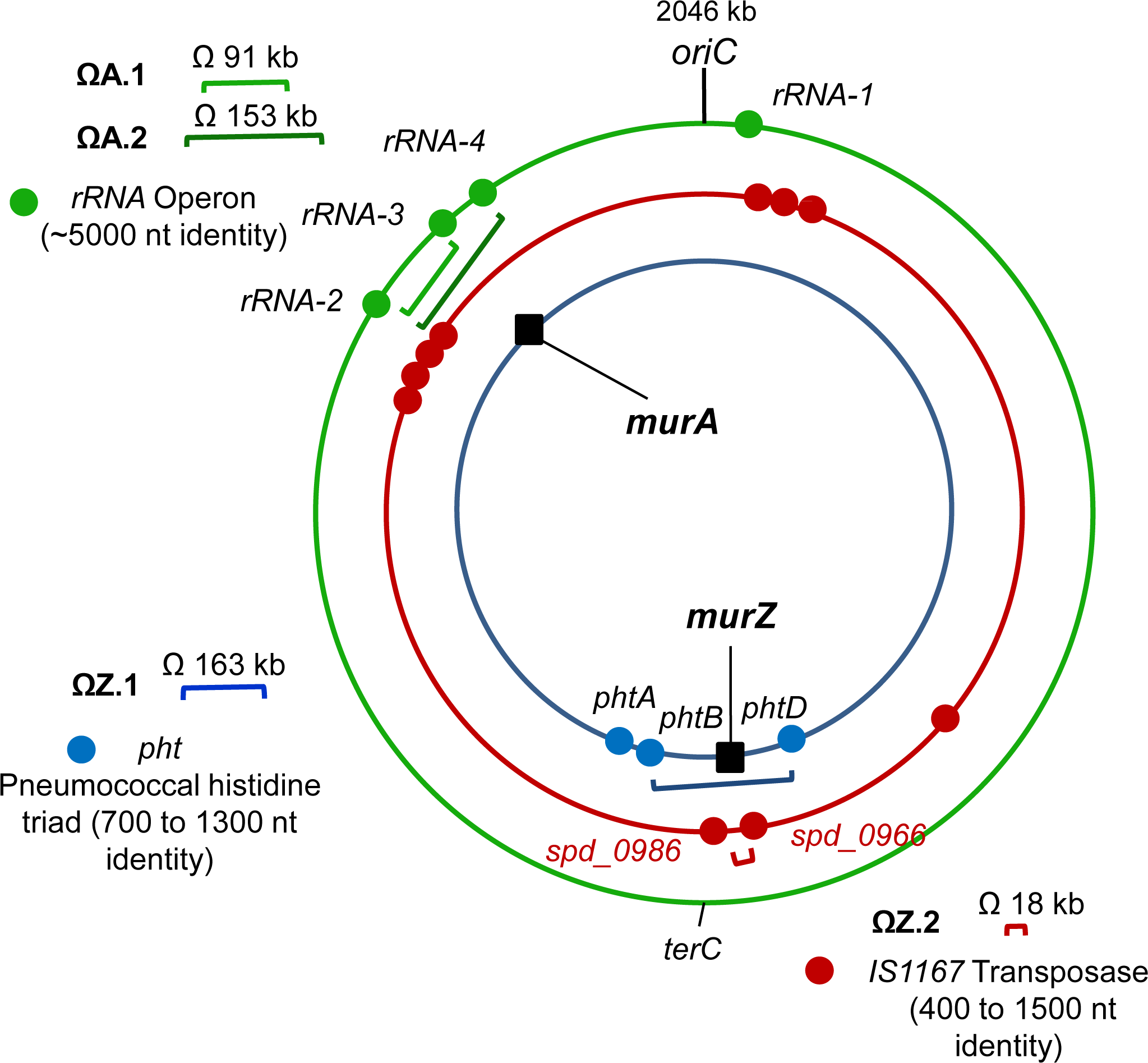
Locations of repeated sequences that anchor chromosomal duplications in *S. pneumoniae* D39. Blue, red, and green dots are locations of *pht* genes, IS1167 transposase, and tRNA/rRNA gene clusters, respectively. Duplications ΩZ.1 and ΩZ.2 result in duplication of *murZ* and surrounding genes, while ΩA.1 and ΩA.2 result in *murA* duplication. ΩZ.1 is present in *sup gpsB*-8. ΩZ.2 is present in *sup gpsB*-9, *sup gpsB*-10, and *sup stkP*-2. ΩA.1 is present in *sup stkP*-3 while ΩA.2 is present in *sup stkP*-4.

**Table 1.**
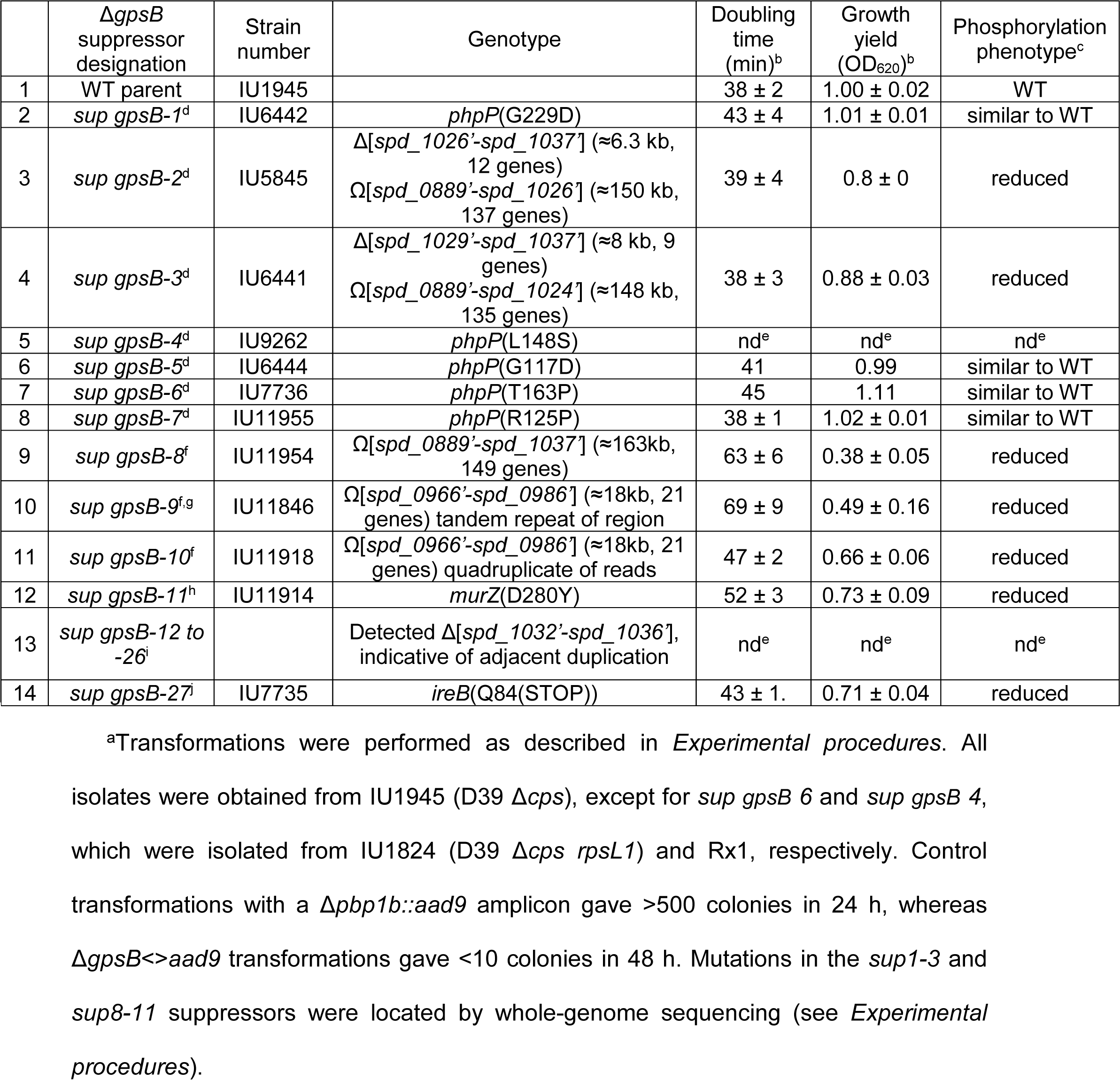
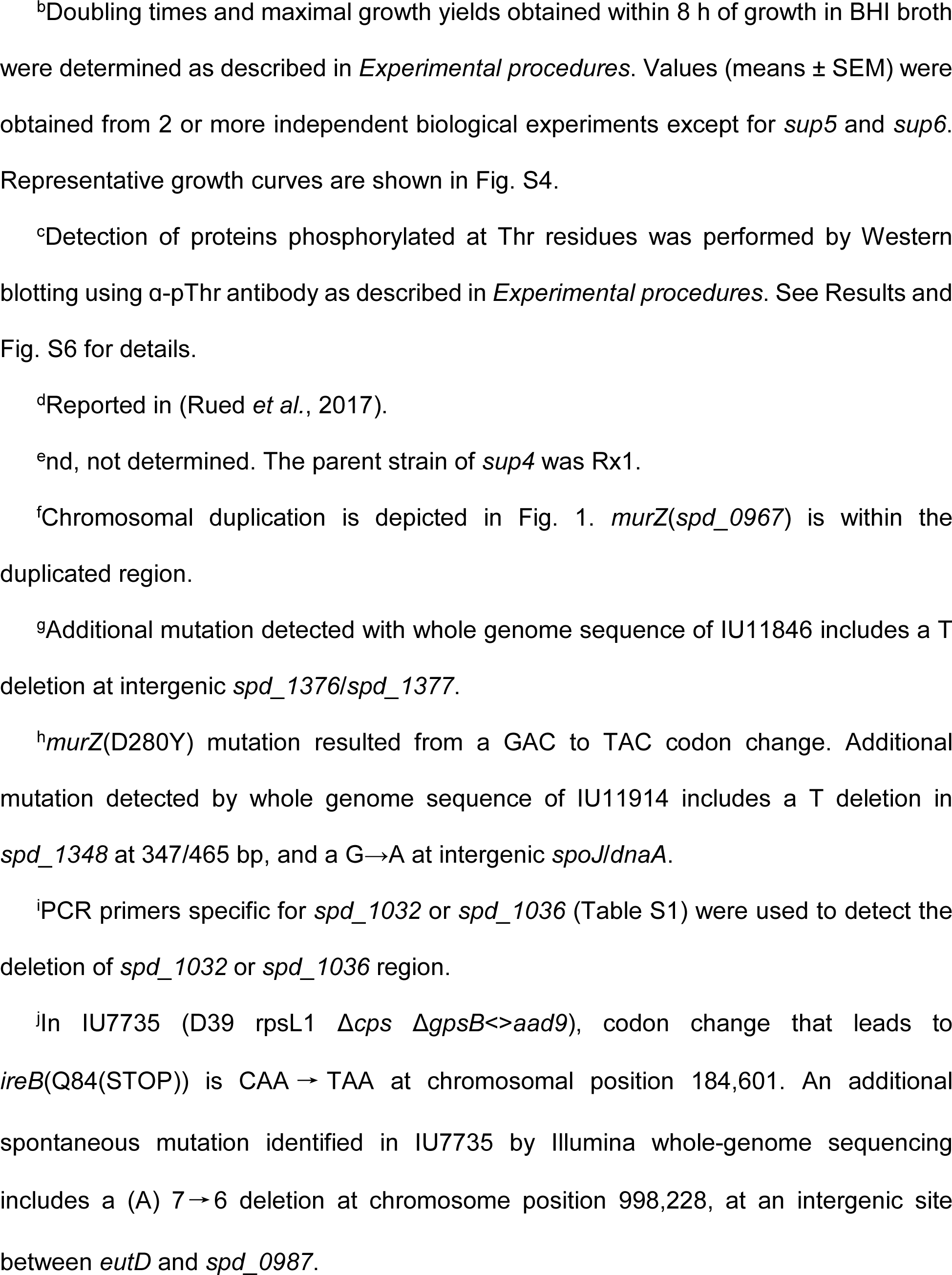
Analysis of spontaneous Δ*gpsB* suppressor mutations that arose in unencapsulated *S. pneumoniae* Δ*cps* D39^a^.

|  | *ΔgpsB* suppressor designation | Strain number | Genotype | Doubling time (min)<sup>b</sup> | Growth yield (OD<sub>620</sub>)<sup>b</sup> | Phosphorylation phenotype<sup>c</sup> |
| --- | --- | --- | --- | --- | --- | --- |
| 1 | WT parent | IU1945 |  | 38 ± 2 | 1.00 ± 0.02 | WT |
| 2 | *sup gpsB-1*<sup>d</sup> | IU6442 | *phpP*(G229D) | 43 ± 4 | 1.01 ± 0.01 | similar to WT |
| 3 | *sup gpsB-2*<sup>d</sup> | IU5845 | Δ[*spd_1026'-spd_1037'*] (≈6.3 kb, 12 genes)<br>Ω[*spd_0889'-spd_1026'*] (≈150 kb, 137 genes) | 39 ± 4 | 0.8 ± 0 | reduced |
| 4 | *sup gpsB-3*<sup>d</sup> | IU6441 | Δ[*spd_1029'-spd_1037'*] (≈8 kb, 9 genes)<br>Ω[*spd_0889'-spd_1024'*] (≈148 kb, 135 genes) | 38 ± 3 | 0.88 ± 0.03 | reduced |
| 5 | *sup gpsB-4*<sup>d</sup> | IU9262 | *phpP*(L148S) | nd<sup>e</sup> | nd<sup>e</sup> | nd<sup>e</sup> |
| 6 | *sup gpsB-5*<sup>d</sup> | IU6444 | *phpP*(G117D) | 41 | 0.99 | similar to WT |
| 7 | *sup gpsB-6*<sup>d</sup> | IU7736 | *phpP*(T163P) | 45 | 1.11 | similar to WT |
| 8 | *sup gpsB-7*<sup>d</sup> | IU11955 | *phpP*(R125P) | 38 ± 1 | 1.02 ± 0.01 | similar to WT |
| 9 | *sup gpsB-8*<sup>f</sup> | IU11954 | Ω[*spd_0889'-spd_1037'*] (≈163kb, 149 genes) | 63 ± 6 | 0.38 ± 0.05 | reduced |
| 10 | *sup gpsB-9*<sup>f,g</sup> | IU11846 | Ω[*spd_0966'-spd_0986'*] (≈18kb, 21 genes) tandem repeat of region | 69 ± 9 | 0.49 ± 0.16 | reduced |
| 11 | *sup gpsB-10*<sup>f</sup> | IU11918 | Ω[*spd_0966'-spd_0986'*] (≈18kb, 21 genes) quadruplicate of reads | 47 ± 2 | 0.66 ± 0.06 | reduced |
| 12 | *sup gpsB-11*<sup>h</sup> | IU11914 | *murZ*(D280Y) | 52 ± 3 | 0.73 ± 0.09 | reduced |
| 13 | *sup gpsB-12 to -26*<sup>i</sup> |  | Detected Δ[*spd_1032'-spd_1036'*], indicative of adjacent duplication | nd<sup>e</sup> | nd<sup>e</sup> | nd<sup>e</sup> |
| 14 | *sup gpsB-27*<sup>j</sup> | IU7735 | *ireB*(Q84(STOP)) | 43 ± 1. | 0.71 ± 0.04 | reduced |

<sup>a</sup>Transformations were performed as described in *Experimental procedures*. All isolates were obtained from IU1945 (D39 *Δcps*), except for *sup gpsB* 6 and *sup gpsB* 4, which were isolated from IU1824 (D39 *Δcps rpsL1*) and Rx1, respectively. Control transformations with a *Δpbp1b::aad9* amplicon gave >500 colonies in 24 h, whereas *ΔgpsB<>aad9* transformations gave <10 colonies in 48 h. Mutations in the *sup1-3* and *sup8-11* suppressors were located by whole-genome sequencing (see *Experimental procedures*).
<sup>b</sup>Doubling times and maximal growth yields obtained within 8 h of growth in BHI broth were determined as described in *Experimental procedures*. Values (means ± SEM) were obtained from 2 or more independent biological experiments except for *sup5* and *sup6*. Representative growth curves are shown in Fig. S4.
<sup>c</sup>Detection of proteins phosphorylated at Thr residues was performed by Western blotting using α-pThr antibody as described in *Experimental procedures*. See Results and Fig. S6 for details.
<sup>d</sup>Reported in (Rued *et al.*, 2017).
<sup>e</sup>nd, not determined. The parent strain of *sup4* was Rx1.
<sup>f</sup>Chromosomal duplication is depicted in Fig. 1. *murZ*(*spd\_0967*) is within the duplicated region.
<sup>g</sup>Additional mutation detected with whole genome sequence of IU11846 includes a T deletion at intergenic *spd\_1376/spd\_1377*.
<sup>h</sup>*murZ*(D280Y) mutation resulted from a GAC to TAC codon change. Additional mutation detected by whole genome sequence of IU11914 includes a T deletion in *spd\_1348* at 347/465 bp, and a G → A at intergenic *spoJ/dnaA*.
<sup>i</sup>PCR primers specific for *spd\_1032* or *spd\_1036* (Table S1) were used to detect the deletion of *spd\_1032* or *spd\_1036* region.
<sup>j</sup>In IU7735 (D39 *rpsL1* $\Delta$ *cps* $\Delta$ *gpsB* ↔ *aad9*), codon change that leads to *ireB*(Q84(STOP)) is CAA → TAA at chromosomal position 184,601. An additional spontaneous mutation identified in IU7735 by Illumina whole-genome sequencing includes a (A) 7 → 6 deletion at chromosome position 998,228, at an intergenic site between *eutD* and *spd\_0987*.

**Table 2.**
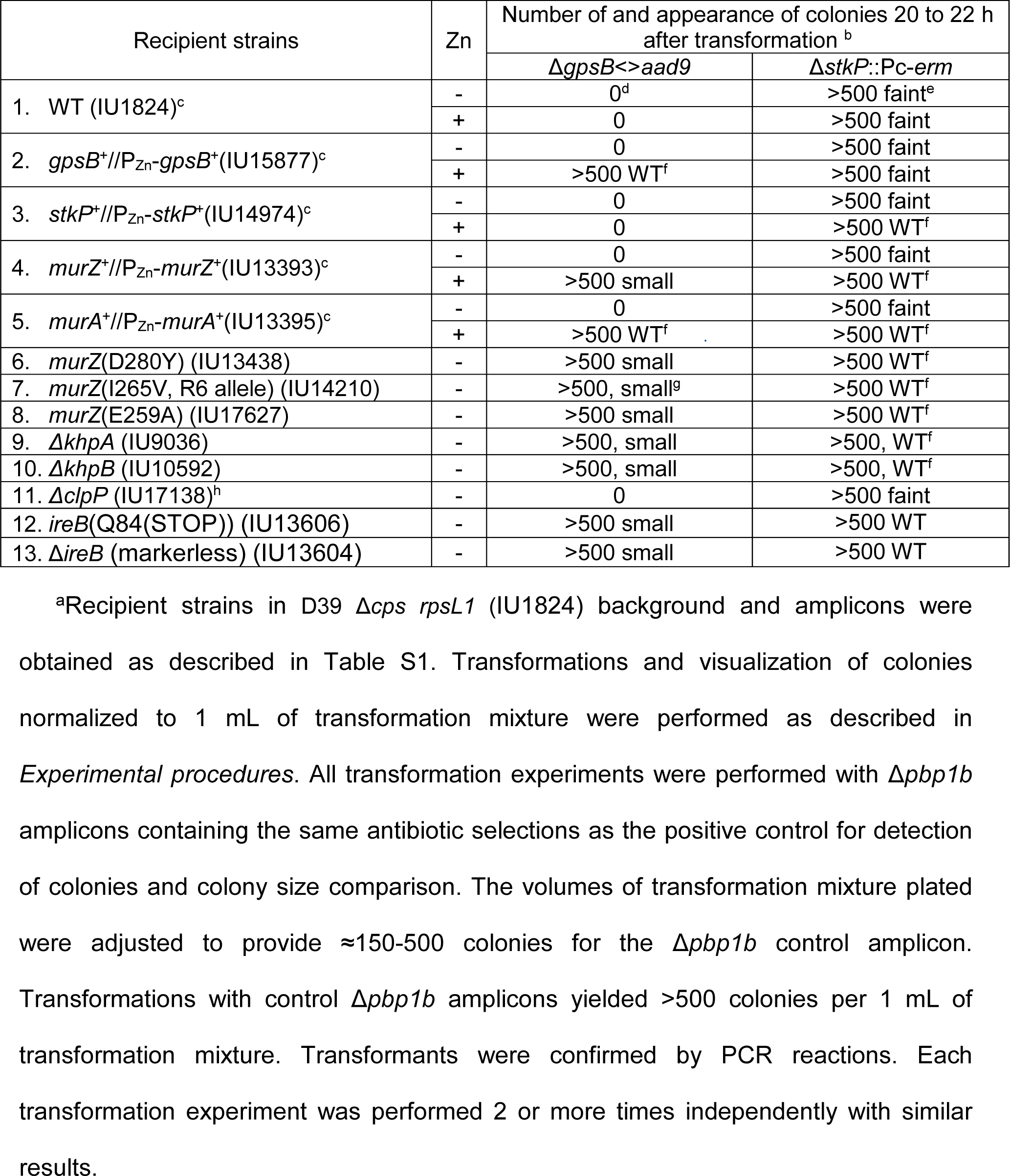
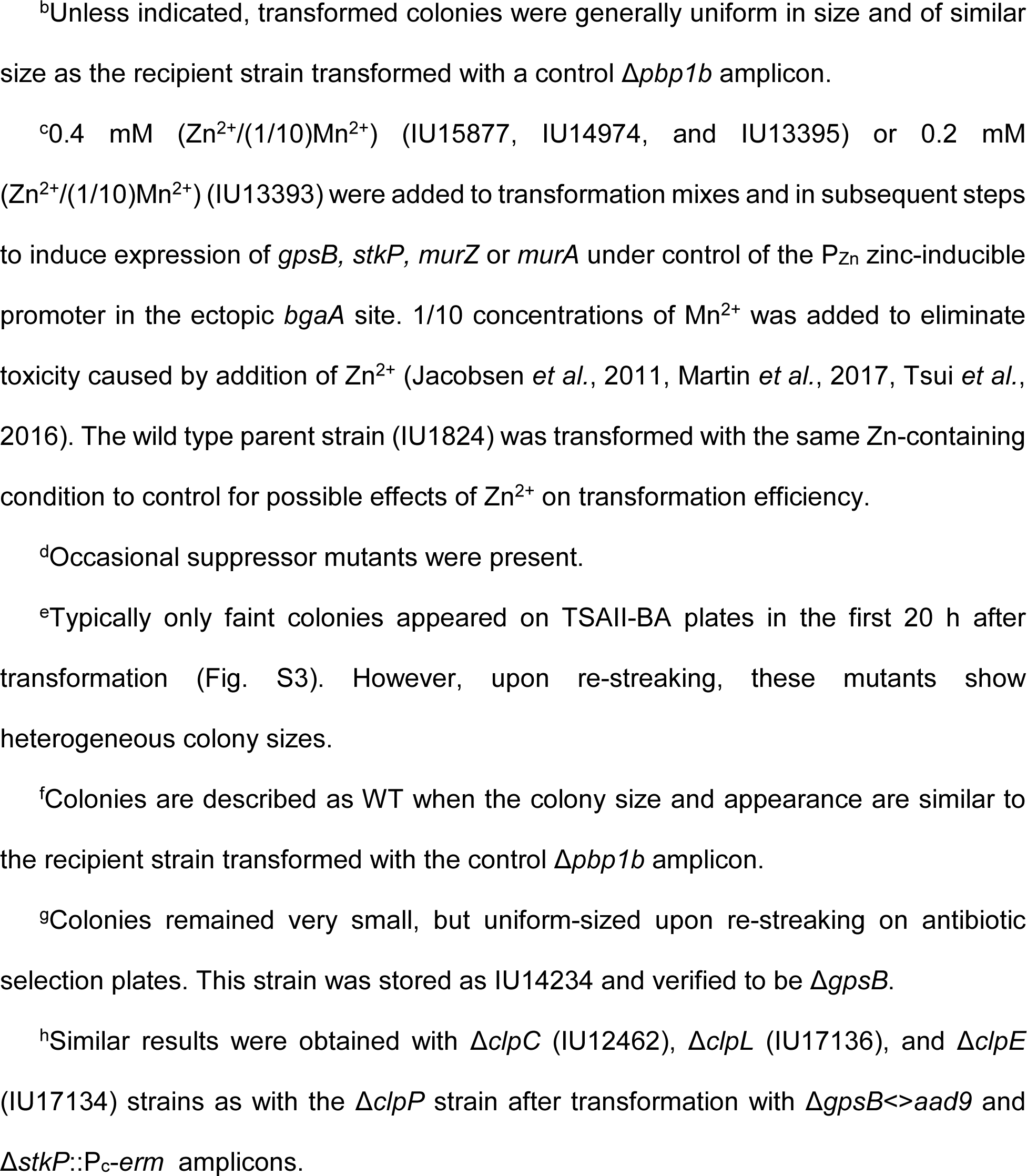
Suppression of Δ*gpsB* or Δ*stkP* mutation in *S. pneumoniae* Δ*cps* D39^a^.

| Recipient strains | Zn | Number of and appearance of colonies 20 to 22 h after transformation <sup>b</sup> |  |
| --- | --- | --- | --- |
| | | $\Delta gpsB \leftrightarrow aad9$ | $\Delta stkP::Pc-erm$ |
| 1. WT (IU1824)<sup>c</sup> | - | 0<sup>d</sup> | >500 faint<sup>e</sup> |
|  | + | 0 | >500 faint |
| 2. *gpsB*<sup>+</sup>//P<sub>Zn</sub>-*gpsB*<sup>+</sup>(IU15877)<sup>c</sup> | - | 0 | >500 faint |
|  | + | >500 WT<sup>f</sup> | >500 faint |
| 3. *stkP*<sup>+</sup>//P<sub>Zn</sub>-*stkP*<sup>+</sup>(IU14974)<sup>c</sup> | - | 0 | >500 faint |
|  | + | 0 | >500 WT<sup>f</sup> |
| 4. *murZ*<sup>+</sup>//P<sub>Zn</sub>-*murZ*<sup>+</sup>(IU13393)<sup>c</sup> | - | 0 | >500 faint |
|  | + | >500 small | >500 WT<sup>f</sup> |
| 5. *murA*<sup>+</sup>//P<sub>Zn</sub>-*murA*<sup>+</sup>(IU13395)<sup>c</sup> | - | 0 | >500 faint |
|  | + | >500 WT<sup>f</sup> | >500 WT<sup>f</sup> |
| 6. *murZ*(D280Y) (IU13438) | - | >500 small | >500 WT<sup>f</sup> |
| 7. *murZ*(I265V, R6 allele) (IU14210) | - | >500, small<sup>g</sup> | >500 WT<sup>f</sup> |
| 8. *murZ*(E259A) (IU17627) | - | >500 small | >500 WT<sup>f</sup> |
| 9. $\Delta khpA$ (IU9036) | - | >500, small | >500, WT<sup>f</sup> |
| 10. $\Delta khpB$ (IU10592) | - | >500, small | >500, WT<sup>f</sup> |
| 11. $\Delta clpP$ (IU17138)<sup>h</sup> | - | 0 | >500 faint |
| 12. *ireB*(Q84(STOP)) (IU13606) | - | >500 small | >500 WT |
| 13. $\Delta ireB$ (markerless) (IU13604) | - | >500 small | >500 WT |

<sup>a</sup>Recipient strains in D39 $\Delta cps$ *rpsL1* (IU1824) background and amplicons were obtained as described in Table S1. Transformations and visualization of colonies normalized to 1 mL of transformation mixture were performed as described in *Experimental procedures*. All transformation experiments were performed with $\Delta pbp1b$ amplicons containing the same antibiotic selections as the positive control for detection of colonies and colony size comparison. The volumes of transformation mixture plated were adjusted to provide $\approx 150$ -500 colonies for the $\Delta pbp1b$ control amplicon. Transformations with control $\Delta pbp1b$ amplicons yielded >500 colonies per 1 mL of transformation mixture. Transformants were confirmed by PCR reactions. Each transformation experiment was performed 2 or more times independently with similar results.
<sup>b</sup>Unless indicated, transformed colonies were generally uniform in size and of similar size as the recipient strain transformed with a control $\Delta pbp1b$ amplicon.
<sup>c</sup>0.4 mM ( $Zn^{2+}/(1/10)Mn^{2+}$ ) (IU15877, IU14974, and IU13395) or 0.2 mM ( $Zn^{2+}/(1/10)Mn^{2+}$ ) (IU13393) were added to transformation mixes and in subsequent steps to induce expression of *gpsB*, *stkP*, *murZ* or *murA* under control of the $P_{Zn}$ zinc-inducible promoter in the ectopic *bgaA* site. 1/10 concentrations of $Mn^{2+}$ was added to eliminate toxicity caused by addition of $Zn^{2+}$ (Jacobsen *et al.*, 2011, Martin *et al.*, 2017, Tsui *et al.*, 2016). The wild type parent strain (IU1824) was transformed with the same Zn-containing condition to control for possible effects of $Zn^{2+}$ on transformation efficiency.
<sup>d</sup>Occasional suppressor mutants were present.
<sup>e</sup>Typically only faint colonies appeared on TSAII-BA plates in the first 20 h after transformation (Fig. S3). However, upon re-streaking, these mutants show heterogeneous colony sizes.
<sup>f</sup>Colonies are described as WT when the colony size and appearance are similar to the recipient strain transformed with the control $\Delta pbp1b$ amplicon.
<sup>g</sup>Colonies remained very small, but uniform-sized upon re-streaking on antibiotic selection plates. This strain was stored as IU14234 and verified to be $\Delta gpsB$ .
<sup>h</sup>Similar results were obtained with $\Delta clpC$ (IU12462), $\Delta clpL$ (IU17136), and $\Delta clpE$ (IU17134) strains as with the $\Delta clpP$ strain after transformation with $\Delta gpsB \rightarrow aad9$ and $\Delta stkP::P_c-erm$ amplicons.

Since GpsB plays a role in activation of pneumococcal StkP Ser/Thr kinase activity (Fleurie *et al*., 2014, Rued *et al*., 2017), we also isolated and characterized suppressor mutations of D39 unencapsulated (Δ*cps*) and encapsulated (*cps*)^+^ strains transformed with a Δ*stkP* or Δ[*phpP-stkP*] amplicon (Tables 2 and 3). Transformants typically appeared as faint, indistinct colonies on TSAII-BA plates after 20 h (Fig. S3A; Table 2). Re-streaking these Δ*stkP* and Δ[*phpP-stkP*] transformants resulted in heterogeneously sized, faster growing colonies, indicative of suppressor accumulation (Rued *et al*., 2017). We interrogated six of these re-streaked transformants for the presence of suppressor mutations (Table 3). Gene sequencing showed that none contained mutations in *murZ*, but one Δ[*phpP*-*stkP*] suppressor contained a 14-bp duplication within the ribosome-binding site (RBS) of *ireB*(*Spn*) (Table 3, line 9). This RBS-mutation will be described further elsewhere. Only one (1/6) of the transformants contained a Δ(*spd_1032’-spd_1036’*)-region deletion (Table 3, line 8), indicative of an adjacent *phtD-phtB* duplication (Fig. 2B). The genomes of the four remaining Δ*stkP* or Δ[*phpP*-*stkP*] transformants were sequenced (Table 3), and all were found to contain chromosomal duplications containing *murZ* or *murA*. (Fig. 2B and S1B-S1D). *sup stkP-1* has a duplication containing *murZ* and unexpectedly, a deletion similar to that of *sup gpsB-2*, except for the deletion junction (Fig. S1B). The deletion in *sup stkP-1* accumulated during propagation of the initial Δ*stkP* isolate, which lacks the deletion accordingly to PCR assays. A similar duplication/deletion was reported previously in a D39 Δ*stkP* mutant (Ulrych *et al*., 2021). *sup stkP-2* contains a 21-gene duplication containing *murZ*, similar to that of *sup gpsB-9* (Fig. S1C). Notably, *sup stkP-3* and *sup* stkP-4 contain duplications of Ω[*spd_1703’-spd_1803’*] and Ω[*spd_1703’-spd_1878*’], respectively, which contain *murA* (*spd_1764*) (Fig. 2B and S1D). Together, these results implicate overexpression of MurZ or MurA in Δ*gpsB* and Δ*stkP* suppression.

**Table 3.**
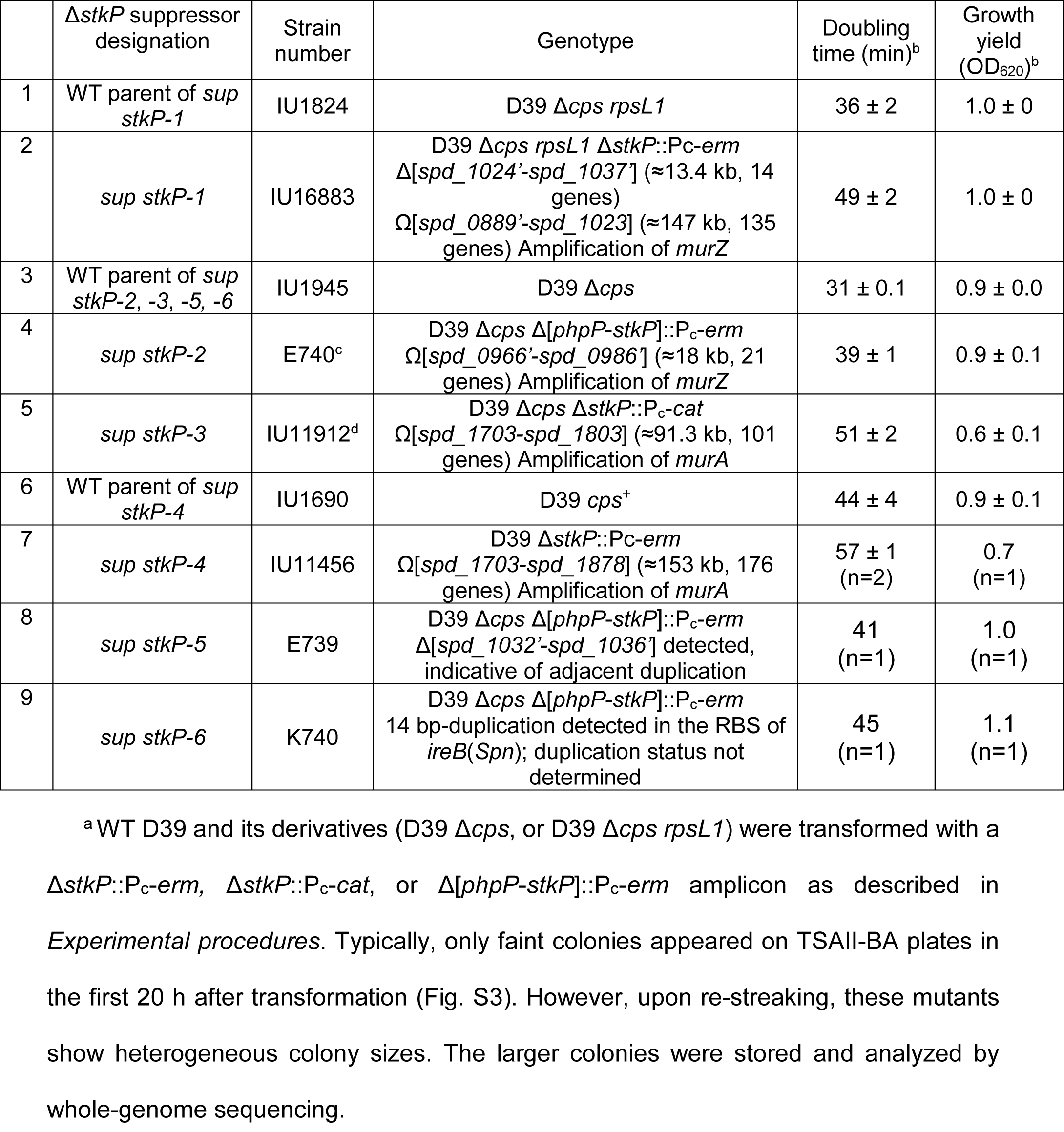
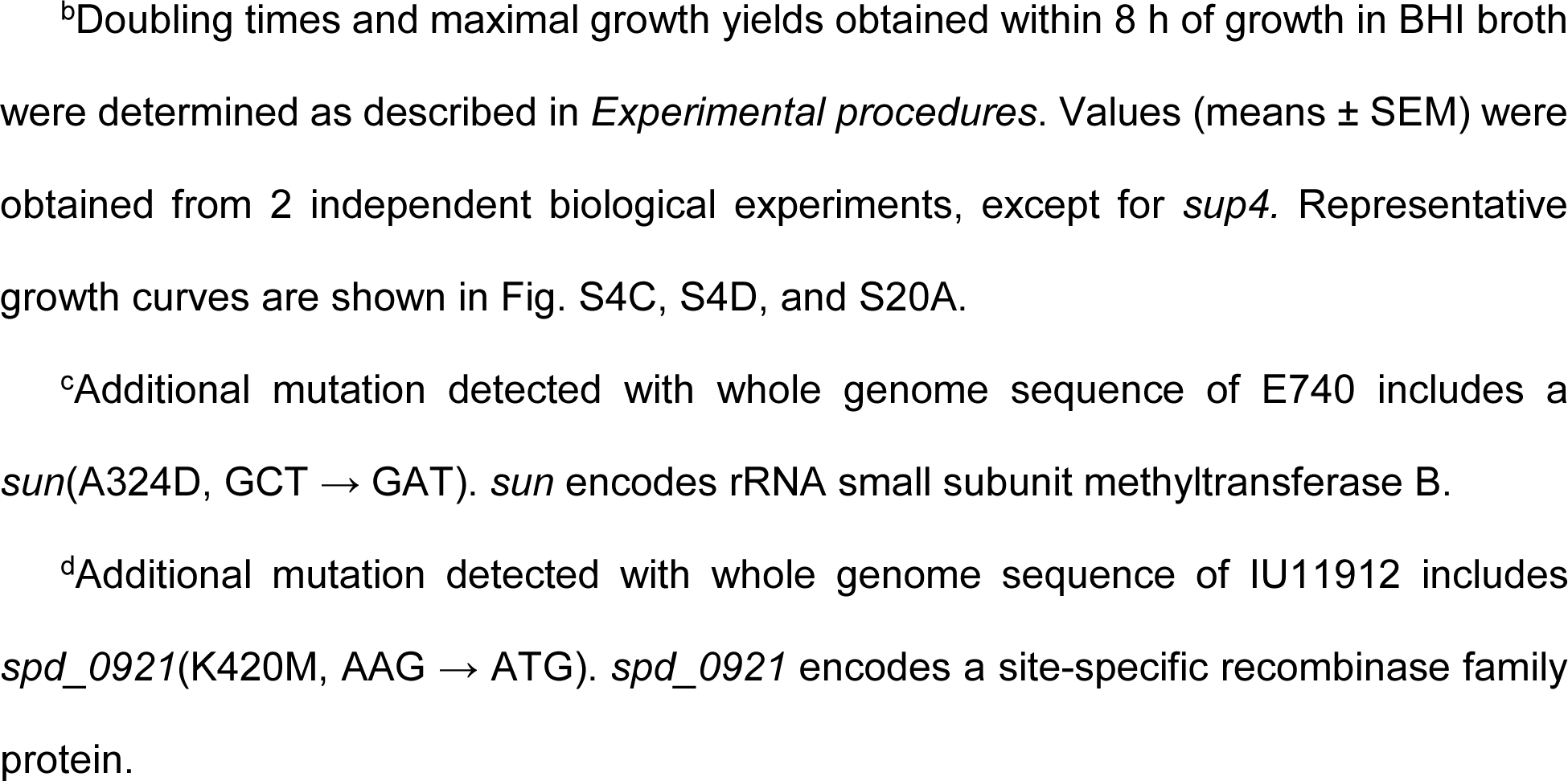
Analysis of spontaneous Δ*stkP* suppressor mutations that arose in unencapsulated (Δ*cps*) and encapsulated D39 *S. pneumoniae* D39^a^

| | $\Delta stkP$ suppressor designation | Strain number | Genotype | Doubling time (min)<sup>b</sup> | Growth yield (OD<sub>620</sub>)<sup>b</sup> |
| --- | --- | --- | --- | --- | --- |
| 1 | WT parent of *sup stkP-1* | IU1824 | D39 $\Delta cps$ *rpsL1* | 36 ± 2 | 1.0 ± 0 |
| 2 | *sup stkP-1* | IU16883 | D39 $\Delta cps$ *rpsL1* $\Delta stkP::P_c-erm$ $\Delta[spd\_1024'-spd\_1037']$ (≈13.4 kb, 14 genes) $\Omega[spd\_0889'-spd\_1023]$ (≈147 kb, 135 genes) Amplification of *murZ* | 49 ± 2 | 1.0 ± 0 |
| 3 | WT parent of *sup stkP-2, -3, -5, -6* | IU1945 | D39 $\Delta cps$ | 31 ± 0.1 | 0.9 ± 0.0 |
| 4 | *sup stkP-2* | E740<sup>c</sup> | D39 $\Delta cps$ $\Delta[phpP-stkP]::P_c-erm$ $\Omega[spd\_0966'-spd\_0986']$ (≈18 kb, 21 genes) Amplification of *murZ* | 39 ± 1 | 0.9 ± 0.1 |
| 5 | *sup stkP-3* | IU11912<sup>d</sup> | D39 $\Delta cps$ $\Delta stkP::P_c-cat$ $\Omega[spd\_1703-spd\_1803]$ (≈91.3 kb, 101 genes) Amplification of *murA* | 51 ± 2 | 0.6 ± 0.1 |
| 6 | WT parent of *sup stkP-4* | IU1690 | D39 *cps*<sup>+</sup> | 44 ± 4 | 0.9 ± 0.1 |
| 7 | *sup stkP-4* | IU11456 | D39 $\Delta stkP::P_c-erm$ $\Omega[spd\_1703-spd\_1878]$ (≈153 kb, 176 genes) Amplification of *murA* | 57 ± 1 (n=2) | 0.7 (n=1) |
| 8 | *sup stkP-5* | E739 | D39 $\Delta cps$ $\Delta[phpP-stkP]::P_c-erm$ $\Delta[spd\_1032'-spd\_1036']$ detected, indicative of adjacent duplication | 41 (n=1) | 1.0 (n=1) |
| 9 | *sup stkP-6* | K740 | D39 $\Delta cps$ $\Delta[phpP-stkP]::P_c-erm$ 14 bp-duplication detected in the RBS of *ireB*(*Spn*); duplication status not determined | 45 (n=1) | 1.1 (n=1) |

<sup>a</sup> WT D39 and its derivatives (D39 $\Delta cps$ , or D39 $\Delta cps$ *rpsL1*) were transformed with a $\Delta stkP::P_c-erm$ , $\Delta stkP::P_c-cat$ , or $\Delta[phpP-stkP]::P_c-erm$ amplicon as described in *Experimental procedures*. Typically, only faint colonies appeared on TSAII-BA plates in the first 20 h after transformation (Fig. S3). However, upon re-streaking, these mutants show heterogeneous colony sizes. The larger colonies were stored and analyzed by whole-genome sequencing.
<sup>b</sup>Doubling times and maximal growth yields obtained within 8 h of growth in BHI broth were determined as described in *Experimental procedures*. Values (means ± SEM) were obtained from 2 independent biological experiments, except for *sup4*. Representative growth curves are shown in Fig. S4C, S4D, and S20A.
<sup>c</sup>Additional mutation detected with whole genome sequence of E740 includes a *sun*(A324D, GCT → GAT). *sun* encodes rRNA small subunit methyltransferase B.
<sup>d</sup>Additional mutation detected with whole genome sequence of IU11912 includes *spd\_0921*(K420M, AAG → ATG). *spd\_0921* encodes a site-specific recombinase family protein.

### 2.2 **Repeats in *phtD* and *phtB,* degenerate IS elements *spd_0966* and *spd_0986,*** or tRNA/rRNA gene clusters contribute to pneumococcal genomic plasticity

To understand their formation, we deduced the flanking sequences of the duplications that suppress Δ*gpsB* and Δ*stkP* (Fig. 2). The duplications were grouped into four patterns: ΩZ.1 or ΩZ.2 for duplication of the *murZ* region and ΩA.1 or ΩA.2 for duplication of the *murA* region (Fig. 3). The flanking sequences of ΩZ.1 duplications are intact and hybrid inverted repeat elements of *phtD* and *phtB*, while ΩZ.2 duplications are bordered by intact direct repeats of degenerate IS elements *spd_0966* and *spd_0986.* The flanking sequences of ΩA.1 or ΩA.2 duplications consist of intact direct repeats of tRNA/rRNA gene clusters (Fig. 2, 3, and S1).

ΩZ.1 duplications (Fig. 3 and S1B; *sup gpsB-2, −3,* and *-8,* and *sup stkP-1*) are bordered by intact or hybrid (*phtB’*/*phtD*’) inverted repeat elements of *phtD* (*spd_0889*) and *phtB* (*spd_1037*) generated by homologous recombination (Fig. 2A, S1B, and S2B; where apostrophes indicate hybrid genes). *phtD* and *phtB* encode 2 of the 3 histidine triad proteins in *S. pneumoniae* D39 and have identical 1,324-bp sequences at their 3’-ends (Table S2). During chromosome replication when there are two copies of the genes between *phtD* and *phtB*, the large *phtD* and *phtB* inverted repeats can recombine to invert the order of intervening genes. Evidence for inversion during duplication formation is presented below for *sup gpsB-3* (Fig. S2C-F).

However, the inverted *phtD* and *phtB* sequences cannot foster direct homologous recombination to form a duplication. Consequently, *phtD* and *phtB* must also contain short direct repeats or other elements that enhance short-junction (SJ) duplication (Reams & Roth, 2015) that keeps the duplication boundaries within *phtD* and *phtB* (Fig. 2A, 2B, S1B, and S2B). Indeed, there are small direct repeats of 8 and 9 bp and shorter clusters of directly repeated bps within inverted *phtD* and *phtB* that could promote SJ duplication. Of the 4 ΩZ.1 duplications, only *sup gpsB-*8 contains intact duplicated regions, which may be aligned in the same or an inverted orientation. The other three ΩZ.1 duplications contain slightly different deletions of duplication junctions (labeled ΩZ.1Δ; Fig. S1B and S2C). Similar remodeling by deletion of duplication junctions often occurs (Reams & Roth, 2015). Interestingly, all ΩZ.1 duplications create a second copy of the *terC* chromosomal replication terminus, including the *dif_SL_* recombination site and *xerC* recombinase gene (star, Fig. S1A), that mediate chromosome dimer resolution in *Streptococci/Lactococci* (Le Bourgeois *et al*., 2007). In ΩZ.1 duplications, the two copies of *dif_SL_* and *xerS* are oppositely oriented (Fig. S1B).

ΩZ.2 duplications are bordered by direct repeats of pseudogenes *spd_0966* and *spd_0986*, which contain IS1167 degenerate transposase sequences (Fig. 2B, 3, and S1C; *sup gpsB-9* and −*10*, and *sup stkP-2*)*. spd_0966* (1,492 bp) shows 91% identity with *spd_0986* (1,477 bp), including 240-bp of identical sequence at their ends (Table S3). The duplications are likely joined by a *spd_0986*’/*spd_0966*’ hybrid element formed by homologous recombination (Fig. S1C). Similarly, ΩA.1 (*sup stkP-3*) and ΩA.2 (*sup stkP*-*4*) duplications are bordered by direct repeats; in this case, of *rRNA* operons that have homologous/heterologous DNA stretching over >5,000 bp (Fig. 2B, 2D, 3, and S1D; Table S4). *sup stkP*-*3* is flanked by direct repeats of the ≈6 kb *rRNA-2* and *rRNA-3* operons, which are 99.9% identical and contain genes for 9 tRNAs, a 5S rRNA, a 23 S rRNA, a tRNA, a 16S rRNA, and a tRNA (Table S4). The internal junction in *sup stkP-3* is likely a *rRNA-3’*/*rRNA*-*2*’ hybrid element (S1D). *sup stkP-4* is flanked by direct repeats of *rRNA-2* and *rRNA-4*, with a hybrid *rRNA-4’*/*rRNA-2’* element in the internal junction (S1D). The ≈5.2 kb *rRNA*-*4* operon contains the same (100% identity) tRNA, 5S RNA, 23S RNA, tRNA, 16S rRNA, and tRNA genes as the distal portion of *rRNA-2* (Table S4). Together these results show that repeats of *phtD* and *phtB*, degenerate IS transposase genes, and tRNA/rRNA gene clusters act as endpoints for duplications of regions ranging from ≈18 kb (21 genes) to >150 kb (176 genes) in the *S. pneumoniae* D39 chromosome.

### 2.3 Deletions in ΩZ.1Δ duplications may enhance fitness of Δ*gpsB* mutants

To provide a model for formation ΩZ.1Δ duplication/deletions (Fig. 2B and S1B), we assumed that the first event was formation of an intact ΩZ.1 inverted duplication between *spd_0889*’ (*phtD’*) and *spd_1037*’ (*phtB’*), such as *sup gpsB-8* (Fig. 2A, S1B, and S2B). The next event would be deletion from *spd_1029*’ on one side of the duplication junction to *spd*_*1024’* on the other side (Fig. S2C). Notably, the endpoints of internal deletions of the duplication junction are slightly different for *sup gpsB-3*, *sup stkP-1,* and *sup gpsB-2* (Fig. S1B and S2C). We obtained results consistent with this model by PCR analysis of *sup gpsB-3* compared to WT (Fig. S2C-S2F). Primer pairs P1/P3, P1/P4, P2/P3, and P2/P4 yielded PCR products of the expected sizes for the arrangement shown for *sup gpsB-3*, but not WT, consistent with formation of an inverted duplication followed by deletion of the rearrangement junction (Fig. S2C).

Different deletions of the *spd_1032’* to *spd_1036’* region were present in most (17/26) Δ*gpsB* suppressors (Table 1, row 13). However, Δ(*spd_1029*-*spd_1037*) by itself had no effect on growth in BHI broth (data not shown). We therefore checked whether Δ*gpsB* suppressor strains that have long (135-137 gene) duplications and short (9-12 gene) deletions, such as *sup gpsB-3* and *sup gpsB*-2, had an apparent fitness advantage over Δ*gpsB* suppressor strains that contain (21-149 gene) duplications, but lack duplication-junction deletions, such as *sup gpsB-8* or *sup gpsB*-9 (Fig. 2A, S1B, and S1C; Appendix A, Tab A). Consistent with this idea, the *sup gpsB-2* and *-3* strains grew similarly to WT in BHI broth with higher growth rates and yields than the *sup gpsB-8* and *-9* strains (Table 1, lines 3-4 and 9-10; Fig. S4A). However, Δ*stkP* suppressors containing a *murZ*-duplication/deletion or a *murZ*-duplication alone grew similarly to each other and the parent strain (Table 3, lines1-4; Fig. S4C and S20), indicating a difference between suppression of Δ*gpsB* and Δ*stkP* that was also detected in other experiments described below.

### 2.4 Overexpression of MurZ or MurA or the presence of MurZ(D280Y) suppresses Δ*gpsB* lethality by a protein phosphorylation-independent mechanism

The mutants described above implicated overexpression of pneumococcal *murZ* or *murA* or mutation in *murZ* in the suppression of pneumococcal Δ*gpsB* or Δ*stkP* (Tables 1 and 3). To test this idea further, we constructed merodiploid strains that overexpress MurZ or MurA under the control of a Zn^2+^-inducible promoter from an ectopic site. Overexpression of MurZ, optimally with 0.2 mM Zn inducer (0.2 mM ZnCl_2_ + 0.02 mM MnSO_4_; 0.2 mM (Zn^2+^/(1/10)Mn^2+^), or MurA, optimally with 0.4 mM Zn inducer (0.4 mM (Zn^2+^/(1/10)Mn^2+^), suppressed Δ*gpsB* in transformation assays (Table 2, lines 4-5; Table S5A, lines 35-39). As a control, overexpression of catalytically inactive MurZ(C116S) or MurA(C120S) did not suppress Δ*gpsB* (Table S5A, lines 15-16, 19-20), indicating a requirement for catalytic activity. Western blot controls indicated that cellular amounts of the catalytically deficient proteins were the same as WT (see below).

Suppression of Δ*gpsB* by MurZ or MurA overexpression was confirmed by growth of Δ*gpsB murZ*^+^//P_Zn_-*murZ*^+^ and Δ*gpsB murA*^+^//P_Zn_-*murA*^+^ merodiploid strains in BHI broth containing a range of Zn inducer concentrations (Fig. 4). Depletion of MurZ or MurA in a Δ*gpsB* mutant led to the formation of large, elongated cells that lysed (Fig. 4B, 4D, and S5; no Zn), as reported previously for Δ*gpsB* mutants (Cleverley *et al*., 2019, Land *et al*., 2013, Rued *et al*., 2017). Surprisingly, suppression of Δ*gpsB* was maximal when MurZ was overexpressed by addition of Zn(0.2) inducer, which led to an ≈3.6-fold increase in cellular MurZ amount (Fig. 5C); but, this level of MurZ Induction did not fully restore WT growth or cell morphology to Δ*gpsB* cells (Fig. 4A, 4B, and S5A). In fact, induction of MurZ above this level led to decreased growth rate and yield in the Δ*gpsB* background (Fig. 4A). In contrast, increasing MurA cellular amount to ≈10-fold above WT suppressed Δ*gpsB* and largely restored WT growth and morphology to Δ*gpsB* cells (Fig. 4C, 4D, 5D, and S5B). Besides overexpression of MurZ or MurA, we tested whether overexpression of 21 other proteins involved in pneumococcal division or peptidoglycan synthesis suppressed Δ*gpsB* (Table S6). Overexpression of these proteins did not suppress Δ*gpsB* in transformation assays, while each ectopic construct complemented its corresponding deletion mutation (data not shown). We conclude that moderate (≈4-fold) overexpression of MurZ or MurA is sufficient to restore growth to a Δ*gpsB* mutant, but higher overexpression of MurZ, but not MurA, is deleterious for growth of Δ*gpsB* mutants in BHI broth.

**Figure 4.**
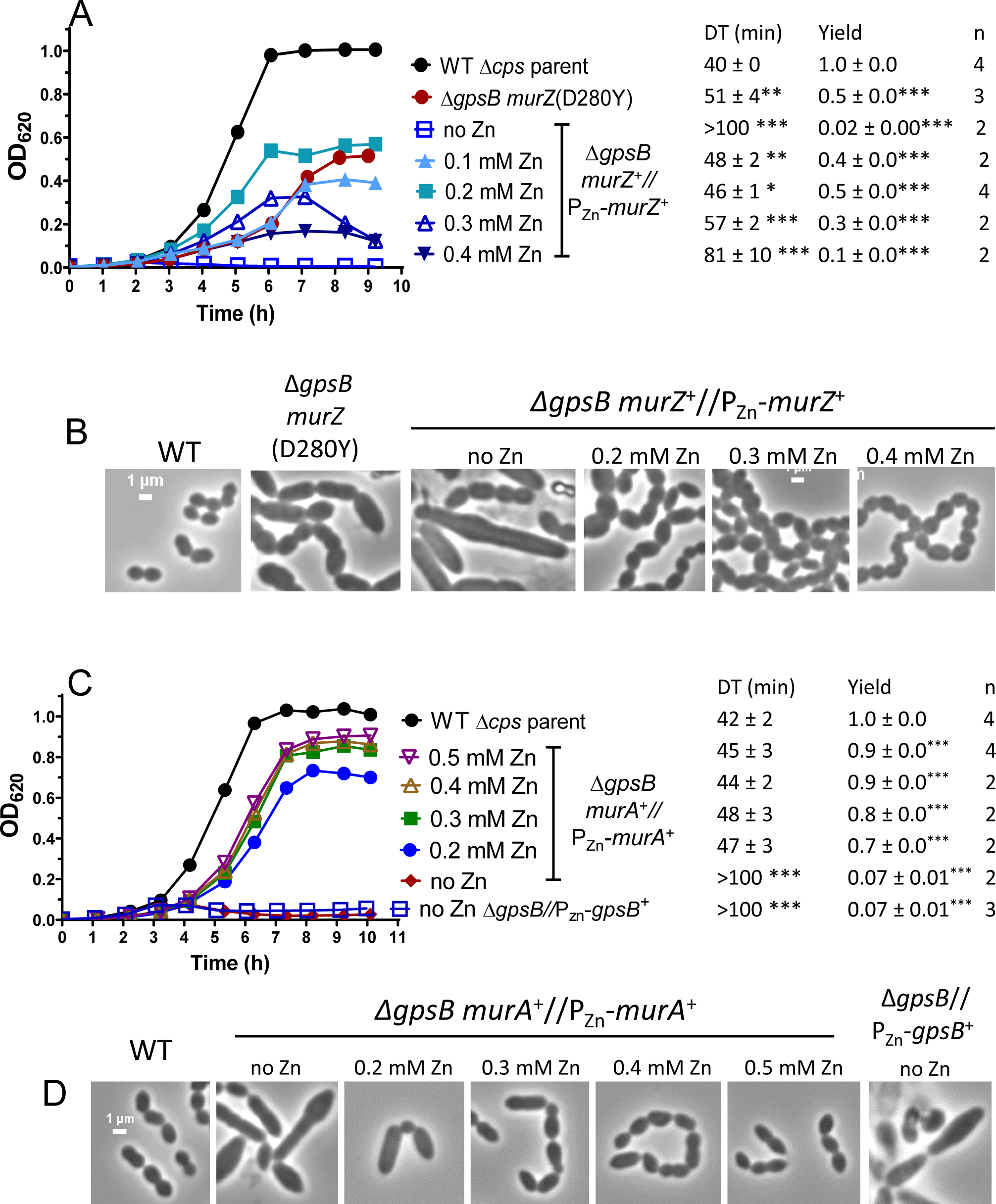
*murZ*(D280Y) and overexpression of *murZ* or *murA* partially suppress Δ*gpsB* growth and morphology phenotypes. (A and B) Parent D39 Δ*cps rpsL1* strain (IU1824), *murZ*(D280Y) Δ*gpsB* strain (IU13509), and Δ*gpsB murZ*^+^//P_Zn_-*murZ*^+^ (IU15860) strain were grown overnight in BHI broth with no (IU1824, IU13509) or 0.2 mM (Zn^2+^/(1/10)Mn^2+^) (IU15860), respectively. Overnight cultures were diluted to OD_620_≈0.003 in the morning in BHI broth for IU1824 and IU13509 and in BHI broth containing Zn^2+^/(1/10)Mn^2+^ for IU15860 as indicated. (A) Left, representative growth curves. Right, averages ± SEMs of doubling times (DT) and maximal growth yields (OD_620_) during 9 h of growth. n denotes number of independent growths. ***, p< 0.001 when compared to WT strain with one-way ANOVA analysis (GraphPad Prism, Dunnett’s test). DTs and growth yields without asterisks were statistically insignificant compared to values obtained from WT. (B) Representative phase-contrast images taken between at 3 to 3.5 h for IU1824, and between 3.5 to 4.5 h for IU13509 and IU15860. Scale bar = 1 µm. (C and D) Parent D39 Δ*cps rpsL1* strain (IU1824), Δ*gpsB murA*^+^//P_Zn_-*murA*^+^ (IU15862), and Δ*gpsB*//P_Zn_-*gpsB*^+^ (IU16370) were grown overnight in BHI broth with no (IU1824) or 0.5 mM (Zn^2+^/(1/10)Mn^2+^) (IU15862 and IU16370). Overnight cultures were diluted to OD_620_≈0.003 in the morning in BHI broth for IU1824 and IU16370 and in BHI broth containing (Zn^2+^/(1/10)Mn^2+^) as indicated for IU15862. Representative growth curves are shown along with averaged DT and growth yields. (D) Representative phase-contrast images taken at 3 h for IU1824 and IU16370 and between 4 to 4.5 h for IU15862. Box-and-whisker plots of cell dimensions of these strains are shown in Fig. S5.

**Figure 5.**
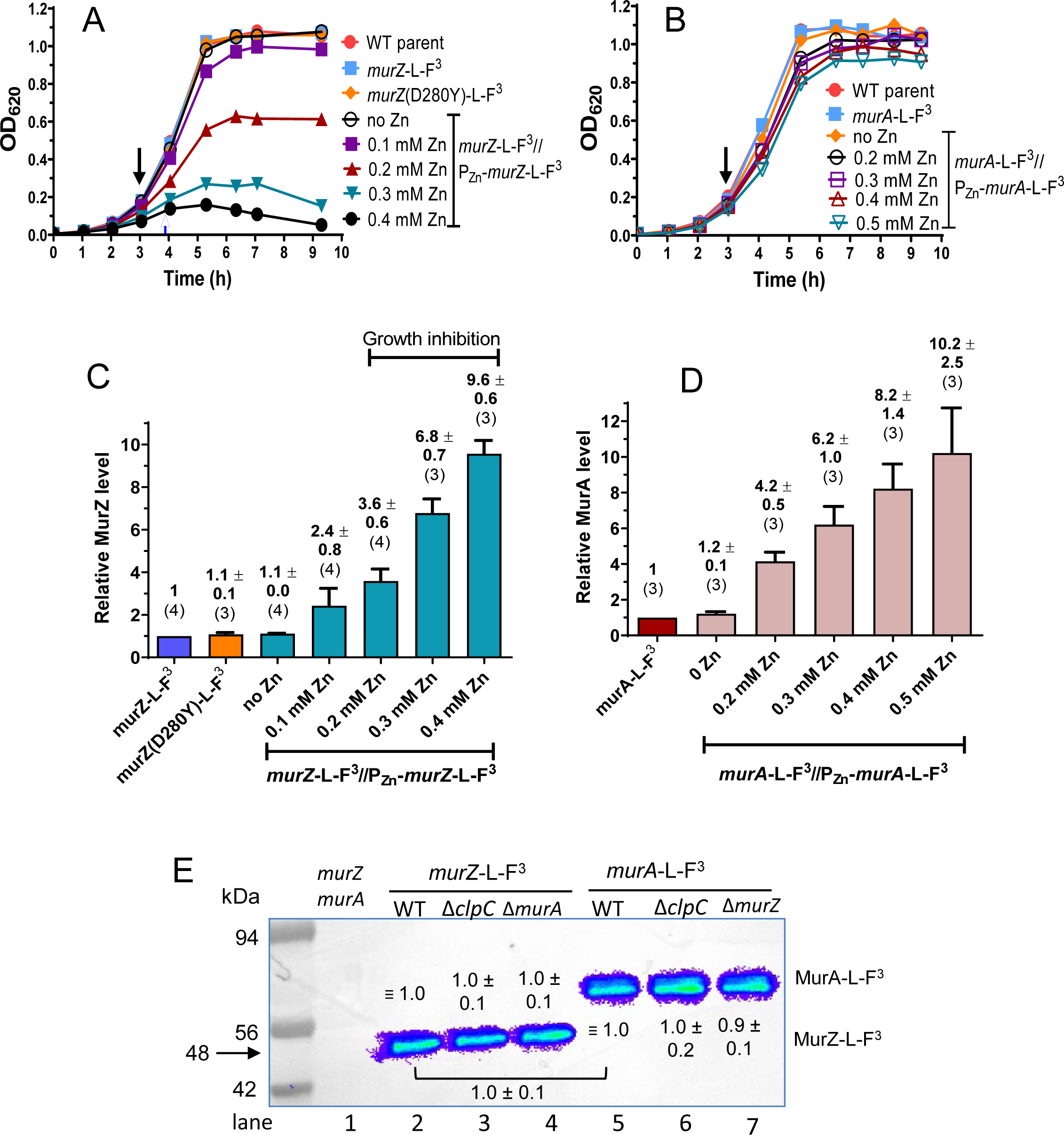
Quantitative western blot assays showing nearly equivalent cellular amounts of MurZ-L-FLAG^3^ (-F^3^), MurZ(D280Y)-L-F^3^, and MurA-L-F^3^, overexpression levels of MurZ-L-FLAG^3^ and MurA-L-FLAG^3^, and lack of change when the other homolog or ClpC is deleted. Strains tested in (A) and (C) were non-FLAG (F) - tagged *murZ* WT (IU1824), *murZ*-L-F^3^ (IU13502), *murZ*(D280Y)-L-F^3^ (IU13600), and *murZ*-L-F^3^//P_Zn_-*murZ*-L-F^3^ (IU13772). Strains tested in (B) and (D) were non-F-tagged *murA* WT (IU1824), *murA*-L-F^3^ (IU14028), and *murA*-L-F^3^//P_Zn_-*murA*-L-F^3^ (IU15983). Strains were grown overnight in BHI broth with no additional (Zn^2+^/(1/10)Mn^2+^), and diluted to OD_620_≈0.005 in the morning in BHI with no additional (Zn^2+^/(1/10)Mn^2+^), or in BHI broth containing 0.1, 0.2, 0.3 or 0.4 mM (Zn^2+^/(1/10)Mn^2+^) for IU13772, or in BHI broth containing 0.2, 0.3, 0.4 or 0.5 mM (Zn^2+^/(1/10)Mn^2+^) for IU15983. Black arrows point to the time (≈3 h) when samples were collected, except for IU13772 grown in the presence of 0.3 or 0.4 mM (Zn^2+^(1/10)Mn^2+^), where samples were collected at 3.6 h (blue arrow). (C) and (D) Quantitative western blotting using anti-FLAG antibody was performed as described in *Experimental procedures.* Calculated averages and SEMs of relative MurZ-L-F^3^ or MurA-L-F^3^ protein amounts were obtained from three or more independent experiments using anti-FLAG antibody. The numbers above each bar are averages ± SEM obtained for the number of independent biological replicates indicated in parentheses. Representative western blots are presented in Fig. S8. (E) Representative western blot showing similar cellular amounts of MurZ-L-F^3^ in Δ*clpC* or Δ*murA* strains as in WT, similar cellular amounts of MurA-L-F^3^ in Δ*clpC* or Δ*murZ* strains as in WT, and similar cellular amounts of WT MurZ-L-F^3^ and WT MurA-L-F^3^. Lane 1, Wild-type (IU1824); lane 2, *murZ*-L-F^3^ (IU13502); lane 3, *murZ*-L-F^3^ Δ*clpC* (IU14082); lane 4, *murZ*-L-F^3^ Δ*murA* (IU14084); lane 5, *murA*-L-F^3^ (IU14028); lane 6, *murA*-L-F^3^ Δ*clpC* (IU14086); lane 7, *murA*-L-F^3^ Δ*murZ* (IU14088). Numbers above MurZ-L-F^3^ or below MurA-L-F^3^ bands are calculated protein amounts (mean ± SEM) relative to *murZ*-L-F^3^ (lane 2) or *murA*-L-F^3^ (lane 5) based on three independent experiments with Δ*clpC* strains and two independent experiments with Δ*murZ or* Δ*murA* strains. 0.67 µg of protein was loaded into each lane. The predicted molecular masses of both MurZ-L-F^3^ and MurA-L-F^3^ are 48 kDa; however, MurA-L-F^3^ (and untagged MurA(*Spn*) (data not shown)) migrate slower than their predicted molecular weights.

We also tested whether the *murZ*(D280Y) mutations identified in the genetic screen suppressed Δ*gpsB*. A constructed isogenic *murZ*(D280Y) mutation suppressed Δ*gpsB* in transformation assays (Table 2, lines 1 and 6). However, the growth rate and yield of the Δ*gpsB murZ*(D280Y) mutant were considerably reduced compared to the WT strain (Fig. 4A), and Δ*gpsB murZ*(D280Y) cells were extremely large and elongated compared to WT cells (Fig. 4B and S5B), similar to Δ*gpsB* cells depleted for MurZ that stop growing (Fig. 4A). We conclude that the *murZ*(D280Y) mutation only partly suppresses the defects caused by Δ*gpsB*.

Finally, we assayed whether overexpression of MurZ or MurA or the presence of MurZ(D280Y) restored general protein phosphorylation in a Δ*gpsB* mutant. It was previously reported that Δ*gpsB* greatly reduces protein phosphorylation by the StkP Ser/Thr kinase in *S. pneumoniae*, leading to the model that GpsB activates StkP function (Fleurie *et al*., 2014, Rued *et al*., 2017). We showed that ΩZ.1Δ *gpsB* suppressors *sup gpsB-2* and *sup gpsB-3* did not restore protein phosphorylation, whereas the *phpP* phosphatase mutation in *sup gpsB-1* restored phosphorylation (Rued *et al*., 2017). Likewise, all new ΩZ.1 and ΩZ.1Δ duplications that suppressed Δ*gpsB* from this study did not restore protein phosphorylation (Fig. S6A), while *phpP* mutations that suppressed Δ*gpsB* did restore phosphorylation (Fig. S6B). Overexpression of MurZ or MurA or *murZ*(D280Y) also failed to restore protein phosphorylation in a Δ*gpsB* mutant (Fig. S7A, lanes 5 and 9; Fig. S7B, lane 6). We conclude that suppression of Δ*gpsB* by overexpression of MurZ or MurA or by MurZ(D280Y) occurs by a phosphorylation-independent mechanism.

### 2.5 Overexpression, absence, or catalytic inactivation of MurZ(*Spn*), but not MurA(*Spn*), results in altered growth, morphology, and sensitivity to fosfomycin or penicillin

The relative contribution of MurZ and MurA in pneumococcal cells is not well understood. Purified MurZ(*Spn*) from strain R6 has a higher catalytic efficiency for UDP-GlcNAc substrate than MurA (*Spn*) (Du *et al*., 2000). By contrast, the MurA-family homolog is essential or catalytically predominant in other Gram-positive species (Fig. 1) (Blake *et al*., 2009, Kedar *et al*., 2008, Kock *et al*., 2004, Mascari *et al*., 2022, Rismondo *et al*., 2017). The growth defects of MurZ(*Spn*) overexpression in the Δ*gpsB* mutant (Fig. 4A-B) prompted us to further characterize the relative roles of MurZ and MurA in WT pneumococcal cells.

The absence of MurZ and MurA was confirmed to be synthetically lethal in *S. pneumoniae* D39 (Table S5B, line 2 and S5C, line 2)(Du *et al*., 2000). Catalytically active MurZ(C116S) and MurA(C120S) also were synthetically lethal with lack of MurA or MurZ, respectively (Table S5B, line 3 and S5C, line 3). Strains expressing MurZ-L-FLAG^3^ or MurA-L-FLAG^3^ from their native chromosomal loci were constructed (Table S1), and expression levels were assayed by quantitative western blotting (Fig. 5). Strains expressing MurZ-L-FLAG^3^ or MurA-L-FLAG^3^ did not show phenotypic differences in growth or transformation assays compared to their WT counterparts, including synthetic lethality (Fig. 5A-B, 6A, and 6C; Table S5B, line 4 and S5C, line 5). Consistent with comparable activities, high overexpression of MurZ-L-FLAG^3^ inhibited growth like MurZ overexpression (Fig. 5A and 6A). MurZ-L-FLAG^3^ and MurA-L-FLAG^3^ amounts were comparable (ratio = 0.95 ± 0.06 (SEM; n= 2)) in bacteria growing exponentially in BHI broth (Fig. 5E). Immunofluorescent microscopy showed that MurZ-L-FLAG^3^ and MurA-L-FLAG^3^ were distributed throughout the cytoplasm, and not localized at division septa or equators (Fig. S9).

**Figure 6.**
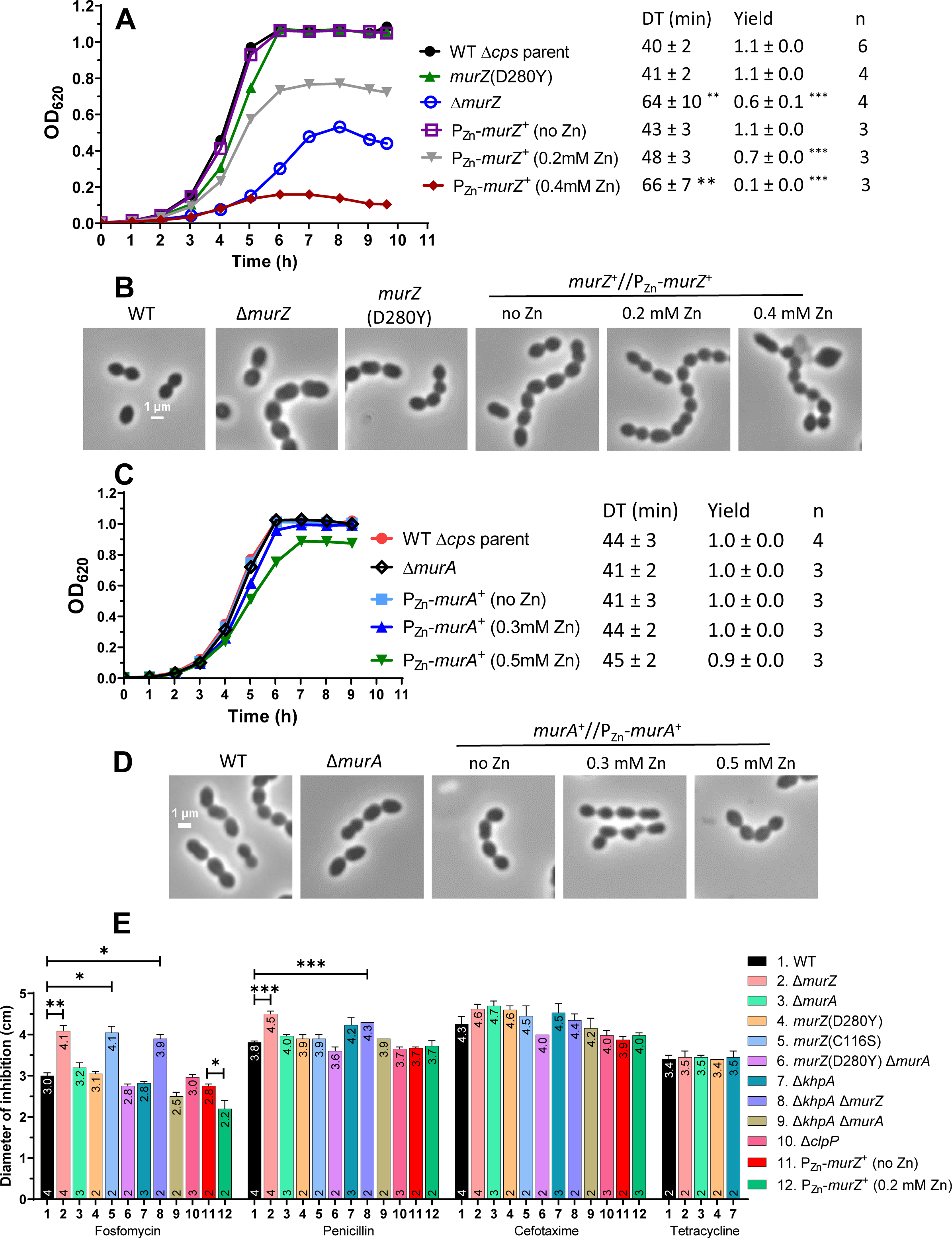
Overexpression or absence of MurZ(*Spn*), but not MurA(*Spn*), alters growth, morphology, and sensitivity to fosfomycin or penicillin. (A and B) Parent D39 Δ*cps rpsL1* strain (IU1824), constructed *murZ*(D280Y) (IU13438), Δ*murZ* (IU13536), and merodiploid *murZ*^+^//P_Zn_-*murZ*^+^ (IU13393) strains were grown overnight in BHI broth with no additional (Zn^2+^/(1/10)Mn^2+^) and diluted to OD_620_ ≈0.003 in the morning in BHI broth with or without (Zn^2+^/(1/10)Mn^2+^) at the concentrations indicated. (A) Representative growth curves and averaged DT and yields. ** p< 0.01; *** p < 0.001 compared to WT strain by one-way ANOVA analysis (GraphPad Prism, Dunnett’s test). (B) Representative phase-contrast images taken between 3.5 to 4 h of growth for all strains and conditions, except for IU13393 with 0.4 mM (Zn^2+^(1/10)Mn^2+^), which was taken at 5 h of growth. (C and D) Parent D39 Δ*cps rpsL1* strain (IU1824), Δ*murA* (IU13538), and merodiploid *murA*^+^//P_Zn_-*murA*^+^ (IU13395) strains were grown similarly to the *murZ* strains described above. The DTs and growth yields of all strains and conditions were not statistically different from the values obtained for the WT strain. (D) Representative phase-contrast images taken at 3 h of growth for all strains and conditions. All micrographs in (B) and (D) are at the same magnification (scale bar = 1 µm). Box-and-whisker plots of cell dimensions of *murZ*(D280Y) and strains overexpressing MurZ or MurA are in Fig. S10. (E) Disc diffusion assays were performed as described in *Experimental procedures* for strains: WT parent (IU1824), Δ*murZ* (IU13536), Δ*murA* (IU13538), *murZ*(D280Y) (IU13438), *murZ*(C116S) (IU15939), *murZ*(D280Y) Δ*murA* (IU17748), Δ*khpA* (IU9036), Δ*khpA* Δ*murZ* (IU13542), Δ*khpA* Δ*murA* (IU13546), Δ*clpP* (IU12462), *murZ*^+^//P_Zn_-*murZ*^+^ (no Zn) (IU13393), and *murZ*^+^//P_Zn_-*murZ*^+^ in 0.2 mM (Zn^2+^/(1/10)Mn^2+^). Mean diameters of zones of inhibition ± SEM are graphed from at least two independent biological replicates. Means and numbers of replicates (n) are shown at the tops and bottoms of bars, respectively. P values were obtained by the Welch t-test (GraphPad Prism). *, **, and *** denote p<0.05, p<0.01, p<0.001, respectively.

However, further experiments indicated that MurZ and MurA function was not equivalent and interchangeable in cells grown in BHI broth. Δ*murZ* mutants grew slower, had a lower growth yield, and formed larger cells than Δ*murA* mutants in exponential cultures (Fig. 6A-D, and S10A-B). While overexpression of MurZ by ≈2-fold did not change growth (Fig. 5A; Zn (0.1)), higher overexpression of MurZ by ≈4-10 fold progressively reduced growth rate and yield and led to smaller cells with increasingly defective morphologies (Fig. 5A-B, 6A-B, and S10A). In contrast, overexpression of MurA by ≈4-10 fold did not affect cell growth or morphology (Fig. 5B, 5D, 6C-D, and S10B). Control experiments showed that the growth and size phenotypes of Δ*murZ* mutants were complemented by ≈2-fold overexpression of MurZ (Fig. S11A-C, Zn(0.1) and Fig. 5C). Δ*murZ* growth and morphology defects were also complemented by overexpression of MurA by ≈4-6-fold (Fig. S12A-C, Zn(0.2) and Fig. 5D)), but not fully at Zn(0.1), indicating that greater induction of MurA than MurZ was required to complement Δ*murZ*.

We looked for other indications of differences in the relative roles of pneumococcal MurZ and MurA. We found that the absence of MurZ or overexpression of MurZ, but not MurA, caused similar growth defects or inhibition, respectively, in the isogenic encapsulated *cps*^+^ D39 progenitor strain as in Δ*cps* mutants (Fig. S13). In the Δ*cps* unencapsulated background, Δ*murZ* and catalytically inactive *murZ*(C116S) mutants were more sensitive to fosfomycin, which covalently binds to the catalytic cysteine of MurA enzymes (Skarzynski *et al*., 1996), than a Δ*murA* mutant in disk-diffusion assays (Fig. 6E). Δ*murZ* mutants were also slightly more sensitive to the β-lactam antibiotic penicillin (Fig. 6E). Conversely, moderate overexpression of MurZ reduced sensitivity to fosfomycin compared to WT. Δ*murZ* and Δ*murA* mutants were equally sensitive as WT to the cephalosporin antibiotics cefotaxime or cefoperazone, and to tetracycline, which inhibits translation (Fig. 6E and data not shown). We also tested whether depletion of MurA in Δ*murZ* or depletion of MurZ in Δ*murA* caused the elongated-cell phenotype characteristic of GpsB depletion in the D39 background (Land *et al*., 2013, Rued *et al*., 2017). To the contrary, reduced amounts of MurZ and MurA inhibited growth and caused formation of rounded, heterogeneously sized cells that began to lyse (Fig. S14A-B). This result is consistent with GpsB having additional roles besides regulating MurZ and MurA function.

Mutants expressing catalytically inactive MurZ(C116S)-L-FLAG^3^ or MurZ(C116S) phenocopied Δ*murZ* by showing impaired growth (Fig. S15A and S16C). By contrast, a mutant expressing catalytically inactive MurA(C120S)-L-FLAG^3^ did not affect growth, similar to Δ*murA* (Fig. S15A). Quantitative western blotting showed that MurZ(C116S)-FLAG^3^ or MurA(C120S)-FLAG^3^ were expressed at the same level as MurZ-L-FLAG^3^ or MurA-L-FLAG^3^, respectively (Fig. S15B). Consistent with its lack of catalytic activity, overexpression of MurZ(C116S) did not cause growth inhibition like WT MurZ (Fig. S15C). This result indicated that MurZ(C116S) is not dominant-negative over WT MurZ, consistent with a MurZ monomer in cells as well as in purified preparations(Du *et al*., 2000). Finally, we tested whether the absence of MurZ inhibited cell growth and caused defective cell morphology in C+Y medium, as occurred in BHI broth (Fig. 6A-B). We found that the absence of MurZ or MurA or their catalytic inactivation did not inhibit growth in C+Y medium (Fig. S16A). However, lack of MurZ or its catalytic activity resulted in longer, wider, and larger cells than WT in C+Y medium (Fig. S16B-C), similar to BHI broth (Fig. 6B), whereas Δ*murA* and WT cells were the same size (data not shown). Altogether, we conclude that MurZ(*Spn*) and MurA(*Spn*) function is not equivalent in exponentially growing D39 cells and that in most cases, phenotypes of *murZ* mutants are more severe than those of *murA* mutants, consistent with a predominant role of MurZ in *S. pneumoniae* D39 cells.

### 2.6 *murZ*(D280Y), *murZ*(I265V) present in laboratory strains R6 and Rx1, and *murZ*(E259A) alleles suppress Δ*gpsB*

*murZ*(D280Y) was isolated as a spontaneous suppressor of Δ*gpsB* (Table 1, line 12), and partial Δ*gpsB* suppression was confirmed in a reconstructed *murZ*(D280Y) mutant (Table 2, line 6). Compared to WT, *murZ*(D280Y) Δ*gpsB* double mutants formed smaller colonies in transformation assays (Table 2, line 6), had reduced growth rate and yield (Fig. 4A), and formed large, aberrantly shaped cells (Fig. 4B and S5A). However, a single *murZ*(D280Y) mutant grew similarly to WT, formed marginally smaller (by 10%-20%) cells than WT in BHI broth, and showed the same sensitivity to fosfomycin or penicillin as WT or a *murZ*(D280Y) Δ*murA* mutant (Fig. 6A, 6B, 6E, S10A, and S17C). Overexpression of MurZ(D280Y) also inhibited growth of *murZ^+^* or *murZ*(D280Y) merodiploid strains, similar to overexpression of MurZ (Fig. S17). MurZ(D280Y) was expressed in approximately the same amount as MurZ in cells growing exponentially in BHI broth (Fig. 5A). Finally, whereas *murZ*(D280Y) partially suppresses Δ*gpsB* in transformation assays (Table 2, line 6; small colonies), it strongly suppressed Δ*stkP* and the requirement for protein phosphorylation (Fig. 9A-C, Fig. S3E and S21 A-B and E; Table 2, line 6; WT colonies). Together, these results suggest that MurZ(D280Y) has comparable enzymatic activity and cellular amount as MurZ, but is not subjected to negative regulation that occurs in Δ*gpsB* or Δ*stkP* mutants.

The MurZ(D280Y) amino-acid change is located in Domain I on a surface distant from the active site of MurZ, which includes C116 (catalysis), N23 (conformation switching), D306 (deprotonation of substrate), and R398 (product release) (Fig. 7) (Jackson *et al*., 2009, Samland *et al*., 2001, Skarzynski *et al*., 1996). Compared to D39 strains (and WT serotype-4 strain TIGR4 (Tettelin *et al*., 2001)), R6 and Rx1 laboratory strains produce mutant MurZ(I265V) (Lanie *et al*., 2007), which has an amino-acid change near MurZ(D280Y) (Fig. 7). Like *murZ*(D280Y), *murZ*(I265V) moved into the Δ*cps* D39 genetic background partially suppressed Δ*gpsB* and strongly suppressed Δ*stkP* in transformation assays (Table 2, lines 6-7; Fig. S3E-F and S18A-D), and D39 Δ*cps murZ*(I265V) partially suppressed Δ*gpsB* in growth and morphology assays (Fig. S18A-B). Both *murZ*(D280Y) Δ*gpsB* and *murZ*(I265V) Δ*gpsB* double mutants formed large, elongated cells (Fig. 4B and S18B), reminiscent of strains depleted for GpsB (Land *et al*., 2013, Rued *et al*., 2017). Both *murZ*(I265V) and *murZ*(D280Y) strains grew similarly to WT (Fig. 5A, 6A, and S18A); however, *murZ*(D280Y) cells were marginally smaller than WT and *murZ*(I265V) cells under these growth conditions (Fig. S10A and S18E). Finally, Δ*gpsB* could not be transformed into an R6 Δ*murZ* mutant, and Δ*murZ* could not be transformed into an R6 Δ*gpsB* mutant, consistent with a requirement for the *murZ*(I265V) allele to suppress Δ*gpsB* in R6-derived strains (Tables S5A, lines 40-42, and S5B, line 7).

**Figure 7.**
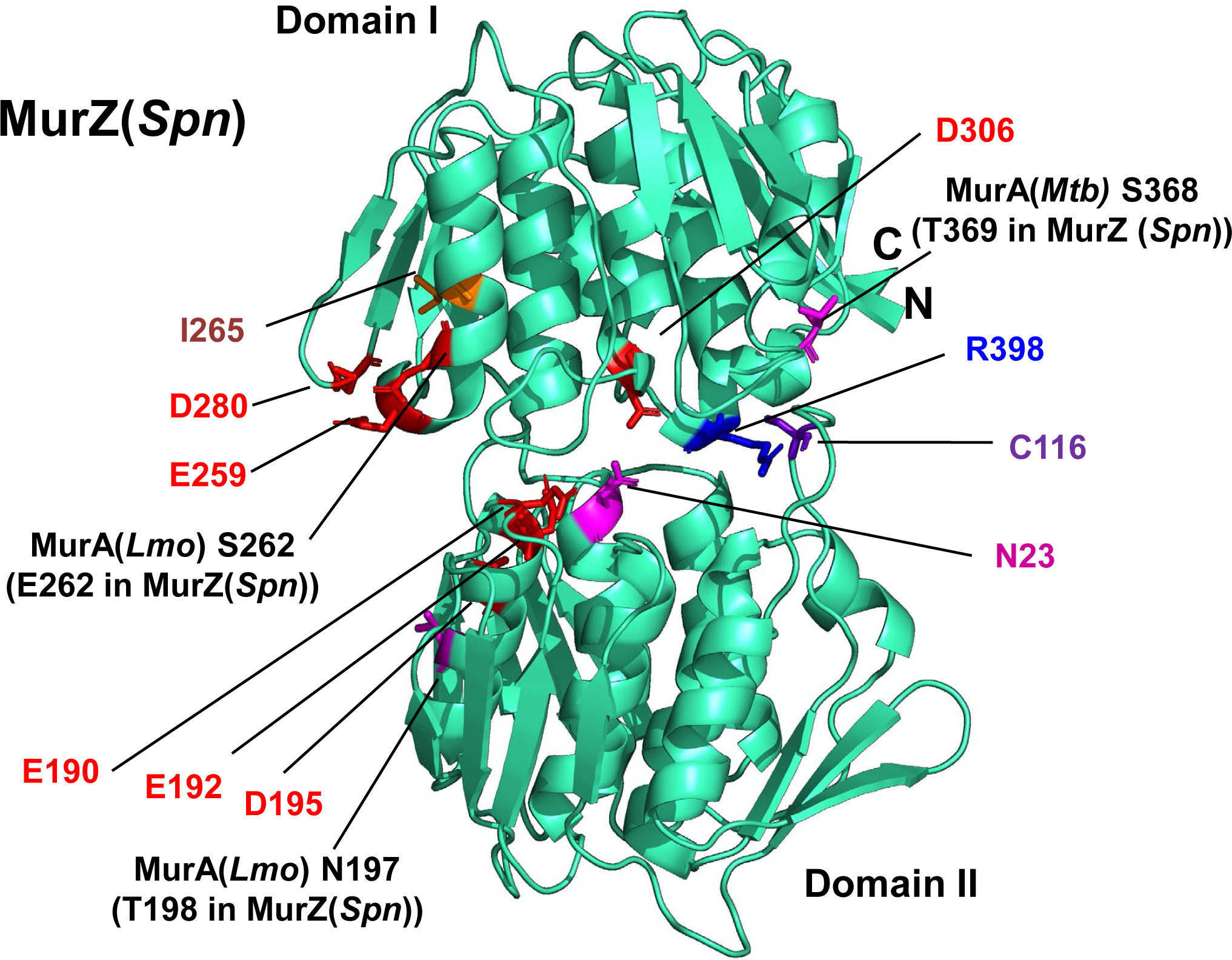
MurZ(D280Y), MurZ(E259A), and MurZ(I265V) that suppress Δ*gpsB* or Δ*stkP* are located on a face of Domain I of MurZ, away from its active site. The predicted 3D-structure of MurZ(*Spn*) from D39 strains generated using the AlphaFold v2.0 webserver is shown in cyan, with important residues illustrated as colored sticks. Catalytic site C116, and other residues important for MurA enzymatic activity include N23 (conformation switching), D306 (initial deprotonation of the UDP substrate), and R398 (product release) (Jackson *et al*., 2009, Samland *et al*., 2001, Skarzynski *et al*., 1996). Although N23 and C116 are in Domain II, and D306 and R398 are in Domain I, these four residues are in close proximity on one side of the molecule. In contrast, D280, E259, and I265, for which amino acid substitutions lead to Δ*gpsB* suppression, are located on the opposite side Domain I compared to C116. E190, E192 and D195 are in Domain II across the cleft from D280 and do not lead to Δ*gpsB* suppression when substituted. Residues T198 and E262 correspond to residues MurA(*Lmo*) N197 and MurA(*Lmo*) S262 respectively. MurA(*Lmo*) N197D and MurA(*Lmo*) S262L are suppressor mutations of Δ*gpsB* and Δ*prkA* mutations in *Listeria monocytogenes* (Wamp et al., 2021).

Based on structure, MurZ(E259) is on the same surface as MurZ(D280Y) and MurZ(I265V) (Fig. 7). *murZ*(E259A) also partly suppressed Δ*gpsB* and strongly suppressed Δ*stkP* in transformation assays (Tables 2, line 8, and S5A, line 7). In contrast, analogous amino acid changes in MurA(D281Y) and MurA(E282Y) did not suppress Δ*gpsB* (Table S5A, lines 11-12). Finally, MurZ(E190A E192A), MurZ(E192A), and MurZ(E195A), which contain amino-acid changes in Domain II on the same side of MurZ as Domain I suppressors MurZ(D280Y), MurZ(I265V), and MurZ(E259A), failed to suppress Δ*gpsB* (Table S5A, lines 8-10). We conclude that the Domain I surface close to MurZ(D280) specifically mediates escape from negative regulation that occurs in Δ*gpsB* mutants.

### 2.7 MurZ and MurA are not degraded by ClpP protease in *S. pneumoniae*

MurA(*Lmo*) (the homolog of MurA(*Spn*); Fig. 1), accumulates to a high level (≈10-fold over WT) in Δ*murZ*(*Lmo*) (the homolog of *murZ*(*Spn*)) or Δ*clpC*(*Lmo*) mutants of *L. monocytogenes* (Rismondo *et al*., 2017, Wamp *et al*., 2020). Likewise, MurAA(*Bsu*) (the homolog of MurA (*Spn*); Fig. 1) is a substrate of the ClpCP protease of *B. subtilis* (Kock *et al*., 2004), although only a marginal increase in MurAA(*Bsu*) amount was detected in a Δ*clpC* mutant in a recent study (Sun *et al*., 2023). Cleavage of MurA(*Lmo*) by the ClpCP protease is central to the model of the regulation of MurA(*Lmo*) cellular amount by MurZ(*Lmo*) and the ReoM and ReoY regulatory proteins in *L. monocytogenes* (Wamp *et al*., 2022, Wamp *et al*., 2020). In support of this model, Δ*clpC*, Δ*murZ*, Δ*reoM*, or Δ*reoY* suppressed Δ*gpsB* or Δ*prkA* (lacking Ser/Thr protein kinase) in *L. monocytogenes* (Rismondo *et al*., 2017, Wamp *et al*., 2020, Wamp *et al*., 2022).

Several different results indicate that MurZ and MurA cellular amounts are not interrelated or regulated by the ClpP protease and its ATPase subunits in *S. pneumoniae.* First, ClpP is not essential in *S. pneumoniae* D39 (Fig. 8)(Robertson *et al*., 2003), and Δ*clpP* did not suppress Δ*gpsB* or Δ*stkP* (Table 2, line 11; Fig. S3B). In addition, Δ*clpC*, Δ*clpE*, or Δ*clpL* mutants, which lack ATPase subunits of ClpP, did not suppress Δ*gpsB* (Table S5A, lines 23-26). Second, a Δ*clpC* mutant did not decrease sensitivity of fosfomycin, which would have been indicative of increased MurZ(*Spn*) or MurA(Spn) amount (Fig. 6E). Third, results presented above demonstrate that MurZ-L-FLAG^3^ and MurA-L-FLAG^3^ expressed from their native chromosomal loci phenocopied WT MurZ and MurA (Fig. 5-6). Quantitative western blotting showed that the cellular amount of MurZ-FLAG^3^ or MurA-FLAG^3^ was not changed by Δ*murA* or Δ*murZ*, respectively (Fig. 5E). Δ*clpE*, Δ*clpL,* Δ*clpP,* Δ*clpC* also did not change cellular MurZ-L-FLAG^3^ or MurA-L-FLAG^3^ amount (Fig. 8).

**Figure 8.**
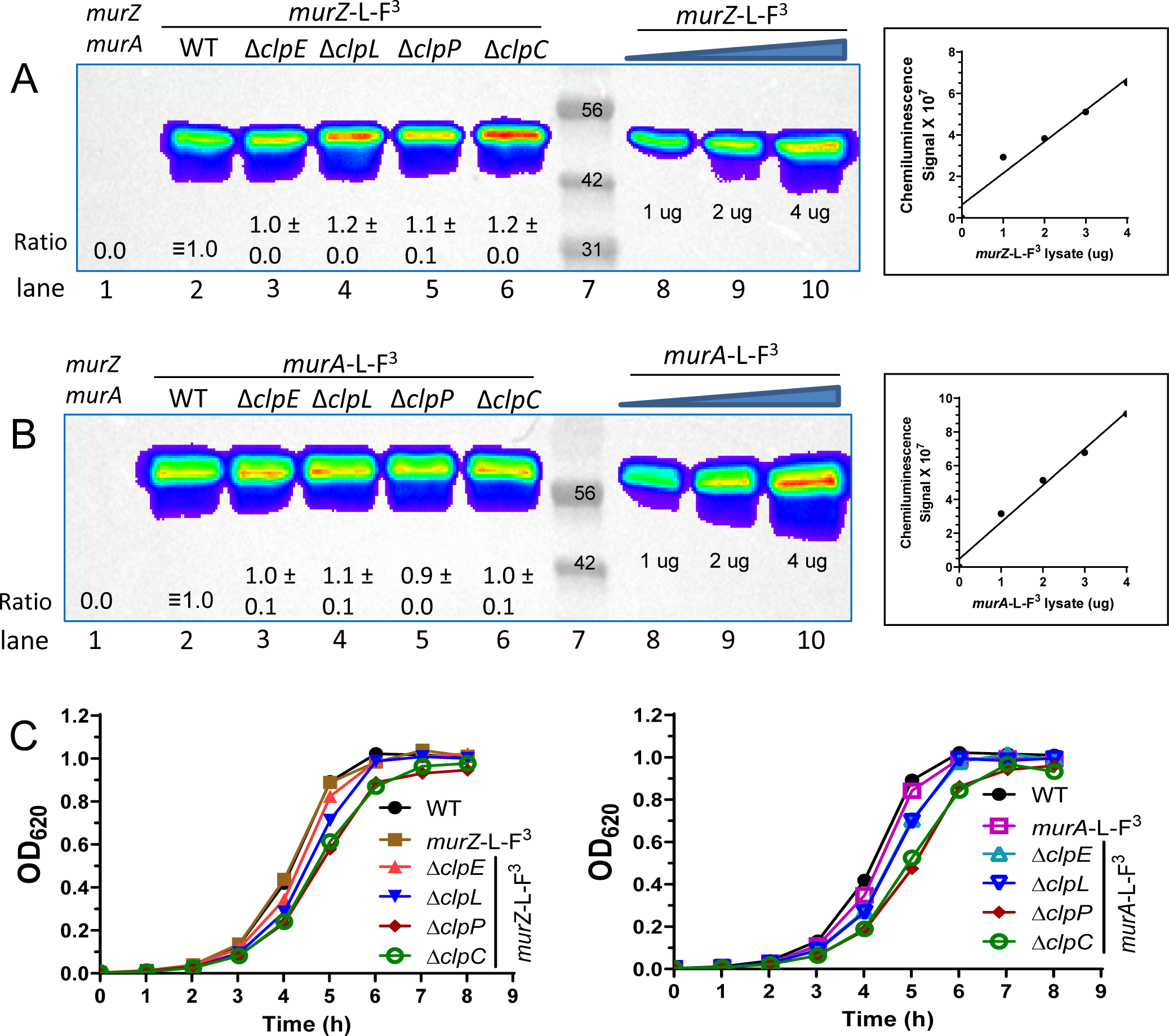
MurZ(*Spn*) and MurA(*Spn*) cellular amounts are unchanged in Δ*clpP*, Δ*clpC*, Δ*clpL*, or Δ*clpE* mutants lacking the ClpP protease or its ATPase subunits. (A) Western blot using anti-FLAG antibody of samples obtained from WT parent (IU1824), *murZ*-L-F3 (IU13502), *murZ*-L-F^3^ Δ*clpE* (IU17150), *murZ*-L-F^3^ Δ*clpL* (IU17152), *murZ*-L-F^3^ Δ*clpP* (IU17154), and *murZ*-L-F^3^ Δ*clpC* (IU14082). (B) Western blot of samples obtained from WT parent (IU1824), *murA*-L-F^3^ (IU14028), *murA*-L-F^3^ Δ*clpE* (IU17158), *murA*-L-F^3^ Δ*clpL* (IU17160), *murA*-L-F^3^ Δ*clpP* (IU17162), and *murA*-L-F^3^ Δ*clpC* (IU14086). 3 µg of each protein was loaded onto lanes 1-6, and 1, 2, or 4 µg of either *murZ*-L-F^3^ (A) or *murA*-L-F^3^ (B) lysates were loaded in lanes 8-10 to generate standard curves for quantitation. Plots of µg of lysate obtained from IU13502 or IU14028 loaded vs chemiluminescence signal intensities are shown to the right of the blots. Calculated protein amounts (mean ± SEM) relative to *murZ*-L-F^3^ (lane 2) or *murA*-L-F^3^ (lane 2) based on two independent experiments are shown. (C) Growth curves of strains used in (A) and (B).

Fourth, it could be argued that the C-terminal epitope tags interfere with degradation of MurZ-L-FLAG^3^ and MurA-L-FLAG^3^ by ClpCP. If this were true, then MurZ-L-FLAG^3^ or MurA-L-FLAG^3^ should suppress Δ*gpsB.* This was found not to be the case in transformation assays (Table S5A, lines 27-28). Fifth, consistent with this conclusion, moving epitope tags from C-termini to N-termini destabilized MurZ and MurA, but the amounts of remaining HT-MurZ or HT-MurA detected did not change in a Δ*clpP* mutant (Fig. S19C). Sixth, polyclonal antibody to purified MurAA(*Efa*) (the closest homolog of MurA(*Spn*); Fig. 1) cross-reacted with overexpressed native MurA(*Spn*). Δ*clpP* or Δ*murZ* did not change the cellular amount of overexpressed native MurA(*Spn*) (data not shown). We conclude that a ClpP-protease dependent mechanism does not regulate the amounts of MurZ and MurA in *S. pneumoniae* D39, in contrast to MurA(*Lmo*) or MurAA(*Bsu*). Furthermore, the relative cellular amounts of MurZ(*Spn*) or MurA (Spn) are unchanged in Δ*murA* or Δ*murZ* mutants, respectively (Fig. 5E).

### 2.8 *murZ*(D280Y) and overexpression of MurZ or MurA strongly suppress primary morphology phenotypes of StkP(*Spn*) depletion

Δ*stkP* mutants have been extensively characterized in R6 and Rx1 laboratory strains that contain *murZ*(I265V), which suppresses Δ*stkP* (Table 2, line 7; Fig. S3F and S18C-D) (Beilharz *et al*., 2012, Echenique *et al*., 2004, Fleurie *et al*., 2012, Novakova *et al*., 2010, Pinas *et al*., 2018, Saskova *et al*., 2007, Ulrych *et al*., 2016, Zucchini *et al*., 2018). Δ*stkP* mutants have also been isolated in D39 and TIGR4 strains (Beilharz *et al*., 2012, Giefing *et al*., 2010, Herbert *et al*., 2015, Kant *et al*., 2023), where chromosomal duplications and other suppressors may have arisen. In our experiments, transformants of a Δ*stkP*::P_c_-*erm* amplicon into D39 Δ*cps* strains resulted in extremely faint colonies that when re-streaked, produced colonies of variable sizes containing suppressor mutations (Table 3; Fig. S3) (Rued *et al*., 2017). The faint-colony phenotype of Δ*stkP* transformants was complemented by ectopic expression of *stkP*^+^ (Fig. S3C-D).

To resolve whether Δ*stkP* is essential in D39 strains, we compared Tn-seq analysis of the unencapsulated WT to a Δ*khpB* mutant that suppresses the requirement for *stkP* (below; Table 2, lines 9-10). Viable insertions in *stkP* were obtained in the WT strain only in the C-terminal 144 amino acid region that contains the third and fourth extracellular PASTA domains (P3 and P4) (Fig. 10A), indicating that the intracellular, transmembrane domain, and the first two PASTA domains (P1 and P2) are essential for exponential growth in BHI broth in 5% CO_2_. The first TA insertion occurs in the WT strain at the TAT(Y515) codon, creating a TAA stop codon (Fig. 10A), and no insertions were detected upstream of TTA(L512) codon, indicating that StkP(M1-L512) are essential under the growth conditions tested. Moreover, the same WT Tn-seq insertion profile was obtained for encapsulated D39 strain IU1781 as for unencapsulated strain IU1824 growing in BHI broth or for IU1824 growing in C+Y, pH 6.9 medium in 5% CO_2_ (Fig. 10A-C; data not shown).

**Figure 9.**
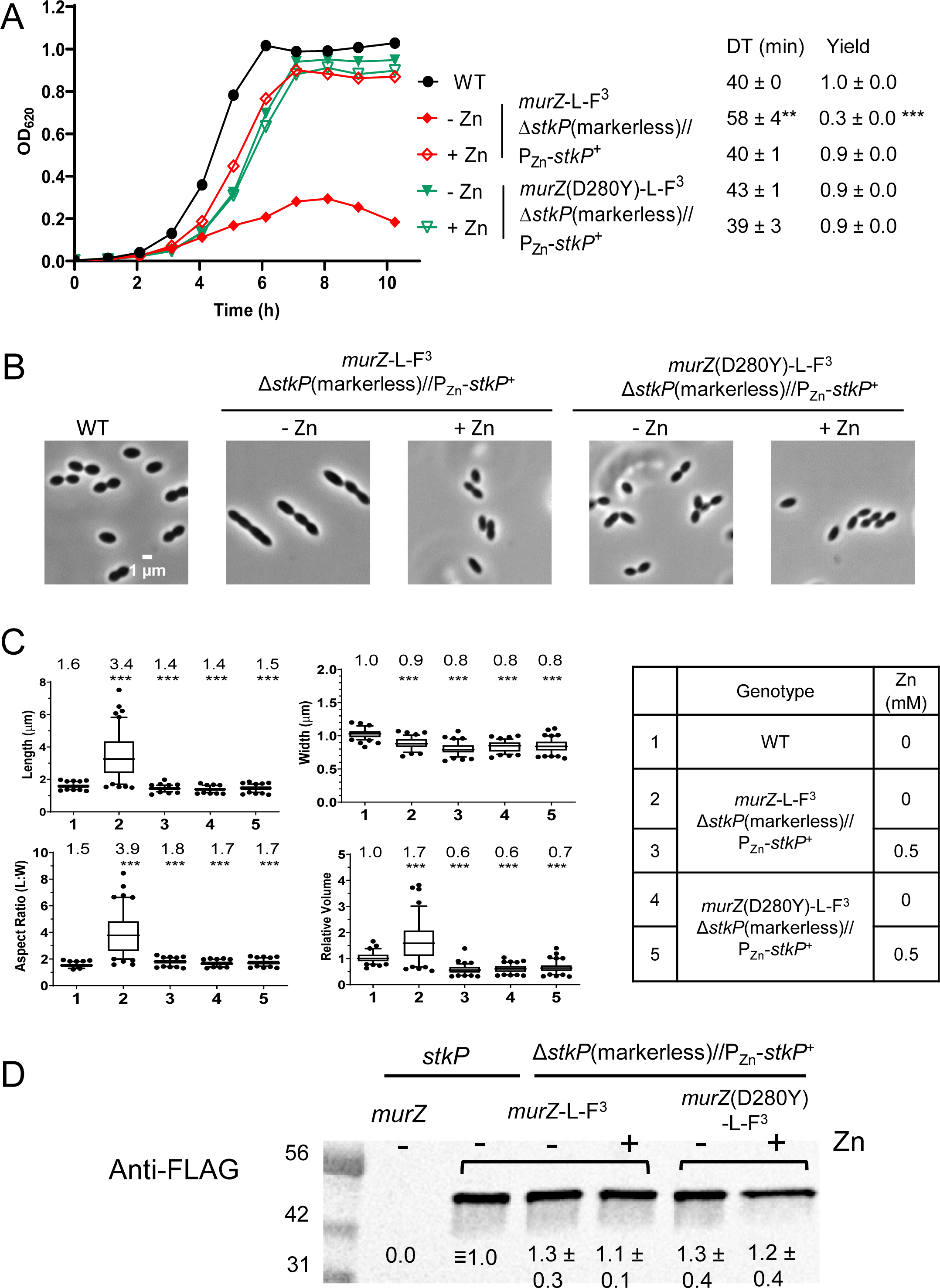
Primary phenotypes of StkP(*Spn*) depletion are strongly suppressed by *murZ*(D280Y). Parent D39 Δ*cps rpsL1* strain (IU1824), and merodiploid Δ*stkP*(markerless)*//*P_Zn_-*stkP*^+^ strains containing *murZ*-L-FLAG^3^ (IU19081) or *murZ*(D280Y)-L-FLAG^3^ (IU19079) were grown overnight in BHI broth with no additional (Zn^2+^/(1/10)Mn^2+^) (IU1824) or with 0.5 mM (Zn^2+^(1/10)Mn^2+^) (IU19081 and IU19079) as described in *Experimental procedures*. Strains were diluted to OD_620_ ≈0.003 in the morning with fresh BHI broth containing no (Zn^2+^/(1/10)Mn^2+^) or 0.5 mM (Zn^2+^/(1/10)Mn^2+^). (A) Growth curves, DT, and maximal growth yields (OD_620_) during 10 h of growth. (B) Representative phase-contrast images taken at ≈3.5 h of growth. Scale bar = 1 µm. Growth curves and microscopy were performed in two independent experiments. (C) Box- and-whisker plots (whiskers, 5 and 95 percentile) of cell lengths, widths, aspect ratios, and relative cell volumes. P values were obtained by one-way ANOVA analysis (GraphPad Prism, Kruskal-Wallis test). *** p<0.001 compared to WT. (D) Representative western blot using anti-FLAG antibody of samples collected after 3.5 h of growth, where – or + indicates the absence of presence of 0.5 mM (Zn^2+^/(1/10)Mn^2+^) in the BHI broth. Western blotting was performed as described in *Experimental procedures*. 6 µL (≈2 µg) of protein samples were loaded in each lane. A standard curve was generated by loading 3, 6, 9 or 12 µL of IU13502 (*murZ*-L-FLAG^3^) samples (lanes not shown). Signal intensities obtained with anti-StkP antibody were normalized in each lane by using Totalstain Q-NC reagent (Azure Biosystems). Calculated protein amounts (mean ± SEM) relative to *stkP*^+^ *murZ*-L-F^3^ (IU13249) are based on two independent experiments.

**Figure 10.**
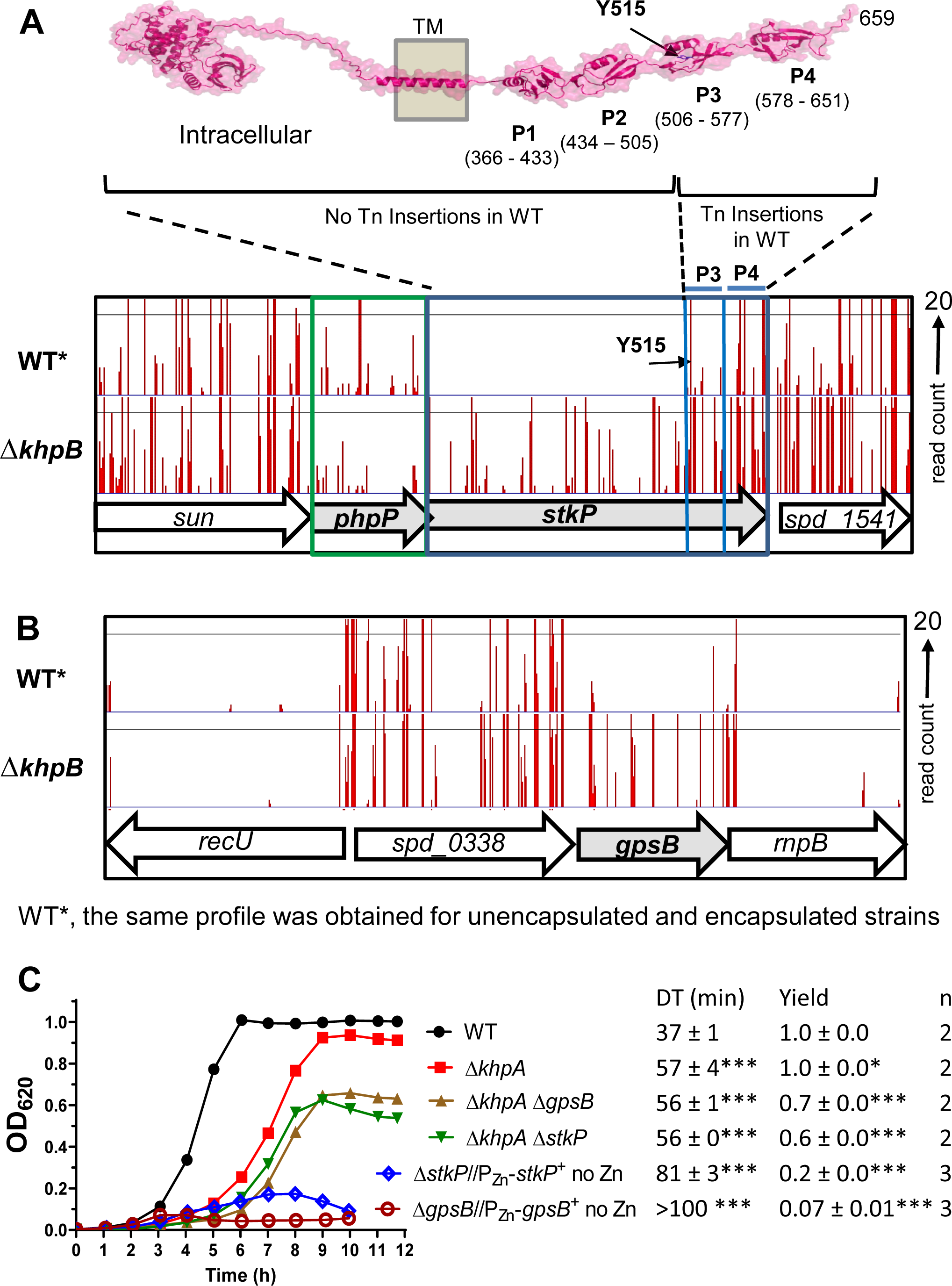
Tn-seq demonstrates the essentiality of StkP(*Spn*) and GpsB(*Spn*) is suppressed by Δ*khpB* in cells growing exponentially in BHI broth in 5% CO_2_. (A) Top: Predicted 3D structure of StkP(*Spn*) generated using the AlphaFold v2.0 webserver. P1, P2, P3 and P4 with indicated amino acid numbers are predicted extracellular PASTA domains. Bottom: Mini-Mariner *Malgellan6* Tn-Seq transposon insertion profile for the genome region covering *sun*, *phpP*, *stkP*, and *spd_1541* in the genomes of the unencapsulated WT parent (D39 Δ*cps rpsL1*, IU1824) or Δ*khpB* (IU10592) strain growing exponentially in BHI broth in 5% CO_2_. The same WT Tn-seq insertion profile was obtained for encapsulated D39 strain IU1781 grown in BHI broth or IU1824 grown in C+Y, pH 6.9 medium in 5% CO_2_ (data not shown). *In vitro* transposition reactions containing purified genomic DNA, *Magellan6* plasmid DNA, and purified MarC9 mariner transposase, transformation, harvesting of transposon-inserted mutants, growth of pooled insertion libraries exponentially in BHI broth or C+Y, pH 6.9 medium, NextSeq 75 high-output sequencing, and analysis were performed as described in *Experimental procedures* based on (Lamanna *et al*., 2022). Sortable data for the profile shown are contained in Appendix A, Tabs C and D. Tn-insertions were recovered for the WT strains in the regions encoding P3 and P4, but not in other regions of *stkP*. The first TA insertion occurs in the WT strain at a TAT (Y515) codon, where the Tn insertion creates a TAA stop codon, while there is no insertion at the upstream TTA (L512) codon, indicating that StkP(M1-L512) is essential for viability. (B) Tn-Seq transposon insertion profiles for the genome region covering *recU*, *spd_0338, gpsB, and rnpB* of in the genomes of the WT parent (D39 Δ*cps rpsL1*, IU1824) or Δ*khpB* (IU10592) strain. (C) Representative growth curves of the WT parent (IU1824), Δ*khpA* (IU9036), Δ*khpA* Δ*gpsB* (IU16196) and Δ*khpA* Δ*stkP* (IU16910) strains. Similar growth results were obtained with Δ*khpB* (IU10592), Δ*khpB* Δ*gpsB* (IU12977), and Δ*khpB* Δ*stkP* (IU16912) strains compared to the strains of Δ*khpA* background. The growths of merodiploid strains Δ*gpsB//*P_Zn_-*gpsB*^+^ (IU16370) and Δ*stkP*::P_c_-*erm//*P_Zn_-*stkP*^+^ (IU16933) grown under conditions that result in depletion of GpsB or StkP were shown for comparison.

We next performed StkP depletion experiments that minimized suppressor accumulation to determine the primary phenotypes caused by lack of StkP. In these experiments, we constructed a merodiploid strain with a non-polar markerless Δ*stkP* at its native site and a zinc-regulatable copy of *stkP*^+^ at an ectopic site (Fig. 9). Depletion of StkP caused cessation of growth followed by a decrease in OD_620_ and substantial increases in the length, aspect ratio, and relative volume, but not width (Fig. 9A-C and S20A-B; no Zn inducer). Markerless Δ*stkP* was nearly completely complemented by an ectopic copy of *stkP^+^* (Fig. 9 and S20; 0.5 mM Zn inducer). Quantitative western blots showed that no StkP was detectable after ≈3-4 h of depletion, and ectopic induction of StkP occurred to ≈50% of the WT level (Fig. S20D). In transformation assays, we used a Δ*stkP*::P_c_-*erm* allele for selection (Table 2). The morphology of markerless Δ*stkP* and Δ*stkP*::P_c_-*erm* cells were slightly different upon StkP deletion (Fig. S20B-C), and unlike markerless Δ*stkP*, Δ*stkP*::P_c_-*erm* was not fully complemented back to WT by ectopic StkP expression (Fig. 20A-D). This lack of full complementation, which was not studied further here, may have been caused by retro-polarity of the insertion construct on expression of upstream *phpP* (phosphatase) or polarity of the constitutive P_c_ promoter on expression of downstream genes, such as *spd_1541* (unknown membrane protein). But together, we conclude that the primary phenotype caused by the absence of StkP is a defect in septum formation in dividing cells, manifested by longer, but not wider, cells compared to WT (Fig. S20B-C).

*murZ*(D280Y) strongly suppressed markerless Δ*stkP* upon StkP depletion (Fig. 9A-C). Notably, cellular MurZ(D280Y) amount was unchanged in the presence or during depletion of StkP (Fig. 9D), when MurZ(D280Y) suppressed the requirement for StkP (Table 2, line 6). In addition, *murZ*(I265V), *murZ*(E259A), and overexpression of MurZ or MurA suppressed Δ*stkP*::P_c_-*erm* in transformation assays (Table 2, lines 4-5 (+Zn inducer) and 7-8; Fig. S3E-J;) and in growth and morphology assays (Fig. S21). In contrast, Δ*clpP*, Δ*clpC*, Δ*clpE,* and Δ*clpL* did not suppress Δ*stkP*::P_c_-*erm* in transformation assays (Table 2, line 11; Fig. S3B; data not shown). We conclude that mutations that suppressed Δ*gpsB* also suppressed Δ*stkP*. Based on transformant colony size, the suppression of Δ*stkP* was generally complete compared to the partial suppression of Δ*gpsB* (Table 2; Fig. S3).

### 2.9 Δ*khpA or* Δ*khpB* suppress Δ*gpsB* by increasing MurZ amount

KhpA and KhpB (EloR/Jag) are KH-domain proteins that form an RNA-binding heterodimer (Stamsas *et al*., 2017, Ulrych *et al*., 2016, Zheng *et al*., 2017, Winther *et al*., 2019). We previously reported that Δ*khpA or* Δ*khpB* suppresses the lethal phenotypes of Δ*pbp2b*, Δ*rodA*, Δ*mreCD*, or Δ*rodZ* elongasome mutants by increasing FtsA expression (Lamanna *et al*., 2022, Zheng *et al*., 2017) (Fig. 11A). We also reported that Δ*khpA* or Δ*khpB* suppresses the lethal phenotypes of Δ*gpsB* (Zheng *et al*., 2017). Tn-seq, transformation, and growth assays confirmed that Δ*khpA* or Δ*khpB* suppressed Δ*gpsB* or Δ*stkP* (Table 2, lines 9-10; Table S5A, lines 30-31; Fig. 10C and S3K-L).

**Figure 11.**
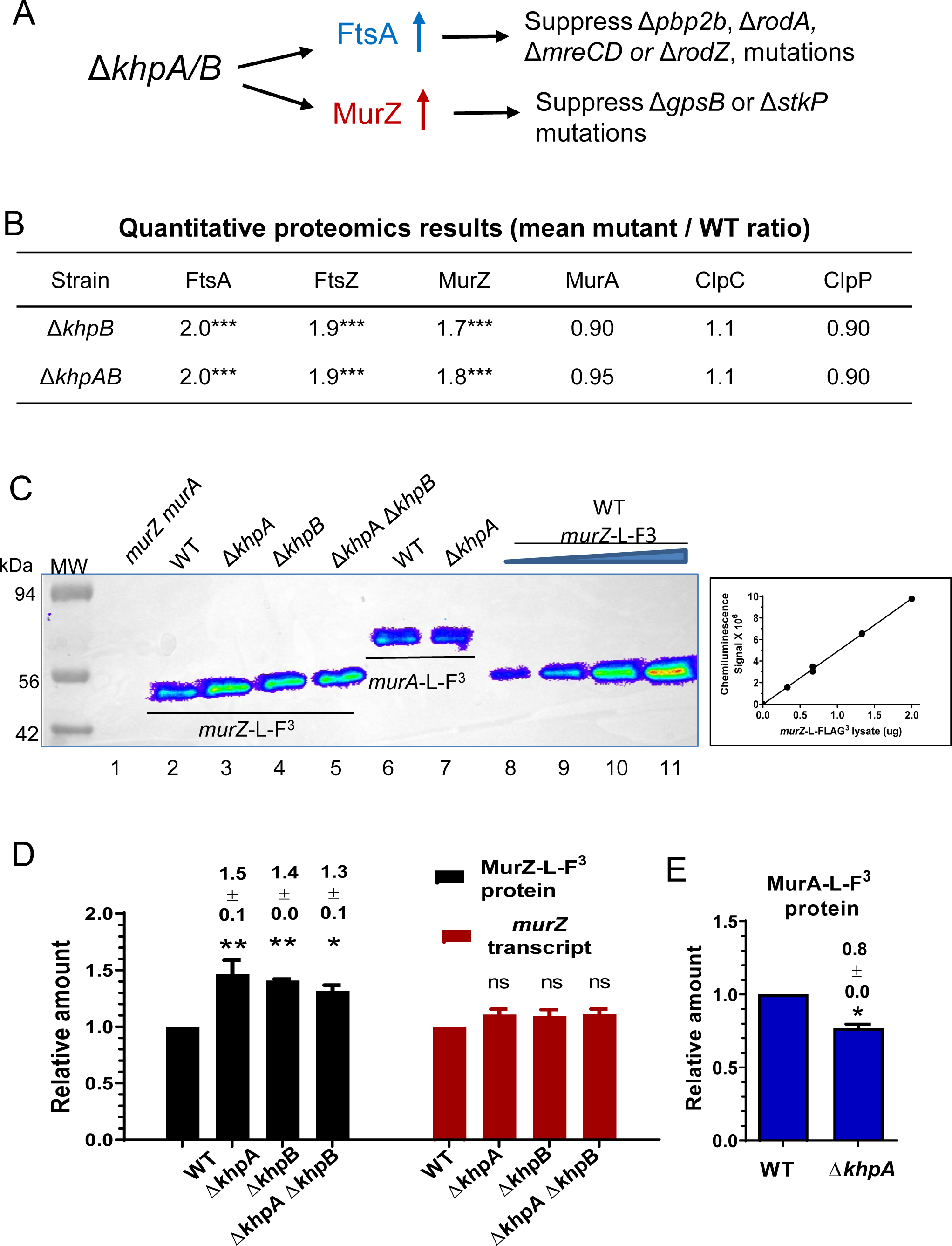
KhpA/B negatively and post-transcriptionally regulates MurZ(*Spn*), but not MurA(*Spn*), cellular amounts. (A) Summary of suppression patterns of Δ*gpsB,* Δ*pbp2b*, Δ*rodA*, and Δ*mreCD* by Δ*khpA/B* mutation. The absence of KhpA and/or KhpB increases the cellular amount of FtsA, which bypasses the requirement for essential PBP2b, RodA, RodZ, and MreCD (Lamanna *et al*., 2022, Zheng *et al*., 2017). The absence of KhpA/B also moderately increases cellular MurZ amount as shown below, which bypasses the requirement for essential GpsB and StkP as described in the text and Fig. 12. (B) Quantitative proteomic results showing relative amounts of FtsA, FtsZ, MurZ, MurA, ClpC, and ClpP in Δ*khpA* Δ*khpB* (IU10596) or Δ*khpB* (IU10592) strains compared to wild-type (IU1824). *** p < 0.001. Proteomics was performed as described in *Experimental procedures*, and data are contained in Appendix A, Tab E. (C) Representative Western blots using anti-FLAG antibody to determine the cellular amounts of MurZ-L-FLAG^3^ and MurA-L-FLAG^3^ in cells growing exponentially in BHI broth. Lane 1, WT parent (IU1824); lane 2, *murZ*-L-F^3^ (IU13502); lane 3, *murZ*-L-F^3^ Δ*khpA* (IU13545); lane 4, *murZ*-L-F^3^ Δ*khpB* (IU14014); lane 5, *murZ*-L-F^3^ Δ*khpA* Δ*khpB* (IU14016); lane 6, *murA*-L-F^3^ (IU14028); lane 7, *murA*-L-F^3^ Δ*khpA* (IU14030). 0.67 µg of total protein from each strain were loaded in lanes 1-7. For lanes 8 to 11, 0.33, 0.67, 1.33, and 2 µg, respectively, of *murZ*-L-FLAG^3^ (IU13502) lysates were loaded to generate the standard curve at right, which showed proportionality between protein amounts and signal intensities over the range of signal intensities obtained. (C) Relative average (± SEM) of cellular amounts of MurZ-L-F^3^ or *murZ* transcripts in mutants compared to WT from 3 independent experiments. P values were obtained relative to WT by one-way ANOVA analysis (Dunnett’s multiple comparison test, GraphPad Prism). * P<0.05; ** P<0.01; ns: not significantly different. (E) Relative average (± SEM) cellular amount of MurA-L-F^3^ protein in a Δ*khpA* mutant compared to WT from 3 independent experiments. P value was obtained relative to WT by one sample t-test (GraphPad Prism). * p<0.05.

Transformation assays strongly implicated MurZ, but not MurA, in Δ*khpA* suppression of Δ*gpsB*. A Δ*khpA* single mutant or a Δ*khpA* Δ*murA* mutant could be transformed by Δ*gpsB*, whereas a Δ*khpA* Δ*murZ* mutant could not be transformed by Δ*gpsB* (Table S5, lines 30, 33-34). Consistent with this result, Δ*murA*, but not Δ*murZ*, could be transformed into a Δ*khpA* Δ*gpsB* suppressed strain (Table S5B, line 5; S5C, line 6). Thus, MurZ is required for Δ*khpA* suppression of Δ*gpsB*.

Consistent with these genetic results, quantitative proteomic analysis detected a ≈1.8-fold (p<0.001) increase in the amount of MurZ, but not MurA, in Δ*khpB* or Δ*khpA* Δ*khpB* mutants compared to WT (Fig. 11B). As controls, the proteomic analysis also confirmed the previous results from quantitative western blotting that FtsA and FtsZ amounts increased ≈2-fold (p< 0.001) in Δ*khpB* and Δ*khpA* Δ*khpB* mutants compared to WT (Fig. 11B; Appendix A, Tab E). Consistent with the proteomic results, quantitative western blotting indicated that MurZ-L-F^3^ amount increased ≈1.4-fold in Δ*khpA*, Δ*khpB*, or Δ*khpA* Δ*khpB* mutants (Fig. 11C-D), whereas MurA-L-F^3^ amount decreased slightly in a Δ*khpA* mutant (Fig. 11E). qRT-PCR showed that the increase in MurZ protein amount was not paralleled by an increase in relative *murZ* transcript amount in Δ*khpA*, Δ*khpB*, or Δ*khpA* Δ*khpB* mutants (Fig. 11D), suggestive of post-transcriptional regulation of MurZ expression. Finally, phosphorylation of KhpB by StkP did not play a role in regulating MurZ expression in cells growing exponentially in BHI broth, since a *khpB*(T89A) phosphoablative mutation did not suppress Δ*gpsB* (Table S5A, line 32). Together, these results suggest that the absence of the KhpAB RNA-binding protein results in a modest (≈2-fold) increase in MurZ(*Spn*), which is sufficient to suppress Δ*gpsB* and Δ*stkP* (Tables 1 and 3, *murZ* duplications; Fig. 5C, 0.1 mM Zn inducer), but not enough to significantly reduce fosfomycin sensitivity (Fig. 6E).

## 3 DISCUSSION

A large majority (25/32) of suppressors of essential Δ*gpsB* or Δ*stkP* in *S. pneumoniae* D39 contained chromosomal duplications that increase the gene dosage of *murZ* or *murA* (Tables 1 and 3). These duplications range from ≈21 to ≈176 genes (Fig. 2 and S1), and suppressors of Δ*gpsB* also suppress Δ*stkP*, and *vice versa* (Table 2). This pattern attests to the extraordinary plasticity of the pneumococcal chromosome, as observed in other studies (Baylay *et al*., 2015, Cowley *et al*., 2018, Johnston *et al*., 2013, Robertson *et al*., 2003, Zheng *et al*., 2017). In this case, the dosage of numerous genes adjoining *murZ* or *murA* is doubled, and in some cases quadrupled, resulting in overexpression of MurZ or MurA and many other essential and nonessential gene products with various functions (Fig. 2; Appendix A, Tabs A1 and A2). Smaller duplications containing *murZ* (3/25) were anchored by direct repeats of degenerate IS elements (Fig. 3 and S1C), and large duplications containing *murA* (2/25) were anchored by direct repeats of tRNA/rRNA gene clusters (Fig. 2 and S1D). Deletions of duplication junctions were not detected in these two classes of duplications, which likely arose by recombination between the long homologous direct repeats of the degenerate IS elements or tRNA/rRNA genes during chromosome replication (Reams & Roth, 2015).

In contrast, the majority (20/25) of large duplications containing *murZ* were anchored by inverted repeats of the redundant *phtD* and *phtB* genes, which encode histidine triad proteins (Fig. 2-3, and S1B). Inverted repeats of redundant copies of genes lead to inversion of the gene order between the repeated genes (Reams & Roth, 2015), which occurred between *phtD* and *phtB* in isolates D39W and D39V of the D39 progenitor strain (Slager *et al*., 2018). However, the results presented here indicate that even though inverted, *phtD* and *phtB* can also anchor large duplications of about ≈150 genes surrounding *murZ*. To do this, *phtD* and *phtB* must contain short direct repeats or other elements that enhance short-junction (SJ) duplication (Reams & Roth, 2015). Indeed, there are small direct repeats of 8 and 9 bp and shorter clusters of directly repeated base pairs within inverted *phtD* and *phtB* that could promote SJ duplication.

Moreover, few large duplications of the *murZ* region (e.g., *sup gpsB-8*) retained an intact duplication junction (Fig. 2A, S1B, and S2B), whereas most of these duplications contained a short deletion of ≈10 genes that removed the junction region (Fig. 2B, S1B, and S2C). Remodeling of chromosome duplications by junction deletion is common and likely arises by a short-junction mechanism involving short, direct repeats or other elements (Reams & Roth, 2015). PCR experiments supported the idea that the deletion/insertion in *sup gpsB-3* arose by a duplication of the *phtD-phtB* region, an inversion within one of the duplicated regions, and last, a short deletion of the duplication junction (Fig. S2C). In Δ*gpsB* mutants, junction deletion is correlated with faster growth compared to long duplication without the deletion (Tables 1; Fig. S4A). Together, these results indicate that the region between inverted *phtD* and *phtB* can readily be duplicated, providing an extra copy of *murZ* that suppresses Δ*gpsB* or Δ*stkP.* This capacity for duplication also raises the potential that the copy of numerous other genes in this region (Appendix A, Tab A1) can be increased in response to other stress conditions.

Besides these *murZ* and *murA* duplications, Δ*gpsB* was suppressed by five separate mutations in *phpP*, which encodes the lone Ser/Thr phosphatase in *S. pneumoniae*, by *murZ*(D280Y), and by *ireB*(Q84(STOP), which truncated the homolog of the IreB(*Efa*) and ReoM(*Lmo*) by four amino acids (Tables 1 and S5; Fig. S4) (Rued *et al*., 2017). Most of the mutations that partly suppressed Δ*gpsB* also almost fully suppressed Δ*stkP* (Table 2; Fig S3). We did not find suppressor mutations that decrease teichoic acid decoration, analogous to those in *L. monocytogenes* (Rismondo *et al*., 2017), because pneumococcal decorations contain GalNAc, which is also an essential component of the teichoic acid core structure (Denapaite *et al*., 2012). As reported previously, *phpP* suppressor mutations restore protein phosphorylation and strongly suppress Δ*gpsB*, whereas the duplication suppressors do not (Rued *et al*., 2017). The new *phpP* and duplication suppressors reported here fit this pattern, and the *murZ*(D280Y) suppressor also did not restore phosphorylation (Fig. S6-S7). The presence of *murZ* or *murA* in all duplication suppressors suggested that overexpression of MurZ or MurA provides a mechanism for phosphorylation-independent suppression of Δ*gpsB* and Δ*stkP*. Consistent with this hypothesis, ectopic overexpression of *murZ* or *murA* was sufficient to partially suppress Δ*gpsB* and strongly suppress Δ*stkP* (Fig. 4 and S21). Likewise, suppression of Δ*gpsB* by Δ*khpAB*, which lacks a regulator that binds to RNA (Zheng *et al*., 2017, Winther *et al*., 2019), depended on MurZ expression and was correlated with MurZ, but not MurA overexpression (Table S5; Fig. 11). In addition, results presented here further confirm that Δ*khpAB* increases the cellular amount of FtsA in exponentially growing pneumococcal cells (Fig. 11A-B), which leads to suppression of peptidoglycan elongasome mutations (Lamanna *et al*., 2022, Zheng *et al*., 2017).

Isolated MurZ(D280Y), constructed MurZ(E259A), and the MurZ(I265V) allele in R6 and Rx1 laboratory strains suppressed Δ*gpsB* and Δ*stkP* (Tables 2 and S5). Notably, the amino acid changes in MurZ(D280Y), MurZ(E259A), and MurZ(I265V) are distant from the catalytic region of MurZ (Fig. 7). MurZ(D280Y) was expressed at the WT MurZ level (Fig. 5C), and Δ*murZ* or Δ*murA* did not change cellular MurA or MurZ amount, respectively (Fig. 5E). These results indicate a third mechanism of suppression, distinct from loss of PhpP activity or MurZ or MurA overexpression. Taken together, these results fit and extend our previous model that GpsB is required for StkP-catalyzed protein phosphorylation, as well as for regulation of peptidoglycan synthesis in exponentially growing cells of *S. pneumoniae* (Rued *et al*., 2017). These new data tie the requirement for StkP-dependent protein phosphorylation to regulation of MurZ and MurA activity, but not amount (Fig. 5C and 9D). According to this updated model, protein phosphorylation drops in the absence of GpsB, which limits MurZ and MurA activity, without changing their amounts. This limitation can be overcome by decreasing PhpP-mediated protein dephosphorylation, by increasing the cellular amounts of MurZ (by ≈2-fold) or MurA (by ≈2-4-fold) by gene duplication or loss of KhpAB, or by altering the interaction of MurZ and MurA with a phosphorylation-dependent regulatory protein. This interaction could potentially be with a phosphorylated positive regulator that activates MurZ and MurA activity or with an unphosphorylated negative regulator that inhibits MurZ and MurA activity (Fig. 12).

**Figure 12.**
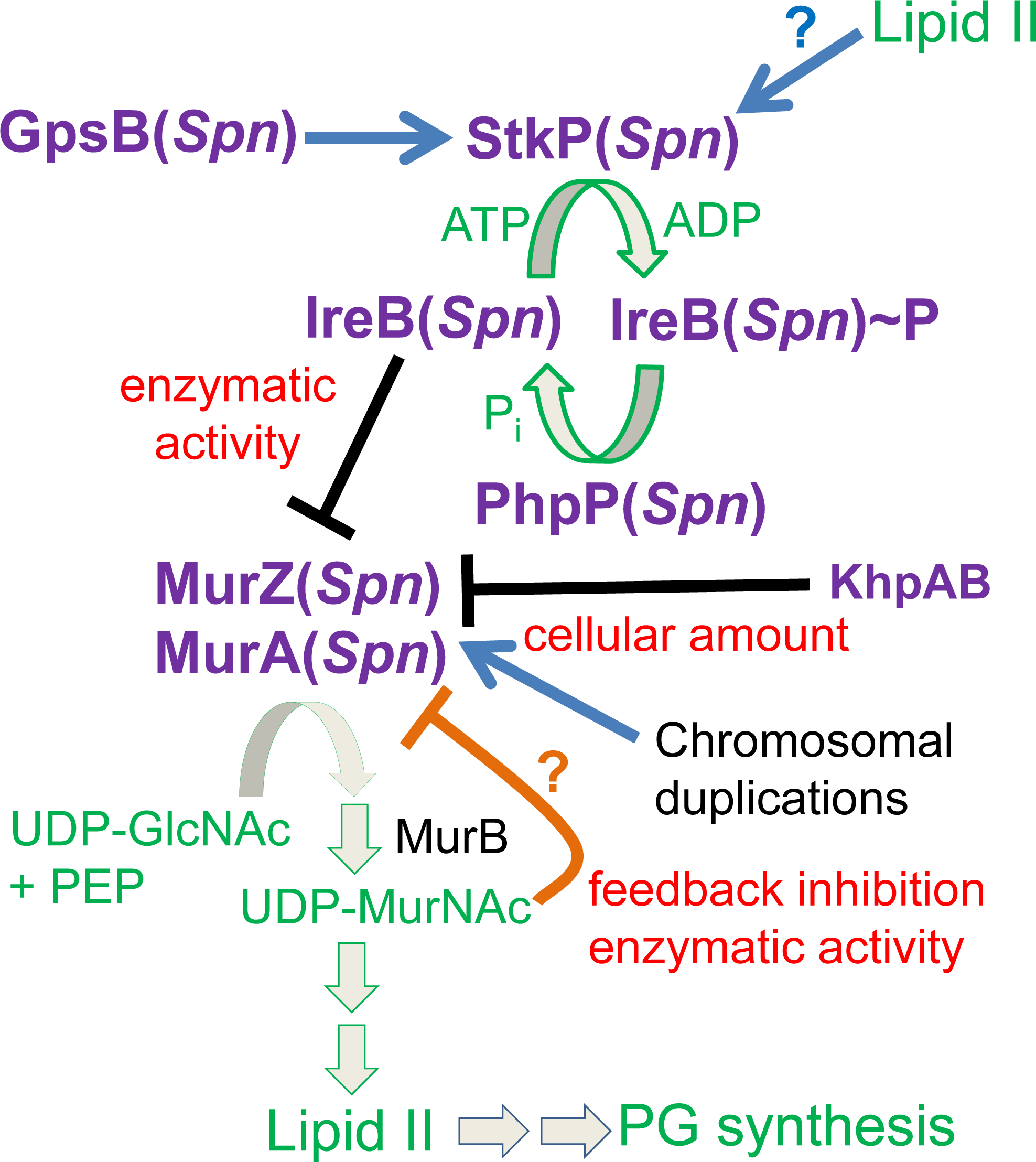
Summary model for regulation of MurZ and MurA enzymatic activities by StkP-mediated phosphorylation in *S. pneumoniae* D39. GpsB(*Spn*) and possibly other ligands, such as Lipid II, stimulate the phosphorylation of a negative regulator of MurZ(*Spn*) and MurA(*Spn*) enzymatic activity, but not their cellular amounts, in the first committed step of Lipid II synthesis for PG synthesis. By genetic criteria presented here, the negative regulator is unphosphorylated IreB(*Spn*). Phosphorylated IreB(*Spn*)∼P does not bind to MurZ(*Spn*) or MurA(*Spn*), resulting in full enzymatic activity in pneumococcal cells growing exponentially in rich media. The absence of GpsB(*Spn*) significantly reduces phosphorylation of IreB(*Spn*) leading to inhibition of MurZ(*Spn*) and MurA(*Spn*) enzymatic activities and no growth. This inhibition can be relieved by inactivation of the cognate PhpP protein phosphatase, which allows residual phosphorylation to IreB(*Spn*)∼P. The absence of the StkP protein kinase and the need for protein phosphorylation in pneumococcal cells growing exponentially in rich media can be suppressed by inactivation or absence of the IreB(*Spn*) negative regulator, by amino-acid changes in a regulatory domain of MurZ(*Spn*), which is enzymatically predominant over MurA(*Spn*), or by overexpression of MurZ(*Spn*) or MurA(*Spn*) in spontaneous chromosomal duplications. Moderate MurZ(*Spn*) overexpression sufficient to suppress the absence of StkP also occurs in the absence of the KhpAB RNA-binding protein, which also negatively regulates FtsA amount. This pathway provides a positive feedback loop, such that cells growing rapidly in rich media produce Lipid II, which may activate StkP(*Spn*) to fully phosphorylate IreB(*Spn*) and maximize MurZ(*Spn*) and MurA(*Spn*) enzymatic activities for the production of even more Lipid II for PG synthesis. Evidence for the direct interaction between unphosphorylated IreB(*Spn*) and MurZ(*Spn*) will be presented elsewhere (Merrin Joseph, unpublished result). Structures predicted by AlphaFold v2.0 also suggest that MurZ(*Spn*) and MurA(*Spn*) enzymatic activity is subject to negative pathway feedback inhibition by binding of UDP-MurNAc (UDP-N-acetylmuramic acid) near the catalytic sites of the enzymes (Mizyed *et al*., 2005, Schonbrunn *et al*., 2000). See text for additional details.

The isolation of the *ireB*(*Spn*)(Q84(STOP)) suppressor implicates IreB(*Spn*) as this regulator, and Δ*ireB* suppressed Δ*gpsB* or Δ*stkP*, consistent with negative regulation (Table 2, line 13). Recent phosphoproteomic analyses show that MurZ or MurA are not phosphorylated by StkP in exponentially growing *S. pneumoniae* D39 cells, whereas IreB(*Spn*) is a prominent phosphorylated protein (Ulrych *et al*., 2021). According to the negative regulation model, the amino acid changes in MurZ(D280Y), MurZ(E259A), and MurZ(I265V) in Domain 1 of MurZ (Fig. 7) weaken an inhibitory interaction between MurZ and unphosphorylated IreB(*Spn*), thereby suppressing the absence of GpsB or StkP (Fig. 12). Details of the interaction between MurZ(*Spn*) and IreB(*Spn*) will be published elsewhere (Merrin Joseph, unpublished results). Moreover, the observation that suppression of Δ*gpsB* is partial, except in *phpP* suppressors, compared to full suppression of Δ*stkP* (Table 2; Fig. 4 and S21) is consistent with GpsB having additional regulatory roles in peptidoglycan synthesis (Cleverley *et al*., 2019, Hammond *et al*., 2022, Minton *et al*., 2022, Rued *et al*., 2017), besides activating StkP. Importantly, these suppression patterns indicate that regulation of MurZ and MurA is the sole essential requirement for protein phosphorylation in unstressed D39 *S. pneumoniae* cells growing exponentially in BHI broth.

Experiments performed parallel to this study and published recently by Wamp and colleagues revealed similar Δ*gpsB* suppression phenotypes in *L. monocytogenes*, with some major differences (Wamp *et al*., 2020). The conditional, temperature-sensitive Δ*gpsB* mutation of *L. monocytogenes* was suppressed by PrpC(*Lmo*) protein phosphatase mutations, by overexpression of MurA(*Lmo*) (the homolog of MurA(*Spn*) (Fig. 1)), and by MurA(*Lmo*)(S262L), which is at a similar position to MurZ(*Spn*)(D280Y) in Domain 1 (Fig. 7) (Wamp *et al*., 2020, Wamp *et al*., 2022). Importantly, several lines of evidence presented here show that the mechanism of MurA homolog regulation is different in *L. monocytogenes* and *S. pneumoniae*. MurA(*Lmo*) stability is regulated by phosphorylation of ReoM(*Lmo*), which is the homolog of IreB(*Efa*) and IreB(*Spn*) (Wamp *et al*., 2020, Wamp *et al*., 2022, Kelliher *et al*., 2021). Unphosphorylated ReoM acts with MurZ(*Lmo*) and ReoY(*Lmo*) as adaptors for degradation of MurA(*Lmo*) by the ClpCP(*Lmo*) protease (Rismondo *et al*., 2017, Wamp *et al*., 2022, Wamp *et al*., 2020). Hence, ReoM(*Lmo*) phosphorylation to ReoM(*Lmo*)∼P makes MurA(*Lmo*) available for peptidoglycan synthesis, including by a special RodA3:PBP3 synthase that contributes to the intrinsic cephalosporin resistance of *L. monocytogenes* (Wamp *et al*., 2022). This mechanism causes Δ*murZ*(*Lmo*) mutants to accumulate MurA(*Lmo*), which is essential in *L. monocytogenes* (Rismondo *et al*., 2017).

In contrast, MurZ(*Spn*) and MurA(*Spn*) share a synthetic lethal relationship (Table S5; Fig. S12) (Du *et al*., 2000), and a Δ*murZ*(*Spn*) or Δ*murA*(*Spn*) mutation does not result in increased cellular amounts of MurA(*Spn*) or MurZ(*Spn*), respectively (Fig. 5E). Several pieces of data in this study demonstrate that MurZ is predominant to MurA in pneumococcal cells. This is the reverse relationship to other Gram-positive bacteria, including *L. monocytogenes*, *E. faecalis*, and *B. subtilis*, where the MurA-family homolog is often essential or predominant to the MurZ-family homolog, which is dispensable and regulatory in the case of MurZ (*Lmo*) (Fig. 1) (Wamp *et al*., 2020). The predominance of MurZ(*Spn*) over MurA(*Spn*) was indicated by the growth and morphology defects of Δ*murZ* mutants in unencapsulated and encapsulated D39 strains grown in BHI broth or C+Y medium (Fig. 6, S10, S13, and S16) and by the increased sensitivity to fosfomycin of Δ*murZ*, but not Δ*murA*, mutants (Fig. 6E). In addition, less overexpression of MurZ (≈2-fold) than MurA (≈4-fold) was required to suppress Δ*gpsB* (Fig. 5), and overexpression of MurZ beyond 2-fold in cells grown in BHI broth led to growth inhibition (Fig. 5-6) that was not observed in C+Y medium (Fig. S10A). This predominance of MurZ compared to MurA in pneumococcal cells is consistent with the greater kinetic efficiency of purified MurZ(*Spn*) compared to MurA(*Spn*) reported earlier by Du and colleagues (Du *et al*., 2000). These combined results show that the relative physiological roles of MurZ(*Spn*) and MurA(*Spn*) are substantially different from those of MurA(*Lmo*) and MurZ(*Lmo*). Likewise, MurZ(*Spn*) and its MurZ-family homolog MurAB(*Bsu*) play very different physiological roles. Remarkably, MurAB(*Bsu*) was discovered to be required for efficient spore engulfment during sporulation of *B. subtilis* (Chan *et al*., 2022). *S. pneumoniae* does not sporulate.

Other evidence strongly argues for a different mechanism of MurA and MurZ regulation by ReoM/IreB homologs in *S. pneumoniae* compared to *L. monocytogenes*, *B. subtilis*, and *E. faecalis. S. pneumoniae* lacks homologs of the ReoY accessory factor required for MurA degradation by ClpCP in *L. monocytogenes* (Wamp *et al*., 2020, Wamp *et al*., 2022). In addition, MurA is essential in *L. monocytogenes* (Rismondo *et al*., 2017) and likely supplies Lipid II precursor to an additional RodA3:PBPB3 synthase that imparts resistance to cephalosporins (Wamp *et al*., 2022). Homologs of ReoY and RodA3:PBPB3 are also present in *E. faecalis* (Wamp *et al*., 2022), while *S. pneumoniae* lacks homologs of these proteins. Most importantly, suppression of Δ*gpsB*(*Spn*) is not dependent on ClpP(*Spn*) (Table 2) or ClpP-associated ATPases, including ClpC(*Spn*) (Table S5), and MurZ(*Spn*) and MurA(*Spn*) cellular amounts remain unchanged in a Δ*clpP*, Δ*clpC*, Δ*clpE*, or Δ*clpL* mutants (Fig. 5E and 8). Together, these results support a model in which protein phosphorylation does not change the amounts of MurZ(*Spn*) or MurA(*Spn*), but rather, regulates their enzymatic activities. Interestingly, amino-acid changes in Domain I of MurZ(*Spn*) (I265V, D280Y, and E259A) and MurA(*Lmo*) (S262L) likely suppress Δ*gpsB* by decreasing interactions with unphosphorylated IreB(*Spn*) and ReoM(*Lmo*), respectively (Wamp *et al*., 2022, Wamp *et al*., 2020). However, amino-acid changes in Domain II of MurZ(*Spn*) (E190A, E192A, and E195A) did not suppress Δ*gpsB* (Table S5; Fig. 7), whereas MurA(*Lmo*)(N197D) did (Wamp *et al*., 2022), consistent with different mechanisms in *S. pneumoniae* and *L. monocytogenes*.

This paper also demonstrates that the StkP Ser/Thr protein kinase is essential, except for its two distal PASTA domains (P3 and P4), in *S. pneumoniae* D39 progenitor strains growing exponentially in BHI broth or C+Y, pH 6.9 medium (Fig. 10). PASTA domains have been shown to bind Lipid II in Ser/Thr protein kinases of other Gram-positive bacteria (Hardt *et al*., 2017, Kaur *et al*., 2019, Sun *et al*., 2023). We also show that the primary phenotype of StkP depletion is the formation of longer, but not wider, non-growing cells (Fig. 9), indicative of a septation defect that may be triggered by decreased cellular Lipid II amount (Fig. 12). Essentiality of *stkP*(*Spn*) has been controversial for two reasons addressed here. First, Δ*stkP* mutants readily accumulate gene duplications of *murZ* or *murA* that compensate for the lack of IreB phosphorylation (Table 3), as do other spontaneous mutations in *murZ* or *ireB* (Table 2). Consequently, Δ*stkP* mutants in D39 strains form unusual-looking, faint colonies with tiny centers containing suppressor mutants (Fig. S3) (Rued *et al*., 2017). Along this line, it was previously noted that the morphology of D39 Δ*stkP* mutant cells seemed to change upon passage (Beilharz *et al*., 2012). Chromosomal duplications do not result in bp changes and are not indicated in standard whole-genome sequencing reports. The recent conclusion that Δ*stkP* is not essential in D39 likely stems from a duplication of the *phtD*-*phtB* region (Fig. 2), as indicated by increased transcript amounts in RNA-seq (Kant *et al*., 2023). Suppression of Δ*stkP* by chromosomal duplications complicates interpretations of mutant phenotypes.

Second, the R6- and Rx1-derived laboratory strains in which previous experiments were performed carry a *murZ*(I265V) mutation (Lanie *et al*., 2007) that suppresses the requirement for Δ*gpsB* or Δ*stkP* (Table 2; Fig. S18). *murZ*(I265V) changes an amino acid in domain I of MurZ, near the *murZ*(D280Y) and *murZ*(E259A) suppressors (Fig. 7). Thus, Δ*gpsB* and Δ*stkP* appeared not to be essential in studies using laboratory strains that contain *murZ*(I265V). Compared to the D39 progenitor strain, these R6- and Rx1-derived laboratory strains contain dozens of additional mutations, besides *murZ*(I265V) (Cuppone *et al*., 2021, Lanie *et al*., 2007, Santoro *et al*., 2019). Mutational variations may account for why the level of Δ*gpsB* and Δ*stkP* suppression by *murZ*(I265V) varies in different R6 and Rx1 isolates (Beilharz *et al*., 2012, Rued *et al*., 2017).

Overall, this study reveals two different evolutionary strategies for the regulation of MurA function in different Gram-positive bacteria. In all cases, MurA function is linked to protein phosphorylation in exponentially growing cells (Fig. 12) (Wamp *et al*., 2020, Wamp *et al*., 2022). Recent biochemical studies by Minton and colleagues and by Doubravová and colleagues demonstrate that purified GpsB directly stimulates the activity of the Ser/Thr protein kinases from *E. faecalis* and *S. pneumoniae*, respectively (Doubravová, unpublished result) (Minton *et al*., 2022). In *L. monocytogenes*, and likely *E. faecalis*, unphosphorylated ReoM/IreB interacts with the MurA-family enzyme, along with adaptors MurZ and ReoY to present MurA to ClpCP protease for degradation, thereby inhibiting peptidoglycan synthesis and growth (Wamp *et al*., 2022, Wamp *et al*., 2020). In *S. pneumoniae,* reduced phosphorylation, likely of IreB(*Spn*) (Merrin Joseph, unpublished results), does not change the cellular amounts of MurZ and MurA, but decreases their enzymatic activity. Thus, binding between MurA homologs and unphosphorylated ReoM/IreB appears to be evolutionary conserved, but *S. pneumoniae* did not evolve or retain the adaptor/ClpCP degradation pathway of MurA regulation present in *L. monocytogenes* and *E. faecalis* (Wamp *et al*., 2022, Wamp *et al*., 2020). It remains to be determined how the relative function and regulation of MurZ(*Spn*) and MurA(*Spn*) change in *S. pneumoniae* cells subjected to stress conditions, which alters protein phosphorylation by StkP (Ulrych *et al*., 2021), besides during the exponential growth conditions used here.

Finally, there is precedent for phosphorylated proteins modulating MurA activity directly. In *Mycobacterium tuberculosis,* MurA(*Mtb*) is inactive until it binds to phosphorylated CwlM(*Mtb*) regulator, which increases MurA(*Mtb*) enzymatic activity by 20-40-fold (Boutte *et al*., 2016). Homologs of CwlM(*Mtb*) are absent from *S. pneumoniae*, *monocytogenes*, *E. faecalis*, and *B. subtilis* (data not shown) (Boutte *et al*., 2016). A MurA(*Mtb*)(S368P) amino-acid change suppressed the lethal phenotype of a phosphoablative change to CwlM(*Mtb*)(T374A), which is unable to be phosphorylated and activate WT MurA(*Mtb*) (Boutte *et al*., 2016). Notably, MurA(*Mtb*)(S368P) is on the opposite side of Domain I of MurA near the active site region (Fig. 7), compared to amino-acid changes in MurA homologs, such as MurZ(*Spn*)(D280Y) and MurA(*Lmo*)(S262L), that likely decrease binding to unphosphorylated homologs of IreB/ReoM. The separate location of amino-acid changes that result in suppression is consistent with the different mechanisms of positive activation of MurA(*Mtb*) activity by phosphorylated CwlM(*Mtb*) (Boutte *et al*., 2016) compared to negative inhibition of MurZ(*Spn*) activity by an unphosphorylated regulator, such as IreB(*Spn*).

## 4 Experimental procedures

### 4.1 Bacterial strains and growth conditions

Strains used in this study are listed in Table S1. Strains were derived from unencapsulated strains IU1824 (D39 Δ*cps rpsL1*) and IU1945 (D39 Δ*cps*), which were derived from the encapsulated serotype-2 D39W progenitor strain IU1690 (Lanie *et al*., 2007, Slager *et al*., 2018). Other strains were derived from unencapsulated laboratory strain R6 (Hoskins *et al*., 2001). A small number of drift mutations that have accumulated in IU1824 and IU1945 compared to IU1690 were determined by whole-genome sequencing and are listed in Appendix A, Tab B. Strains containing antibiotic markers were constructed by transformation of CSP1-induced competent pneumococcal cells with linear DNA amplicons synthesized by overlapping fusion PCR (Ramos-Montanez *et al*., 2008, Tsui *et al*., 2016, Tsui *et al*., 2014). Strains containing markerless alleles in native chromosomal loci were constructed using allele replacement via the P_c_-[*kan*-*rpsL*^+^] (Janus cassette) (Sung *et al*., 2001). Primers used to synthesize different amplicons are listed in Table S1. Bacteria were grown on plates containing trypticase soy agar II (modified; Becton-Dickinson), and 5% (vol/vol) defibrinated sheep blood (TSAII-BA). Plates were incubated at 37°C in an atmosphere of 5% CO_2_. TSAII-BA plates for selections contained antibiotics at concentrations described previously (Tsui *et al*., 2016, Tsui *et al*., 2014). Bacteria were cultured statically in Becton-Dickinson brain heart infusion (BHI) broth at 37°C in an atmosphere of 5% CO_2_, and growth was monitored by OD_620_ as described before (Tsui *et al*., 2016). Mutant constructs were confirmed by PCR and DNA sequencing of chromosomal regions corresponding to the amplicon region used for transformation. Ectopic expression of various genes was achieved with a P_Zn_ zinc-inducible promoter in the ectopic *bgaA* site. 0.2 to 0.5 mM (Zn^2+^/(1/10)Mn^2+^) was added to TSAII-BA plates or BHI broth for inducing conditions. Mn^2+^ was added with Zn^2+^ to prevent zinc toxicity (Jacobsen *et al*., 2011, Tsui *et al*., 2016, Rued *et al*., 2017).

In all experiments, cells were inoculated from frozen glycerol stocks into BHI broth, serially diluted, and incubated 12–15 h statically at 37°C in an atmosphere of 5% CO_2_. Parallel cultures were set up for each strain and condition for generation of growth curves and collections of samples for Western blot or microscopy. For culturing merodiploid strains that require Zn^2+^ for overexpressing *murZ, murA, gpsB*, or *stkP* from a Zn-dependent promoter (P_Zn_) placed at an ectopic *bgaA* site (Tsui *et al*., 2016), 0.2 to 0.5 mM (Zn^2+^/(1/10)Mn^2+^) were added to BHI broth in the overnight cultures. BHI was supplemented with 0.2 mM (Zn^2+^/(1/10)Mn^2+^) for overnight growth of IU15860 (Δ*gpsB murZ*^+^//P_Zn_-*murZ*^+^) and IU16897 (Δ*stkP murZ*^+^//P_Zn_-*murZ*^+^), with 0.5 mM (Zn^2+^/(1/10)Mn^2+^) for growth of IU15862 (Δ*gpsB murA*^+^//P_Zn_-*murA*^+^) and IU16933 (Δ*stkP*//P_Zn_-*stkP*^+^), and with 0.4 mM (Zn^2+^/(1/10)Mn^2+^) for growth of IU16915 (Δ*stkP murA*^+^//P_Zn_-*murA*^+^). The next day, cultures at OD_620_ ≈0.1–0.4 were diluted to OD_620_ ≈0.003 in BHI broth with no additional (Zn^2+^/(1/10)Mn^2+^) or the amounts of (Zn^2+^/(1/10)Mn^2+^) indicated for each experiment. Doubling time determination was performed by first examining the growth curves on a log scale to determine the time points when growth was in exponential phase. Doubling times were determined with GraphPad Prism exponential growth equation using only data points that exhibit exponential growth. Maximal growth yields were determined by the highest OD_620_ values obtained within 9 h of growth. Doubling times and maximal growth yields were compared to WT strain with one-way ANOVA analysis (GraphPad Prism, Dunnett’s test). Cultures were sampled for microscopy or western analysis at OD_620_ ≈0.1–0.2 (early to mid-exponential phase).

### 4.2 Transformation assays

Transformations were performed as previously described (Rued *et al*., 2017, Tsui *et al*., 2016). Δ*gpsB*<>*aad9,* Δ*murZ*::P_c_-*erm*, Δ*murA*::P_c_-*erm,* Δ*stkP*::P_c_-*erm* amplicons, and positive control Δ*pbp1b*::P_c_-*aad9* or Δ*pbp1b*::P_c_-*erm* amplicon were synthesized by PCR using the primers and templates listed in Table S1, and contain ≈1 kb of flanking chromosomal DNA. All transformation experiments were performed with no added DNA as the negative control, and with respective Δ*pbp1b* amplicons containing the same antibiotic selections as the positive control for competence efficiency and colony size comparison. The volumes of transformation mixture plated (50 to 300 µL) were adjusted to provide ≈150 to 300 colonies with the Δ*pbp1b* amplicons. Transformations with control Δ*pbp1b* amplicons with unencapsulated or encapsulated strains typically yielded >500, or ≈300 colonies per 1 mL of transformation mixture. Transformants were confirmed by PCR reactions. Each transformation experiment was performed 2 or more times. The sizes of colonies indicated in Table 2 were relative to colonies transformed with the same recipient strain with a control Δ*pbp1b* amplicon. For transformations in 0.2 mM or 0.4 mM (Zn^2+^/(1/10)Mn^2+^), ZnCl_2_ and MnSO_4_ stock solutions were added to transformation mixes and soft agar for plating and spread onto blood plates containing (Zn^2+^/(1/10)Mn^2+^) to induce gene expression under control of the P_Zn_ zinc-inducible promoter in the ectopic *bgaA* site (Jacobsen *et al*., 2011, Rued *et al*., 2017). For Δ*stkP* transformation experiments, a volume (≈100 to 150 µL) of transformation mix so that ≈100 colonies appeared on each plate. We ensured that there were similar numbers of the Δ*stkP* and positive control transformants, and that all the colonies appeared similar on each plate. Pictures of colony morphologies of strains transformed with Δ*stkP*::P_c_-*erm* and the control Δ*pbp1b*::P_c_-*erm* amplicon were taken from transformation plates after 20 h of incubation at 37°C post-transformation, with illumination source from under the plates.

### 4.3 Whole-genome DNA sequencing

Whole-genome sequencing was used to identify suppressor mutations and to verify the genomes of constructed mutants. Strains listed in Table 1 containing suppressor mutations that allowed growth of a Δ*gpsB* mutant were isolated as described previously (Rued *et al*., 2017, Tsui *et al*., 2016). Genomic DNA preparation, DNA library construction, Illumina MiSeq or NextSeq DNA sequencing, and bioinformatics analyses were performed as described previously (Rued *et al*., 2017, Tsui *et al*., 2016). Reads were adapter trimmed and quality filtered using Trimmomatic ver. 0.38 (http://www.usadellab.org/cms/?page=trimmomatic), with the cutoff threshold for average base quality score set at 20 over a window of 3 bases. Reads shorter than 20 bases post-trimming were excluded. More than 95% of the sequenced reads passed quality filters. Cleaned reads were mapped to *Streptococcus pneumoniae* D39 genome sequence (CP000410.2) using bowtie2 version 2.3.2. More than 97.5% of the cleaned reads mapped to the genome. Variants in the libraries with each group against the D39 reference were called and compared using Breseq version 0.35.1 (Deatherage & Barrick, 2014) https://barricklab.org/twiki/bin/view/Lab/ToolsBacterialGenomeResequencing). Several spontaneous drift mutations (Appendix A, Tab B) that do not cause detectable phenotypes in the IU1824 (D39 Δ*cps rpsL1*) and IU1945 (D39 Δ*cps*) unencapsulated parent strains (Table S1) (Lanie *et al*., 2007) were eliminated manually as new variants. The number of reads of each base was also mapped to the D39 reference genome by using the JBrowse program (Skinner *et al*., 2009, Westesson *et al*., 2013) to detect regions containing chromosomal duplications or large deletions (Rued *et al*., 2017).

### 4.4 Cell length and width measurements

Cell lengths and widths of strain growing exponentially in BHI broth were measured as previously described (Tsui *et al*., 2016). For *gpsB*^+^ strains, only ovoid-shape predivisional cells were measured. For analysis that include Δ*gpsB* strain, all separated cells, including cells that were constricted or narrower at midcell, were measured. Unless indicated in the figure legends, more than 100 cells from at least 2 independent experiments were measured and plotted with box and whiskers plot (5 to 95 percentile whiskers). P values were obtained by one-way ANOVA analysis by using the nonparametric Kruskal-Wallis test in GraphPad Prism program.

### 4.5 RNA preparation and qRT-PCR

RNA preparation, qRT-PCR were performed as previously described (Tsui *et al*., 2016, Zheng *et al*., 2017). Primers used for qRT-PCR are listed in Table S1.

### 4.6 Quantitative western blotting

Cell lysate preparations using SEDS lysis buffer (0.1% deoxycholate (vol/vol), 150 mM NaCl, 0.2% SDS (vol/vol), 15 mM EDTA pH 8.0) and western blotting was performed as previously described (Cleverley *et al*., 2019, Lamanna *et al*., 2022). Briefly, bacteria were grown exponentially in 5 ml BHI broth to an OD_620_ ≈ 0.15–0.2. Frozen pellets collected from 1.8 mL of cultures at OD_620_ ≈0.16 were suspended in 80 μL of SEDS lysis buffer. The volume of SEDS buffer was adjusted proportional to the OD_620_ values. Protein assays were performed with the lysates and the μg amounts of protein lysates loaded on each lane were listed in the figure legends of each blot. The sources of antibodies used for western blotting are as below. Primary antibodies used are anti-HaloTag monoclonal antibody (Promega, G921A, 1:1000), and polyclonal rabbit antibodies: anti-FLAG (Sigma, F7425, 1:2000); anti-HA (Invitrogen, 71–5500, 1:1000); α-pThr antibody (Cell Signaling, #9381) (Rued *et al*., 2017) and anti-StkP (1:10,000) (Beilharz *et al*., 2012, Rued *et al*., 2017). Secondary antibodies used were anti-mouse IgG conjugated to horseradish peroxidase (Invitrogen, SZ-100, 1:3300), anti-rabbit IgG conjugated to horseradish peroxidase (GE healthcare NA93AV, 1:10,000), or Licor IR Dye800 CW goat anti-rabbit (926–32,211, 1:14,000). Chemiluminescence signals obtained with secondary HRP-conjugated antibodies were detected using IVIS imaging system (Fig. 5, 8, 11, S6, S7, S8, S15), or Azure biosystem 600 (Fig. S19C) as described previously (Lamanna *et al*., 2022). IR signals obtained with Licor IR Dye800 CW secondary antibody was detected with Azure biosystem 600 (Fig. 9, S19A, S19B S20).

The relative expression levels of MurZ and MurA were measured with MurZ-L-FLAG^3^ or MurA-L-FLAG^3^ expressed from their native chromosomal locus (Fig. 5, 8, 9, 11, S8, S15). To ensure linearity of western signal values vs protein amounts, a range of protein samples of IU13502 (*murZ*-L-FLAG^3^) or IU14028 (*murA*-L-FLAG^3^) were loaded on the same gel as the experimental samples to provide a standard curve of µg protein amounts versus signal intensities. These plots were performed for each western quantitation experiment (see Fig. 5, 8, 9, 11, S8, S15, S20), and were used to calculate the relative protein amounts in each sample lane by extrapolation. To avoid intensity values beyond the linear range, lower µg amounts of proteins from the induced *murZ*-L-FLAG^3^ or *murA*-L-FLAG^3^ overexpression strains (IU13772 or IU15983, respectively) were loaded per lane in order for the intensity signals of these samples to stay within the linear range (Fig. S8). For Fig. 9D, 6 µL (≈2 µg) of protein samples were loaded in each sample lane for comparison. A standard curve was generated by loading 3, 6, 9 or 12 µL of IU13502 (*murZ*-L-FLAG^3^) samples (lanes not shown). For Fig. S20D, 10 µL (≈3 µg) of protein samples were loaded in each sample lane for comparison. A standard curve was generated by loading 5, 7.5, 10 or 15 µL of WT samples. Signal intensities obtained with the anti-Flag or anti-StkP antibody were normalized with total protein stain in each lane using Totalstain Q-NC reagent from Azure biosystems in these two experiments.

### 4.7 2D-immunofluorescence microscopy (2D-IFM)

2D-IFM was performed to examine the localization pattern of MurZ and MurA as described in (Land *et al*., 2013) using a primary anti-FLAG antibody (Sigma, F7425, 1:100 dilution) and secondary Alexa Fluor 488 goat anti-rabbit IgG (Life Technologies, Z1034, 1:100 dilution) with strains IU13502 (*murZ*-L-FLAG^3^) and IU14028 (*murA*-L-FLAG^3^). Nucleoid DNA was labeled with mounting media SlowFade gold antifade reagent with DAPI (Life Technologies, S36936).

### 4.8 Antibiotic disk-diffusion assay

Strains were inoculated in 3 mL BHI broth from frozen glycerol stocks and grown at 37°C until early exponential phase (OD_620_ ≈0.09-0.15). Cells were then diluted to OD_620_ ≈0.009 in 1 mL BHI, and 50 µL of diluted culture was then mixed into 3 mL nutrient-broth soft agar [0.8% (w/v) nutrient broth and 0.7% (w/v) Bacto Agar (Difco)] and poured onto TSAII-BA plates. After 15 min, antibiotic Sensi-Disc^TM^ (Becton Dickinson Pty Ltd., Fosfomycin; cat# 231709, Cefotaxime; cat# 231606, Tetracycline; cat# 230998, penicillin; cat# 230918, Cefoperazone; cat# 231612 (data not shown)), were placed at the middle of plates that were incubated 37°C for 16 h prior to measurement of zone of inhibition. Images of plates were taken using the Azure imaging system, and diameters of the zones of inhibition were measured using the Java program AntibiogramJ (Alonso *et al*., 2017).

### 4.9 3D structure and residue alignment

The MurZ structure from *S. pneumoniae* D39 was generated using AlphaFold v2.0 (Jumper *et al*., 2021) on the Carbonate Research supercomputer at Indiana University, and images were generated using PyMOL (Schrödinger, LLC). For amino acid sequence comparisons, amino acid sequences of MurZ and MurA from *S. pneumoniae* D39 and MurA from *E. coli* K12 were obtained from the protein PubMed database (https://www.ncbi.nlm.nih.gov/protein/) and aligned using the Clustal Omega web server to determine locations of the catalytic Cys, and other residues demonstrated to be important for MurA function in other bacterial species.

### 4.10 Proteomic analysis

Triplicate 30-mL cultures of wild-type (IU1824), Δ*khpA* Δ*khpB* (IU10596) and Δ*khpB* (IU10592) strains were grown in BHI broth to an OD_620_ ≈0.1-0.15. Cultures were then collected by centrifugation at 16,000 x *g* for 5 min at 4°C. Cell pellets were resuspended in 1 mL of cold PBS, centrifuged at 16,100 x *g* for 5 min at 4°C, and resuspended in 1 mL of lysis buffer (8 M Urea, 100 mM ammonium bicarbonate (pH 7.8), 0.5 % sodium deoxycholate, and protease inhibitor (1 mini tablet (Pierce™ A32955) per 10 mL). Resuspended cells were transferred to lysing matrix B tubes and lysed in a FastPrep homogenizer (MP Biomedicals) at a rate of 6 m/s for 40 s three times. Samples were centrifuged at 16,100 x *g* for 5 min at 4°C. 700 µL supernatant was transferred to a new 1.5-mL tube and concentrated using Amicon Ultra 1 mL 10K membrane filters (Millipore, catalog number: UFC501096) to ≈40 µL by centrifuging at room temperature at 14,000 x *g* for ≈45 min. Samples were washed in the spin filter by adding 200 µL of wash buffer (8 M Urea, 100 mM ammonium bicarbonate (pH 7.8), 0.1% sodium deoxycholate) in the spin filter and centrifuged at room temperature at 14,000 x *g* for ≈1 h until ≈40 µL remains in the column. 3 x volumes (≈120 µL) of 100 mM ammonium bicarbonate were added to the samples to produce a final urea concentration of 2M. Samples were concentrated by centrifugation in the spin column to ≈40 µL, which were transferred to fresh 1.5 mL microfuge tubes. Spin filters were rinsed twice with 200 μL of 25 mM ammonium bicarbonate and added to the sample tubes. The protein concentration was quantified by a Bio-Rad DC protein assay (catalog number: 5000111) using BSA in 0.2M urea and 25 mM ammonium bicarbonate (pH 7.8) as standards. Typical protein yields were 270 to 430 μg per 30-mL culture. 100 μg of protein were dried in SpeedVac concentrator for ≈15 h followed by in-solution protein digestion.

Samples were denatured in 8 M urea, 100 mM ammonium bicarbonate solution, then incubated for 45 min at 56°C with 10 mM dithiothreitol (DTT) to reduce cysteine residues. The free cysteine residue side chains were then alkylated with 40 mM iodoacetamide for 1 h in the dark at room temperature. The solution was diluted to 1 M urea and 1:100 (wt/wt) ratio of trypsin was added and the samples were digested at 37°C for 16 h. Peptides were desalted by Zip-tip.

LC-MS/MS Analysis was performed by injection of peptides into an Easy-nLC HPLC system coupled to an Orbitrap Fusion Lumos mass spectrometer (Thermo Scientific, Bremen, Germany). Peptide samples were loaded onto a 75 µm x 2 cm Acclaim PepMap 100 C18 trap column (Thermo Scientific) in 0.1% formic acid. The peptides were separated using a 75 µm x 25 cm Acclaim PepMap C18 analytical column using an acetonitrile-based gradient (Solvent A: 0% acetonitrile, 0.1% formic acid; Solvent B: 80% acetonitrile, 0.1% formic acid) at a flow rate of 300 nL/min. Peptides were separated using a 120 min gradient. The initial solvent was 2% B. This was ramped to 4% B over 30 sec. The gradient then ramped up to 32% B over 114 min, then up to 100% B over 30 sec and held there for the remaining five min. The electrospray ionization was carried out with a nanoESI source at a 260°C capillary temperature and 1.8 kV spray voltage. The mass spectrometer was operated in data-dependent acquisition mode with mass range 400 to 1600 m/z. The precursor ions were selected for tandem mass (MS/MS) analysis in the Orbitrap with 3 sec cycle time using HCD at 35% collision energy. Intensity threshold was set at 1e4. The dynamic exclusion was set with a repeat count of 1 and exclusion duration of 30 s.

The resulting data were searched against a *Streptococcus pneumoniae* D39 database (Uniprot UP000001452 with 1,915 entries, downloaded on 02/2020) using MaxQuant version 1.6. Carbamidomethylation of cysteine residues was set as a fixed modification. Protein N-terminal acetylation and oxidation of methionine were set as variable modifications. Trypsin digestion specificity with two missed cleavage was allowed. The first and main search peptide tolerances were set to 20 and 4.5 ppm, respectively.

Perseus Version 2.0.3.0 was used for statistical analysis of the data (Aguilan *et al*., 2020, Turapov *et al*., 2018). The fractional abundance of each protein is calculated relative to the total lysate (protein area/total lysate area) and used to estimate the fold-change. Statistical data analyzation was done in Perseus by applying the following workflow: (a) log2 data transformation and imputation based on normal distribution to eliminate division by zero, (b) removing proteins only identified in one of replicates, (c) calculating the mean of replicates, and (d) performing a t-test to determine proteins that were statistically different between wild-type and mutant. Average values reported in this study were calculated based on 5 replicates of wild-type and 3 replicates of mutant strains. Pairwise Pearson correlation coefficients among replicates of the same strain were ≥ 0.986 for all three strains. Data from the proteomic analysis is contained in Appendix A, Tab E.

### 4.11 Tn-seq transposon library generation and insertion sequencing

Tn-seq transposon library generation and insertion sequencing of WT D39 Δ*cps rpsL1* (IU1824) and isogenic Δ*khpB* (IU10592) are as reported in (Lamanna *et al*., 2022). Tn-seq primary data for the region between *sun* (*spd_1544*) and *spd_1541,* which are upstream and downstream of *phpP*(*spd_1543*)*-stkP*(*spd_1542*), respectively, are contained in Appendix A, Tabs C and D, including run summaries, number of reads per TA site in each gene, and count ratios for each gene in the indicated mutants compared with WT. P values for comparisons of the number of reads per TA site in each gene were calculated by the nonparametric Mann-Whitney test using GraphPad Prism (9.2.0).

## AUTHOR CONTRIBUTIONS THAT MET ICMJE CRITERIA FOR AUTHORSHIP

HCTT, MJ, JJZ, AJP, BER, and MEW contributed to the conception or design of this study. HCTT, MJ, JJZ, AJP, IM, and BER contributed to the acquisition and analysis of the data. HCTT, MJ, PB, LD, OM, and MEW contributed further analysis and interpretation of the data. HCTT, MJ, PB, LD, OM, and MEW contributed to the writing of the manuscript with input from the other authors.

## ACKNOWLEDGMENTS

We thank Ziyun April Ye and Bobby Walker for technical assistance in strain construction, Jonathon Trinidad and Aleš Ulrych for advice on interpretation of proteomic data, Doug Rusch and Ram Podicheti for bioinformatic assistance of whole-genome sequences, Ulf Gerth and Chris Kristich for polyclonal antibodies against MurAA(*Bsu*) and MurAA(*Efa*), respectively, and Kevin Bruce and other members of the Winkler lab for discussions about this work. This work was supported by NIH Grant R35GM131767 (to MEW), grants 18-07748S (to LD) and 19-03269S (to PB) from the Czech Science Foundation, grant LTAUSA18112 (to LD and PB) from the Ministry of Education, Youth, and Sports of the Czech Republic, and by institutional research funds from the CIBIO Department of the University of Trento (to OM). Work done on the Carbonate Research supercomputer was supported in part by the Lilly Endowment, Inc., through its support of the Indiana University Pervasive Technology Institute.

## ETHICS STATEMENT

This work did not include animal or human experimental subjects requiring formal approval or consent. Antibodies used in this study are available commercially, were published previously, or were prepared by companies approved by the Indiana University Bloomington Institutional Animal Care and Use Committee.

## DATA AVAILABILITY STATEMENT

All data that support the findings of this study are reported with indicated statistical analyses and numbers of biological repeats in the main text, Supplemental Information, and Appendix A. Primary data from experiments are available from the corresponding authors upon reasonable request.

## REFERENCES

Aguilan, J.T., Kulej, K., and Sidoli, S. (2020) Guide for protein fold change and p-value calculation for non-experts in proteomics. Mol Omics 16: 573–582.

Alonso, C.A., Dominguez, C., Heras, J., Mata, E., Pascual, V., Torres, C., and Zarazaga, (2017) Antibiogramj: A tool for analysing images from disk diffusion tests. Comput Methods Programs Biomed 143: 159–169.

Baylay, A.J., Ivens, A., and Piddock, L.J. (2015) A novel gene amplification causes upregulation of the PatAB ABC transporter and fluoroquinolone resistance in *Streptococcus pneumoniae*. Antimicrob Agents Chemother 59: 3098–3108.

Beilharz, K., Novakova, L., Fadda, D., Branny, P., Massidda, O., and Veening, J.W. (2012) Control of cell division in *Streptococcus pneumoniae* by the conserved Ser/Thr protein kinase StkP. Proc Natl Acad Sci U S A 109: E905–913.

Blake, K.L., O’Neill, A.J., Mengin-Lecreulx, D., Henderson, P.J., Bostock, J.M., Dunsmore, C.J., Simmons, K.J., Fishwick, C.W., Leeds, J.A., and Chopra, I. (2009) The nature of *Staphylococcus aureus* MurA and MurZ and approaches for detection of peptidoglycan biosynthesis inhibitors. Mol Microbiol 72: 335–343.

Booth, S., and Lewis, R.J. (2019) Structural basis for the coordination of cell division with the synthesis of the bacterial cell envelope. Protein Sci 28: 2042–2054.

Boutte, C.C., Baer, C.E., Papavinasasundaram, K., Liu, W., Chase, M.R., Meniche, X., Fortune, S.M., Sassetti, C.M., Ioerger, T.R., and Rubin, E.J. (2016) A cytoplasmic peptidoglycan amidase homologue controls mycobacterial cell wall synthesis. Elife 5: e14590.

Briggs, N.S., Bruce, K.E., Naskar, S., Winkler, M.E., and Roper, D.I. (2021) The pneumococcal divisome: dynamic control of *Streptococcus pneumoniae* cell division. Front Microbiol 12: e737396.

Brown, E.D., Vivas, E.I., Walsh, C.T., and Kolter, R. (1995) MurA (MurZ), the enzyme that catalyzes the first committed step in peptidoglycan biosynthesis, is essential in *Escherichia coli*. J Bacteriol 177: 4194–4197.

Bush, K., and Bradford, P.A. (2016) β-Lactams and β-lactamase inhibitors: an overview. Cold Spring Harbor Perspect Med 6: a025247.

CDC, (2019) Antibiotic resistance threats in the United States, 2019. Atlanta, GA;U.S. Department of Health and Human Services, CDC. Available from: http://www.cdc.gov/drugresistance/Biggest-Threats.html.

Chan, H., Taib, N., Gilmore, M.C., Mohamed, A.M.T., Hanna, K., Luhur, J., Nguyen, H., Hafiz, E., Cava, F., Gribaldo, S., Rudner, D., and Rodrigues, C.D.A. (2022) Genetic screens identify additional genes implicated in envelope remodeling during the engulfment stage of *Bacillus subtilis* sporulation. mBio 13: e0173222.

Claessen, D., Emmins, R., Hamoen, L.W., Daniel, R.A., Errington, J., and Edwards, D.H. (2008) Control of the cell elongation-division cycle by shuttling of PBP1 protein in *Bacillus subtilis*. Mol Microbiol 68: 1029–1046.

Cleverley, R.M., Rutter, Z.J., Rismondo, J., Corona, F., Tsui, H.T., Alatawi, F.A., Daniel, R.A., Halbedel, S., Massidda, O., Winkler, M.E., and Lewis, R.J. (2019) The cell cycle regulator GpsB functions as cytosolic adaptor for multiple cell wall enzymes. Nat Commun 10: 261.

Cowley, L.A., Petersen, F.C., Junges, R., Jimson, D.J.M., Morrison, D.A., and Hanage, W.P. (2018) Evolution via recombination: cell-to-cell contact facilitates larger recombination events in *Streptococcus pneumoniae*. PLoS Genet 14: e1007410.

Cox, M.J., Loman, N., Bogaert, D., and O’Grady, J. (2020) Co-infections: potentially lethal and unexplored in COVID-19. The Lancet Microbe 1: e11.

Cuppone, A.M., Colombini, L., Fox, V., Pinzauti, D., Santoro, F., Pozzi, G., and Iannelli, F. (2021) Complete genome sequence of *Streptococcus pneumoniae* strain Rx1, a Hex mismatch repair-deficient standard transformation recipient. Microbiol Resour Announc 10: e0079921.

Deatherage, D.E., and Barrick, J.E. (2014) Identification of mutations in laboratory-evolved microbes from next-generation sequencing data using breseq. Methods Mol Biol 1151: 165–188.

Denapaite, D., Brückner, R., Hakenbeck, R., and Vollmer, W. (2012) Biosynthesis of teichoic acids in *Streptococcus pneumoniae* and closely related species: lessons from genomes. Microb Drug Resist 18: 344–358.

Dias, R., Felix, D., Canica, M., and Trombe, M.C. (2009) The highly conserved serine threonine kinase StkP of *Streptococcus pneumoniae* contributes to penicillin susceptibility independently from genes encoding penicillin-binding proteins. BMC Microbiol 9: 121.

Du, W., Brown, J.R., Sylvester, D.R., Huang, J., Chalker, A.F., So, C.Y., Holmes, D.J., Payne, D.J., and Wallis, N.G. (2000) Two active forms of UDP-N-acetylglucosamine enolpyruvyl transferase in gram-positive bacteria. J Bacteriol 182: 4146–4152.

Echenique, J., Kadioglu, A., Romao, S., Andrew, P.W., and Trombe, M.C. (2004) Protein serine/threonine kinase StkP positively controls virulence and competence in *Streptococcus pneumoniae*. Infect Immun 72: 2434–2437.

Egan, A.J.F., Errington, J., and Vollmer, W. (2020) Regulation of peptidoglycan synthesis and remodelling. Nat Rev Microbiol 18: 446–460.

Eswara, P.J., Brzozowski, R.S., Viola, M.G., Graham, G., Spanoudis, C., Trebino, C., Jha, J., Aubee, J.I., Thompson, K.M., Camberg, J.L., and Ramamurthi, K.S. (2018) An essential *Staphylococcus aureus* cell division protein directly regulates FtsZ dynamics. Elife 7: 38856.

Fenton, A.K., Manuse, S., Flores-Kim, J., Garcia, P.S., Mercy, C., Grangeasse, C., Bernhardt, T.G., and Rudner, D.Z. (2018) Phosphorylation-dependent activation of the cell wall synthase PBP2a in *Streptococcus pneumoniae* by MacP. Proc Natl Acad Sci U S A 115: 2812–2817.

Fleurie, A., Cluzel, C., Guiral, S., Freton, C., Galisson, F., Zanella-Cleon, I., Di Guilmi, A.M., and Grangeasse, C. (2012) Mutational dissection of the S/T-kinase StkP reveals crucial roles in cell division of *Streptococcus pneumoniae*. Mol Microbiol 83: 746–758.

Fleurie, A., Manuse, S., Zhao, C., Campo, N., Cluzel, C., Lavergne, J.P., Freton, C., Combet, C., Guiral, S., Soufi, B., Macek, B., Kuru, E., VanNieuwenhze, M.S., Brun, Y.V., Di Guilmi, A.M., Claverys, J.P., Galinier, A., and Grangeasse, C. (2014) Interplay of the serine/threonine-kinase StkP and the paralogs DivIVA and GpsB in pneumococcal cell elongation and division. PLoS Genet 10: e1004275.

Garde, S., Chodisetti, P.K., and Reddy, M. (2021) Peptidoglycan: structure, synthesis, and regulation. EcoSal Plus 9: ESP-0010-2020.

Giefing, C., Jelencsics, K.E., Gelbmann, D., Senn, B.M., and Nagy, E. (2010) The pneumococcal eukaryotic-type serine/threonine protein kinase StkP co-localizes with the cell division apparatus and interacts with FtsZ in vitro. Microbiol 156: 1697–1707.

Grangeasse, C. (2016) Rewiring the pneumococcal cell cycle with serine/threonine- and tyrosine-kinases. Trends Microbiol 24: 713–724.

Halbedel, S., and Lewis, R.J. (2019) Structural basis for interaction of DivIVA/GpsB proteins with their ligands. Mol Microbiol 111: 1404–1415.

Hammond, L.R., Sacco, M.D., Khan, S.J., Spanoudis, C., Hough-Neidig, A., Chen, Y., and Eswara, P.J. (2022) GpsB coordinates cell division and cell surface decoration by wall teichoic acids in *Staphylococcus aureus*. Microbiol Spect 10: e01413–01422.

Hammond, L.R., White, M.L., and Eswara, P.J. (2019) ¡vIVA la DivIVA! J Bacteriol 201: e00245-00219.

Hardt, P., Engels, I., Rausch, M., Gajdiss, M., Ulm, H., Sass, P., Ohlsen, K., Sahl, H.G., Bierbaum, G., Schneider, T., and Grein, F. (2017) The cell wall precursor lipid II acts as a molecular signal for the Ser/Thr kinase PknB of *Staphylococcus aureus*. Int J Med Microbiol 307: 1–10.

Herbert, J.A., Mitchell, A.M., and Mitchell, T.J. (2015) A serine-threonine kinase (StkP) regulates expression of the pneumococcal pilus and modulates bacterial adherence to human epithelial and endothelial cells *in vitro*. Plos One 10: e0127212.

Hirschfeld, C., Gomez-Mejia, A., Bartel, J., Hentschker, C., Rohde, M., Maass, S., Hammerschmidt, S., and Becher, D. (2019) Proteomic investigation uncovers potential targets and target sites of pneumococcal serine-threonine kinase StkP and phosphatase PhpP. Front Microbiol 10: 3101.

Holeckova, N., Doubravova, L., Massidda, O., Molle, V., Buriankova, K., Benada, O., Kofronova, O., Ulrych, A., and Branny, P. (2014) LocZ is a new cell division protein involved in proper septum placement in *Streptococcus pneumoniae*. mBio 6: e01700–01714.

Hoskins, J., Alborn, W.E., Jr., Arnold, J., Blaszczak, L.C., Burgett, S., DeHoff, B.S., Estrem, S.T., Fritz, L., Fu, D.J., Fuller, W., Geringer, C., Gilmour, R., Glass, J.S., Khoja, H., Kraft, A.R., Lagace, R.E., LeBlanc, D.J., Lee, L.N., Lefkowitz, E.J., Lu, J., Matsushima, P., McAhren, S.M., McHenney, M., McLeaster, K., Mundy, C.W., Nicas, T.I., Norris, F.H., O’Gara, M., Peery, R.B., Robertson, G.T., Rockey, P., Sun, P.M., Winkler, M.E., Yang, Y., Young-Bellido, M., Zhao, G., Zook, C.A., Baltz, R.H., Jaskunas, S.R., Rosteck, P.R., Jr., Skatrud, P.L., and Glass, J.I. (2001) Genome of the bacterium *Streptococcus pneumoniae* strain R6. J Bacteriol 183: 5709–5717.

Hummels, K.R., Berry, S.P., Li, Z., Taguchi, A., Min, J.K., Walker, S., Marks, D.S., and Bernhardt, T.G. (2023) Coordination of bacterial cell wall and outer membrane biosynthesis. Nature 615: 300–304.

Jackson, S.G., Zhang, F., Chindemi, P., Junop, M.S., and Berti, P.J. (2009) Evidence of kinetic control of ligand binding and staged product release in MurA (enolpyruvyl UDP-GlcNAc synthase)-catalyzed reactions. Biochem 48: 11715–11723.

Jacobsen, F.E., Kazmierczak, K.M., Lisher, J.P., Winkler, M.E., and Giedroc, D.P. (2011) Interplay between manganese and zinc homeostasis in the human pathogen *Streptococcus pneumoniae*. Metallomics: integrat biometal sci 3: 38–41.

Johnston, C., Caymaris, S., Zomer, A., Bootsma, H.J., Prudhomme, M., Granadel, C., Hermans, P.W., Polard, P., Martin, B., and Claverys, J.P. (2013) Natural genetic transformation generates a population of merodiploids in *Streptococcus pneumoniae*. PLoS Genet 9: e1003819.

Jumper, J., Evans, R., Pritzel, A., Green, T., Figurnov, M., Ronneberger, O., Tunyasuvunakool, K., Bates, R., Zidek, A., Potapenko, A., Bridgland, A., Meyer, C., Kohl, S.A.A., Ballard, A.J., Cowie, A., Romera-Paredes, B., Nikolov, S., Jain, R., Adler, J., Back, T., Petersen, S., Reiman, D., Clancy, E., Zielinski, M., Steinegger, M., Pacholska, M., Berghammer, T., Silver, D., Vinyals, O., Senior, A.W., Kavukcuoglu, K., Kohli, P., and Hassabis, D. (2021) Applying and improving AlphaFold at CASP14. Proteins 89: 1711–1721.

Kant, S., Sun, Y., and Pancholi, V. (2023) StkP- and PhpP-mediated posttranslational modifications modulate the *S. pneumoniae* metabolism, polysaccharide capsule, and virulence. Infect Immun: e0029622.

Kaur, P., Rausch, M., Malakar, B., Watson, U., Damle, N.P., Chawla, Y., Srinivasan, S., Sharma, K., Schneider, T., Jhingan, G.D., Saini, D., Mohanty, D., Grein, F., and Nandicoori, V.K. (2019) LipidII interaction with specific residues of *Mycobacterium tuberculosis* PknB extracytoplasmic domain governs its optimal activation. Nat Commun 10: 1231.

Kedar, G.C., Brown-Driver, V., Reyes, D.R., Hilgers, M.T., Stidham, M.A., Shaw, K.J., Finn, J., and Haselbeck, R.J. (2008) Comparison of the essential cellular functions of the two *murA* genes of *Bacillus anthracis*. Antimicrob Agents Chemother 52: 2009–2013.

Kelliher, J.L., Grunenwald, C.M., Abrahams, R.R., Daanen, M.E., Lew, C.I., Rose, W.E., and Sauer, J.D. (2021) PASTA kinase-dependent control of peptidoglycan synthesis via ReoM is required for cell wall stress responses, cytosolic survival, and virulence in *Listeria monocytogenes*. PLoS Pathog 17: e1009881.

Kock, H., Gerth, U., and Hecker, M. (2004) MurAA, catalysing the first committed step in peptidoglycan biosynthesis, is a target of Clp-dependent proteolysis in *Bacillus subtilis*. Mol Microbiol 51: 1087–1102.

Kumar, S., Mollo, A., Kahne, D., and Ruiz, N. (2022) The bacterial cell wall: from Lipid II flipping to polymerization. Chem Rev 122: 8884–8910.

Lamanna, M.M., Manzoor, I., Joseph, M., Ye, Z.A., Benedet, M., Zanardi, A., Ren, Z., Wang, X., Massidda, O., Tsui, H.T., and Winkler, M.E. (2022) Roles of RodZ and class A PBP1b in the assembly and regulation of the peripheral peptidoglycan elongasome in ovoid-shaped cells of *Streptococcus pneumoniae* D39. Mol Microbiol 118: 336–368.

Land, A.D., Tsui, H.C., Kocaoglu, O., Vella, S.A., Shaw, S.L., Keen, S.K., Sham, L.T., Carlson, E.E., and Winkler, M.E. (2013) Requirement of essential Pbp2x and GpsB for septal ring closure in *Streptococcus pneumoniae* D39. Mol Microbiol 90: 939–955.

Lanie, J.A., Ng, W.L., Kazmierczak, K.M., Andrzejewski, T.M., Davidsen, T.M., Wayne, K.J., Tettelin, H., Glass, J.I., and Winkler, M.E. (2007) Genome sequence of Avery’s virulent serotype 2 strain D39 of *Streptococcus pneumoniae* and comparison with that of unencapsulated laboratory strain R6. J Bacteriol 189: 38–51.

Le Bourgeois, P., Bugarel, M., Campo, N., Daveran-Mingot, M.L., Labonté, J., Lanfranchi, D., Lautier, T., Pagès, C., and Ritzenthaler, P. (2007) The unconventional Xer recombination machinery of *Streptococci/Lactococci*. PLoS Genet 3: e117.

Manuse, S., Fleurie, A., Zucchini, L., Lesterlin, C., and Grangeasse, C. (2016) Role of eukaryotic-like serine/threonine kinases in bacterial cell division and morphogenesis. FEMS Microbiol Rev 40: 41–56.

Martin, J.E., Edmonds, K.A., Bruce, K.E., Campanello, G.C., Eijkelkamp, B.A., Brazel, E.B., McDevitt, C.A., Winkler, M.E., and Giedroc, D.P. (2017) The zinc efflux activator SczA protects *Streptococcus pneumoniae* serotype 2 D39 from intracellular zinc toxicity. Mol Microbiol 104: 636–651.

Mascari, C.A., Djorić, D., Little, J.L., and Kristich, C.J. (2022) Use of an interspecies chimeric receptor for inducible gene expression reveals that metabolic flux through the peptidoglycan biosynthesis pathway is an important driver of cephalosporin resistance in *Enterococcus faecalis*. J Bacteriol 204: e0060221.

Massidda, O., Novakova, L., and Vollmer, W. (2013) From models to pathogens: how much have we learned about *Streptococcus pneumoniae* cell division? Environ Microbiol 15: 3133–3157.

Minton, N.E., Djorić, D., Little, J., and Kristich, C.J. (2022) Gpsb promotes pasta kinase signaling and cephalosporin resistance in *Enterococcus faecalis*. J Bacteriol 204: e0030422.

Mizyed, S., Oddone, A., Byczynski, B., Hughes, D.W., and Berti, P.J. (2005) UDP-N-acetylmuramic acid (UDP-MurNAc) is a potent inhibitor of MurA (enolpyruvyl-UDP-GlcNAc synthase). Biochem 44: 4011–4017.

Mobegi, F.M., Cremers, A.J., de Jonge, M.I., Bentley, S.D., van Hijum, S.A., and Zomer, A. (2017) Deciphering the distance to antibiotic resistance for the pneumococcus using genome sequencing data. Sci Rep 7: 42808.

Novakova, L., Bezouskova, S., Pompach, P., Spidlova, P., Saskova, L., Weiser, J., and Branny, P. (2010) Identification of multiple substrates of the StkP Ser/Thr protein kinase in *Streptococcus pneumoniae*. J Bacteriol 192: 3629–3638.

Novakova, L., Saskova, L., Pallova, P., Janecek, J., Novotna, J., Ulrych, A., Echenique, J., Trombe, M.C., and Branny, P. (2005) Characterization of a eukaryotic type serine/threonine protein kinase and protein phosphatase of *Streptococcus pneumoniae* and identification of kinase substrates. FEBS J 272: 1243–1254.

Perez, A.J., Cesbron, Y., Shaw, S.L., Bazan Villicana, J., Tsui, H.T., Boersma, M.J., Ye, Z.A., Tovpeko, Y., Dekker, C., Holden, S., and Winkler, M.E. (2019) Movement dynamics of divisome proteins and PBP2x:FtsW in cells of *Streptococcus pneumoniae*. Proc Natl Acad Sci U S A 116: 3211–3220.

Pinas, G.E., Reinoso-Vizcaino, N.M., Yandar Barahona, N.Y., Cortes, P.R., Duran, R., Badapanda, C., Rathore, A., Bichara, D.R., Cian, M.B., Olivero, N.B., Perez, D.R., and Echenique, J. (2018) Crosstalk between the serine/threonine kinase StkP and the response regulator ComE controls the stress response and intracellular survival of *Streptococcus pneumoniae*. PLoS Pathog 14: e1007118.

Pompeo, F., Foulquier, E., Serrano, B., Grangeasse, C., and Galinier, A. (2015) Phosphorylation of the cell division protein GpsB regulates PrkC kinase activity through a negative feedback loop in *Bacillus subtilis*. Mol Microbiol 97: 139–150.

Ramos-Montanez, S., Tsui, H.C., Wayne, K.J., Morris, J.L., Peters, L.E., Zhang, F., Kazmierczak, K.M., Sham, L.T., and Winkler, M.E. (2008) Polymorphism and regulation of the spxB (pyruvate oxidase) virulence factor gene by a CBS-HotDog domain protein (SpxR) in serotype 2 *Streptococcus pneumoniae*. Mol Microbiol 67: 729–746.

Reams, A.B., and Roth, J.R. (2015) Mechanisms of gene duplication and amplification. Cold Spring Harb Perspect Biol 7: a016592.

Rismondo, J., Bender, J.K., and Halbedel, S. (2017) Suppressor Mutations Linking *gpsB* with the first committed step of peptidoglycan biosynthesis in *Listeria monocytogenes*. J Bacteriol 199: e00393–16.

Rismondo, J., Cleverley, R.M., Lane, H.V., Grosshennig, S., Steglich, A., Moller, L., Mannala, G.K., Hain, T., Lewis, R.J., and Halbedel, S. (2016) Structure of the bacterial cell division determinant GpsB and its interaction with penicillin-binding proteins. Mol Microbiol 99: 978–998.

Robertson, G.T., Ng, W.L., Gilmour, R., and Winkler, M.E. (2003) Essentiality of *clpX*, but not *clpP, clpL*, clpC, or clpE, in Streptococcus pneumoniae R6. J Bacteriol 185: 2961-2966.

Rohs, P.D.A., and Bernhardt, T.G. (2021) Growth and Division of the peptidoglycan matrix. Ann Rev Microbiol 75: 315–336.

Rued, B.E., Zheng, J.J., Mura, A., Tsui, H.T., Boersma, M.J., Mazny, J.L., Corona, F., Perez, A.J., Fadda, D., Doubravova, L., Buriankova, K., Branny, P., Massidda, O., and Winkler, M.E. (2017) Suppression and synthetic-lethal genetic relationships of Δ*gpsB* mutations indicate that GpsB mediates protein phosphorylation and penicillin-binding protein interactions in *Streptococcus pneumoniae* D39. Mol Microbiol 103: 931–957.

Sacco, M.D., Hammond, L.R., Noor, R.E., Bhattacharya, D., Madsen, J.J., Zhang, X., Butler, S.G., Kemp, M.T., Jaskolka-Brown, A.C., Khan, S.J., Gelis, I., Eswara, P.J., and Chen, Y. (2022) *Staphylococcus aureus* FtsZ and PBP4 bind to the conformationally dynamic N-terminal domain of GpsB. bioRxiv: 2022.2010.2025.513704.

Samland, A.K., Etezady-Esfarjani, T., Amrhein, N., and Macheroux, P. (2001) Asparagine 23 and aspartate 305 are essential residues in the active site of UDP-N-acetylglucosamine enolpyruvyl transferase from *Enterobacter cloacae*. Biochem 40: 1550–1559.

Santoro, F., Iannelli, F., and Pozzi, G. (2019) Genomics and genetics of S*treptococcus pneumoniae*. Microbiol Spectr 7: GPP3-0025-2018.

Saskova, L., Novakova, L., Basler, M., and Branny, P. (2007) Eukaryotic-type serine/threonine protein kinase StkP is a global regulator of gene expression in *Streptococcus pneumoniae*. J Bacteriol 189: 4168–4179.

Schonbrunn, E., Eschenburg, S., Luger, K., Kabsch, W., and Amrhein, N. (2000) Structural basis for the interaction of the fluorescence probe 8-anilino-1-naphthalene sulfonate (ANS) with the antibiotic target MurA. Proc Natl Acad Sci USA 97: 6345–6349.

Sender, V., Hentrich, K., and Henriques-Normark, B. (2021) Virus-induced changes of the respiratory tract environment promote secondary infections with *Streptococcus pneumoniae*. Front Cell Infect Microbiol 11: 643326.

Skarzynski, T., Mistry, A., Wonacott, A., Hutchinson, S.E., Kelly, V.A., and Duncan, K. (1996) Structure of UDP-N-acetylglucosamine enolpyruvyl transferase, an enzyme essential for the synthesis of bacterial peptidoglycan, complexed with substrate UDP-N-acetylglucosamine and the drug fosfomycin. Structure 4: 1465–1474.

Skinner, M.E., Uzilov, A.V., Stein, L.D., Mungall, C.J., and Holmes, I.H. (2009) JBrowse: a next-generation genome browser. Genome Res 19: 1630–1638.

Slager, J., Aprianto, R., and Veening, J.W. (2018) Deep genome annotation of the opportunistic human pathogen *Streptococcus pneumoniae* D39. Nucleic Acids Res 46: 9971–9989.

Stamsas, G.A., Straume, D., Ruud Winther, A., Kjos, M., Frantzen, C.A., and Havarstein, L.S. (2017) Identification of EloR (Spr1851) as a regulator of cell elongation in Streptococcus pneumoniae. Mol Microbiol 105: 954–967.

Sun, X., Ge, F., Xiao, C.L., Yin, X.F., Ge, R., Zhang, L.H., and He, Q.Y. (2010) Phosphoproteomic analysis reveals the multiple roles of phosphorylation in pathogenic bacterium Streptococcus pneumoniae. J Proteome Res 9: 275–282.

Sun, Y., Hürlimann, S., and Garner, E. (2023) Growth rate is modulated by monitoring cell wall precursors in *Bacillus subtilis*. Nat Microbiol 8: 469–480.

Sung, C.K., Li, H., Claverys, J.P., and Morrison, D.A. (2001) An *rpsL* cassette, Janus, for gene replacement through negative selection in *Streptococcus pneumoniae*. Appl Environ Microbiol 67: 5190–5196.

Taguchi, A., Page, J.E., Tsui, H.T., Winkler, M.E., and Walker, S. (2021) Biochemical reconstitution defines new functions for membrane-bound glycosidases in assembly of the bacterial cell wall. Proc Natl Acad Sci U S A 118.

Tettelin, H., Nelson, K.E., Paulsen, I.T., Eisen, J.A., Read, T.D., Peterson, S., Heidelberg, J., DeBoy, R.T., Haft, D.H., Dodson, R.J., Durkin, A.S., Gwinn, M., Kolonay, J.F., Nelson, W.C., Peterson, J.D., Umayam, L.A., White, O., Salzberg, S.L., Lewis, M.R., Radune, D., Holtzapple, E., Khouri, H., Wolf, A.M., Utterback, T.R., Hansen, C.L., McDonald, L.A., Feldblyum, T.V., Angiuoli, S., Dickinson, T., Hickey, E.K., Holt, I.E., Loftus, B.J., Yang, F., Smith, H.O., Venter, J.C., Dougherty, B.A., Morrison, D.A., Hollingshead, S.K., and Fraser, C.M. (2001) Complete genome sequence of a virulent isolate of *Streptococcus pneumoniae*. Science 293: 498–506.

Tsui, H.C., Zheng, J.J., Magallon, A.N., Ryan, J.D., Yunck, R., Rued, B.E., Bernhardt, T.G., and Winkler, M.E. (2016) Suppression of a deletion mutation in the gene encoding essential PBP2b reveals a new lytic transglycosylase involved in peripheral peptidoglycan synthesis in *Streptococcus pneumoniae* D39. Mol Microbiol 100: 1039–1065.

Tsui, H.T., Boersma, M.J., Vella, S.A., Kocaoglu, O., Kuru, E., Peceny, J.K., Carlson, E.E., VanNieuwenhze, M.S., Brun, Y.V., Shaw, S.L., and Winkler, M.E. (2014) Pbp2x localizes separately from Pbp2b and other peptidoglycan synthesis proteins during later stages of cell division of *Streptococcus pneumoniae* D39. Mol Microbiol 94: 21–40.

Turapov, O., Forti, F., Kadhim, B., Ghisotti, D., Sassine, J., Straatman-Iwanowska, A., Bottrill, A.R., Moynihan, P.J., Wallis, R., Barthe, P., Cohen-Gonsaud, M., Ajuh, P., Vollmer, W., and Mukamolova, G.V. (2018) Two faces of CwlM, an essential PknB substrate, in *Mycobacterium tuberculosis*. Cell Rep 25: 57–67 e55.

Ulrych, A., Fabrik, I., Kupčík, R., Vajrychová, M., Doubravová, L., and Branny, P. (2021) Cell wall stress stimulates the activity of the protein kinase StkP of *Streptococcus pneumoniae*, leading to multiple phosphorylation. J Mol Biol 433: 167319.

Ulrych, A., Holeckova, N., Goldova, J., Doubravova, L., Benada, O., Kofronova, O., Halada, P., and Branny, P. (2016) Characterization of pneumococcal Ser/Thr protein phosphatase *phpP* mutant and identification of a novel PhpP substrate, putative RNA binding protein Jag. BMC Microbiol 16: 247.

Vesic, D., and Kristich, C.J. (2012) MurAA is required for intrinsic cephalosporin resistance of *Enterococcus faecalis*. Antimicrob Agents Chemother 56: 2443–2451.

Vollmer, W., Massidda, O., and Tomasz, A. (2019) The cell wall of *Streptococcus pneumoniae*. Microbiol Spectr 7: GPP3-0018-2018.

Wamp, S., Rothe, P., Stern, D., Holland, G., Döhling, J., and Halbedel, S. (2022) MurA escape mutations uncouple peptidoglycan biosynthesis from PrkA signaling. PLoS Pathog 18: e1010406.

Wamp, S., Rutter, Z.J., Rismondo, J., Jennings, C.E., Möller, L., Lewis, R.J., and Halbedel, S. (2020) PrkA controls peptidoglycan biosynthesis through the essential phosphorylation of ReoM. Elife 9: e56048.

Weiser, J.N., Ferreira, D.M., and Paton, J.C. (2018) *Streptococcus pneumoniae*: transmission, colonization and invasion. Nat Rev Microbiol 16: 355–367.

Westesson, O., Skinner, M., and Holmes, I. (2013) Visualizing next-generation sequencing data with JBrowse. Brief Bioinform 14: 172–177.

WHO (2017) List of bacteria for which new antibiotics are urgently needed. World Health Organization, Geneva, Switzerland: http://www.who.int/mediacentre/news/releases/2017/bacteria-antibiotics-needed/en.

Winther, A.R., Kjos, M., Stamsas, G.A., Havarstein, L.S., and Straume, D. (2019) Prevention of EloR/KhpA heterodimerization by introduction of site-specific amino acid substitutions renders the essential elongasome protein PBP2b redundant in Streptococcus pneumoniae. Sci Rep 9: 3681.

Zheng, J.J., Perez, A.J., Tsui, H.T., Massidda, O., and Winkler, M.E. (2017) Absence of the KhpA and KhpB (JAG/EloR) RNA-binding proteins suppresses the requirement for PBP2b by overproduction of FtsA in *Streptococcus pneumoniae* D39. Mol Microbiol 106: 793–814.

Zhou, J., Cai, Y., Liu, Y., An, H., Deng, K., Ashraf, M.A., Zou, L., and Wang, J. (2022) Breaking down the cell wall: Still an attractive antibacterial strategy. Frontiers in Microbiology 13: 952633.

Zucchini, L., Mercy, C., Garcia, P.S., Cluzel, C., Gueguen-Chaignon, V., Galisson, F., Freton, C., Guiral, S., Brochier-Armanet, C., Gouet, P., and Grangeasse, C. (2018) PASTA repeats of the protein kinase StkP interconnect cell constriction and separation of *Streptococcus pneumoniae*. Nature Microbiology 3: 197–209.

