## Supplemental Information for "Chromosomal Duplications of MurZ (MurA2) or MurA (MurA1), Amino Acid Substitutions in MurZ (MurA2), and Absence of KhpAB Obviate the Requirement for Protein Phosphorylation in *Streptococcus pneumoniae* D39"

### Contents:

**Supplemental Tables S1-S6**

**Supplemental References**

**Supplemental Figures S1-S21 with Legends**

**Table S1.** *Streptococcus pneumoniae* strains and oligonucleotide primers used in this study

| **Strains used in this study** | | | |
|---|---|---|---|
| Strain number | Genotype (description)[a] | Antibiotic resistance[b] | Reference or source |
| EL59 | R6 | None | (Hoskins *et al.*, 2001) |
| IU1690 | D39 *cps*$^+$ (D39W) | None | (Lanie *et al.*, 2007, Slager *et al.*, 2018) |
| IU1781 | D39 *cps*$^+$ *rpsL1* | Str$^R$ | (Lanie *et al.*, 2007) |
| IU1824[c] | D39 *rpsL1* Δ*cps2A'-cps2H'* = D39 *rpsL1* Δ*cps* | Str$^R$ | (Lanie *et al.*, 2007) |
| IU1945 | D39 Δ*cps2A'-cps2H'*= D39 Δ*cps* | None | (Lanie *et al.*, 2007) |
| E193 | D39 Δ*cps* Δ*pbp1b*::$P_c$-*erm* | Erm$^R$ | (Land *et al.*, 2013) |
| E655 | D39 Δ*cps* Δ*rodZ*::$P_c$-*erm* | Erm$^R$ | (Tsui *et al.*, 2016) |
| K180 | D39 Δ*cps* Δ*pbp1b*::$P_c$-[*kan-rpsL*$^+$] | Kan$^R$ | (Tsui *et al.*, 2014) |
| E740 | D39 Δ*cps* Δ[*phpP-stkP*]::$P_c$-*erm sup2* (IU1945 transformed with fusion Δ[*phpP-stkP*]::$P_c$-*erm* amplicon) | Erm$^R$ | This Study |
| E765 | D39 Δ*cps* Δ*murA*::$P_c$-*erm* (IU1945 X fusion Δ*murA*::$P_c$-*erm*) | Erm$^R$ | This Study |
| E767 | D39 Δ*cps* Δ*murZ*::$P_c$-*erm* (IU1945 X fusion Δ*murZ*::$P_c$-*erm*) | Erm$^R$ | This Study |
| E780 | D39 Δ*cps* Δ*clpC*::$P_c$-*erm* (IU1945 X fusion Δ*clpC*::$P_c$-*erm*) | Erm$^R$ | This Study |
| K761 | D39 Δ*cps* Δ*khpB*::$P_c$-[*kan-rpsL*$^+$] | Kan$^R$ | (Zheng *et al.*, 2017) |
| K765 | D39 Δ*cps* Δ*murA*::$P_c$-[*kan-rpsL*$^+$] (IU1945 X fusion Δ*murA*::$P_c$-[*kan-rpsL*$^+$]) | Kan$^R$ | This study |
| K767 | D39 Δ*cps* Δ*murZ*::$P_c$-[*kan-rpsL*$^+$] (IU1945 X fusion Δ*murZ*::$P_c$-[*kan-rpsL*$^+$]) | Kan$^R$ | This Study |
| K779 | D39 Δ*cps* Δ*clpC*::$P_c$-[*kan-rpsL*$^+$] (IU1945 X fusion Δ*clpC*::$P_c$-[*kan-rpsL*$^+$]) | Kan$^R$ | This Study |
| K787 | D39 Δ*cps* Δ*ireB*::$P_c$-[*kan-rpsL*$^+$] (IU1945 X fusion Δ*ireB*::$P_c$-[*kan-rpsL*$^+$]) | Kan$^R$ | This Study |
| IU4355 | *D39 rpsL1* Δ*cps* Δ*bgaA*::*kan*-t1t2-$P_{fcsk}$-*secA*-L-FLAG$^3$ | Str$^R$ Kan$^R$ | (Tsui *et al.*, 2011) |
| IU4888 | D39 Δ*cps* Δ*gpsB*<>*aad9*//Δ*bgaA*::*kan*-t1t2-$P_{fcsK}$-*gpsB*$^+$ | Kan$^R$ Spc$^R$ | (Land *et al.*, 2013) |
| IU4970 | D39 Δ*cps* *mreC*-L-FLAG$^3$-$P_c$-*erm* | Erm$^R$ | (Land & Winkler, 2011) |

| | | | |
|---|---|---|---|
| IU5845 | D39 Δ*cps* Δ*gpsB*<>*aad9 sup2* | Spc$^{R}$ | (Rued *et al.*, 2017) |
| IU6441 | D39 Δ*cps* Δ*gpsB*<>*aad9 sup3* | Spc$^{R}$ | (Rued *et al.*, 2017) |
| IU6442 | D39 Δ*cps* Δ*gpsB*<>*aad9 sup1* | Spc$^{R}$ | (Rued *et al.*, 2017) |
| IU6444 | D39 Δ*cps* Δ*gpsB*<>*aad9 sup5* | Spc$^{R}$ | (Rued *et al.*, 2017) |
| IU7397 | D39 Δ*cps* Δ*pbp2b*<>*aad9* //Δ*bga*::*kan*-t1t2-P$_{fcsk}$-*pbp2b*$^{+}$ | Spc$^{R}$ Kan$^{R}$ | (Tsui *et al.*, 2014) |
| IU7673 | D39 Δ*cps rpsL1 phpP*$^{+}$-P$_{c}$-[*kan-rpsL*$^{+}$]-*stkP*$^{+}$ | Kan$^{R}$ | (Rued *et al.*, 2017) |
| IU7735 | D39 Δ*cps rpsL1* Δ*gpsB*<>*aad9 sup27* with spontaneous *ireB*(Q84(STOP)) mutation (IU1824 X Δ*gpsB*<>*aad9* amplicon from IU4888). Original Δ*gpsB* suppressor strain in IU1824 background. | Str$^{R}$ Spc$^{R}$ | This study |
| IU7736 | D39 Δ*cps rpsL1* Δ*gpsB*<>*aad9 sup6* | Str$^{R}$ Spc$^{R}$ | (Rued *et al.*, 2017) |
| IU7824 | D39 Δ*cps* Δ[*spd_1031-1037*]::P$_{c}$-*erm* | Erm$^{R}$ | (Rued *et al.*, 2017) |
| IU7923 | D39 Δ*cps* Δ*stkP*::P$_{c}$-*erm* | Erm$^{R}$ | (Rued *et al.*, 2017) |
| IU8108 | D39 Δ*cps* Δ[*spd_1029-1030*]::P$_{c}$-*erm* (IU1945 X fusion Δ[*spd_1029-1030*]::P$_{c}$-*erm*) | Erm$^{R}$ | This study |
| IU8122 | D39 Δcps Δ*bgaA*::*tet*-P$_{Zn}$-RBS$^{ftsA}$-*ftsZ*$^{+}$ | Tet$^{R}$ | (Zheng *et al.*, 2017) |
| IU8224 | R6 Δ*gpsB*<>*aad9* | Spc$^{R}$ | (Rued *et al.*, 2017) |
| IU8271 | D39 Δ*cps* Δ[*spd_1029-1037*]::P$_{c}$-[*kan-rpsL*$^{+}$] | Kan$^{R}$ | (Rued *et al.*, 2017) |
| IU8742 | D39 Δ*cps* Δ*gpsB*<>*aad9 phpP*(G229D) Δ*bgaA*::*tet*-P$_{Zn}$-RBS$^{ftsA}$-*phpP*$^{+}$ (IU6442 X fusion Δ*bgaA*::*tet*-P$_{Zn}$-RBS$^{ftsA}$-*phpP*$^{+}$ amplicon) | Spc$^{R}$ Tet$^{R}$ | This Study |
| IU8791 | D39 Δ*cps* Δ*mltG*::P$_{c}$-*aad9*//Δ*bgaA*::*kan*-t1t2-P$_{fcsK}$-*mltG*$^{+}$ | Kan$^{R}$ Spc$^{R}$ | (Tsui *et al.*, 2016) |
| IU8872 | D39 Δ*cps* Δ*bgaA*::*tet*-P$_{Zn}$-RBS$^{mltG}$–*mltG*$^{+}$ | Tet$^{R}$ | (Tsui *et al.*, 2016) |
| IU9036 | D39 Δ*cps rpsL1* Δ*khpA* | Str$^{R}$ | (Zheng *et al.*, 2017) |
| IU9262 | Rx1 Δ*gpsB*<>*aad9 sup4* | Spc$^{R}$ | (Rued *et al.*, 2017) |
| IU9600 | D39 Δ*khpA*::P$_{c}$-*erm* | Erm$^{R}$ | (Zheng *et al.*, 2017) |
| IU9613 | D39 Δ*cps rpsL1* Δ*bgaA*::*tet*-P$_{Zn}$-RBS$^{ftsA}$-*rodZ*$^{+}$ (IU1824 X fusion Δ*bgaA*::*tet*-P$_{Zn}$-RBS$^{ftsA}$-*rodZ*$^{+}$) | Str$^{R}$ Tet$^{R}$ | This Study |
| IU9765 | D39 Δ*cps* Δ*bgaA*::*tet*-P$_{Zn}$-RBS$^{ftsA}$-*rodZ*$^{+}$ | Tet$^{R}$ | (Tsui *et al.*, 2016) |

| | | | |
|---|---|---|---|
| IU9805 | D39 Δ*cps* Δ*bgaA*::*kan*-t1t2-$P_{Zn}$-RBS$^{ftsA}$-*sepF*$^+$ (IU1945 X fusion Δ*bgaA*::*kan*-t1t2-$P_{Zn}$-RBS$^{ftsA}$-*sepF*$^+$) | Kan$^R$ | This Study |
| IU9931 | D39 Δ*cps* Δ*rodZ*<>*aad9*//Δ*bgaA*::*tet*-$P_{Zn}$-RBS$^{ftsA}$-*rodZ*$^+$ | Spc$^R$ Tet$^R$ | (Tsui *et al.*, 2016) |
| IU9990 | D39 Δ*cps* Δ*bgaA*::*tet*-$P_{Zn}$-RBS$^{ftsA}$-*pbp2b*$^+$ | Tet$^R$ | (Zheng *et al.*, 2017) |
| IU9992 | D39 Δ*cps* Δ*bgaA*::*tet*-$P_{Zn}$-RBS$^{ftsA}$-*pbp1b*$^+$ (IU1945 X fusion Δ*bgaA*::tet-$P_{Zn}$-RBS$^{ftsA}$-*pbp1b*$^+$) | Tet$^R$ | This Study |
| IU10063 | D39 Δ*cps* Δ*bgaA*::*tet*-$P_{Zn}$-RBS$^{ftsA}$-*pbp2x*$^+$ | Tet$^R$ | (Perez *et al.*, 2019) |
| IU10220 | D39 Δ*cps rpsL1* Δ*bgaA*::*tet*-$P_{Zn}$-RBS$^{ftsA}$-*mreC*$^+$ (IU1824 X fusion Δ*bgaA*::tet-$P_{Zn}$-RBS$^{ftsA}$-*mreC*$^+$) | Str$^R$ Tet$^R$ | This Study |
| IU10592 | D39 Δ*cps rpsL1* Δ*khpB* | Str$^R$ | (Zheng *et al.*, 2017) |
| IU10596 | D39 Δ*cps rpsL1* Δ*khpA* Δ*khpB* | Str$^R$ | (Zheng *et al.*, 2017) |
| IU10659 | D39 Δ*cps rpsL1* Δ*khpA* Δ*rodZ*<>*aad9* | Str$^R$ Spc$^R$ | (Zheng *et al.*, 2017) |
| IU10922 | D39 Δ*cps* Δ*bgaA*::*tet*-$P_{Zn}$-RBS$^{ftsA}$-*rodA*$^+$ | Tet$^R$ | (Tsui *et al.*, 2016) |
| IU11049 | D39 Δ*cps* Δ*bgaA*::*kan*-t1t2-$P_{Zn}$-*murG*$^+$ (IU1945 X fusion Δ*bgaA*::*kan*-t1t2-$P_{Zn}$-*murG*$^+$) | Kan$^R$ | This Study |
| IU11077 | D39 Δ*cps* Δ*bgaA*::*kan*-t1t2-$P_{Zn}$-*murZ*$^+$ (IU1945 X fusion Δ*bgaA*::*kan*-t1t2-$P_{Zn}$-*murZ*$^+$) | Kan$^R$ | This Study |
| IU11079 | D39 Δ*cps* Δ*bgaA*::*kan*-t1t2-$P_{Zn}$-*murA*$^+$ (IU1945 X fusion Δ*bgaA*::*kan*-t1t2-$P_{Zn}$-*murA*$^+$) | Kan$^R$ | This Study |
| IU11083 | D39 Δ*cps* Δ*bgaA*::*kan*-t1t2-$P_{Zn}$-*mraY*$^+$ (IU1945 X fusion Δ*bgaA*::*kan*-t1t2-$P_{Zn}$-*mraY*$^+$) | Kan$^R$ | This Study |
| IU11094 | D39 Δ*cps* Δ*bgaA*::*kan*-t1t2-$P_{Zn}$-*uppS*$^+$ (IU1945 X fusion Δ*bgaA*::*kan*-t1t2-$P_{Zn}$-*uppS*$^+$) | Kan$^R$ | This Study |
| IU11286 | D39 Δ*cps* Δ*bgaA*::*tet*-$P_{Zn}$-RBS$^{ftsA}$-*gpsB*$^+$ | Tet$^R$ | (Cleverley *et al.*, 2019) |
| IU11456 | D39 Δ*stkP*::$P_c$-*erm sup4* | Erm$^R$ | (Rued *et al.*, 2017) |
| IU11628 | D39 Δ*cps* Δ*bgaA*::*kan*-t1t2-$P_{Zn}$-*mapZ*$^+$ (IU1945 X fusion Δ*bgaA*::*kan*-t1t2-$P_{Zn}$-*mapZ*$^+$) | Kan$^R$ | This Study |
| IU11846 | D39 Δ*cps* Δ*gpsB*<>*aad9 sup9* | Spc$^R$ | This Study |
| IU11912 | D39 Δ*cps* Δ*stkP*::$P_c$-*cat sup3* (IU1945 X fusion Δ*stkP*::$P_c$-*cat*) | Cm$^R$ | This Study |
| IU11914 | D39 Δ*cps* Δ*gpsB*<>*aad9 sup11* | Spc$^R$ | This Study |
| IU11918 | D39 Δ*cps* Δ*gpsB*<>*aad9 sup10* | Spc$^R$ | This Study |
| IU11954 | D39 Δ*cps* Δ*gpsB*<>*aad9 sup8* | Spc$^R$ | This Study |
| IU11955 | D39 Δ*cps* Δ*gpsB*<>*aad9 sup7* | Spc$^R$ | (Rued *et al.*, 2017) |
| IU12192 | D39 Δ*cps rpsL1* Δ*bgaA*::*tet*-$P_{Zn}$-RBS$^{ftsA}$-*ftsW*$^+$ (IU1824 X fusion Δ*bgaA*::*tet*-$P_{Zn}$-RBS$^{ftsA}$-*ftsW*$^+$) | Str$^R$ Tet$^R$ | This Study |
| IU12286 | D39 Δ*cps rpsL1* Δ*bgaA*::*tet*-$P_{Zn}$-RBS$^{ftsA}$-*ftsZ*$^+$ | Str$^R$ Tet$^R$ | (Zheng *et al.*, 2017) |

| | | | |
|---|---|---|---|
| IU12307 | D39 Δ*cps* Δ*bgaA*::*tet*-$P_{Zn}$-RBS$^{ftsA}$-*ftsA*$^{+}$ | Str$^{R}$ Tet$^{R}$ | (Mura *et al.*, 2017) |
| IU12310 | D39 Δ*cps* *rpsL1* Δ*bgaA*::*tet*-$P_{Zn}$-RBS$^{ftsA}$-*ftsA*$^{+}$ | Str$^{R}$ Tet$^{R}$ | (Mura *et al.*, 2017) |
| IU12428 | D39 Δ*cps* Δ*bgaA*::*kan*-t1t2-$P_{Zn}$-RBS$^{ftsA}$-*murA*$^{+}$ Δ*gpsB*<>*aad9* (IU11079 X Δ*gpsB*<>*aad9* amplicon from IU4888) | Kan$^{R}$ Spec$^{R}$ | This Study |
| IU12462 | D39 Δ*cps* *rpsL1* Δ*clpC::*$P_{c}$-*kan-rpsL*$^{+}$ (IU1824 X Δ*clpC::*$P_{c}$-*kan-rpsL*$^{+}$ amplicon from K779) | Kan$^{R}$ | This Study |
| IU12678 | D39 Δ*cps* Δ*bgaA*::*tet*-$P_{Zn}$-RBS$^{ftsA}$-*cozE*$^{+}$ (IU1945 X fusion Δ*bgaA*::tet-$P_{Zn}$-RBS$^{ftsA}$-*cozE*$^{+}$) | Tet$^{R}$ | This Study |
| IU12704 | D39 Δ*cps* Δ*bgaA*::*tet*-$P_{Zn}$-RBS$^{ftsA}$-*ftsA*$^{+}$ Δ*pbp2b*<>*aad9* (IU12307 X Δ*pbp2b*<>*aad9* from IU7397) | Tet$^{R}$ Spec$^{R}$ | This Study |
| IU12707 | D39 Δ*cps* *rpsL1* Δ*bgaA*::*tet*-$P_{Zn}$-RBS$^{ftsA}$-*ftsA*$^{+}$ Δ*pbp2b*<>*aad9* | Tet$^{R}$ Spec$^{R}$ | (Zheng *et al.*, 2017) |
| IU12712 | D39 Δ*cps* Δ*bgaA*::*kan*-t1t2-$P_{ftsA}$-RBS$^{ftsA}$-*ftsA*$^{+}$ (IU1945 X fusion Δ*bgaA*::*kan*- t1t2-$P_{ftsA}$-RBS$^{ftsA}$-*ftsA*) | Kan$^{R}$ | This study |
| IU12719 | D39 Δ*cps* *rpsL1* Δ*bgaA*::*kan*-t1t2-$P_{ftsA}$-RBS$^{ftsA}$-*ftsA* (IU1824 X fusion Δ*bgaA*::*kan*- t1t2-$P_{ftsA}$-RBS$^{ftsA}$-*ftsA*) | Str$^{R}$ Kan$^{R}$ | This study |
| IU12883 | D39 Δ*cps* *rpsL1* Δ*khpA* Δ*gpsB*<>*aad9* | Str$^{R}$ Spc$^{R}$ | (Zheng *et al.*, 2017) |
| IU12977 | D39 Δ*cps* *rpsL1* Δ*khpB* Δ*gpsB*<>*aad9* | Str$^{R}$ Spc$^{R}$ | (Zheng *et al.*, 2017) |
| IU13249 | D39 Δ*cps* *rpsL1* *murZ*-L-FLAG$^{3}$-$P_{c}$-*erm* (IU1824 X fusion *murZ*-L-FLAG$^{3}$-$P_{c}$-*erm*) | Erm$^{R}$ Str$^{R}$ | This Study |
| IU13251 | D39 Δ*cps* *rpsL1* *murA*-L-FLAG$^{3}$-$P_{c}$-*erm* (IU1824 X fusion *murA*-L-FLAG$^{3}$-$P_{c}$-*erm*) | Erm$^{R}$ Str$^{R}$ | This study |
| IU13283 | D39 Δ*cps* *rpsL1* Δ*khpA* *murZ*-L-FLAG$^{3}$-$P_{c}$-*erm* (IU9036 X *murZ*-L-FLAG$^{3}$-$P_{c}$-*erm* from IU13249) | Erm$^{R}$ Str$^{R}$ | This Study |
| IU13285 | D39 Δ*cps* *rpsL1* Δ*khpA* *murA*-L-FLAG$^{3}$-$P_{c}$-*erm* (IU9036 X *murA*-L-FLAG$^{3}$-$P_{c}$-*erm* from IU13251) | Erm$^{R}$ Str$^{R}$ | This Study |
| IU13327 | D39 Δ*cps* *rpsL1* CEP::$P_{Zn}$-*ezrA*$^{+}$ Δ*bgaA*::*kan*-t1t2-$P_{Zn}$-RBS$^{ftsA}$-*ezrA*$^{+}$ | Kan$^{R}$ Str$^{R}$ | (Perez *et al.*, 2021) |
| IU13393 | D39 Δ*cps* *rpsL1* Δ*bgaA*::*kan*-t1t2-$P_{Zn}$-RBS$^{ftsA}$-*murZ*$^{+}$ (IU1824 X Δ*bgaA*::*kan*-t1t2-$P_{Zn}$-*murZ*$^{+}$ amplicon from IU11077) | Kan$^{R}$ Str$^{R}$ | This Study |
| IU13395 | D39 Δ*cps* *rpsL1* Δ*bgaA*::*kan*-t1t2-$P_{Zn}$-RBS$^{ftsA}$ -*murA*$^{+}$ (IU1824 X Δ*bgaA*::*kan*-t1t2-$P_{Zn}$-*murA*$^{+}$ amplicon from IU11079) | Kan$^{R}$ Str$^{R}$ | This Study |
| IU13396 | D39 Δ*cps* *rpsL1* Δ*murZ*::$P_{c}$-[*kan-rpsL*$^{+}$] (IU1824 X Δ*murZ*::$P_{c}$-[*kan-rpsL*$^{+}$] amplicon from K767) | Kan$^{R}$ | This Study |
| IU13438 IU13439 | D39 Δ*cps* *rpsL1* *murZ*(D280Y) (IU13396 X *murZ*(D280Y) amplicon from IU11914) | Str$^{R}$ | This Study |
| IU13485 | D39 Δ*cps* *rpsL1* *murZ*(D280Y) Δ*gpsB*<>*aad9* (IU13438 X Δ*gpsB*<>*aad9* amplicon from IU4888) | Str$^{R}$ Spc$^{R}$ | This Study |

| | | | |
|---|---|---|---|
| IU13491 | D39 Δ*cps rpsL1* Δ*murA*::P$_{c}$-[*kan-rpsL*$^{+}$] (IU1824 X Δ*murA*::P$_{c}$-[*kan-rpsL*$^{+}$] amplicon from K765) | Kan$^{R}$ | This Study |
| IU13493 | D39 Δ*cps rpsL1* Δ*khpA* Δ*murZ*::P$_{c}$-[*kan-rpsL*$^{+}$] (IU9036 X Δ*murZ*::P$_{c}$-[*kan-rpsL*$^{+}$] amplicon from K767) | Kan$^{R}$ | This Study |
| IU13495 | D39 Δ*cps rpsL1* Δ*khpA* Δ*murA*::P$_{c}$-[*kan-rpsL*$^{+}$] (IU9036 X Δ*murA*::P$_{c}$-[*kan-rpsL*$^{+}$] amplicon from K765) | Kan$^{R}$ | This Study |
| IU13502 | D39 Δ*cps rpsL1 murZ*-L-FLAG$^{3}$ (IU13396 X fusion *murZ*-L-FLAG$^{3}$) | Str$^{R}$ | This Study |
| IU13505 | D39 Δ*cps rpsL1 murZ*(D280Y) Δ*gpsB*<>*aad9* (IU13438 X Δ*gpsB*<>*aad9* amplicon from IU4888) | Str$^{R}$ Spc$^{R}$ | This Study |
| IU13509 | D39 Δ*cps rpsL1 murZ*(D280Y) Δ*gpsB*<>*aad9* (IU13438 X Δ*gpsB*<>*aad9* amplicon from IU4888) | Str$^{R}$ Spc$^{R}$ | This Study |
| IU13536 | D39 Δ*cps rpsL1* Δ*murZ* (IU13396 X fusion Δ*murZ*) | Str$^{R}$ | This Study |
| IU13538 | D39 Δ*cps rpsL1* Δ*murA* (IU13491 X fusion Δ*murA*) | Str$^{R}$ | This Study |
| IU13542 | D39 Δ*cps rpsL1* Δ*khpA* Δ*murZ* (IU13493 X fusion Δ*murZ*) | Str$^{R}$ | This Study |
| IU13545 | D39 Δ*cps rpsL1* Δ*khpA murZ*-L-FLAG$^{3}$ (IU13493 X fusion *murZ*-L-FLAG$^{3}$) | Str$^{R}$ | This Study |
| IU13546 | D39 Δ*cps rpsL1* Δ*khpA* Δ*murA* (IU13495 X fusion Δ*murA*) | Str$^{R}$ | This Study |
| IU13590 | D39 Δ*cps rpsL1* Δ*ireB*::P$_{c}$-[*kan-rpsL*$^{+}$] (IU1824 X Δ*ireB*::P$_{c}$-[*kan-rpsL*$^{+}$] amplicon from K787) | Kan$^{R}$ | This Study |
| IU13600 | D39 Δ*cps rpsL1 murZ*(D280Y)-L-FLAG$^{3}$ (IU13396 X fusion *murZ*(D280Y)-L-FLAG$^{3}$) | Str$^{R}$ | This Study |
| IU13604 | D39 Δ*cps rpsL1* Δ*ireB* markerless (IU13590 X fusion Δ*ireB* markerless amplicon) | Str$^{R}$ | This Study |
| IU13606 | D39 Δ*cps rpsL1 ireB*(Q84(STOP)) (IU13590 X *ireB*(Q84(STOP)) amplicon from IU7735) | Str$^{R}$ | This Study |
| IU13680 | D39 Δ*cps* Δ*pbp1b*::P$_{c}$-*aad9* (IU1945 X fusion Δ*pbp1b*::P$_{c}$-*aad9*) | Spc$^{R}$ | This Study |
| IU13756 | D39 Δ*cps* Δ*bgaA*::*kan*-t1t2-P$_{Zn}$-RBS$^{ftsA}$-*murZ*$^{+}$ Δ*gpsB*<>*aad9* (IU11077 X Δ*gpsB*<>*aad9* amplicon from IU4888) | Kan$^{R}$ Spc$^{R}$ | This Study |
| IU13757 | D39 Δ*cps* Δ*bgaA*::*kan*-t1t2-P$_{Zn}$-RBS$^{ftsA}$-*murA*$^{+}$ Δ*gpsB*<>*aad9* (IU11079 X Δ*gpsB*<>*aad9* amplicon from IU4888) | Kan$^{R}$ Spc$^{R}$ | This Study |
| IU13772 | D39 Δ*cps rpsL1 murZ*-L-FLAG$^{3}$ Δ*bgaA*::*kan*-t1t2-P$_{Zn}$-RBS$^{ftsA}$-*murZ*-L-FLAG$^{3}$ (IU13502 X fusion Δ*bgaA*::*kan*-t1t2-P$_{Zn}$-RBS$^{ftsA}$-*murZ*-L-FLAG$^{3}$) | Str$^{R}$ Kan$^{R}$ | This Study |
| IU13794 | D39 Δ*cps rpsL1* Δ*bgaA*::*tet*-P$_{Zn}$-RBS$^{ftsA}$-*divIVA*$^{+}$ (R6 annotation) (IU1824 X fusion Δ*bgaA*::*tet*-P$_{Zn}$-RBS$^{ftsA}$–*divIVA*$^{+}$) | Tet$^{R}$ Str$^{R}$ | This Study |

| | | | |
|---|---|---|---|
| IU13987 | D39 Δ*cps rpsL1* Δ*khpB*::$P_c$-[*kan-rpsL*$^+$] *murZ*-L-FLAG$^3$ (IU13502 X Δ*khpB*::$P_c$-[*kan-rpsL*$^+$] from K761) | Kan$^R$ | This Study |
| IU13989 | D39 Δ*cps rpsL1* Δ*khpA* Δ*khpB*::$P_c$-[*kan-rpsL*$^+$] *murZ*-L-FLAG$^3$ (IU13545 X Δ*khpB*::$P_c$-[*kan-rpsL*$^+$] from K761) | Kan$^R$ | This Study |
| IU14014 | D39 Δ*cps rpsL1* Δ*khpB murZ*-L-FLAG$^3$ (IU13987 X Δ*khpB* from IU10592) | Str$^R$ | This Study |
| IU14016 | D39 Δ*cps rpsL1* Δ*khpA* Δ*khpB murZ*-L-FLAG$^3$ (IU13989 X Δ*khpB* from IU10592) | Str$^R$ | This Study |
| IU14028 | D39 Δ*cps rpsL1 murA*-L-FLAG$^3$ (IU13491 X fusion *murA*-L-FLAG$^3$) | Str$^R$ | This Study |
| IU14030 | D39 Δ*cps rpsL1* Δ*khpA murA*-L-FLAG$^3$ (IU13495 X fusion *murA*-L-FLAG$^3$) | Str$^R$ | This Study |
| IU14082 | D39 Δ*cps rpsL1* Δ*clpC*::$P_c$-*erm murZ*-L-FLAG$^3$ (IU13502 X Δ*clpC*::$P_c$-*erm* from E780) | Str$^R$ Erm$^R$ | This Study |
| IU14084 | D39 Δ*cps rpsL1* Δ*murA*::$P_c$-*erm murZ*-L-FLAG$^3$ (IU13502 X Δ*murA*::$P_c$-*erm* from E765) | Str$^R$ Erm$^R$ | This Study |
| IU14086 | D39 Δ*cps rpsL1* Δ*clpC*::$P_c$-*erm murA*-L-FLAG$^3$ (IU14028 X Δ*clpC*::$P_c$-*erm* from E780) | Str$^R$ Erm$^R$ | This Study |
| IU14088 | D39 Δ*cps rpsL1* Δ*murZ*::$P_c$-*erm murA*-L-FLAG$^3$ (IU14028 X Δ*murZ*::$P_c$-*erm* from E767) | Str$^R$ Erm$^R$ | This Study |
| IU14210 | D39 Δ*cps rpsL1 murZ*(I265V) (IU13396 X *murZ*(I265V) amplicon from EL59) | Str$^R$ | This Study |
| IU14234 | D39 Δ*cps rpsL1 murZ*(I265V) Δ*gpsB<>aad9* (IU14210 X Δ*gpsB<>aad9* amplicon from IU4888) | Str$^R$ Spc$^R$ | This Study |
| IU14270 | D39 Δ*cps* Δ*mraY<>aad9//*Δ*bgaA*::*kan*-t1t2-$P_{Zn}$-*mraY*$^+$ (IU11083 X fusion Δ*mraY<>aad9*) | Spc$^R$ Kan$^R$ | This Study |
| IU14272 | D39 Δ*cps* Δ*uppS<>aad9//* Δ*bgaA*::*kan*-t1t2-$P_{Zn}$-*uppS*$^+$ (IU11094 X fusion Δ*mraY<>aad9*) | Spc$^R$ Kan$^R$ | This Study |
| IU14274 | D39 Δ*cps ΔmurG<>aad9//*Δ*bgaA*::*kan*-t1t2-$P_{Zn}$-RBS$^{ftsA}$-*murG*$^+$ (IU11049 X fusion *ΔmurG<>aad9*) | Spc$^R$ Kan$^R$ | This Study |
| IU14312 | D39 Δ*cps rpsL1* Δ*bgaA*::*tet*- $P_{Zn}$-RBS$^{ftsA}$-*pbp1a*$^+$ (IU1824 X fusion Δ*bgaA*::*tet*-$P_{Zn}$- RBS$^{ftsA}$-*pbp1a*$^+$) | Str$^R$ Tet$^R$ | This Study |
| IU14318 | D39 Δ*cps rpsL1* Δ*bgaA*::*tet*-$P_{Zn}$-RBS$^{ftsA}$-*pbp2a*$^+$ (IU1824 X fusion Δ*bgaA*::*tet*-$P_{Zn}$-RBS$^{ftsA}$-*pbp2a*$^+$) | Str$^R$ Tet$^R$ | (Cleverley *et al.*, 2019) |
| IU14738 | D39 Δ*cps rpsL1 iht*-$L_6$-*mapZ* markerless | Str$^R$ | (Perez *et al.*, 2019) |
| IU14974 | D39 Δ*cps rpsL1* Δ*bgaA*::*kan*-t1t2-$P_{Zn}$-RBS$^{ftsA}$-*stkP*$^+$ (IU1824 X fusion Δ*bgaA*::*kan*-t1t2-$P_{Zn}$-*stkP*$^+$) | Str$^R$ Kan$^R$ | This Study |
| IU15124 | D39 Δ*cps rpsL1 murZ*(I265V) Δ*gpsB<>aad9* (IU14210 X Δ*gpsB<>aad9* amplicon from IU4888) | Str$^R$ Spc$^R$ | This Study |
| IU15143 | D39 Δ*cps rpsL1 murA*(D281Y) (IU13491 X fusion amplicon *murA*(D281Y)) | Str$^R$ | This Study |
| IU15145 | D39 Δ*cps rpsL1 murA*(E282Y) (IU13491 X fusion amplicon *murA*(E282Y)) | Str$^R$ | This Study |

| | | | |
|---|---|---|---|
| IU15355 | D39 Δ*cps rpsL1* Δ*rodZ*<>*aad9*//Δ*bgaA*::*tet*-$P_{Zn}$-RBS$^{ftsA}$-*rodZ*$^{+}$ (IU9613 X Δ*rodZ*<>*aad9* from IU9931) | Str$^{R}$ Tet$^{R}$ Spc$^{R}$ | This Study |
| IU15357 | D39 Δ*cps rpsL1* Δ*rodZ*::$P_{c}$-*aad9*//Δ*bgaA*::*tet*-$P_{Zn}$-RBS$^{ftsA}$-*rodZ*$^{+}$ (IU9613 X Δ*rodZ*::$P_{c}$-*aad9* from IU6987) | Str$^{R}$ Tet$^{R}$ Spc$^{R}$ | This Study |
| IU15361 | D39 Δ*cps rpsL1* Δ*rodZ*::$P_{c}$-*aad9*//Δ*bgaA*::*tet*-$P_{Zn}$-RBS$^{ftsA}$-*ftsA*$^{+}$ (IU12310 X Δ*rodZ*::$P_{c}$-*aad9* from IU6987) | Str$^{R}$ Tet$^{R}$ Spc$^{R}$ | This Study |
| IU15371 | D39 Δ*cps* Δ*rodZ*::$P_{c}$-*aad9*//Δ*bgaA*::*tet*-$P_{Zn}$-RBS$^{ftsA}$-*rodZ*$^{+}$ (IU9765 X Δ*rodZ*::$P_{c}$-*aad9* from IU6987) | Tet$^{R}$ Spc$^{R}$ | This Study |
| IU15386 | D39 Δ*cps rpsL1* Δ*rodZ*::$P_{c}$-*aad9* Δ*bgaA*::*tet*-$P_{ftsA}$-*ftsA* (IU12719 X Δ*rodZ*::$P_{c}$-*aad9* from IU6987) | Str$^{R}$ Tet$^{R}$ Spc$^{R}$ | This Study |
| IU15531 | D39 Δ*cps rpsL1* Δ*khpA* Δ*rodZ*::$P_{c}$-*erm* (IU9036 X Δ*rodZ*::$P_{c}$-*erm* from E655) | Str$^{R}$ Erm$^{R}$ | This Study |
| IU15636 | D39 Δ*cps rpsL1* Δ*rodZ*::$P_{c}$-*erm*//Δ*bgaA*::*tet*-$P_{Zn}$-RBS$^{ftsA}$-*rodZ*$^{+}$ (IU9613 X Δ*rodZ*::$P_{c}$-*erm* from E655) | Str$^{R}$ Tet$^{R}$ Erm$^{R}$ | This Study |
| IU15641 | D39 Δ*cps* Δ*rodZ*::$P_{c}$-*erm*//Δ*bgaA*::*tet*-$P_{ftsA}$-*ftsA* (IU12712 X Δ*rodZ*::$P_{c}$-*erm* from E655) | Str$^{R}$ Tet$^{R}$ Erm$^{R}$ | This Study |
| IU15860 | D39 Δ*cps rpsL1* Δ*bgaA*::*kan*-t1t2-$P_{Zn}$-RBS$^{ftsA}$ -*murZ*$^{+}$ Δ*gpsB*<>*aad9* (IU13393 X Δ*gpsB*<>*aad9* amplicon from IU4888) | Kan$^{R}$ Str$^{R}$ Spc$^{R}$ | This Study |
| IU15862 | D39 Δ*cps rpsL1* Δ*bgaA*::*kan*-t1t2-$P_{Zn}$-RBS$^{ftsA}$ -*murA*$^{+}$ Δ*gpsB*<>*aad9* (IU13395 X Δ*gpsB*<>*aad9* amplicon from IU4888) | Kan$^{R}$ Str$^{R}$ Spc$^{R}$ | This Study |
| IU15873 | D39 Δ*bgaA*::*tet*-$P_{Zn}$-RBS$^{ftsA}$-*gpsB*$^{+}$ (IU1690 X Δ*bgaA*::*tet*-$P_{Zn}$-RBS$^{ftsA}$-*gpsB*$^{+}$ from IU11286) | Tet$^{R}$ | This Study |
| IU15875 | D39 *rpsL1* Δ*bgaA*::*tet*-$P_{Zn}$-RBS$^{ftsA}$-*gpsB*$^{+}$ (IU1781 X Δ*bgaA*::*tet*-$P_{Zn}$-RBS$^{ftsA}$-*gpsB*$^{+}$ from IU11286) | Str$^{R}$ Tet$^{R}$ | This Study |
| IU15877 | D39 Δ*cps rpsL* Δ*bgaA*::*tet*-$P_{Zn}$-RBS$^{ftsA}$-*gpsB*$^{+}$ (IU1824 X Δ*bgaA*::*tet*-$P_{Zn}$-RBS$^{ftsA}$-*gpsB*$^{+}$ from IU11286) | Str$^{R}$ Tet$^{R}$ | This Study |
| IU15879 | D39 Δ*bgaA*::*kan*-t1t2-$P_{Zn}$-RBS$^{ftsA}$-*murZ*$^{+}$ (IU1690 X Δ*bgaA*::*kan*-t1t2-$P_{Zn}$-*murZ*$^{+}$ amplicon from IU11077) | Kan$^{R}$ | This Study |
| IU15880 | D39 Δ*bgaA*::*kan*-t1t2-$P_{Zn}$-RBS$^{ftsA}$-*murA*$^{+}$ (IU1690 X Δ*bgaA*::*kan*-t1t2-$P_{Zn}$-*murA*$^{+}$ amplicon from IU11079) | Kan$^{R}$ | This Study |
| IU15882 | D39 *rpsL1* Δ*bgaA*::*kan*-t1t2-$P_{Zn}$-RBS$^{ftsA}$-*murZ*$^{+}$ (IU1781 X Δ*bgaA*::*kan*-t1t2-$P_{Zn}$-*murZ*$^{+}$ amplicon from IU11077) | Str$^{R}$ Kan$^{R}$ | This Study |
| IU15884 | D39 *rpsL1* Δ*bgaA*::*kan*-t1t2-$P_{Zn}$-RBS$^{ftsA}$-*murA*$^{+}$ (IU1781 X Δ*bgaA*::*kan*-t1t2-$P_{Zn}$-*murA*$^{+}$ amplicon from IU11079) | Str$^{R}$ Kan$^{R}$ | This Study |
| IU15899 | D39 *rpsL1* Δ*murZ*::$P_{c}$-[*kan-rpsL*$^{+}$] (IU1781 X Δ*murZ*::$P_{c}$-[*kan-rpsL*$^{+}$] from K767) | Kan$^{R}$ | This Study |
| IU15917 | D39 *rpsL1 murZ*(D280Y) (IU15899 X *murZ*(D280Y) from IU13438) | Str$^{R}$ | This Study |

| | | | |
|---|---|---|---|
| IU15939 | D39 Δ*cps rpsL1 murZ*(C116S) (IU13396 X fusion *murZ*(C116S)) | $Str^R$ | This Study |
| IU15941 | D39 Δ*cps rpsL1 murZ*(C116S)-L-$FLAG^3$ (IU13396 X fusion *murZ*(C116S)-L-$FLAG^3$) | $Str^R$ | This Study |
| IU15943 | D39 Δ*cps rpsL1* Δ*bgaA*::*kan*-t1t2-$P_{Zn}$- $RBS^{ftsA}$-*murZ*(C116S) (IU1824 X fusion Δ*bgaA*::*kan*-t1t2-$P_{Zn}$-$RBS^{ftsA}$-*murZ*(C116S)) | $Str^R$ $Kan^R$ | This Study |
| IU15949 | D39 Δ*cps rpsL1 murA*(C120S) (IU13491 X fusion *murA*(C120S)) | $Str^R$ | This Study |
| IU15951 | D39 Δ*cps rpsL1 murA*(C120S)-L-$FLAG^3$ (IU13491 X fusion *murA*(C120S)-L-$FLAG^3$) | $Str^R$ | This Study |
| IU15954 | D39 Δ*cps rpsL1* Δ*bgaA*::*kan*-t1t2-$P_{Zn}$-$RBS^{ftsA}$ -*murA*(C120S) (IU1824 X fusion Δ*bgaA*::*kan*-t1t2-$P_{Zn}$-$RBS^{ftsA}$-*murA*(C120S)) | $Str^R$ $Kan^R$ | This Study |
| IU15955 | D39 Δ*cps rpsL1* Δ*bgaA*::tet-Pzn-*$phpP^+$* (IU1824 X *bgaA*::tet-Pzn-*$phpP^+$* amplicon from IU8742) | $Tet^R$ | This Study |
| IU15983 | D39 Δ*cps rpsL1 murA*-L-$FLAG^3$// Δ*bgaA*::*kan*-t1t2-$P_{Zn}$-$RBS^{ftsA}$-*$murA^+$*-L-$FLAG^3$ (IU14028 X fusion Δ*bgaA*::*kan*-t1t2-$P_{Zn}$-$RBS^{ftsA}$-*$murA^+$*-L-$FLAG^3$) | $Str^R$ $Kan^R$ | This Study |
| IU16176 | D39 Δ*murZ*::$P_c$-*erm* (IU1690 X Δ*murZ*::$P_c$-*erm* amplicon from E767) | $Erm^R$ | This Study |
| IU16178 | D39 Δ*murA*::$P_c$-*erm* (IU1690 X Δ*murA*::$P_c$-*erm* amplicon from E765) | $Erm^R$ | This Study |
| IU16196 | D39 Δ*cps rpsL1* Δ*khpA* Δ*gpsB*<>*aad9* (IU9036 X Δ*gpsB*<>*aad9* from IU4888) | $Str^R$ $Spc^R$ | This Study |
| IU16259 | D39 Δ*cps rpsL1* Δ*murZ*// Δ*bgaA*::*kan*-t1t2-$P_{Zn}$-*$murZ^+$* (IU13536 X Δ*bgaA*::*kan*-t1t2-$P_{Zn}$-*$murZ^+$* amplicon from IU11077) | $Str^R$ $Kan^R$ | This Study |
| IU16262 | D39 Δ*cps rpsL1* Δ*murZ* // Δ*bgaA*::*kan*-t1t2-$P_{Zn}$-*$murA^+$* (IU13536 X Δ*bgaA*::*kan*-t1t2-$P_{Zn}$-*$murA^+$* amplicon from IU11079) | $Str^R$ $Kan^R$ | This Study |
| IU16265 | R6 Δ*murZ*::$P_c$-*erm* (EL59 X Δ*murZ*::$P_c$-*erm* amplicon from E767) | $Erm^R$ | This Study |
| IU16267 | R6 Δ*murA*::$P_c$-*erm* (EL59 X Δ*murZ*::$P_c$-*erm* amplicon from E765) | $Erm^R$ | This Study |
| IU16295 | D39 Δ*cps rpsL1* Δ[*spd_1029-1030*]::$P_c$-*erm* Δ*bgaA*::*kan*-t1t2-$P_{Zn}$-$RBS^{ftsA}$-*$murZ^+$* (IU13393 X Δ[*spd_1029-1030*]::$P_c$-*erm* amplicon from IU8108) | $Kan^R$ $Erm^R$ | This Study |
| IU16298 | D39 Δ*cps rpsL1* Δ[*spd_1031-1037*]::$P_c$-*erm* Δ*bgaA*::*kan*-t1t2-$P_{Zn}$-$RBS^{ftsA}$-*$murZ^+$* (IU13393 X Δ[*spd_1031-1037*]::$P_c$-*erm* amplicon from IU7824) | $Kan^R$ $Erm^R$ | This Study |
| IU16330 | D39 Δ*cps rpsL1* Δ*murZ*// Δ*bgaA::kan*- t1t2-$P_{Zn}$-$RBS^{ftsA}$ -*murZ* Δ*murA*::$P_c$-*erm* (IU16259 X Δ*murA*::$P_c$-*erm* from E765) | $Erm^R$ $Kan^R$ | This Study |
| IU16332 | D39 Δ*cps rpsL1* Δ*murZ* Δ*murA*::$P_c$-*erm*// Δ*bgaA::kan*- t1t2-$P_{Zn}$-$RBS^{ftsA}$ -*murA* (IU16262 X Δ*murA*::$P_c$-*erm* from E765) | $Erm^R$ $Kan^R$ | This Study |

| | | | |
|---|---|---|---|
| IU16334 | D39 Δ*cps rpsL1* Δ*bgaA::kan*- t1t2-P$_{Zn}$-RBS$^{ftsA}$ -*murZ*(D280Y) (IU1824 X fusion Δ*bgaA::kan*- t1t2-P$_{Zn}$-RBS$^{ftsA}$ -*murZ*(D280Y)) | Str$^R$ Kan$^R$ | This Study |
| IU16336 | D39 Δ*cps rpsL1 murZ*(D280Y)// Δ*bgaA::kan*- t1t2-P$_{Zn}$-RBS$^{ftsA}$ -*murZ*(D280Y) (IU13438 X fusion Δ*bgaA::kan*- t1t2-P$_{Zn}$-RBS$^{ftsA}$ -*murZ*(D280Y)) | Str$^R$ Kan$^R$ | This Study |
| IU16370 | D39 Δ*cps rpsL1* Δ*gpsB*<>*aad9*//Δ*bgaA*::*tet*-P$_{Zn}$-RBS$^{ftsA}$-*gpsB*$^+$ (IU15877 X Δ*gpsB*<>*aad9* amplicon from IU4888) | Spc$^R$ Tet$^R$ | This Study |
| IU16883 | D39 Δ*cps rpsL1* Δ*stkP*::P$_c$-*erm sup1* (IU1824 X Δ*stkP*::P$_c$-*erm* from IU7923) | Erm$^R$ | This Study |
| IU16885, IU16895 | D39 Δ*cps rpsL1 murZ*(D280Y) Δ*stkP*::P$_c$-*erm* (IU13438 X Δ*stkP*::P$_c$-*erm* amplicon from IU7923) | Str$^R$ Erm$^R$ | This Study |
| IU16897 | D39 Δ*cps rpsL1* Δ*bgaA*::*kan*-t1t2-P$_{Zn}$-RBS$^{ftsA}$-*murZ*$^+$ Δ*stkP*::P$_c$-*erm* (IU13393 X Δ*stkP*::P$_c$-*erm* amplicon from IU7923) | Erm$^R$ Kan$^R$ | This Study |
| IU16910 | D39 Δ*cps rpsL1* Δ*khpA* Δ*stkP*::P$_c$-*erm* (IU9036 X Δ*stkP*::P$_c$-*erm* amplicon from IU7923) | Str$^R$ Erm$^R$ | This Study |
| IU16912 | D39 Δ*cps rpsL1* Δ*khpB* Δ*stkP*::P$_c$-*erm* (IU10592 X Δ*stkP*::P$_c$-*erm* amplicon from IU7923) | Str$^R$ Erm$^R$ | This Study |
| IU16915 | D39 Δ*cps rpsL1* Δ*bgaA*::*kan*-t1t2-P$_{Zn}$-RBS$^{ftsA}$-*murA*$^+$ Δ*stkP*::P$_c$-*erm* (IU13395 X Δ*stkP*::P$_c$-*erm* amplicon from IU7923) | Erm$^R$ Kan$^R$ | This Study |
| IU16933, IU16934 | D39 Δ*cps rpsL1* Δ*stkP*::P$_c$-*erm*//Δ*bgaA*::*kan*-t1t2-P$_{Zn}$-RBS$^{ftsA}$-*stkP*$^+$ (IU14974 X Δ*stkP*::P$_c$-*erm* amplicon from IU7923) | Erm$^R$ Kan$^R$ | This Study |
| IU17134 | D39 Δ*cps rpsL1* Δ*clpE*::P$_c$-[*kan-rpsL*$^+$] (IU1824 X fusion Δ*clpE*::P$_c$-[*kan-rpsL*$^+$) | Str$^R$ Kan$^R$ | This Study |
| IU17136 | D39 Δ*cps rpsL1* Δ*clpL*::P$_c$-[*kan-rpsL*$^+$] (IU1824 X fusion Δ*clpL*::P$_c$-[*kan-rpsL*$^+$]) | Str$^R$ Kan$^R$ | This Study |
| IU17138 | D39 Δ*cps rpsL1* Δ*clpP*::P$_c$-[*kan-rpsL*$^+$] (IU1824 X fusion Δ*clpP*::P$_c$-[*kan-rpsL*$^+$]) | Str$^R$ Kan$^R$ | This Study |
| IU17150 | D39 Δ*cps rpsL1* Δ*clpE*::P$_c$-*erm murZ*-L-FLAG$^3$ (IU13502 X fusion Δ*clpE*::P$_c$-*erm*) | Str$^R$ Erm$^R$ | This Study |
| IU17152 | D39 Δ*cps rpsL1* Δ*clpL*::P$_c$-*erm murZ*-L-FLAG$^3$ (IU13502 X fusion Δ*clpL*::P$_c$-*erm*) | Str$^R$ Erm$^R$ | This Study |
| IU17154 | D39 Δ*cps rpsL1* Δ*clpP*::P$_c$-*erm murZ*-L-FLAG$^3$ (IU13502 X fusion Δ*clpP*::P$_c$-*erm*) | Str$^R$ Erm$^R$ | This Study |
| IU17158 | D39 Δ*cps rpsL1* Δ*clpE*::P$_c$-*erm murA*-L-FLAG$^3$ (IU14028 X fusion Δ*clpE*::P$_c$-*erm*) | Str$^R$ Erm$^R$ | This Study |
| IU17160 | D39 Δ*cps rpsL1* Δ*clpL*::P$_c$-*erm murA*-L-FLAG$^3$ (IU14028 X fusion Δ*clpL*::P$_c$-*erm*) | Str$^R$ Erm$^R$ | This Study |
| IU17162 | D39 Δ*cps rpsL1* Δ*clpP*::P$_c$-*erm murA*-L-FLAG$^3$ (IU14028 X fusion Δ*clpP*::P$_c$-*erm*) | Str$^R$ Erm$^R$ | This Study |
| IU17170 | D39 Δ*cps rpsL1 murZ*-HA (IU13396 X fusion *murZ*-HA) | Str$^R$ | This Study |
| IU17469, IU17475 | D39 Δ*cps rpsL1 murZ*(I265V) Δ*stkP*::P$_c$-*erm* (IU14210 X Δ*stkP*::P$_c$-*erm* amplicon from IU7923) | Str$^R$ Erm$^R$ | This Study |

| | | | |
|---|---|---|---|
| IU17603 | D39 Δ*cps rpsL1* Δ*bgaA*::*tet*-$P_{ftsA}$-*ftsA* Δ*pbp2b*<>*aad9* (IU12719 X Δ*pbp2b*<>*aad9* from IU7397) | $Str^R$ $Tet^R$ $Spc^R$ | This Study |
| IU17605 | D39 Δ*cps* Δ*bgaA*::*tet*-$P_{ftsA}$-*ftsA* Δ*rodZ*::$P_c$-*aad9* (IU12712 X Δ*rodZ*::$P_c$-*aad9* from IU6987) | $Str^R$ $Tet^R$ $Spc^R$ | This Study |
| IU17607 | D39 Δ*cps* Δ*bgaA*::*tet*-$P_{ftsA}$-*ftsA* Δ*pbp2b*<>*aad9* (IU12712 X Δ*pbp2b*<>*aad9* from IU7397) | $Str^R$ $Tet^R$ $Spc^R$ | This Study |
| IU17609 | D39 Δ*cps* Δ*bgaA*::*tet*-$P_{Zn}$-$RBS^{ftsA}$-$ftsA^+$ Δ*pbp2b*<>*aad9* (IU12307 X Δ*rodZ*::$P_c$-*aad9* from IU6987) | $Str^R$ $Tet^R$ $Spc^R$ | This Study |
| IU17619 | D39 Δ*cps rpsL1 murZ*(E190A E192A) (IU13396 X fusion *murZ*(E192A)) | $Str^R$ | This Study |
| IU17622 | D39 Δ*cps rpsL1 murZ*(E192A) (IU13396 X fusion *murZ*(E192A)) | $Str^R$ | This Study |
| IU17623 | D39 Δ*cps rpsL1 murZ*(D195A) (IU13396 X fusion *murZ*(D195A)) | $Str^R$ | This Study |
| IU17627 | D39 Δ*cps rpsL1 murZ*(E259A) (IU13396 X fusion *murZ*(E259A)) | $Str^R$ | This Study |
| IU17764 | D39 Δ*cps rpsL1* F-*murZ* (IU13396 X fusion F-*murZ*) | $Str^R$ | This Study |
| IU17766 | D39 Δ*cps rpsL1* HA-*murZ* (IU13396 X fusion HA-*murZ*) | $Str^R$ | This Study |
| IU17768 | D39 Δ*cps rpsL1* F-*murA* (IU13491 X fusion F-*murA*) | $Str^R$ | This Study |
| IU17770 | D39 Δ*cps rpsL1* HA-*murA* (IU13491 X fusion HA-*murA*) | $Str^R$ | This Study |
| IU17838 | D39 Δ*cps rpsL1 iht*-$L_6$-*murZ* with spontaneous L88F mutation in *iht* (IU13396 X fusion *iht*-$L_6$-*murZ*) | $Str^R$ | This Study |
| IU17840 | D39 Δ*cps rpsL1 iht*-$L_6$-*murZ* with spontaneous Z21F mutation in *iht* (IU13396 X fusion *iht*-$L_6$-*murZ*) | $Str^R$ | This Study |
| IU17841 | D39 Δ*cps rpsL1 iht*-$L_6$-*murA* (IU13491 X fusion *iht*-$L_6$-*murA*) | $Str^R$ | This Study |
| IU17865 | D39 Δ*cps rpsL1* Δ*clpP*::$P_c$-*erm iht*-$L_6$-*murZ* with spontaneous L88F mutation in *iht* (IU17838 X Δ*clpP*::$P_c$-*erm* from IU17154) | $Str^R$ $Erm^R$ | This Study |
| IU17867 | D39 Δ*cps rpsL1* Δ*clpP*::$P_c$-*erm iht*-$L_6$-*murZ* with spontaneous Z21F mutation in *iht* (IU17840 X Δ*clpP*::$P_c$-*erm* from IU17154) | $Str^R$ $Erm^R$ | This Study |
| IU17869 | D39 Δ*cps rpsL1 iht*-$L_6$-*murA* Δ*clpP*::$P_c$-*erm* (IU17841 X Δ*clpP*::$P_c$-*erm* from IU17154) | $Str^R$ $Erm^R$ | This Study |
| IU17957 | D39 Δ*cps rpsL1 murZ*-L-$FLAG^3$-$P_c$-*erm* Δ*bgaA*::*kan*-t1t2-$P_{Zn}$-$RBS^{ftsA}$-$stkP^+$ (IU14974 X *murZ*-L-$FLAG^3$-$P_c$-*erm* from IU13249) | $Erm^R$ $Kan^R$ | This Study |
| IU17959 | D39 Δ*cps rpsL1 murA*-L-$FLAG^3$-$P_c$-*erm* Δ*bgaA*::*kan*-t1t2-$P_{Zn}$-$RBS^{ftsA}$-$stkP^+$ (IU14974 X *murA*-L-$FLAG^3$-$P_c$-*erm* from IU13251) | $Erm^R$ $Kan^R$ | This Study |

| | | | |
|---|---|---|---|
| IU17961 | D39 Δ*cps rpsL1 murZ*-L-FLAG$^{3}$-P$_{c}$-*erm* Δ*bgaA*::*tet*-P$_{Zn}$-RBS$^{ftsA}$-*phpP*$^{+}$ (IU15955 X *murZ*-L-FLAG$^{3}$-P$_{c}$-*erm* from IU13249) | Erm$^{R}$ Tet$^{R}$ | This Study |
| IU17963 | D39 Δ*cps rpsL1 murA*-L-FLAG$^{3}$-P$_{c}$-*erm* Δ*bgaA*::*tet*-P$_{Zn}$-RBS$^{ftsA}$-*phpP*$^{+}$ (IU15955 X *murA*-L-FLAG$^{3}$-P$_{c}$-*erm* from IU13251) | Erm$^{R}$ Tet$^{R}$ | This Study |
| IU18555 | D39 Δ*cps rpsL1* Δ*bgaA*::*tet*-P$_{Zn}$-RBS$^{ftsA}$-*stkP*$^{+}$ (IU1824 X fusion Δ*bgaA*::*tet*-P$_{Zn}$-RBS$^{ftsA}$-*stkP*$^{+}$) | Tet$^{R}$ | This Study |
| IU18643 | D39 Δ*cps rpsL1 phpP*$^{+}$-P$_{c}$-[*kan-rpsL*$^{+}$]-*stkP*$^{+}$ //Δ*bgaA*::*tet*-P$_{Zn}$-RBS$^{ftsA}$-*stkP*$^{+}$ (IU18555 X *phpP*$^{+}$-P$_{c}$-[*kan-rpsL*$^{+}$]-*stkP*$^{+}$ from IU7673) | Kan$^{R}$ Tet$^{R}$ | This Study |
| IU18665 | D39 Δ*cps rpsL1* Δ*stkP* markerless//Δ*bgaA*::*tet*-P$_{Zn}$-RBS$^{ftsA}$-*stkP*$^{+}$ (IU18643 X fusion Δ*stkP* markerless) | Str$^{R}$ Tet$^{R}$ | This Study |
| IU19079 | D39 Δ*cps rpsL1 murZ*(D280Y)-L-FLAG$^{3}$-P$_{c}$-*erm* Δ*stkP* markerless//Δ*bgaA*::*tet*-P$_{Zn}$-RBS$^{ftsA}$-*stkP*$^{+}$ (IU18665 X fusion *murZ*(D280Y)-L-FLAG$^{3}$-P$_{c}$-*erm*) | Erm$^{R}$ Str$^{R}$ Tet$^{R}$ | This Study |
| IU19081 | D39 Δ*cps rpsL1 murZ*-L-FLAG$^{3}$-P$_{c}$-*erm* Δ*stkP* markerless//Δ*bgaA*::*tet*-P$_{Zn}$-RBS$^{ftsA}$-*stkP*$^{+}$ (IU18665 X *murZ*-L-FLAG$^{3}$-P$_{c}$-*erm* from IU13249) | Erm$^{R}$ Str$^{R}$ Tet$^{R}$ | This Study |
| IU19821 | D39 Δ*cps rpsL1* Δ*spd_0567*::P$_{c}$-[*sacB-kan-rpsL*$^{+}$] (IU1824 X fusion Δ*spd_0567*::P$_{c}$-[*sacB-kan-rpsL*$^{+}$]) | Kan$^{R}$ | This Study |
| IU19835 | D39 Δ*cps rpsL1 spd_0567*$^{+}$ (IU19821 X *spd_0567*$^{+}$ from IU1690) | Str$^{R}$ | This Study |

| **Primers used to construct strains** | | | |
|---|---|---|---|
| Primer | Sequence (5' to 3') | Template[c] | Amplicon Product |
| **For construction of E740 (Δ[*phpP-stkP*]::P$_{c}$-*erm*)** | | | |
| P1485 | CCAAGCCTTGTTGGAGGCGAATAATTCCCT | D39 | 5' fragment with 60 bp of 5' *phpP* |
| P1486 | CATTATCCATTAAAAATCAAACGGATCCTAGA CATA GTCTTGGTTATTTGTTCGTTTCTG | | |
| Kan rpsL forward | TAGGATCCGTTTGATTTTTAATGGATAATG | Pc-*erm* cassette[d] | Pc-*erm* |
| Kan rpsL reverse | GGGCCCCTTTCCTTATGCTTTTG | | |
| P1497 | CAAAAGCATAAGGAAAGGGGCCCAATAAGAC TAG AGTCAAGATTTCAATCTACAAACCTA | D39 | 3' fragment with 60 bp of 3' *stkP* |
| P1496 | CAATACCAAGGCGACAGAAGTTCCTGCCCC | | |
| **For construction of E765 (Δ*murA*::P$_{c}$-*erm*)** | | | |
| P1558 | TCAGGAGACTACAGGTGGTTCTTCCGATGT | D39 | 5' fragment with 60 bp of 5' *murA* |
| P1560 | CATTATCCATTAAAAATCAAACGGATCCTACT CGATCGTCACGCTTCCTACCAGACGATT | | |
| Kan rpsL forward | TAGGATCCGTTTGATTTTTAATGGATAATG | P$_{c}$-*erm* cassette[d] | P$_{c}$-*erm* |
| Kan rpsL reverse | GGGCCCCTTTCCTTATGCTTTTG | | |

| | | | |
|---|---|---|---|
| P1561 | AAACGTCCAAAAGCATAAGGAAAGGGGCCC AAGTTGGCGCAGCTAGGTGCTAAGATTCAG | D39 | 3' fragment with 60 bp of 3' *murA* |
| P1559 | CTTAGTACCTGTTCTAGCCCTGCTTAAACT | | |
| **For construction of E767 (∆*murZ*::$P_c$-*erm*)** | | | |
| P1554 | GATTTTGTGGTACGACGGGCATGTATAGCG | D39 | 5' fragment with 60 bp of 5' *murZ* |
| P1556 | CATTATCCATTAAAAATCAAACGGATCCTAAC CACTAATAGTGATTTCACCTTGCAGTGG | | |
| Kan rpsL forward | TAGGATCCGTTTGATTTTTAATGGATAATG | $P_c$-*erm* cassette[d] | $P_c$-*erm* |
| Kan rpsL reverse | GGGCCCCTTTCCTTATGCTTTTG | | |
| P1557 | AAACGTCCAAAAGCATAAGGAAAGGGGCCCT CTGATATTATCGAAAAATTACGTAATTTA | D39 | 3' fragment with 60 bp of 3' *murZ* |
| P1555 | TGAACCTGAAATCCCCCTGTAACCAGAACT | | |
| **For construction of E780 (∆*clpC*::$P_c$-*erm*)** | | | |
| P1663 | GACTAGAGCACGTCAGTTATGCCTATGGTC | D39 | 5' fragment with 60 bp of 5' *clpC* |
| P1665 | CATTATCCATTAAAAATCAAACGGATCCTAAT GTCCAGCAACCATGTAGGCACTTTCGAT | | |
| Kan rpsL forward | TAGGATCCGTTTGATTTTTAATGGATAATG | $P_c$-*erm* cassette[d] | $P_c$-*erm* |
| Kan rpsL reverse | GGGCCCCTTTCCTTATGCTTTTG | | |
| P1666 | AAACGTCCAAAAGCATAAGGAAAGGGGCCC GCAGGCAGCATACTTAAGATTGGTGTCAAA | D39 | 3' fragment with 60 bp of 3' *clpC* |
| P1664 | AAATCCACTGTTACATCCTGATATCGCCAA | | |
| **For construction of K765 (∆*murA*::$P_c$-[*kan-rpsL*$^+$])** | | | |
| P1558 | TCAGGAGACTACAGGTGGTTCTTCCGATGT | D39 | 5' fragment with 60 bp of 5' *murA* |
| P1560 | CATTATCCATTAAAAATCAAACGGATCCTACT CGATCGTCACGCTTCCTACCAGACGATT | | |
| Kan rpsL forward | TAGGATCCGTTTGATTTTTAATGGATAATG | $P_c$-[*kan-rpsL*$^+$] cassette[d] | $P_c$-[*kan-rpsL*$^+$] |
| Kan rpsL reverse | GGGCCCCTTTCCTTATGCTTTTG | | |
| P1561 | AAACGTCCAAAAGCATAAGGAAAGGGGCCC AAGTTGGCGCAGCTAGGTGCTAAGATTCAG | D39 | 3' fragment with 60 bp of 3' *murA* |
| P1559 | CTTAGTACCTGTTCTAGCCCTGCTTAAACT | | |
| **For construction of K767 (∆*murZ*::$P_c$-[*kan-rpsL*$^+$])** | | | |
| P1554 | GATTTTGTGGTACGACGGGCATGTATAGCG | D39 | 5' fragment with 60 bp of 5' *murZ* |
| P1556 | CATTATCCATTAAAAATCAAACGGATCCTAAC CACTAATAGTGATTTCACCTTGCAGTGG | | |
| Kan rpsL forward | TAGGATCCGTTTGATTTTTAATGGATAATG | $P_c$-[*kan-rpsL*$^+$] cassette[d] | $P_c$-[*kan-rpsL*$^+$] |
| Kan rpsL reverse | GGGCCCCTTTCCTTATGCTTTTG | | |
| P1557 | AAACGTCCAAAAGCATAAGGAAAGGGGCCCT CTGATATTATCGAAAAATTACGTAATTTA | D39 | 3' fragment with 60 bp of 3' *murZ* |
| P1555 | TGAACCTGAAATCCCCCTGTAACCAGAACT | | |
| **For construction of K779 (∆*clpC*::$P_c$-[*kan-rpsL*$^+$])** | | | |
| P1663 | GACTAGAGCACGTCAGTTATGCCTATGGTC | D39 | |

| | | | |
|---|---|---|---|
| P1665 | CATTATCCATTAAAAATCAAACGGATCCTAAT GTCCAGCAACCATGTAGGCACTTTCGAT | | 5' fragment with 60 bp of 5' *clpC* |
| Kan rpsL forward | TAGGATCCGTTTGATTTTTAATGGATAATG | $P_c$-[*kan-rpsL*$^+$] cassette[d] | $P_c$-[*kan-rpsL*$^+$] |
| Kan rpsL reverse | GGGCCCCTTTCCTTATGCTTTTG | | |
| P1666 | AAACGTCCAAAAGCATAAGGAAAGGGGCCC GCAGGCAGCATACTTAAGATTGGTGTCAAA | D39 | 3' fragment with 60 bp of 3' *clpC* |
| P1664 | AAATCCACTGTTACATCCTGATATCGCCAA | | |
| **For construction of K787 (Δ*ireB*(*spd_0180*)::$P_c$-[*kan-rpsL*$^+$])** | | | |
| P1711 | GAGTGTCAATGAAGTTCTCAATCTGATTATGG AAACACC | D39 | 5' upstream of *ireB* + 30 bp of 5' *ireB* |
| P1713 | CATTATCCATTAAAAATCAAACGGATCCTAAA AACGTACTGTTTCTTCAGTAAATCCCAT | | |
| Kan rpsL forward | TAGGATCCGTTTGATTTTTAATGGATAATG | $P_c$-[*kan-rpsL*$^+$] cassette[f] | $P_c$-[*kan-rpsL*$^+$] |
| Kan rpsL reverse | GGGCCCCTTTCCTTATGCTTTTG | | |
| P1714 | AACGTCCAAAAGCATAAGGAAAGGGGCCCTA TCTCAAAGGACAAGGAGTCGATCTATAAC | D39 | 30 bp of 3' *ireB* and downstream of *ireB* |
| P1712 | CCACTGGACGTTCCAACTCTTCCCCATTTC | | |
| **For construction of IU8108 (Δ[*spd_1029-1030*]::$P_c$-*erm*)** | | | |
| P1514 | GCTGGTCAAATCTGGGAGCCTTTTACTGAT | D39 | 5' fragment with 60 bp of 5' *spd_1030* |
| P1513 | CATTATCCATTAAAAATCAAACGGATCCTAAC AAACTTGATCCAAACCAGACTTGG | | |
| Kan rpsL forward | TAGGATCCGTTTGATTTTTAATGGATAATG | $P_c$-*erm* cassette[d] | $P_c$-*erm* |
| Kan rpsL reverse | GGGCCCCTTTCCTTATGCTTTTG | | |
| P1512 | CAAAAGCATAAGGAAAGGGGCCCCGTTGGC GTTTAACTGTGATTATGAA | D39 | 3' fragment with 60 bp of 3' *spd_1029* |
| P1510 | ACCATTGCCACTGCGAACATGGTCTACAGC | | |
| **For construction of IU8742 (Δ*bgaA::tet-P$_{Zn}$-RBS$^{ftsA}$-phpP$^+$*)** | | | |
| TT657 | CGCCCCAAGTTCATCACCAATGACATCAAC | IU8122 | *bgaA'* *tet*-$P_{Zn}$-*RBS*$^{ftsA}$ |
| BR01 | TTCCTTCCTAATCCGATATCTTGTAATAGATTT ATGAACACCTTGTTCATTATCATTATC | | |
| BR02 | AATGAACAAGGTGTTCATAAATCTATTACAAG ATATCGGATTAGGAAGGAACTGACAC | D39 | *phpP*$^+$ |
| BR03 | CAACTGGTTTATGAGAAAGTAAGTTCTTTCAT TCTGCATCCTCCTCGTTCA | | |
| BR04 | ACGAGGAGGATGCAGAATGAAAGAACTTACT TTCTCATAAACCAGTTGCTG | D39 | *bgaA'* to downstream |
| CS121 | GCTTTCTTGAGGCAATTCACTTGGTGC | | |
| **For construction of IU9613 (Δ*bgaA::tet-P$_{Zn}$-RBS$^{ftsA}$-rodZ$^+$*)** | | | |
| TT657 | CGCCCCAAGTTCATCACCAATGACATCAAC | IU8122 | *bgaA'* |

| | | | |
|---|---|---|---|
| TT769 | CCTCTCCAATTGTTTTTTTTCTCATTACATCGC TTCCTCTCTATCTTCCTTGT | | *tet*-P$_{Zn}$-*RBS*$^{ftsA}$ |
| TT770 | GGAAGATAGAGAGGAAGCGATGTAATGAGAA AAAAAACAATTGGAGAGGTTTTAC | D39 | *rodZ*$^{+}$ |
| TT771 | ACTGGTTTATGAGAAAGTAAGTTCTTTTAATTT TTAGTAAAGGTTACAGTGATTTGTCCA | | |
| TT772 | AAATCACTGTAACCTTTACTAAAAATTAAAAG AACTTACTTTCTCATAAACCAGTTGCTG | D39 | *bgaA'* to downstream |
| CS121 | GCTTTCTTGAGGCAATTCACTTGGTGC | | |
| **For construction of IU9805 (Δ*bgaA*::*kan*-t1t2-P$_{Zn}$-*sepF*$^{+}$)** | | | |
| P146 | TGGCCATTCATCGCTGGTCGTGCTGAAAT | IU9689 | 5' Δ*bgaA*::*kan*-t1t2-P$_{Zn}$-RBS$_{ftsA-}$ |
| AJP32 | ACATCGCTTCCTCTCTATCTTCCTTGTTATAA TAGATTTATGAACACCTTGTTCATTATC | | |
| AJP107 | GGAAGATAGAGAGGAAGCGATGTAATGTCTT TAAAAGATAGATTCGATAGATTTATAGAT | D39 | *sepF*$^{+}$ |
| AJP108 | CAACTGGTTTATGAGAAAGTAAGTTCTTTTAT CGTACTCTATTTCGCTTCATATCAAAA | | |
| AJP109 | GATATGAAGCGAAATAGAGTACGATAAAAGA ACTTACTTTCTCATAAACCAGTTGCTG | D39 | *bgaA'* to downstream |
| CS121 | GCTTTCTTGAGGCAATTCACTTGGTGC | | |
| **For construction of IU9992 (Δ*bgaA*::tet-P$_{Zn}$-RBS$^{ftsA}$–*pbp1b*$^{+}$)** | | | |
| P146 | TGGCCATTCATCGCTGGTCGTGCTGAAAT | IU8122 | 5' Δ*bgaA*::tet-t1t2-P$_{Zn}$-RBS$^{ftsA}$ |
| BR74 | TGATTTTGCATGGATTTCCTCACTACATCGCT TCCTCTCTATCTTCCTTGTTATA | | |
| BR73 | AGGAAGATAGAGAGGAAGCGATGTAGTGAG GAAATCCATGCAAAATCAATTAA | D39 | *pbp1b*$^{+}$ |
| BR76 | CAACTGGTTTATGAGAAAGTAAGTTCTTTTAT CGTCTCGCCCTTGAAGAAGAAG | | |
| BR75 | TCTTCAAGGGCGAGACGATAAAAGAACTTAC TTTCTCATAAACCAGTTGCTGC | IU8122 | *bgaA'* to downstream |
| CS121 | GCTTTCTTGAGGCAATTCACTTGGTGC | | |
| **For construction of IU10220 (Δ*bgaA*::*tet*-P$_{Zn}$-RBS$^{ftsA}$-*mreC*$^{+}$)** | | | |
| TT657 | CGCCCCAAGTTCATCACCAATGACATCAAC | IU9613 | *bgaA*' *tet*-P$_{Zn}$-*RBS*$^{ftsA}$ |
| TT865 | GACATATTTTGATTTTTTAAAACGGTTCATTA CATCGCTTCCTCTCTATCTTCCTTGTTA | | |
| TT866 | ACAAGGAAGATAGAGAGGAAGCGATGTAAT GAACCGTTTTAAAAAATCAAAATATGTCAT | D39 | *mreC*$^{+}$ |
| TT867 | AACTGGTTTATGAGAAAGTAAGTTCTTTTATG AATTCCCCACTAATTCTATCACATCTAC | | |
| TT868 | ATGTGATAGAATTAGTGGGGAATTCATAAAA GAACTTACTTTCTCATAAACCAGTTGCTG | | *bgaA'* to downstream |
| CS121 | GCTTTCTTGAGGCAATTCACTTGGTGC | | |
| **For construction of IU11049 (Δ*bgaA*::*kan*-t1t2-P$_{Zn}$-*murG*$^{+}$)** | | | |
| P146 | TGGCCATTCATCGCTGGTCGTGCTGAAAT | IU9805 | 5' containing Δ*bgaA*::*kan*-t1t2-P$_{Zn}$ |
| JQ199 | CCCCCCACCTGTAAAGACAATTTTTTTCATTA CATCGCTTCCTCTCTATCTTCCTTGTTA | | |
| JQ200 | TAACAAGGAAGATAGAGAGGAAGCGATGTAA TGAAAAAAATTGTCTTTACAGGTGGGGGG | D39 | *murG*$^{+}$ |

| | | | |
|---|---|---|---|
| JQ201 | AGCAACTGGTTTATGAGAAAGTAAGTTCTTTT ATGATAAATCTTTTTTCAACAATTGATA | | |
| JQ202 | TATCAATTGTTGAAAAAAGATTTATCATAAAA GAACTTACTTTCTCATAAACCAGTTGCT | IU9805 | *bgaA'* to downstream |
| CS121 | GCTTTCTTGAGGCAATTCACTTGGTGC | | |
| **For construction of IU11077 (Δ*bgaA*::*kan*-t1t2-$P_{Zn}$-*murZ*$^+$)** | | | |
| P146 | TGGCCATTCATCGCTGGTCGTGCTGAAAT | IU9805 | 5' containing Δ*bgaA*::*kan*-t1t2-$P_{Zn}$ |
| JQ222 | TAATCCACCATTGATAACAATTTTTCTCATTA CATCGCTTCCTCTCTATCTTCCTTGTTA | | |
| JQ223 | TAACAAGGAAGATAGAGAGGAAGCGATGTAA TGAGAAAAATTGTTATCAATGGTGGATTA | D39 | *murZ*$^+$ |
| JQ224 | AGCAACTGGTTTATGAGAAAGTAAGTTCTTTT AATCCTCAACAAGTCTAATATCCGCTCC | | |
| JQ225 | GGAGCGGATATTAGACTTGTTGAGGATTAAA AGAACTTACTTTCTCATAAACCAGTTGCT | IU9805 | *bgaA'* to downstream |
| CS121 | GCTTTCTTGAGGCAATTCACTTGGTGC | | |
| **For construction of IU11079 (Δ*bgaA*::*kan*-t1t2-$P_{Zn}$-*murA*$^+$)** | | | |
| P146 | TGGCCATTCATCGCTGGTCGTGCTGAAAT | IU9805 | 5' containing Δ*bgaA*::*kan*-t1t2-$P_{Zn}$ |
| JQ226 | ATCGCCACCTTGAACCACAATTTTATCCATTA CATCGCTTCCTCTCTATCTTCCTTGTTA | | |
| JQ227 | TAACAAGGAAGATAGAGAGGAAGCGATGTAA TGGATAAAATTGTGGTTCAAGGTGGCGAT | D39 | *murA*$^+$ |
| JQ228 | AGCAACTGGTTTATGAGAAAGTAAGTTCTTTT ATTCATCTTCATCATTTGCCTCAATCCG | | |
| JQ229 | CGGATTGAGGCAAATGATGAAGATGAATAAA AGAACTTACTTTCTCATAAACCAGTTGCT | IU9805 | *bgaA'* to downstream |
| CS121 | GCTTTCTTGAGGCAATTCACTTGGTGC | | |
| **For construction of IU11083 (Δ*bgaA*::*kan*-t1t2-$P_{Zn}$-*mraY*$^+$)** | | | |
| P146 | TGGCCATTCATCGCTGGTCGTGCTGAAAT | IU9805 | 5' containing Δ*bgaA*::*kan*-t1t2-$P_{Zn}$ |
| JQ195 | CACAATTCCAGCACTGATGGAAATAAACATTA CATCGCTTCCTCTCTATCTTCCTTGTTA | | |
| JQ196 | TAACAAGGAAGATAGAGAGGAAGCGATGTAA TGTTTATTTCCATCAGTGCTGGAATTGTG | D39 | *mraY*$^+$ |
| JQ197 | AGCAACTGGTTTATGAGAAAGTAAGTTCTTTT ACATCAAATACAAAATTGCGAGGGTCAG | | |
| JQ198 | CTGACCCTCGCAATTTTGTATTTGATGTAAAA GAACTTACTTTCTCATAAACCAGTTGCT | IU9805 | *bgaA'* to downstream |
| CS121 | GCTTTCTTGAGGCAATTCACTTGGTGC | | |
| **For construction of IU11094 (Δ*bgaA*::*kan*-t1t2-$P_{Zn}$-*uppS*$^+$)** | | | |
| P146 | TGGCCATTCATCGCTGGTCGTGCTGAAAT | IU9805 | 5' containing Δ*bgaA*::*kan*-t1t2-$P_{Zn}$ |
| JQ191 | AGCCTTATCTTTCTTAAAAAATCCAAACATTA CATCGCTTCCTCTCTATCTTCCTTGTTA | | |
| JQ192 | TAACAAGGAAGATAGAGAGGAAGCGATGTAA TGTTTGGATTTTTTAAGAAAGATAAGGCT | D39 | *uppS*$^+$ |
| JQ193 | AGCAACTGGTTTATGAGAAAGTAAGTTCTTCT AAACTCCTCCAAATCGGCGATGACGACG | | |
| JQ194 | CGTCGTCATCGCCGATTTGGAGGAGTTTAGA AGAACTTACTTTCTCATAAACCAGTTGCT | IU9805 | *bgaA'* to downstream |
| CS121 | GCTTTCTTGAGGCAATTCACTTGGTGC | | |

| | | | |
|---|---|---|---|
| **For construction of IU11628 (Δ*bgaA*::*kan*-t1t2-$P_{Zn}$-RBS$^{ftsA}$ -*mapZ*$^{+}$)** | | | |
| P146 | TGGCCATTCATCGCTGGTCGTGCTGAAAT | IU9805 | Δ*bgaA*::*kan*-t1t2-$P_{Zn}$-RBS$^{ftsA}$ |
| AJP32 | ACATCGCTTCCTCTCTATCTTCCTTGTTATAAT AGATTTATGAACACCTTGTTCATTATC | | |
| AJP223 | AAGGAAGATAGAGAGGAAGCGATGTAATGAG TAAAAAAAGACGAAATCGTCATAAA | D39 | *mapZ*$^{+}$ |
| AJP224 | CAACTGGTTTATGAGAAAGTAAGTTCTTTTAG TAGTCCAAGTCATCCGCATGAC | | |
| AJP225 | ATGCGGATGACTTGGACTACTAAAAGAACTTA CTTTCTCATAAACCAGTTGCTG | D39 | *bgaA'* to downstream |
| CS121 | GCTTTCTTGAGGCAATTCACTTGGTGC | | |
| **For construction of IU11912 (Δ*stkP*::$P_c$-*cat*)** | | | |
| TT546 | AGAGAGTCATCCCGAGTTCGAGCAGGTAAA | D39 | 5' fragment with 60 bp of 5' *stkP* |
| TT654 | CATTATCCATTAAAAATCAAACGGATCCTATC GACCAATCTGTTTGACAATCCG | | |
| kanrpsL forward | TAGGATCCGTTTGATTTTTAATGGATAATG | IU11119[e] | $P_c$-*cat* |
| kanrpsL reverse | GGGCCCCTTTCCTTATGCTTTTG | | |
| P1497 | CAAAAGCATAAGGAAAGGGGCCCAATAAGAC TAGAGTCAAGATTTCAATCTACAAACCTA | D39 | 3' fragment with 60 bp of 3' *stkP* |
| P1496 | CAATACCAAGGCGACAGAAGTTCCTGCCCC | | |
| **For construction of IU12192 (Δ*bgaA*::tet- $P_{Zn}$- RBS$^{ftsA}$-*ftsW*$^{+}$)** | | | |
| TT657 | CGCCCCAAGTTCATCACCAATGACATCAAC | IU8122 | Δ*bgaA*::tet- $P_{Zn}$-RBS$^{ftsA}$ |
| YT50 | ATAATTTAATAAGTGCCTCTTACTAATCTTCAT TACATCGCTTCCTCTCTATCTTCCTTG | | |
| YT51 | GGAAGATAGAGAGGAAGCGATGTAATGAAGA TTAGTAAGAGGCACTTATTAAATTATTCC | D39 | *ftsW*$^{+}$ |
| YT52 | CAGCAACTGGTTTATGAGAAAGTAAGTTCTTC TACTTCAACAGAAGGTTCATTGGTTGAT | | |
| YT53 | ATCAACCAATGAACCTTCTGTTGAAGTAGAAG AACTTACTTTCTCATAAACCAGTTGCTG | IU8122 | *bgaA'* to downstream |
| CS121 | GCTTTCTTGAGGCAATTCACTTGGTGC | | |
| **For construction of IU12678 (Δ*bgaA*::tet- $P_{Zn}$- RBS$^{ftsA}$-*cozE*$^{+}$)** | | | |
| P146 | TGGCCATTCATCGCTGGTCGTGCTGAAAT | IU8122 | Δ*bgaA*::tet- $P_{Zn}$-RBS$^{ftsA}$ |
| TT968 | CAAAAAAATAATTTATTTCTACGAAACATTACA TCGCTTCCTCTCTATCTTCCTTGTTAT | | |
| TT969 | AAGGAAGATAGAGAGGAAGCGATGTAATGTT TCGTAGAAATAAATTATTTTTTTGGACCA | D39 | *cozE*$^{+}$ |
| TT970 | CTGGTTTATGAGAAAGTAAGTTCTTTTACTTA GCTAATTCTCTTTCTCGTTCTTTCATTA | | |
| TT971 | AAGAACGAGAAAGAGAATTAGCTAAGTAAAA GAACTTACTTTCTCATAAACCAGTTGCTG | IU8122 | *bgaA'* to downstream |
| CS121 | GCTTTCTTGAGGCAATTCACTTGGTGC | | |
| **For construction of IU12712 and IU12719 (Δ*bgaA*::*kan*-t1t2-$P_{ftsA}$-RBS$^{ftsA}$-*ftsA*)** | | | |

| | | | |
|---|---|---|---|
| P146 | TGGCCATTCATCGCTGGTCGTGCTGAAAT | IU9621 | 5’ *bgaA*’-Kan-T1T2 |
| SC484 | GAGCAAAAAAGAAAGCTCTGTGGTAGAAAC GCAAAAAGGCCATCCGTCAGG | | |
| SC483 | GACGGATGGCCTTTTTGCGTTTCTACCACA GAGCTTTCTTTTTTGCTCTTAGAGAG | D39 | $P_{ftsA}$-*ftsA*$^{+}$ |
| AJP49 | CAACTGGTTTATGAGAAAGTAAGTTCTTTTA TTCGTCAAACATGCTTCCGATC | | |
| AJP50 | CGGAAGCATGTTTGACGAATAAAAGAACTT ACTTTCTCATAAACCAGTTGC | D39 | *bgaA’* to downstream |
| CS121 | GCTTTCTTGAGGCAATTCACTTGGTGC | | |
| **For construction of IU13249 (*murZ*-L-FLAG$^{3}$-$P_{c}$-*erm*)** | | | |
| P1554 | GATTTTGTGGTACGACGGGCATGTATAGCG | D39 | Upstream of *murZ* + *murZ* |
| JQ315 | GCCAGAACCAGCAGCGGAGCCAGCGGAACC ATCCTCAACAAGTCTAATATCCGCTCCTAA | | |
| JQ179 | GGTTCCGCTGGCTCCGCTGCTGGTTCTGGC | IU4970 | L-FLAG$^{3}$-$P_{c}$-*erm* |
| JQ184 | TTATTTCCTCCCGTTAAATAATAGATAACTAT | | |
| JQ316 | ATAGTTATCTATTATTTAACGGGAGGAAATAA ACCGTAGAGGTGTTTATGAATATTTGGA | D39 | Downstream *murZ* |
| P1555 | TGAACCTGAAATCCCCCTGTAACCAGAACT | | |
| **For construction of IU13251 (*murA*-L-FLAG$^{3}$-$P_{c}$-*erm*)** | | | |
| P1558 | TCAGGAGACTACAGGTGGTTCTTCCGATGT | D39 | Upstream of *murA* + *murA* |
| JQ317 | GCCAGAACCAGCAGCGGAGCCAGCGGAACC TTCATCTTCATCATTTGCCTCAATCCGCTG | | |
| JQ179 | GGTTCCGCTGGCTCCGCTGCTGGTTCTGGC | IU4970 | L-FLAG$^{3}$-$P_{c}$-*erm* |
| JQ184 | TTATTTCCTCCCGTTAAATAATAGATAACTAT | | |
| JQ318 | ATAGTTATCTATTATTTAACGGGAGGAAATAA GAAATCAAGCTACGTAGTCAAGCGTTTA | D39 | Downstream *murA* |
| P1559 | CTTAGTACCTGTTCTAGCCCTGCTTAAACT | | |
| **For construction of IU13502, IU13545 (*murZ*-L-FLAG$^{3}$)** | | | |
| P1554 | GATTTTGTGGTACGACGGGCATGTATAGCG | IU13249 | *murZ*-L-FLAG$^{3}$ |
| JQ338 | GGTCCAAATATTCATAAACACCTCTACGGTTT ATTTATCATCATCATCTTTATAATCTTT | | |
| JQ339 | AAAGATTATAAAGATGATGATGATAAATAAAC CGTAGAGGTGTTTATGAATATTTGGACC | D39 | Downstream *murZ* |
| P1555 | TGAACCTGAAATCCCCCTGTAACCAGAACT | | |
| **For construction of IU13536, IU13542 (Δ*murZ*)** | | | |
| P1554 | GATTTTGTGGTACGACGGGCATGTATAGCG | D39 | Upstream of *murZ* + 60 bp 5’ *murZ* |
| JQ344 | TAAATTACGTAATTTTTCGATAATATCAGAACC ACTAATAGTGATTTCACCTTGCAGTGG | | |
| JQ345 | CCACTGCAAGGTGAAATCACTATTAGTGGTT CTGATATTATCGAAAAATTACGTAATTTA | | 60 bp 3’ of *murZ* + downstream *murZ* |
| P1555 | TGAACCTGAAATCCCCCTGTAACCAGAACT | | |
| **For construction of IU13538, IU13546 (Δ*murA*)** | | | |

| | | | |
|---|---|---|---|
| P1558 | TCAGGAGACTACAGGTGGTTCTTCCGATGT | D39 | Upstream of *murA* + 60 bp 5’ *murA* |
| JQ346 | CTGAATCTTAGCACCTAGCTGCGCCAACTTC TCGATCGTCACGCTTCCTACCAGACGATT | | |
| JQ347 | AATCGTCTGGTAGGAAGCGTGACGATCGAGA AGTTGGCGCAGCTAGGTGCTAAGATTCAG | | 60 bp 3’ of *murA* + downstream *murA* |
| P1559 | CTTAGTACCTGTTCTAGCCCTGCTTAAACT | | |
| **For construction of IU13600 (*murZ*(D280Y)-L-FLAG$^3$)** | | | |
| P1554 | GATTTTGTGGTACGACGGGCATGTATAGCG | IU13438 | Upstream of *murZ* + *murZ*(D280Y) |
| JQ315 | GCCAGAACCAGCAGCGGAGCCAGCGGAACC ATCCTCAACAAGTCTAATATCCGCTCCTAA | | |
| JQ179 | GGTTCCGCTGGCTCCGCTGCTGGTTCTGGC | IU13502 | L-FLAG$^3$ + downstream of *murZ* |
| P1555 | TGAACCTGAAATCCCCCTGTAACCAGAACT | | |
| **For construction of IU13604 (Δ*ireB* markerless)** | | | |
| P1711 | GAGTGTCAATGAAGTTCTCAATCTGATTATGG AAACACC | D39 | 5’ upstream of *ireB* + 15 bp of 5’ *ireB* |
| TT1030 | CATTATCCATTAAAAATCAAACGGATCCTAAA AACGTACTGTTTCTTCAGTAAATCCCAT | | |
| TT1031 | AACGTCCAAAAGCATAAGGAAAGGGGCCCTA TCTCAAAGGACAAGGAGTCGATCTATAAC | D39 | 18 bp of 3’ *ireB* and downstream of *ireB* |
| P1712 | CCACTGGACGTTCCAACTCTTCCCCATTTC | | |
| **For construction of IU13680 (Δ*pbp1b*::$P_c$-*aad9*)** | | | |
| P222 | CGTTCGTGTGGCGCTGCTTCAAATTGTT | D39 | Upstream of *pbp1b* and 100 bp of 5’ *pbp1b* |
| P456 | CATTATCCATTAAAAATCAAACGGATCCTATT GAACCTTTCTTGCCAGGTCTAGCTGATT | | |
| KanrpsL forward | TAGGATCCGTTTGATTTTTAATGGATAATG | IU8791 | $P_c$-*aad9* |
| KanrpsL reverse | CAAAAGCATAAGGAAAGGGGCCC | | |
| P225 | CAAAAGCATAAGGAAAGGGGCCCTCTAGCGA TAGCAGTAACTCAAGTACTACACGACCTT | D39 | 60 bp of 3’ *pbp1b* and downstream of *pbp1b* |
| P522 | AACGGCAACCACCAAAGGAGAAACCAAGGA | | |
| **For construction of IU13772 (Δ*bgaA*::*kan*-t1t2-$P_{Zn}$-*murZ*-L-FLAG$^3$)** | | | |
| P146 | TGGCCATTCATCGCTGGTCGTGCTGAAAT | IU11077 | Δ*bgaA*::*kan*-t1t2-$P_{Zn}$-*murZ* |
| JQ315 | GCCAGAACCAGCAGCGGAGCCAGCGGAACC ATCCTCAACAAGTCTAATATCCGCTCCTAA | | |
| JQ179 | GGTTCCGCTGGCTCCGCTGCTGGTTCTGGC | IU4355 | L-FLAG$^3$-*bgaA*’ |
| CS121 | GCTTTCTTGAGGCAATTCACTTGGTGC | | |
| **For construction of IU13794 (Δ*bgaA*::*tet*-$P_{Zn}$-$RBS^{ftsA}$–$divIVA^+$ (R6))** | | | |
| P146 | TGGCCATTCATCGCTGGTCGTGCTGAAAT | IU8122 | Δ*bgaA*::*tet*-t1t2-$P_{Zn}$-$RBS^{ftsA}$ |
| YT72 | TAATGATGTAATTGGCATTCTATTCCTCACTA CATCGCTTCCTCTCTATCTTCCTTGTTA | | |

| | | | |
|---|---|---|---|
| YT73 | TAACAAGGAAGATAGAGAGGAAGCGATGTAGTGAGGAATAGAATGCCAATTACATCATTA | D39 | *divIVA*$^{+}$ (R6 annotation) |
| YT62 | AGCAACTGGTTTATGAGAAAGTAAGTTCTTCTACTTCTGGTTCTTCATACATTGGGCCAA | | |
| YT63 | GGCCCAATGTATGAAGAACCAGAAGTAGAAGAACTTACTTTCTCATAAACCAGTTGCTGC | IU8122 | *bgaA'* to downstream |
| CS121 | GCTTTCTTGAGGCAATTCACTTGGTGC | | |
| **For construction of IU14028, IU14030 (*murA*-L-FLAG$^3$)** | | | |
| P1558 | TCAGGAGACTACAGGTGGTTCTTCCGATGT | IU13251 | Upstream of *murA* + *murA*-L-FLAG$^3$ |
| JQ340 | CTGAATCTTAGCACCTAGCTGCGCCAACTTTTATTTATCATCATCATCTTTATAATCTTT | | |
| JQ341 | AAAGATTATAAAGATGATGATGATAAATAAAAGTTGGCGCAGCTAGGTGCTAAGATTCAG | D39 | Downstream of *murA* |
| P1559 | CTTAGTACCTGTTCTAGCCCTGCTTAAACT | | |
| **For construction of IU14270 (Δ*mraY*<>*aad9*)** | | | |
| TT345 | GTGACCCAGACGCAAATGATTCGTGCCTTT | D39 | upstream of *mraY* |
| TT1076 | TATTCAAATATATCCTCCTCATATTAGTCTCCTAAAGTTAATGTAATTTTTTTAATGTCC | | |
| TT1077 | AAATTACATTAACTTTAGGAGACTAATATGAGGAGGATATATTTGAATACATACGAACAA | IU4888 | *aad9* ORF |
| TT1078 | ATCAGGGTGCCATTCTTATAATTTTTTTAATCTGTTATTTAAATAGTTTATAGTTAAATT | | |
| TT1079 | TATAAACTATTTAAATAACAGATTAAAAAAATTATAAGAATGGCACCCTGATGTTTCAGG | D39 | downstream of *mraY* |
| TT1080 | CTGCTGTCAAGTTTCGACCCAGTTTAGCAAGG | | |
| **For construction of IU14272 (Δ*uppS*<>*aad9*)** | | | |
| TT1070 | GCCATTCTGACGATCATCCGAGACCTTGGT | D39 | upstream of *uppS* |
| TT1071 | GTATGTATTCAAATATATCCTCCTCATGATCTTATTCCTATTCAAAAATCTATCGTTTCA | | |
| TT1072 | CGATAGATTTTTGAATAGGAATAAGATCATGAGGAGGATATATTTGAATACATACGAACA | IU4888 | *aad9* ORF |
| TT1073 | GGGTCATATTTCCTCTTATAATTTTTTTAATCTGTTATTTAAATAGTTTATAGTTAAATT | | |
| TT1074 | ACTATTTAAATAACAGATTAAAAAAATTATAAGAGGAAATATGACCCAGGATTTACAGAA | D39 | downstream of *uppS* |
| TT1075 | GGTTAAAATTCCGAGCATAGCGTTTCCTCCGTC | | |
| **For construction of IU14274 (Δ*murG*<>*aad9*)** | | | |
| TT1064 | CCAACCTCATGCCAACTCATATCGACTACCATG | D39 | upstream of *murG* |
| TT1065 | CGTATGTATTCAAATATATCCTCCTCATATTTTATTCTTTTAACTCCGCTACTGTGTCG | | |
| TT1066 | ACAGTAGCGGAGTTAAAAGAATAAAATATGAGGAGGATATATTTGAATACATACGAACA | IU4888 | *aad9* ORF |
| TT1067 | TTGACATTTACTTTCCTTATAATTTTTTTAATCTGTTATTTAAATAGTTTATAGTTAAAT | | |

| | | | |
|---|---|---|---|
| TT1068 | TTAAATAACAGATTAAAAAAATTATAAGGAAA GTAAATGTCAAAAGATAAGAAAAATGAG | D39 | downstream of *murG* |
| TT1069 | GCCGCCTTGAGTTCTGGGCTAATTTGAGCA | | |
| **For construction of IU14312 (Δ*bgaA*::tet- $P_{Zn}$- RBS$^{ftsA}$-*pbp1a*$^{+}$)** | | | |
| P146 | TGGCCATTCATCGCTGGTCGTGCTGAAAT | IU8122 | Δ*bgaA*::tet- $P_{Zn}$- RBS$^{ftsA}$ |
| BR62 | CGCAGAATCGTTGGTTTGTTCATTACATCGCT TCCTCTCTATCTTCCTTGTTATAATA | | |
| BR61 | ACAAGGAAGATAGAGAGGAAGCGATGTAATG AACAAACCAACGATTCTGCGC | D39 | *pbp1a*$^{+}$ |
| BR64 | CAGCAACTGGTTTATGAGAAAGTAAGTTCTTT TATGGTTGTGCTGGTTGAGGATTCTG | | |
| BR63 | GAATCCTCAACCAGCACAACCATAAAAGAAC TTACTTTCTCATAAACCAGTTGCTGC | D39 | *bgaA'* to downstream |
| CS121 | GCACCAAGTGAATTGCCTCAAGAAAGC | | |
| **For construction of IU14974 (Δ*bgaA*::*kan*-t1t2-$P_{Zn}$-RBS$^{ftsA}$-*stkP*$^{+}$)** | | | |
| P146 | TGGCCATTCATCGCTGGTCGTGCTGAAAT | IU9805 | Δ*bgaA*::*kan*-t1t2-$P_{Zn}$ |
| BR12 | AAATCTTGCCGATTTGGATCATTACATCGCTT CCTCTCTATCTTCCTTGT | | |
| BR13 | GGAAGATAGAGAGGAAGCGATGTAATGATCC AAATCGGCAAGATTTTTG | D39 | *stkP*$^{+}$ |
| BR14 | GCAGCAACTGGTTTATGAGAAAGTAAGTTCTT TTAAGGAGTAGCTGAAGTTGTTTTAGGT | | |
| BR15 | CAATCTACAAACCTAAAACAACTTCAGCTACT CCTTAAAAGAACTTACTTTCTCATAAAC | IU9805 | *bgaA'* to downstream |
| CS121 | GCACCAAGTGAATTGCCTCAAGAAAGC | | |
| **For construction of IU15143 (*murA*(D281Y))** | | | |
| P1558 | TCAGGAGACTACAGGTGGTTCTTCCGATGT | D39 | Upstream *murA* + 5' *murA*(D281Y) |
| TT1145 | ACGAACACGAATTCCTTCGTATTCTTCAATTA CTTCAACA | | |
| TT1146 | TGTTGAAGTAATTGAAGAATACGAAGGAATTC GTGTTCGT | D39 | 3' m*urA*(D281Y) + downstream *murA* |
| P1559 | CTTAGTACCTGTTCTAGCCCTGCTTAAACT | | |
| **For construction of IU15145 (murA(E282Y))** | | | |
| P1558 | TCAGGAGACTACAGGTGGTTCTTCCGATGT | D39 | Upstream *murA* + 5' *murA*(E282Y) |
| TT1147 | GAGAACGAACACGAATTCCGTAGTCTTCTTC AATTACTTCA | | |
| TT1148 | TGAAGTAATTGAAGAAGACTACGGAATTCGT GTTCGTTCTC | D39 | 3' m*urA*(E282Y) + downstream *murA* |
| P1559 | CTTAGTACCTGTTCTAGCCCTGCTTAAACT | | |
| **For construction of IU15939 (*murZ*(C116S))** | | | |
| P1554 | GATTTTGTGGTACGACGGGCATGTATAGCG | D39 | Upstream *murZ* + 5' *murZ*(C116S) |
| TT1203 | CGGACGAGGACCAAGATCAGATCCTCCCGG TAGACCAA | | |

| | | | |
|---|---|---|---|
| TT1204 | TTGGTCTACCGGGAGGATCTGATCTTGGTCC TCGTCCG | D39 | 3’ *murZ*(C116S) + downstream *murZ* |
| P1555 | TGAACCTGAAATCCCCCTGTAACCAGAACT | | |
| **For construction of IU15941 (*murZ*(C116S)-L-FLAG$^3$)** | | | |
| P1554 | GATTTTGTGGTACGACGGGCATGTATAGCG | D39 | 5’ *murZ*(C116S) |
| TT1203 | CGGACGAGGACCAAGATCAGATCCTCCCGG TAGACCAA | | |
| TT1204 | TTGGTCTACCGGGAGGATCTGATCTTGGTCC TCGTCCG | IU13502 | 3’ *murZ*(C116S) -L-FLAG$^3$ |
| P1555 | TGAACCTGAAATCCCCCTGTAACCAGAACT | | |
| **For construction of IU15943 (Δ*bgaA*::*kan*-t1t2-P$_{Zn}$- RBS$^{ftsA}$ -*murZ*(C116S))** | | | |
| P146 | TGGCCATTCATCGCTGGTCGTGCTGAAAT | IU13393 | Δ*bgaA::kan*-t1t2-P$_{Zn}$-*murZ*(C116S) |
| TT1203 | CGGACGAGGACCAAGATCAGATCCTCCCGG TAGACCAA | | |
| TT1204 | TTGGTCTACCGGGAGGATCTGATCTTGGTCC TCGTCCG | IU13393 | *murZ*(C116S) and 3’ bgaA |
| CS121 | GCTTTCTTGAGGCAATTCACTTGGTGC | | |
| **For construction of IU15949 (*murA*(C120S))** | | | |
| P1558 | TCAGGAGACTACAGGTGGTTCTTCCGATGT | D39 | Upstream *murA* + 5’ *murA*(C120S) |
| TT1205 | AGGACGGCTACCAATCGTAGAACCACCTGGC ATGGATA | | |
| TT1206 | TATCCATGCCAGGTGGTTCTACGATTGGTAG CCGTCCT | D39 | 3’ m*urA*(C120S) + downstream *murA* |
| P1559 | CTTAGTACCTGTTCTAGCCCTGCTTAAACT | | |
| **For construction of IU15951 (*murA*(C120S)-L-FLAG$^3$)** | | | |
| P1558 | TCAGGAGACTACAGGTGGTTCTTCCGATGT | D39 | Upstream *murA* + 5’ *murA*(C120S) |
| TT1205 | AGGACGGCTACCAATCGTAGAACCACCTGGC ATGGATA | | |
| TT1206 | TATCCATGCCAGGTGGTTCTACGATTGGTAG CCGTCCT | IU14028 | 3’ *murA*(C120S) -L-FLAG$^3$ + downstream *murA* |
| P1559 | CTTAGTACCTGTTCTAGCCCTGCTTAAACT | | |
| **For construction of IU15954 (Δ*bgaA*::*kan*-t1t2-P$_{Zn}$- RBS$^{ftsA}$ -*murA*(C120S))** | | | |
| P146 | TGGCCATTCATCGCTGGTCGTGCTGAAAT | IU13395 | Δ*bgaA*::*kan*-t1t2-P$_{Zn}$-5’ *murA*(C120S) |
| TT1205 | AGGACGGCTACCAATCGTAGAACCACCTGGC ATGGATA | | |
| TT1206 | TATCCATGCCAGGTGGTTCTACGATTGGTAG CCGTCCT | IU13395 | *3’* *murA*(C120S)-3’ *bgaA’* |
| CS121 | GCTTTCTTGAGGCAATTCACTTGGTGC | | |
| **For construction of IU15983 (Δ*bgaA*::*kan*-t1t2-P$_{Zn}$- RBS$^{ftsA}$ -*murA*-L-FLAG$^3$)** | | | |
| P146 | TGGCCATTCATCGCTGGTCGTGCTGAAAT | IU13395 | Δ*bgaA*::*kan*-t1t2-P$_{Zn}$-*murA* |
| JQ317 | GCCAGAACCAGCAGCGGAGCCAGCGGAACC TTCATCTTCATCATTTGCCTCAATCCGCTG | | |

| | | | |
|---|---|---|---|
| JQ179 | GGTTCCGCTGGCTCCGCTGCTGGTTCTGGC | IU4355 | L-FLAG[3]-*bgaA*’ |
| CS121 | GCTTTCTTGAGGCAATTCACTTGGTGC | | |
| **For construction of IU16334, IU16336 (Δ*bgaA*::*kan*-t1t2-$P_{Zn}$- RBS$^{ftsA}$ -*murZ*(D280Y))** | | | |
| P146 | TGGCCATTCATCGCTGGTCGTGCTGAAAT | IU13393 | Δ*bgaA*::*kan*-t1t2-$P_{Zn}$-5’ *murZ*(D280Y) |
| TT1230 | TTCCTCGACAAAAATGCTGTATTCAGATACAG<br>TCATTCTCA | | |
| TT1231 | TGAGAATGACTGTATCTGAATACAGCATTTTT<br>GTCGAGGAA | IU13393 | *3’ murZ*(D280Y)-*bgaA*’ |
| CS121 | GCTTTCTTGAGGCAATTCACTTGGTGC | | |
| **For construction of IU17134 (Δ*clpE*::$P_c$-[*kan-rpsL*$^+$])** | | | |
| P1730 | ACGAACAATCTCCGAAACATAAGCACCACT | D39 | Upstream of *clpE* + 60 bp of 5’ *clpE* |
| P1727 | CATTATCCATTAAAAATCAAACGGATCCTAA<br>TTGAGATTGGTGTAAAGATGAATTGTTGA | | |
| Kan rpsL forward | TAGGATCCGTTTGATTTTTAATGGATAATG | $P_c$-[*kan-rpsL*$^+$] cassette[d] | $P_c$-[*kan-rpsL*$^+$] |
| Kan rpsL reverse | GGGCCCCTTTCCTTATGCTTTTG | | |
| P1728 | AAACGTCCAAAAGCATAAGGAAAGGGGCCC<br>AACATTCAGATTAAATCTGCCAAAAAAGCT | D39 | 60 bp of 3’ *clpE* + downstream |
| P1729 | TTCTTATGGCATATTCAATAGATTTTCGTA | | |
| **For construction of IU17136 (Δ*clpL*::$P_c$-[*kan-rpsL*$^+$])** | | | |
| P1726 | ATTAGTTTGTTTGCCTATGGAGTTATTGCC | D39 | Upstream of *clpL* + 60 bp of 5’ *clpL* |
| P1723 | CATTATCCATTAAAAATCAAACGGATCCTAA<br>CCCATCAATTGGTTAAATAAATCATCCAT | | |
| Kan rpsL forward | TAGGATCCGTTTGATTTTTAATGGATAATG | $P_c$-[*kan-rpsL*$^+$] cassette[d] | $P_c$-[*kan-rpsL*$^+$] |
| Kan rpsL reverse | GGGCCCCTTTCCTTATGCTTTTG | | |
| P1724 | CAAAAGCATAAGGAAAGGGGCCCGCTAAAC<br>ATCTGGAAGCAGATATGGAAGAT | D39 | 60 bp of 3’ *clpL* + downstream |
| P1725 | TTCGTAAACTGGGTATCAACGTAACCTTTG | | |
| **For construction of IU17138 (Δ*clpP*::$P_c$-[*kan-rpsL*$^+$])** | | | |
| P1722 | CGAATGGACGACTACGCCCAATACCTTTAT | D39 | Upstream of *clpP* + 90 bp of 5’ *clpP* |
| P1719 | CATTATCCATTAAAAATCAAACGGATCCTAC<br>AGCATAATGATGCGGTCTTTGAGAAGACG | | |
| Kan rpsL forward | TAGGATCCGTTTGATTTTTAATGGATAATG | $P_c$-[*kan-rpsL*$^+$] cassette[d] | $P_c$-[*kan-rpsL*$^+$] |
| Kan rpsL reverse | GGGCCCCTTTCCTTATGCTTTTG | | |
| P1720 | AAACGTCCAAAAGCATAAGGAAAGGGGCCC<br>CAGGAAACACTTGAATATGGCTTTATTGAT | D39 | 60 bp of 3’ *clpP* + downstream |
| P1721 | GTGTAAAGAACAACTTTCTTAGCATTTAAT | | |
| **For construction of IU17150, IU17158 (Δ*clpE*::$P_c$-*erm*)** | | | |
| P1730 | ACGAACAATCTCCGAAACATAAGCACCACT | D39 | Upstream of *clpE* + 60 bp of 5’ *clpE* |
| P1727 | CATTATCCATTAAAAATCAAACGGATCCTAAT<br>TGAGATTGGTGTAAAGATGAATTGTTGA | | |
| Kan rpsL forward | TAGGATCCGTTTGATTTTTAATGGATAATG | $P_c$-*erm* cassette[d] | $P_c$-*erm* |

| | | | |
|---|---|---|---|
| Kan rpsL reverse | GGGCCCCTTTCCTTATGCTTTTG | | |
| P1728 | AAACGTCCAAAAGCATAAGGAAAGGGGCCC AACATTCAGATTAAATCTGCCAAAAAAGCT | D39 | 60 bp of 3' *clpE* + downstream |
| P1729 | TTCTTATGGCATATTCAATAGATTTTCGTA | | |
| **For construction of IU17152, IU17160 (Δ*clpL*::P$_c$-*erm*)** | | | |
| P1726 | ATTAGTTTGTTTGCCTATGGAGTTATTGCC | D39 | Upstream of *clpL* + 60 bp of 5' *clpL* |
| P1723 | CATTATCCATTAAAAATCAAACGGATCCTAA CCCATCAATTGGTTAAATAAATCATCCAT | | |
| Kan rpsL forward | TAGGATCCGTTTGATTTTTAATGGATAATG | P$_c$-*erm* cassette[d] | P$_c$-*erm* |
| Kan rpsL reverse | GGGCCCCTTTCCTTATGCTTTTG | | |
| P1724 | CAAAAGCATAAGGAAAGGGGCCCGCTAAAC ATCTGGAAGCAGATATGGAAGAT | D39 | 60 bp of 3' *clpL* + downstream |
| P1725 | TTCGTAAACTGGGTATCAACGTAACCTTTG | | |
| **For construction of IU17154, IU17162 (Δ*clpP*::P$_c$-*erm*)** | | | |
| P1722 | CGAATGGACGACTACGCCCAATACCTTTAT | D39 | Upstream of *clpP* + 90 bp of 5' *clpP* |
| P1719 | CATTATCCATTAAAAATCAAACGGATCCTAC AGCATAATGATGCGGTCTTTGAGAAGACG | | |
| Kan rpsL forward | TAGGATCCGTTTGATTTTTAATGGATAATG | P$_c$-*erm* cassette[d] | P$_c$-*erm* |
| Kan rpsL reverse | GGGCCCCTTTCCTTATGCTTTTG | | |
| P1720 | AAACGTCCAAAAGCATAAGGAAAGGGGCCC CAGGAAACACTTGAATATGGCTTTATTGAT | D39 | 60 bp of 3' *clpP* + downstream |
| P1721 | GTGTAAAGAACAACTTTCTTAGCATTTAAT | | |
| **For construction of IU17170 (*murZ*-HA)** | | | |
| P1554 | GATTTTGTGGTACGACGGGCATGTATAGCG | D39 | Upstream of *murZ* + *murZ*-HA |
| MJ062 | GCATAATCTGGAACATCATATGGATAATCCT CAACAAGTCTAATATCCGCTCCTAA | | |
| MJ063 | GGATTATCCATATGATGTTCCAGATTATGCT TAAACCGTAGAGGTGTTTATGAATATTTG | D39 | HA + downstream of *murZ* |
| P1555 | TGAACCTGAAATCCCCCTGTAACCAGAACT | | |
| **For construction of IU17619 (*murZ*(E190A E192A))** | | | |
| P1554 | GATTTTGTGGTACGACGGGCATGTATAGCG | D39 | Upstream of *murA* + *murA*(E190A E192A) |
| TT1360 | AGCTACATCAATAATCGCAGGTGCACGGGCTG CATTTTCA | | |
| TT1361 | TGAAAATGCAGCCCGTGCACCTGCGATTATTG ATGTAGCT | D39 | *murA*(E190A E192A) + downstream of *murA* |
| P1555 | TGAACCTGAAATCCCCCTGTAACCAGAACT | | |
| **For construction of IU17622 (*murZ*(E192A))** | | | |
| P1554 | GATTTTGTGGTACGACGGGCATGTATAGCG | D39 | Upstream of *murA* + *murA*(E192A) |
| TT1358 | GTAGCTACATCAATAATCGCAGGTTCACGGGC TGCATTT | | |
| TT1359 | AAATGCAGCCCGTGAACCTGCGATTATTGATG TAGCTAC | D39 | |

| | | | |
|---|---|---|---|
| P1555 | TGAACCTGAAATCCCCCTGTAACCAGAACT | | *murA*(E192A) + downstream of *murA* |
| **For construction of IU17623 (*murZ*(D195A))** | | | |
| P1554 | GATTTTGTGGTACGACGGGCATGTATAGCG | D39 | Upstream of *murA* + *murA*(D195A) |
| TT1356 | TATTCAAGAGAGTAGCTACAGCAATAATCTCAG GTTCACGGG | | |
| TT1357 | CCCGTGAACCTGAGATTATTGCTGTAGCTACTC TCTTGAATA | D39 | *murA*(E195A) + downstream of *murA* |
| P1555 | TGAACCTGAAATCCCCCTGTAACCAGAACT | | |
| **For construction of IU17627 (*murZ*(E259A))** | | | |
| P1554 | GATTTTGTGGTACGACGGGCATGTATAGCG | D39 | Upstream of *murA* + *murA*(E259A) |
| TT1362 | GCAATAAACCCTTCCAGGTGTGCGTAAAGAAC ATTATTTA | | |
| TT1363 | TAAATAATGTTCTTTACGCACACCTGGAAGGGT TTATTGC | D39 | *murA*(E259A) + downstream of *murA* |
| P1555 | TGAACCTGAAATCCCCCTGTAACCAGAACT | | |
| **For construction of IU17764 (F-*murZ*)** | | | |
| P1554 | GATTTTGTGGTACGACGGGCATGTATAGCG | D39 | Upstream of *murZ* + F |
| TT1366 | CTTTTATCATCATCATCTTTATAATCCATTCT AAGTTTTCAATACTCTTTCAAGATTTCT | | |
| TT1367 | TAGAATGGATTATAAAGATGATGATGATAAA AGAAAAATTGTTATCAATGGTGGATTACC | D39 | F-*murZ* + downstream of *murZ* |
| P1555 | TGAACCTGAAATCCCCCTGTAACCAGAACT | | |
| **For construction of IU17766 (HA-*murZ*)** | | | |
| P1554 | GATTTTGTGGTACGACGGGCATGTATAGCG | D39 | Upstream of *murZ* + HA |
| TT1368 | AGCATAATCTGGAACATCATATGGATACATT CTAAGTTTTCAATACTCTTTCAAGATTTC | | |
| TT1369 | AATGTATCCATATGATGTTCCAGATTATGCT AGAAAAATTGTTATCAATGGTGGATTACC | D39 | HA-*murZ* + downstream of *murZ* |
| P1555 | TGAACCTGAAATCCCCCTGTAACCAGAACT | | |
| **For construction of IU17768 (F-*murA*)** | | | |
| P1558 | TCAGGAGACTACAGGTGGTTCTTCCGATGT | D39 | Upstream of *murA* + F |
| TT1370 | ATCTTTATCATCATCATCTTTATAATCCATAC TCGTTTCCTTTACTCTTGATTTCATAAT | | |
| TT1371 | AAACGAGTATGGATTATAAAGATGATGATGA TAAAGATAAAATTGTGGTTCAAGGTGGCG | D39 | F-*murA* + downstream of *murA* |
| P1559 | CTTAGTACCTGTTCTAGCCCTGCTTAAACT | | |
| **For construction of IU17770 (HA-*murA*)** | | | |
| P1558 | TCAGGAGACTACAGGTGGTTCTTCCGATGT | D39 | Upstream of *murA* + HA |
| TT1372 | AGCATAATCTGGAACATCATATGGATACATACTC GTTTCCTTTACTCTTGATTTCATAAT | | |
| TT1373 | CGAGTATGTATCCATATGATGTTCCAGATTATGC TGATAAAATTGTGGTTCAAGGTGGCG | D39 | HA-*murA* + downstream of *murA* |
| P1559 | CTTAGTACCTGTTCTAGCCCTGCTTAAACT | | |
| **For construction of IU17838, IU17840 (*iht*-$L_6$-*murZ*)** | | | |

| | | | |
|---|---|---|---|
| P1554 | GATTTTGTGGTACGACGGGCATGTATAGCG | D39 | Upstream of *murZ* |
| TT1392 | AAAAATTTCCAAACCTTTTTATCCATTCTAA GTTTTCAATACTCTTTCAAGATTTCTAA | | |
| TT1393 | AATCTTGAAAGAGTATTGAAAACTTAGAATG GATAAAAAAGGTTTGGAAATTTTTTTGGC | IU14738 | *iht*-$L_6$ |
| TT1394 | TTGCAGTGGTAATCCACCATTGATAACAATT TTTCTACCAGAACCTTGACCAGATCCTGG | | |
| TT1395 | CAAGGACCAGGATCTGGTCAAGGTTCTGGT AGAAAAATTGTTATCAATGGTGGATTACCA | D39 | *murZ* + downstream |
| P1555 | TGAACCTGAAATCCCCCTGTAACCAGAACT | | |
| **For construction of IU17841 (*iht*-$L_6$-*murA*)** | | | |
| P1558 | TCAGGAGACTACAGGTGGTTCTTCCGATGT | D39 | Upstream of *murA* |
| TT1396 | CAAAAAAATTTCCAAACCTTTTTTATCCATAC TCGTTTCCTTTACTCTTGATTTCATAAT | | |
| TT1397 | TATGAAATCAAGAGTAAAGGAAACGAGTATG GATAAAAAAGGTTTGGAAATTTTTTTGGC | IU14738 | *iht*-$L_6$ |
| TT1398 | CAGACGATTATCGCCACCTTGAACCACAATT TTATCACCAGAACCTTGACCAGATCCTGG | | |
| TT1399 | CAGGACAAGGACCAGGATCTGGTCAAGGTT CTGGTGATAAAATTGTGGTTCAAGGTGGCG | D39 | *murA* + downstream |
| P1559 | CTTAGTACCTGTTCTAGCCCTGCTTAAACT | | |
| **For construction of IU18555 (Δ*bgaA*::*tet*-$P_{Zn}$-$RBS^{ftsA}$-*stkP*$^+$)** | | | |
| TT657 | CGCCCCAAGTTCATCACCAATGACATCAAC | IU9990 | Δ*bgaA*::*tet*-$P_{Zn}$-$RBS^{ftsA}$ |
| TT1435 | AAATCTTGCCGATTTGGATCATTACATCGCT TCCTCTCTATCTTCCTTGTTATAATAGAT | | |
| TT1436 | ATAACAAGGAAGATAGAGAGGAAGCGATGT AATGATCCAAATCGGCAAGATTTTTGCCGG | IU14974 | *stkP*$^+$- *bgaA'* to downstream |
| CS121 | GCTTTCTTGAGGCAATTCACTTGGTGC | | |
| **For construction of IU18665 (Δ*stkP* markerless)** | | | |
| TT546 | AGAGAGTCATCCCGAGTTCGAGCAGGTAAA | D39 | Upstream *stkP* + 60 bp 5' *stkP* |
| TT1315 | TGTAGATTGAAATCTTGACTCTAGTCTTATTT CGACCAATCTGTTTGACAATCCG | | |
| TT1316 | GGATTGTCAAACAGATTGGTCGAAATAAGAC TAGAGTCAAGATTTCAATCTACAAACC | D39 | 60 bp of 3' *stkP* + downstream |
| P1496 | CAATACCAAGGCGACAGAAGTTCCTGCCCC | | |
| **For construction of IU19079 (*murZ*(D280Y)-L-FLAG$^3$-$P_c$-*erm*)** | | | |
| P1554 | GATTTTGTGGTACGACGGGCATGTATAGCG | IU13438 | *murZ*(D280Y) |
| JQ315 | GCCAGAACCAGCAGCGGAGCCAGCGGAAC CATCCTCAACAAGTCTAATATCCGCTCCTAA | | |
| JQ179 | GGTTCCGCTGGCTCCGCTGCTGGTTCTGGC | IU13249 | L-FLAG$^3$-$P_c$-*erm* + downstream |
| P1555 | TGAACCTGAAATCCCCCTGTAACCAGAACT | | |
| **For construction of IU19821 (Δ*spd-0567*::$P_c$-[*sacB-kan-rpsL*$^+$])** | | | |
| TT1522 | GTCCCTATTGATGCGGAATTTGACTGTCCC | D39 | 5' fragment with 90 bp of 5' *spd_0567* |
| TT1527 | CATTATCCATTAAAAATCAAACGGATCCTATC CCATAGTCGCATCCACTACGACATCCTC | | |
| Kan rpsL forward | TAGGATCCGTTTGATTTTTAATGGATAATG | | $P_c$-[*sacB-kan-rpsL*$^+$]$^f$ |

| | | | |
|---|---|---|---|
| Kan rpsL reverse | GGGCCCCTTTCCTTATGCTTTTG | $P_c$-[*kan-rpsL*$^+$] cassette[d] | |
| TT1528 | CAAAAGCATAAGGAAAGGGGCCCGTCAACAACCCGCCGTTTTAGTGATGATT | D39 | 3’ fragment with 60 bp of 3’ *spd_0567* |
| TT1520 | CCAGAAGCATCATTCAAGAGTCCTTCGCCC | | |

| **Templates and primers used to generate amplicons for transformation assays** | | | |
|---|---|---|---|
| TT196 | GCCAAGCCCTGAGACAAATAGTAGTCGTTGGT | IU4888 | Δ*gpsB*<>*aad9* |
| TT197 | TTTGATACGATCTGCTGCCCGAAGCCAAAGGT | | |
| P1554 | GATTTTGTGGTACGACGGGCATGTATAGCG | E767 | Δ*murZ*::$P_c$-*erm* |
| P1555 | TGAACCTGAAATCCCCCTGTAACCAGAACT | | |
| P1558 | TCAGGAGACTACAGGTGGTTCTTCCGATGT | E765 | Δ*murA*::$P_c$-*erm* |
| P1559 | CTTAGTACCTGTTCTAGCCCTGCTTAAACT | | |
| TT571 | GAGCGAGTGCTTGATGCCTGTGCGGCTCCA | IU7923 | Δ*stkP*::$P_c$-*erm* |
| P1496 | CAATACCAAGGCGACAGAAGTTCCTGCCCC | | |
| TT329 | CAACTGATATAGTTGGAAGTGAGGAGTCCATTTCCC | IU9931 | Δ*rodZ* <>*aad9* |
| P1385 | ACAACACCTGCAATGGCCACACGTTGCTTT | | |
| TT329 | CAACTGATATAGTTGGAAGTGAGGAGTCCATTTCCC | IU6987 | Δ*rodZ* ::$P_c$-*aad9* |
| P1385 | ACAACACCTGCAATGGCCACACGTTGCTTT | | |
| TT329 | CAACTGATATAGTTGGAAGTGAGGAGTCCATTTCCC | E655 | Δ*rodZ* ::$P_c$-*erm* |
| P1385 | ACAACACCTGCAATGGCCACACGTTGCTTT | | |
| TT452 | GGAGGGTTGGCTGTGGGTGGCTACAAGAAC | IU7397 | Δ*pbp2b* <>*aad9* |
| TT352 | TGAAGGACTGGAAAGACCACTGCACCTTCT | | |
| P104 | AATGAGACGTGTTGCCATTGCAGG | IU1751 | Δ*mreCD* <>*aad9* |
| P107 | TGTCGCTTTCTCAGCAGCAAGACT | | |
| P222 | CGTTCGTGTGGCGCTGCTTCAAATTGTT | E193 | Δ*pbp1b*::$P_c$*erm* |
| P522 | AACGGCAACCACCAAAGGAGAAACCAAGGA | | |
| P222 | CGTTCGTGTGGCGCTGCTTCAAATTGTT | IU13680 | Δ*pbp1b* ::$P_c$-*aad9* |
| P522 | AACGGCAACCACCAAAGGAGAAACCAAGGA | | |
| P222 | CGTTCGTGTGGCGCTGCTTCAAATTGTT | K180 | Δ*pbp1b*::$P_c$-[*kan-rpsL*$^+$] |
| P522 | AACGGCAACCACCAAAGGAGAAACCAAGGA | | |

| **Primers used to confirm deletion junction in Δ*gpsB sup3*** | |
|---|---|
| Primer name | Sequence (5’ to 3’) |
| P1, P1510 | ACCATTGCCACTGCGAACATGGTCTACAGC |
| P2, TT1345 | GCACCAAGGTTCCCAGCATCAAGGTCAGC |

| | | |
|---|---|---|
| P3, TT1346 | TGGCAAACGTGACTCAGTCAATGTCGCTGC | |
| P4, TT1347 | CTAGTCTTTACAAGTATCTAACCGAGGAGGTTGAAA ACGATCAG | |
| **Primers used for detection of *spd_1033 to spd_1035* in Δ*gpsB* suppressor strains** | | |
| Primer name | Sequence (5' to 3') | Product |
| P1481 | TTATGTAGGAGGAACCGAGGGCGGAGGAAT | *3' spd_1036 to 5' spd_1032* |
| P1482 | AGACGAGTGTTCCATAGCCGACTCCTTCATTT | |
| **Primers used for qRT-PCR** | | |
| Primer name | Sequence (5' to 3') | Gene name |
| JQ342 | GGAGCTACTGTTAAGCGTTATG | *murZ* |
| JQ343 | CGCCTTAAGGTGTAAGTCAATC | |
| KK489 | AAAGGTCGTGGTGGTAAGGGAATG | *gyrA* |
| KK490 | GCATCTTGATCCAGGCGCATTACT | |

[a]FLAG-tag fusions ((C)-L-FLAG$^3$) were made to the carboxyl-ends (C) of reading frames. The amino acid sequence of the FLAG epitope is DYKDDDDK (Ramos-Montanez *et al.*, 2008, Wayne *et al.*, 2010). The FLAG-tag used in this study contained a linker sequence (L; GSAGSAAGSG) followed by three tandem copies of the FLAG epitope (FLAG$^3$).

[b]Antibiotic resistance markers: Erm$^R$, erythromycin; Kan$^R$, kanamycin; Spc$^R$, spectinomycin; Str$^R$, streptomycin; Cm$^R$, chloramphenicol; Tet$^R$, tetracycline.

[c]Genomic DNA of indicated *S. pneumoniae* strains was used as templates for PCR reactions, except for P$_c$-[*kan-rpsL*$^+$] and P$_c$-*erm* cassettes.

[d]P$_c$-*erm* and P$_c$-[*kan-rpsL*$^+$] cassettes are described in (Tsui *et al.*, 2011).

[e]Genotype of IU11119 is *ezrA*-L$_0$-*sfgfp*-P$_c$-*cat*, as described in (Perez *et al.*, 2019).

[f]P$_c$-[*sacB-kan-rpsL*$^+$] is described in (Li *et al.*, 2014)

**Table S2**. Blastn results using *phtD* as query sequence against *S. pneumoniae* D39 database.

| *spd #* | % identity | alignment length | Mis matches | gap opens | q. start | q. end | s. start | s. end | evalue | bit score |
|---|---|---|---|---|---|---|---|---|---|---|
| ***spd_0889, phtD*** | 100 | 2562 | 0 | 0 | 899901 | 902462 | 899901 | 902462 | 0 | 4732 |
| ***spd_1037, phtB*** | 100 | 1324 | 0 | 0 | 900760 | 902083 | 1063403 | 1062080 | 0 | 2446 |
| ***spd_1037, phtB*** | 78.7 | 700 | 125 | 14 | 899901 | 900585 | 1064304 | 1063614 | 2.6E-125 | 446 |
| *spd_1038, phtA* | 77.3 | 699 | 137 | 12 | 899901 | 900585 | 1066912 | 1066222 | 1.2E-108 | 390 |
| *spd_1038, phtA* | 89.1 | 258 | 28 | 0 | 900760 | 901017 | 1066011 | 1065754 | 4.6E-88 | 322 |

**Table S3**. Blastn results using *spd_0966* as query sequence against *S. pneumoniae* D39 database[a]

| *spd #* | % identity | alignment length | Mis matches | gap opens | q. start | q. end | s. start | s. end | evalue | bit score |
|---|---|---|---|---|---|---|---|---|---|---|
| ***spd_0966*** | 100 | 1492 | 0 | 0 | 978724 | 980215 | 978724 | 980215 | 0 | 2691 |
| *spd_0758* | 93 | 1518 | 84 | 4 | 978724 | 980215 | 768573 | 770089 | 0 | 2244 |
| *spd_1641* | 91 | 1479 | 109 | 3 | 978724 | 980185 | 1656439 | 1654961 | 0 | 2102 |
| ***spd_0986*** | 91 | 1477 | 119 | 3 | 978724 | 980183 | 998197 | 999673 | 0 | 2053 |
| *spd_1666* | 88 | 1489 | 127 | 5 | 978724 | 980187 | 1679982 | 1678514 | 0 | 1929 |
| *spd_0048* | 86 | 1486 | 132 | 4 | 978724 | 980183 | 40190 | 41625 | 0 | 1793 |
| *spd_1708* | 86 | 831 | 98 | 6 | 979359 | 980188 | 1709287 | 1708478 | 0 | 951 |
| *spd_1708* | 89 | 595 | 42 | 1 | 978724 | 979294 | 1709869 | 1709275 | 0 | 793 |
| *spd_0034* | 87 | 404 | 51 | 2 | 979781 | 980183 | 29738 | 30140 | 1.43E-136 | 483 |
| *spd_1681* | 98 | 284 | 5 | 1 | 979905 | 980187 | 1697825 | 1697542 | 1.43E-136 | 482 |
| *spd_1681* | 92 | 319 | 24 | 1 | 978724 | 979041 | 1698142 | 1697824 | 1.63E-129 | 460 |
| *spd_0022* | 84 | 386 | 61 | 1 | 979463 | 979848 | 21517 | 21901 | 6.08E-116 | 414 |

[a]The reading frame of *spd_0966* was assigned to be from 978757 to 980059 on the complementary strand of D39 genome. The blastn analysis was performed with sequence 978724 to 980215, corresponding to 156 bp upstream to 33 bp downstream of *spd_0966*.

**Table S4**. Blastn results using *spd_1690* to *spd_1703* (*rRNA*) as query sequence against *S. pneumoniae* D39 database[a]

| *spd* # | % identity | alignment length | Mis matches | gap opens | q. start | q. end | s. start | s. end | evalue | bit score |
|---|---|---|---|---|---|---|---|---|---|---|
| **(*rRNA-2*) *spd_1690* to *spd_1703*** | 100 | 5998 | 0 | 0 | 1699037 | 1705034 | 1699037 | 1705034 | 0 | 10817 |
| **(*rRNA-3*) *spd_1804* to *spd_1817*** | 99.9 | 5998 | 5 | 0 | 1699037 | 1705034 | 1796588 | 1802585 | 0 | 10795 |
| **(*rRNA-4*) *spd_1889* to *spd_1894*** | 100.0 | 5289 | 1 | 0 | 1699746 | 1705034 | 1859222 | 1864510 | 0 | 9534 |
| (*rRNA-1*) *spd_0015* to *spd_0019* | 99.9 | 5216 | 6 | 0 | 1699819 | 1705034 | 20042 | 14827 | 0 | 9380 |

[a]The blastn analysis was performed with sequence 1699037 to 1705034, corresponding to *spd_1690* to *spd_1703*. The genes from *spd_1690* to *spd_1703* are tRNA-pro (*spd_1690*); tRNA-arg (*spd_1691*); tRNA-leu (*spd_1692*); tRNA-gly (*spd_1693*); tRNA-thr (*spd_1694*); tRNA-leu (*spd_1695*); tRNA-lys (*spd_1696*); tRNA-asp (*spd_1697*); tRNA-val (*spd_1698*); *rrfB* (5S rRNA, *spd_1699*); *rrlB* (23S rRNA, *spd_1700*), tRNA-ala (*spd_1701*); *rrsB* (16S rRNA, *spd_1702*) and tRNA-glu (*spd_1703*). Alignment of *spd_1804* to *spd_1817* covers all sequence from *spd_1690* to *spd_1703.* Alignment of *spd_1889* to *spd_1894* covers tRNA-val to tRNA-glu, and alignment of *spd_0015* to *spd_0019* covers from 5S rRNA to tRNA-glu.

**Table S5.** Suppression of Δ*gpsB* lethality in *S. pneumoniae* D39 or R6 strains[a].

A. Transformation with Δ*gpsB*<>*aad9* amplicon

| Genetic background | Recipient strains | Number of colonies 22 h after transformation |
|---|---|---|
| D39 Δ*cps* *rpsL1* | 1. WT (IU1824) | 0 |
| | 2. WT + Zn[b,c] | 0 |
| | 3. *gpsB*$^+$//$P_{Zn}$-*gpsB*$^+$(IU15877) | 0 |
| | 4. *gpsB*$^+$//$P_{Zn}$-*gpsB*$^+$ + Zn[c] | >500 |
| | 5. *murZ*(D280Y) (IU13438) | >500 small |
| | 6. *murZ*(I265V, R6 allele) (IU14210) | >500 small |
| | 7. *murZ*(E259A) (IU17627) | >500 small |
| | 8. *murZ*(E190A E192A) (IU17619) | 0 |
| | 9. *murZ*(E192A) (IU17622) | 0 |
| | 10. *murZ*(E195A) (IU17623) | 0 |
| | 11. *murA*(D281Y) (IU15143) | 0 |
| | 12. *murA*(E282Y) (IU15145) | 0 |
| | 13. *murZ*$^+$//$P_{Zn}$-*murZ*$^+$(IU13393) | 0 |
| | 14. *murZ*$^+$//$P_{Zn}$-*murZ*$^+$ + Zn[b] | >500 small |
| | 15. *murZ*$^+$//$P_{Zn}$-*murZ*(C116S)(IU15943) | 0 |
| | 16. *murZ*$^+$//$P_{Zn}$-*murZ*(C116S) + Zn[b] | 0 |
| | 17. *murA*$^+$//$P_{Zn}$-*murA*$^+$(IU13395) | 0 |
| | 18. *murA*$^+$//$P_{Zn}$-*murA*$^+$ + Zn[c] | >500 |
| | 19. *murA*$^+$//$P_{Zn}$-*murA*(C120S)(IU15954) | 0 |
| | 20. *murA*$^+$//$P_{Zn}$-*murA*(C120S) + Zn[c] | 0 |
| | 21. Δ*murZ* (IU13536) | 0 |
| | 22. Δ*murA* (IU13538) | 0 |
| | 23. Δ*clpC*::$P_c$-[*kan-rpsL*$^+$] (IU12462) | 0 |
| | 24. Δ*clpP*::$P_c$-[*kan-rpsL*$^+$] (IU17138) | 0 |
| | 25. Δ*clpE*::$P_c$-[*kan-rpsL*$^+$] (IU17134) | 0 |
| | 26. Δ*clpL*::$P_c$-[*kan-rpsL*$^+$] (IU17136) | 0 |
| | 27. *murZ*-L-FLAG$^3$ (IU13502) | 0 |
| | 28. *murA*-L-FLAG$^3$ (IU14028) | 0 |
| | 29. *murZ*(D280Y)-L-FLAG$^3$ (IU13600) | >500 small |
| | 30. Δ*khpA* (IU9036) | >500, small |
| | 31. Δ*khpB* (IU10592) | >500, small |
| | 32. *khpB*(T89A)(IU12744) | 0 |
| | 33. Δ*khpA* Δ*murZ* (IU13542) | 0 |
| | 34. Δ*khpA* Δ*murA* (IU13546) | >500, very small |
| D39 Δ*cps* | 35. WT (IU1945) | 0 |
| | 36. *murZ*$^+$//$P_{Zn}$-*murZ*$^+$(IU11077) | 0 |
| | 37. *murZ*$^+$//$P_{Zn}$-*murZ*$^+$ + Zn[b] | >500 small |
| | 38. *murA*$^+$//$P_{Zn}$-*murA*$^+$(IU11079) | 0 |
| | 39. *murA*$^+$//$P_{Zn}$-*murA*$^+$ + Zn[c] | >500 |
| R6[d] | 40. WT, EL59 (*murZ*(I265V)) | >500 small |
| | 41. Δ*murZ*::$P_c$-*erm* (IU16265) | 0 |
| | 42. Δ*murA*::$P_c$-*erm* (IU16267) | >500 small |

B. Transformation with Δ*murZ*::$P_c$-*erm* amplicon

| Genetic background | Recipient strains | Number of colonies 22 h after transformation |
|---|---|---|
| D39 Δ*cps rpsL1* | 1. WT (IU1824) | >500 |
| | 2. Δ*murA* (IU13538) | 0 |
| | 3. *murA*(C120S)(IU15949) | 0 |
| | 4. *murA*-L-FLAG$^3$ (IU14028) | >500 |
| | 5. Δ*khpA* Δ*gpsB*<>*aad9* (IU12883, IU16196) | 0 |
| R6[d] | 6. WT, EL59 (*murZ*(I265V)) | >500 |
| | 7. Δ*gpsB*<>*aad9* (IU8224) | 0 |

C. Transformation with Δ*murA*::$P_c$-*erm* amplicon

| Genetic background | Recipient strains | Number of colonies 22 h after transformation |
|---|---|---|
| D39 Δ*cps rpsL1* | 1. WT (IU1824) | >500 |
| | 2. Δ*murZ* (IU13536) | 0 |
| | 3. *murZ*(C116S)(IU15939) | 0 |
| | 4. *murZ*(D280Y)(IU13438) | >500 |
| | 5. *murZ*-L-FLAG$^3$ (IU13502) | >500 |
| | 6. Δ*khpA* Δ*gpsB*<>*aad9* (IU12883, IU16196) | ~ 25 to 50, small[e] |
| R6[d] | 7. WT, EL59, (*murZ*(I265V)) | >500 |
| | 8. Δ*gpsB*<>*aad9* (IU8224) | >500 small |

[a]Transformations and visualization of colonies were performed as described in Experimental procedures and footnote to Table 3. Colony sizes are relative to colonies transformed with positive control Δ*pbp1b* amplicons containing the same antibiotic selection marker.

[b]0.2 mM $ZnCl_2$ + 0.02 mM $MnSO_4$ were added to transformation mixes and in subsequent steps to induce expression of *murZ* under control of the $P_{Zn}$ zinc-inducible promoter in the ectopic *bgaA* site.

[c]0.4 mM $ZnCl_2$ + 0.04 mM $MnSO_4$ were added to transformation mixes and in subsequent steps to induce ectopic expression of *gpsB* or *murA.*

[d]R6 strain contains a spontaneous *murZ*(I265V) mutation compared to D39 strain (Lanie *et al.*, 2007).

[e]Both IU12883 and IU16196, two independent Δ*khpA* Δ*gpsB*<>*aad9* isolates obtained from independent transformations, have very low transformation efficiency. Transformation of these

strains with a positive control Δ*pbp1b* amplicon also yielded the same low number (25 to 50) of transformants.

**Table S6**. Overexpression strains in D39 Δ*cps* backgrounds that did not suppress Δ*gpsB* essentiality[a]

In IU1824 (D39 Δ*cps rpsL1*) background

| Recipient strain genotype[a] | Strain[b] |
|---|---|
| $P_{Zn}$-*stkP*$^+$ | IU14974 |
| $P_{Zn}$-*pbp1a*$^+$ | IU14312 |
| $P_{Zn}$-*pbp2a*$^+$ | IU14318 |
| $P_{Zn}$-*mreC*$^+$ | IU10220 |
| $P_{Zn}$-*rodZ*$^+$ | IU9613 |
| CEP- $P_{Zn}$-*ezrA*$^+$ Δ*bgaA*:: $P_{Zn}$-*ezrA*$^+$ | IU13327 |
| $P_{Zn}$-*divIVA*$^+$ (R6 annotation) | IU13794 |
| $P_{Zn}$-*ftsA*$^+$ | IU12310 |
| $P_{Zn}$-*ftsZ*$^+$ | IU12286 |
| $P_{Zn}$-*ftsW*$^+$ | IU12192 |

In IU1945 (D39 Δ*cps*) background

| Recipient strain genotype[a] | Strain |
|---|---|
| $P_{Zn}$-*pbp2x*$^+$ | IU10063 |
| $P_{Zn}$-*pbp2b*$^+$ | IU9990 |
| $P_{Zn}$-*pbp1b*$^+$ | IU9992 |
| $P_{Zn}$-*mltG*$^+$ | IU8872 |
| $P_{Zn}$-*rodA*$^+$ | IU10922 |
| $P_{Zn}$-*mraY*$^+$ | IU11083 |
| $P_{Zn}$-*uppS*$^+$ | IU10094 |
| $P_{Zn}$-*murG*$^+$ | IU11049 |
| $P_{Zn}$-*cozE*$^+$ | IU12678 |
| $P_{Zn}$-*mapZ*$^+$ | IU11628 |
| $P_{Zn}$-*sepF*$^+$ | IU9805 |

[a]Recipient strains and Δ*gpsB*<>*aad9* were obtained as described in Table S1. Transformations with 1 mL of transformation mixture were performed as described in *Experimental procedures*. Final concentrations of 0.4 mM $ZnCl_2$ + 0.04 mM $MnSO_4$ were present in the transformation mixes and in subsequent steps to induce gene expression mediated by the $P_{Zn}$ zinc-inducible promoter in the ectopic *bgaA* site for all strains except for IU10063 ($P_{Zn}$-*pbp2x*), which was transformed in the presence of 0.2 mM $ZnCl_2$ + 0.02 mM $MnSO_4$. IU14974 ($P_{Zn}$-*stkP*) was tested with 0.1, 0.2 and 0.4 mM $ZnCl_2$ and 1/10 concentration of $MnSO_4$. No colonies were obtained with these overexpression strains when transformed with a Δ*gpsB* amplicon in the

presence of $ZnCl_2$, while more than 500 colonies were obtained with strains that overexpressed *gpsB*, *murZ*, or *murA* (see Table 1).

[b]The Zn-induced expression of the ectopic genes in these strains have been shown to complement the respective deletions in the native site, except for IU9992 ($P_{Zn}$-*pbp1b*$^+$) because of the lack of overt phenotypes caused by Δ*pbp1b*.

**SUPPLEMENTAL REFERENCES**

Cleverley, R.M., Rutter, Z.J., Rismondo, J., Corona, F., Tsui, H.T., Alatawi, F.A., Daniel, R.A., Halbedel, S., Massidda, O., Winkler, M.E., and Lewis, R.J. (2019) The cell cycle regulator GpsB functions as cytosolic adaptor for multiple cell wall enzymes. *Nat Comm* **10**:261. doi: 10.1038/s41467-018-08056-2.

Hoskins, J., Alborn, W.E., Jr., Arnold, J., Blaszczak, L.C., Burgett, S., DeHoff, B.S., Estrem, S.T., Fritz, L., Fu, D.J., Fuller, W., Geringer, C., Gilmour, R., Glass, J.S., Khoja, H., Kraft, A.R., Lagace, R.E., LeBlanc, D.J., Lee, L.N., Lefkowitz, E.J., Lu, J., Matsushima, P., McAhren, S.M., McHenney, M., McLeaster, K., Mundy, C.W., Nicas, T.I., Norris, F.H., O'Gara, M., Peery, R.B., Robertson, G.T., Rockey, P., Sun, P.M., Winkler, M.E., Yang, Y., Young-Bellido, M., Zhao, G., Zook, C.A., Baltz, R.H., Jaskunas, S.R., Rosteck, P.R., Jr., Skatrud, P.L., and Glass, J.I. (2001) Genome of the bacterium *Streptococcus pneumoniae* strain R6. *J Bacteriol* **183**: 5709-5717.

Land, A.D., Tsui, H.C., Kocaoglu, O., Vella, S.A., Shaw, S.L., Keen, S.K., Sham, L.T., Carlson, E.E., and Winkler, M.E. (2013) Requirement of essential Pbp2x and GpsB for septal ring closure in *Streptococcus pneumoniae* D39. *Molec Microbiol* **90**: 939-955.

Land, A.D., and Winkler, M.E. (2011) The requirement for pneumococcal MreC and MreD is relieved by inactivation of the gene encoding PBP1a. *J Bacteriol* **193**: 4166-4179.

Lanie, J.A., Ng, W.L., Kazmierczak, K.M., Andrzejewski, T.M., Davidsen, T.M., Wayne, K.J., Tettelin, H., Glass, J.I., and Winkler, M.E. (2007) Genome sequence of Avery's virulent serotype 2 strain D39 of *Streptococcus pneumoniae* and comparison with that of unencapsulated laboratory strain R6. *J Bacteriol* **189**: 38-51.

Li, Y., Thompson, C.M., and Lipsitch, M. (2014) A modified Janus cassette (Sweet Janus) to improve allelic replacement efficiency by high-stringency negative selection in Streptococcus pneumoniae. *PloS One* **9**: e100510.

Mura, A., Fadda, D., Perez, A.J., Danforth, M.L., Musu, D., Rico, A.I., Krupka, M., Denapaite, D., Tsui, H.T., Winkler, M.E., Branny, P., Vicente, M., Margolin, W., and

Massidda, O. (2017) Roles of the essential protein FtsA in cell growth and division in *Streptococcus pneumoniae*. *J Bacteriol* **199**.

Perez, A.J., Cesbron, Y., Shaw, S.L., Bazan Villicana, J., Tsui, H.T., Boersma, M.J., Ye, Z.A., Tovpeko, Y., Dekker, C., Holden, S., and Winkler, M.E. (2019) Movement dynamics of divisome proteins and PBP2x:FtsW in cells of *Streptococcus pneumoniae*. *Proc Natl Acad Sci USA* **116**: 3211-3220.

Ramos-Montanez, S., Tsui, H.C., Wayne, K.J., Morris, J.L., Peters, L.E., Zhang, F., Kazmierczak, K.M., Sham, L.T., and Winkler, M.E. (2008) Polymorphism and regulation of the spxB (pyruvate oxidase) virulence factor gene by a CBS-HotDog domain protein (SpxR) in serotype 2 *Streptococcus pneumoniae*. *Molec Microbiol* **67**: 729-746.

Rued, B.E., Zheng, J.J., Mura, A., Tsui, H.T., Boersma, M.J., Mazny, J.L., Corona, F., Perez, A.J., Fadda, D., Doubravova, L., Buriankova, K., Branny, P., Massidda, O., and Winkler, M.E. (2017) Suppression and synthetic-lethal genetic relationships of Δ*gpsB* mutations indicate that GpsB mediates protein phosphorylation and penicillin-binding protein interactions in *Streptococcus pneumoniae* D39. *Molec Microbiol* **103**: 931-957.

Slager, J., Aprianto, R., and Veening, J.W. (2018) Deep genome annotation of the opportunistic human pathogen *Streptococcus pneumoniae* D39. *Nuc Acids Res* **46**: 9971-9989.

Tsui, H.C., Keen, S.K., Sham, L.T., Wayne, K.J., and Winkler, M.E. (2011) Dynamic distribution of the SecA and SecY translocase subunits and septal localization of the HtrA surface chaperone/protease during *Streptococcus pneumoniae* D39 cell division. *mBio* **2**: e00202-00211.

Tsui, H.C., Zheng, J.J., Magallon, A.N., Ryan, J.D., Yunck, R., Rued, B.E., Bernhardt, T.G., and Winkler, M.E. (2016) Suppression of a deletion mutation in the gene encoding essential PBP2b reveals a new lytic transglycosylase involved in peripheral peptidoglycan synthesis in *Streptococcus pneumoniae* D39. *Molec Microbiol* **100**: 1039-1065.

Tsui, H.T., Boersma, M.J., Vella, S.A., Kocaoglu, O., Kuru, E., Peceny, J.K., Carlson, E.E., VanNieuwenhze, M.S., Brun, Y.V., Shaw, S.L., and Winkler, M.E. (2014) Pbp2x localizes separately from Pbp2b and other peptidoglycan synthesis proteins during later stages of cell division of *Streptococcus pneumoniae* D39. *Molec Microbiol* **94**: 21-40.

Wayne, K.J., Sham, L.T., Tsui, H.C., Gutu, A.D., Barendt, S.M., Keen, S.K., and Winkler, M.E. (2010) Localization and cellular amounts of the WalRKJ (VicRKX) two-component regulatory system proteins in serotype 2 *Streptococcus pneumoniae*. *J Bacteriol* **192**: 4388-4394.

Zheng, J.J., Perez, A.J., Tsui, H.T., Massidda, O., and Winkler, M.E. (2017) Absence of the KhpA and KhpB (JAG/EloR) RNA-binding proteins suppresses the requirement for PBP2b by overproduction of FtsA in *Streptococcus pneumoniae* D39. *Molec Microbiol* **106**: 793-814.

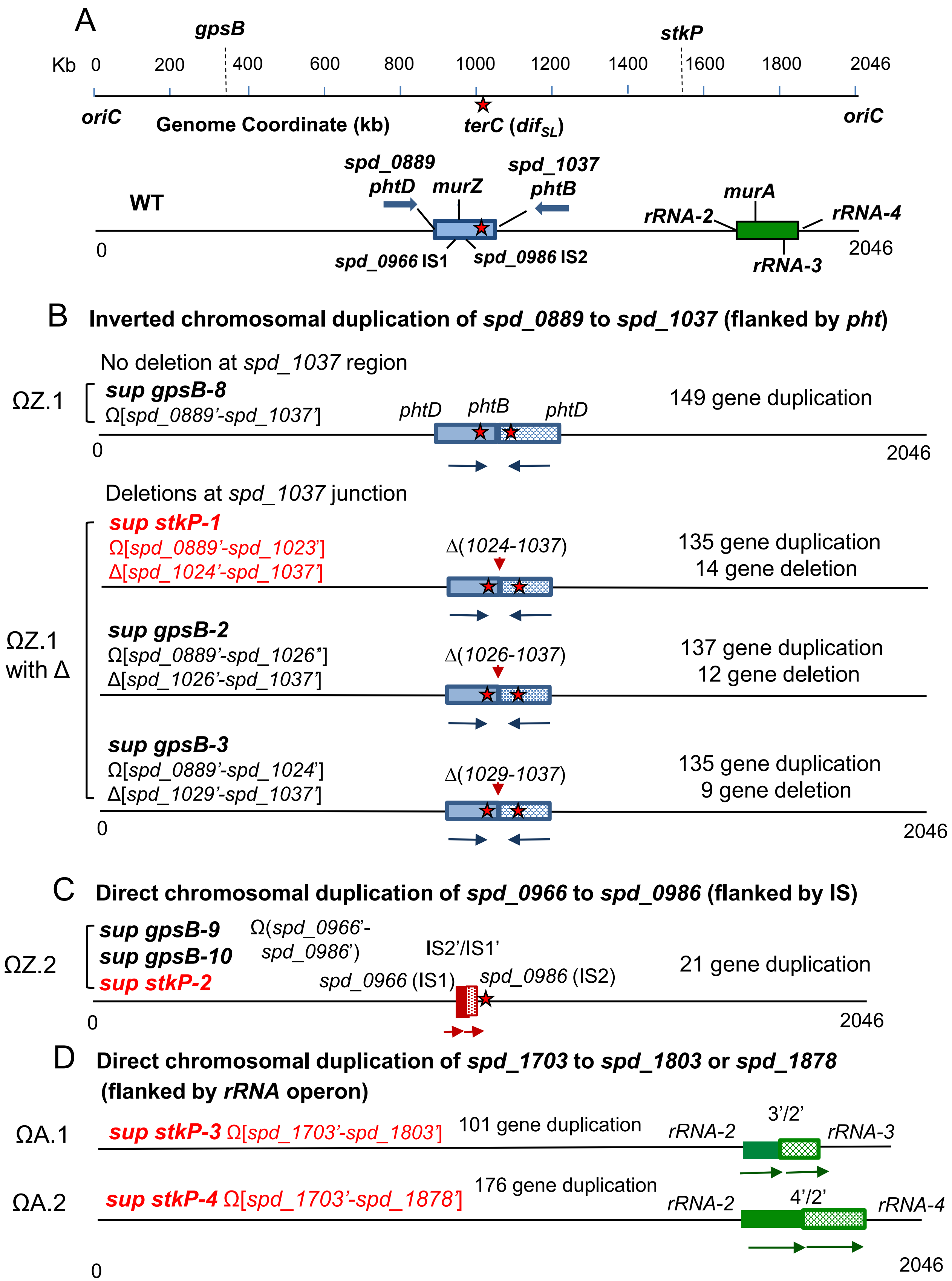
A
gpsB
stkP
Kb 0 200 400 600 800 1000 1200 1400 1600 1800 2046
oriC
Genome Coordinate (kb)
terC (difSL)
oriC
spd_0889
phtD
murZ
spd_1037
phtB
WT
murA
rRNA-2
rRNA-4
0
spd_0966 IS1
spd_0986 IS2
rRNA-3
2046
B Inverted chromosomal duplication of spd_0889 to spd_1037 (flanked by pht)
No deletion at spd_1037 region
ΩZ.1
sup gpsB-8
Ω[spd_0889'-spd_1037']
phtD phtB phtD
149 gene duplication
0
2046
Deletions at spd_1037 junction
sup stkP-1
Ω[spd_0889'-spd_1023']
Δ[spd_1024'-spd_1037']
Δ(1024-1037)
135 gene duplication
14 gene deletion
ΩZ.1 with Δ
sup gpsB-2
Ω[spd_0889'-spd_1026"]
Δ[spd_1026'-spd_1037']
Δ(1026-1037)
137 gene duplication
12 gene deletion
sup gpsB-3
Ω[spd_0889'-spd_1024']
Δ[spd_1029'-spd_1037']
Δ(1029-1037)
135 gene duplication
9 gene deletion
0
2046
C Direct chromosomal duplication of spd_0966 to spd_0986 (flanked by IS)
ΩZ.2
sup gpsB-9
sup gpsB-10
sup stkP-2
Ω(spd_0966'-spd_0986')
IS2'/IS1'
spd_0966 (IS1)
spd_0986 (IS2)
21 gene duplication
0
2046
D Direct chromosomal duplication of spd_1703 to spd_1803 or spd_1878 (flanked by rRNA operon)
ΩA.1
sup stkP-3 Ω[spd_1703'-spd_1803']
101 gene duplication
rRNA-2
3'/2'
rRNA-3
ΩA.2
sup stkP-4 Ω[spd_1703'-spd_1878']
176 gene duplication
rRNA-2
4'/2'
rRNA-4
0
2046

Fig. S1

**Figure S1. Duplication and duplication/deletion regions in Δ*gpsB* and Δ*stkP* suppressor strains.** A. Chromosome coordinates representation in a linear scheme in kb. *terC* (*dif*$_{SL}$ sequence, red star) is located at 83 to 53 bp upstream of *xerS* (*spd_1023*) as described by (Le Bourgeois *et al.*, 2007). (B) The duplication patterns are grouped into ΩZ.1 or ΩZ.2 (duplication of *murZ* region) or ΩA.1 or ΩA.2 (duplication of *murA* region). ΩZ.1 duplications are flanked by *phtD* and *phtB*, while ΩZ.2 duplications are bordered by degenerate IS elements *spd_0966* and *spd_0986.* ΩA.1 or ΩA.2 are bordered by tRNA/rRNA gene clusters. In ΩZ.1, represented by *sup gpsB-8,* large inverted duplications (shaded region, >135 genes) are flanked by *phtB* (*spd_1037*), and *phtD* (*spd_0889*), two oppositely transcribed genes with identical 1324 nt sequence at the 3' end. No flanking deletion is found in this strain. In group ΩZ.1 with Δ, duplication of the *phtD* to *phtB* region is accompanied by gene deletions in the *spd_1037 region.* In *sup stkP-1,* the regions from ≈50 bp upstream of *spd_1024* to *spd_1037* at both duplications are deleted, leading to the resulting genotype of Ω[*spd_0889'*-*spd_1023'*] Δ[*spd_1024*-*spd_1037'*]. In *sup gpsB-2*, the deletion junction is within *spd_1026,* leading to the resulting genotype of Ω[*spd_0889'*-*spd_1026'*] Δ[*spd_1026'*-*spd_1037'*]. In *sup gpsB-3*, the deletion occurs between *spd_1029* and *spd_1037* in one segment and *spd_1037* to *spd_1024* in the other segment, leading to Ω[*spd_0889'*-*spd_1024'*] Δ[*spd_1029'*-*spd_1037'*] (see Figure 3 for detail). Two copies of *terC* (red star) are present in ΩZ.1 and 'ΩZ.1 with Δ' classes. (C) ΩZ.2 suppressors contain 21-gene duplication flanked by *spd_0966* to *spd_0986,* two degenerate transposase IS1167 genes. *Sup gpsB-9* and *sup stkP-2* contain tandem duplications, while *sup gpsB-10* contains a higher level amplification. (D) In ΩA.1 and ΩA.2 suppressors, the duplications are flanked by tRNA/rRNA gene clusters, *rRNA-2, rRNA-3*, or *rRNA-4.* Duplicated fragments are represented by shaded segments. Arrows represent the sequence directions.

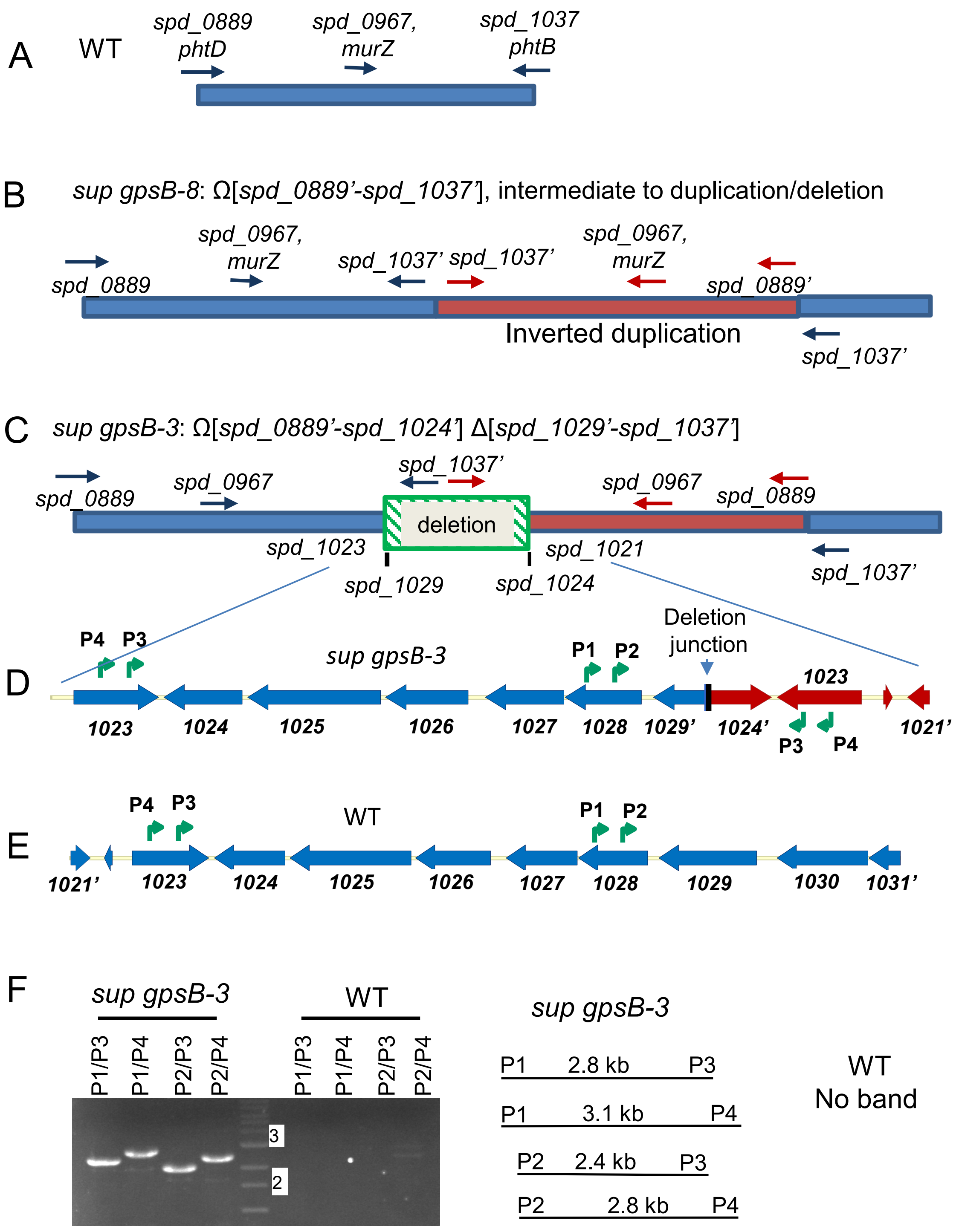

A
WT
spd_0889
phtD
spd_0967,
murZ
spd_1037
phtB
B
sup gpsB-8: Ω[spd_0889'-spd_1037'], intermediate to duplication/deletion
spd_0889
spd_0967,
murZ
spd_1037'
spd_1037'
spd_0967,
murZ
spd_0889'
Inverted duplication
spd_1037'
C
sup gpsB-3: Ω[spd_0889'-spd_1024'] Δ[spd_1029'-spd_1037']
spd_0889
spd_0967
spd_1037'
deletion
spd_0967
spd_0889
spd_1023
spd_1021
spd_1029
spd_1024
spd_1037'
D
P4
P3
sup gpsB-3
P1
P2
Deletion
junction
1023
1023
1024
1025
1026
1027
1028
1029'
1024'
P3
P4
1021'
E
P4
P3
WT
P1
P2
1021'
1023
1024
1025
1026
1027
1028
1029
1030
1031'
F
sup gpsB-3
WT
P1/P3
P1/P4
P2/P3
P2/P4
P1/P3
P1/P4
P2/P3
P2/P4
3
2
sup gpsB-3
P1 2.8 kb P3
P1 3.1 kb P4
P2 2.4 kb P3
P2 2.8 kb P4
WT
No band

Fig. S2

**Figure S2. Model for formation of the chromosomal *sup gpsB-3* duplication/deletion (ΩZ.1 with Δ) that suppresses Δ*gpsB* by large inverted duplication followed by small deletion of the duplication junction.** (A) Arrangement of *spd_0889* to *spd_1037* in a WT strain. (B) Chromosomal arrangement of *sup gpsB-8* which contains an inverted duplication (orange segment) from *spd_0889* (*phtD*) to *spd_1037* (*phtB*). (C) In *sup gpsB-3*, inverted duplication was followed by the deletion between *spd_1029* and *spd_1037* in one segment and *spd_1037* to *spd_1024* in the other segment, leading to Ω[*spd_0889*'-*spd_1024'*] Δ[*spd_1029*'-*spd_1037'*]. (D) Enlargement of the deletion junction in *sup gpsB-3* between *spd_1023* of one segment and *spd_1021* of the other segment. Primers P1, P2, P3, and P4 are used to confirm the rearrangement. (E) Location of primers P1 to P4 in the WT strain. (F) PCR analysis to confirm the chromosomal arrangement shown in C. Bands of expected sizes were obtained with four sets of primers using *sup gpsB-3* strain as DNA template, while no bands was obtained using DNA obtained from the WT parent (IU1945).

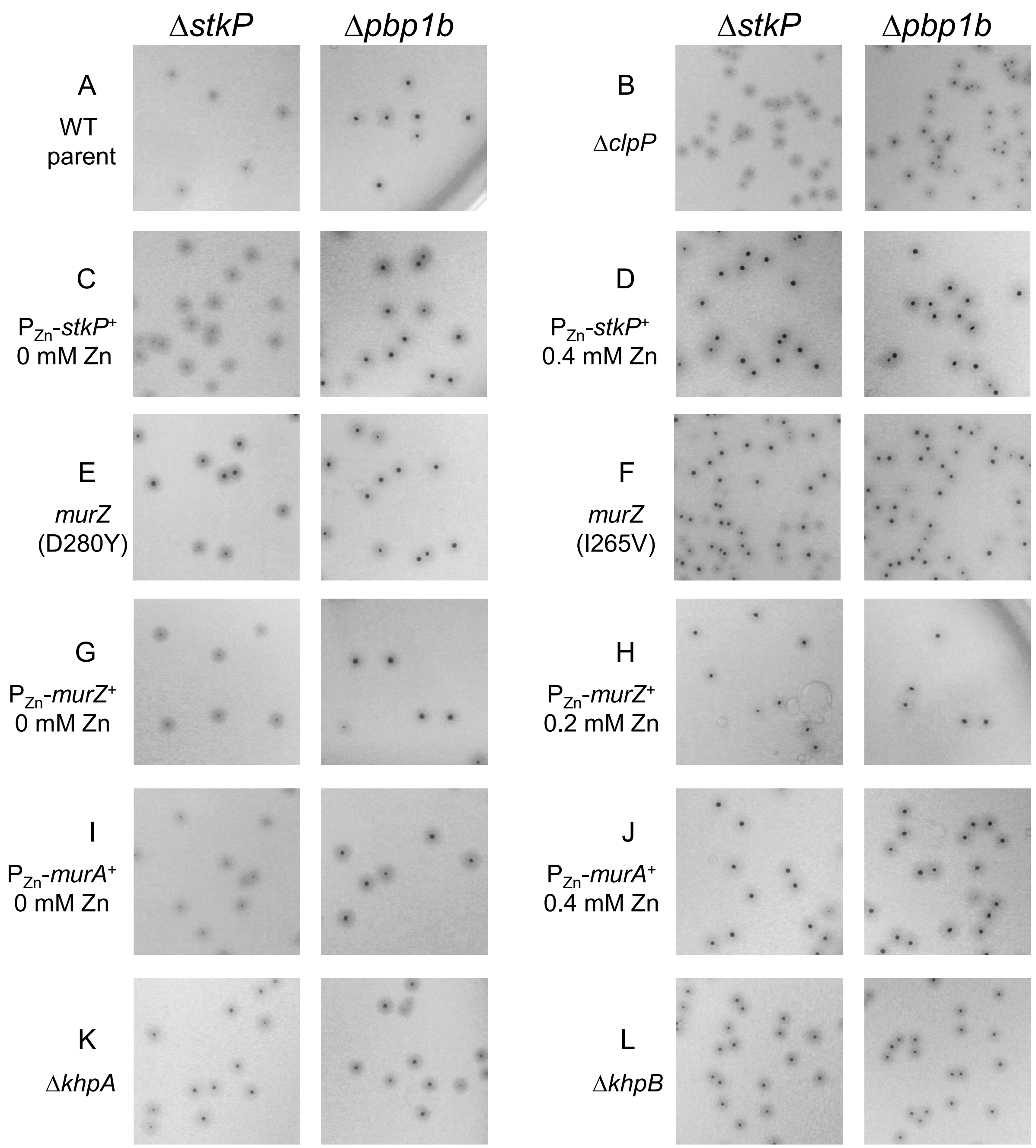

**Figure S3. *murZ*(D280Y), *murZ*(I265V)*, ΔkhpA/B* mutations, and overexpression of *murZ* or *murA* suppress the faint colony phenotype of Δ*stkP* on TSAII-BA plates.** (A) Parent D39 Δ*cps rpsL1* strain (IU1824), (B) Δ*clpP* (IU17138), (C and D) *stkP*$^+$//P$_{Zn}$-*stkP*$^+$ (IU14974), (E) *murZ*(D280Y (IU13438), (F) *murZ*(I265V (IU14210), (G and H) *murZ*$^+$//P$_{Zn}$-*murZ*$^+$ (IU13393), (I and J) *murA*$^+$//P$_{Zn}$-*murA*$^+$ (IU13395), (K) Δ*khpA* (IU9036)*,* and (L) Δ*khpB* (IU10592) were transformed with a Δ*stkP*::P$_c$-*erm* or a positive control amplicon Δ*pbp1b*::P$_c$-*erm* as described in *Experimental procedures.* Images of colonies on the TSAII-BA transformation plates were taken with a light source under the plates after 20h incubation in a 37°C 5% $CO_2$ incubator.

Fig. S3

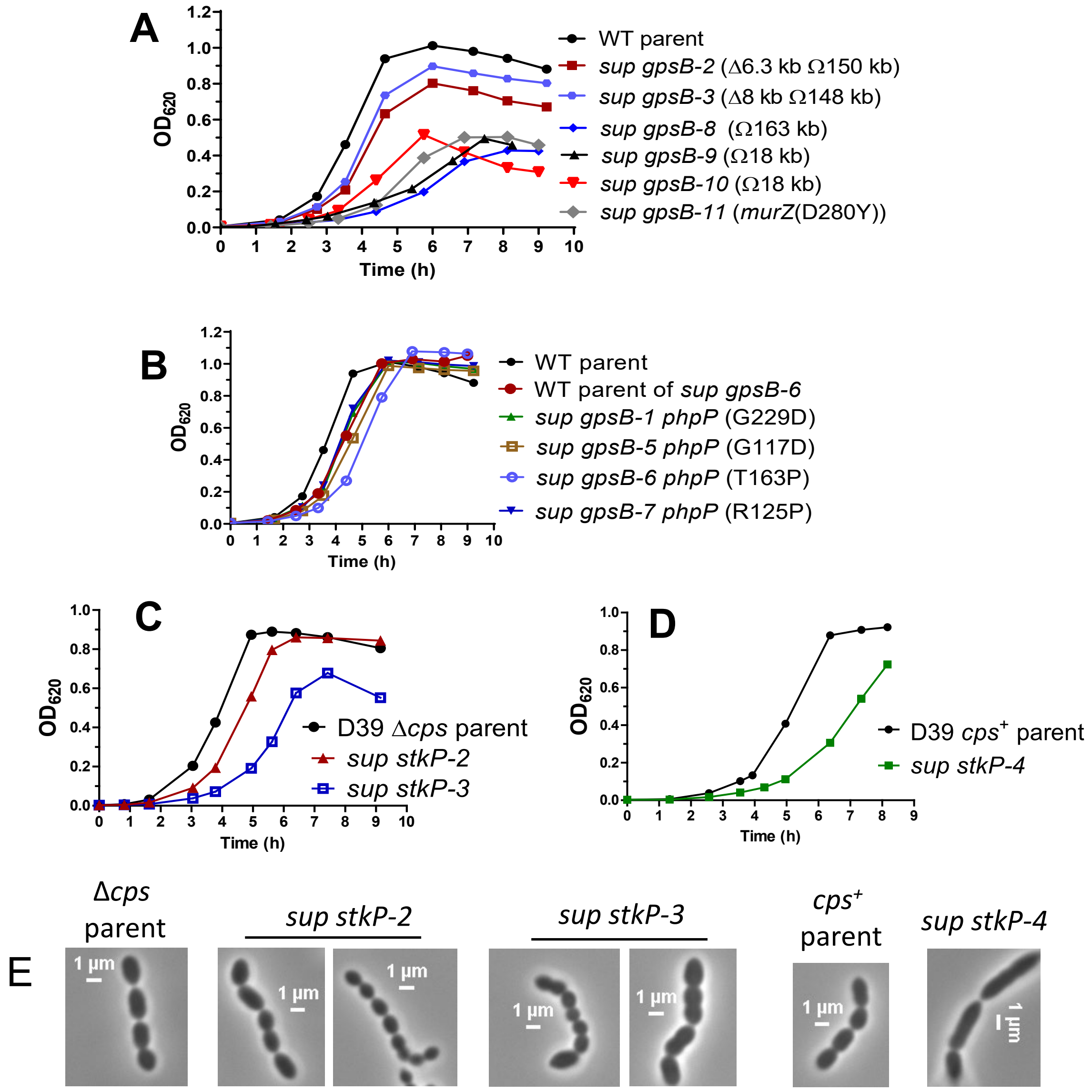

**Figure S4. Growth profiles of Δ*gpsB* and Δ*stkP* suppressor strains.** (A) and (B), Δ*gpsB* suppressor strains with mutations in *phpP* exhibit better growth profile compared to Δ*gpsB* suppressor strains with large chromosomal deletion. (A) Growth curves of D39 Δ*cps* WT parent (IU1945) and Δ*gpsB* suppressor strains with large chromosomal duplication and deletion (*sup gpsB-2 and -3*), strains with large duplications (*sup gpsB -8* to *-10)* and *sup gpsB -11* which contains a *murZ*(D280Y) mutation. (B) Growth curves of WT parents D39 Δ*cps* (IU1945), and D39 Δ*cps rpsL1* (IU1824), and Δ*gpsB* suppressor strains containing mutations in *phpP*. (C) Growth curves of D39 Δ*cps* WT parent (IU1945) and *sup stkP-2* and *-3* strains. (D) Growth curves of D39 WT parent (IU1690) and *sup stkP-4* strain. (E) Phase contrast images of parents, and *sup stkP-2 to -4* strains. Growth curve and micrographs of *sup stkP-1* are shown in Fig. S20. Detailed summaries of genotypes, growth rates and growth yields of various Δ*gpsB* and Δ*stkP* *sup* strains are listed in Table 1 and Table 3, respectively.

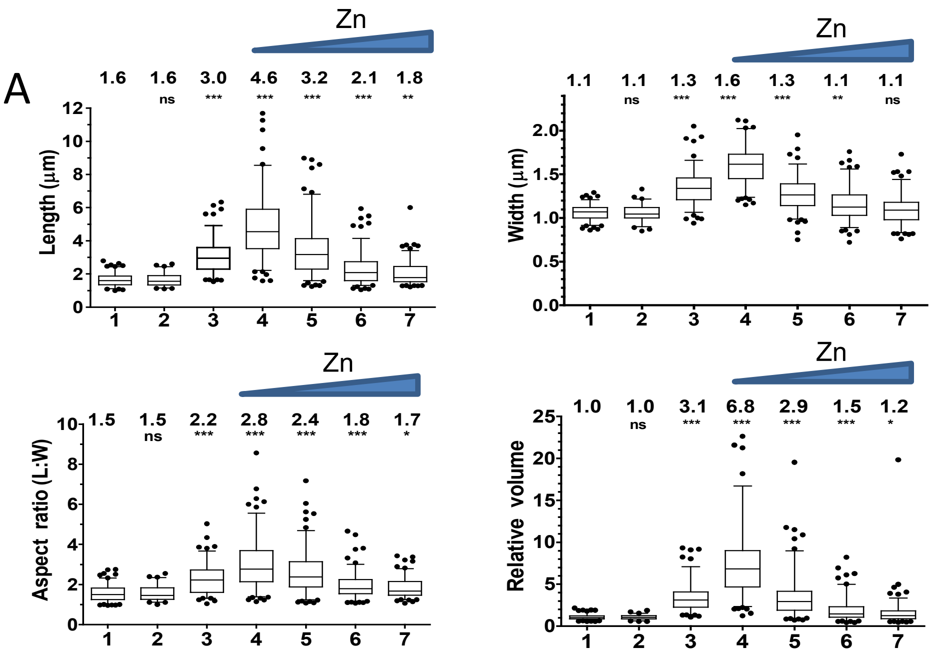

| | Genotype | Zn (mM) |
|---|---|---|
| 1 | WT | 0 |
| 2 | | 0.4 |
| 3 | Δ*gpsB murZ* (D280Y) | 0 |
| 4 | Δ*gpsB* $P_{Zn}$-*murZ*$^{+}$ | 0 |
| 5 | | 0.1 |
| 6 | | 0.2 |
| 7 | | 0.3 |

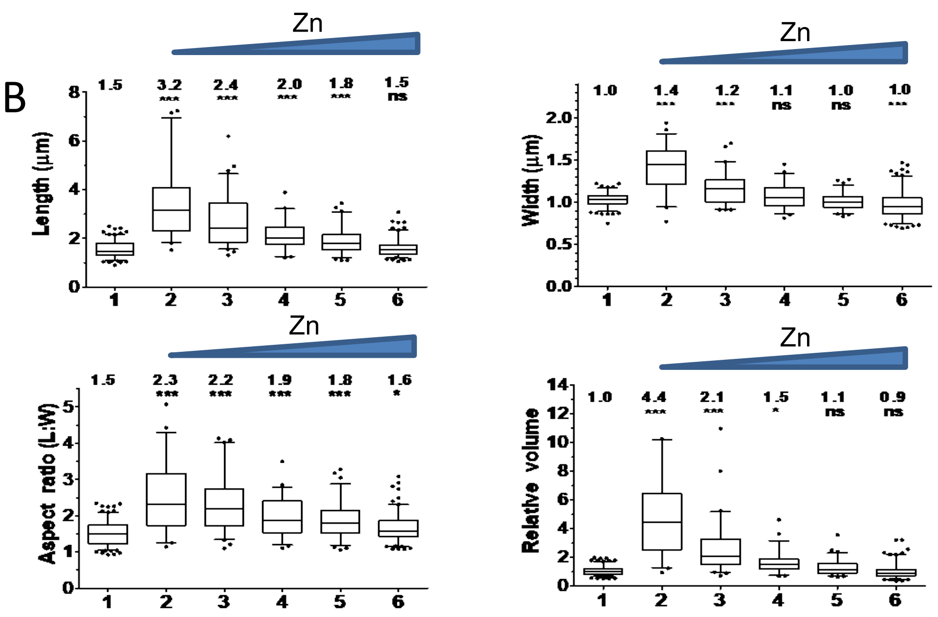

| | Genotype | Zn (mM) |
|---|---|---|
| 1 | WT | 0 |
| 2 | Δ*gpsB* $P_{Zn}$-*murA*$^{+}$ | 0 |
| 3 | | 0.2 |
| 4 | | 0.3 |
| 5 | | 0.4 |
| 6 | | 0.5 |

**Figure S5. Box-and-whisker plots of cell dimensions of *murZ*(D280Y) and overexpression strains of *murZ* and *murA* in a Δ*gpsB* background.** (A) Box-and-whisker plots (whiskers, 5 and 95 percentile) of cell lengths, widths, aspect ratios (cell length to width) and relative cell volumes of strains grown without or with indicated ($Zn^{2+}$/(1/10)$Mn^{2+}$) shown in Fig. 4. For both (A) and (B), P values were obtained by one-way ANOVA analysis (GraphPad Prism, Kruskal-Wallis test). *, **, *** and ns denote p<0.05, p<0.01, p<0.001, not significant, respectively when compared to WT.

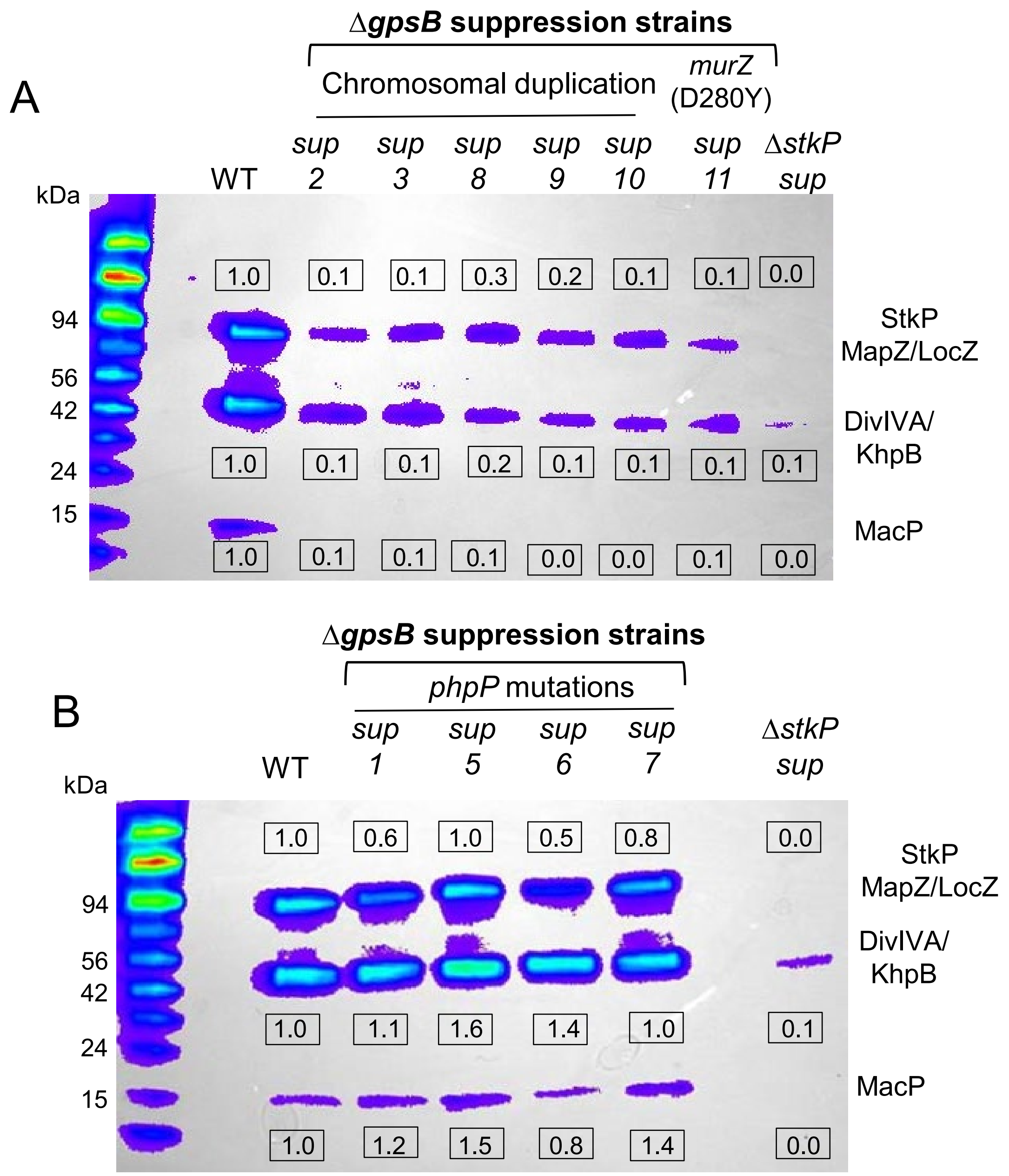

**Figure S6. Protein phosphorylation profiles in Δ*gpsB* suppression strains.** Western blot with α-pThr antibody to detect protein phosphorylation on Thr resides for WT D39 Δ*cps* parent strain (IU1945), Δ*gpsB* suppressor strains listed in Table 1, and a Δ*stkP* strain that contains uncharacterized suppressor mutation(s) (IU7923). Mean relative values of band intensities (±SEM) compared to the wild-type (WT) strain indicated for the phosphorylated MapZ/StkP, DivIVA/KhpB or MacP bands are shown in boxes above or below the blots. (A) Western blot for Δ*gpsB* suppressor strains containing chromosomal duplications and deletions (*sup2* and *sup3*), chromosomal duplications (*sup8*, *sup9* and *sup10*), or *murZ*(D280Y) mutation (*sup11*). (B) Western blot for Δ*gpsB* suppressor strains containing mutations in *phpP*.

Fig. S6

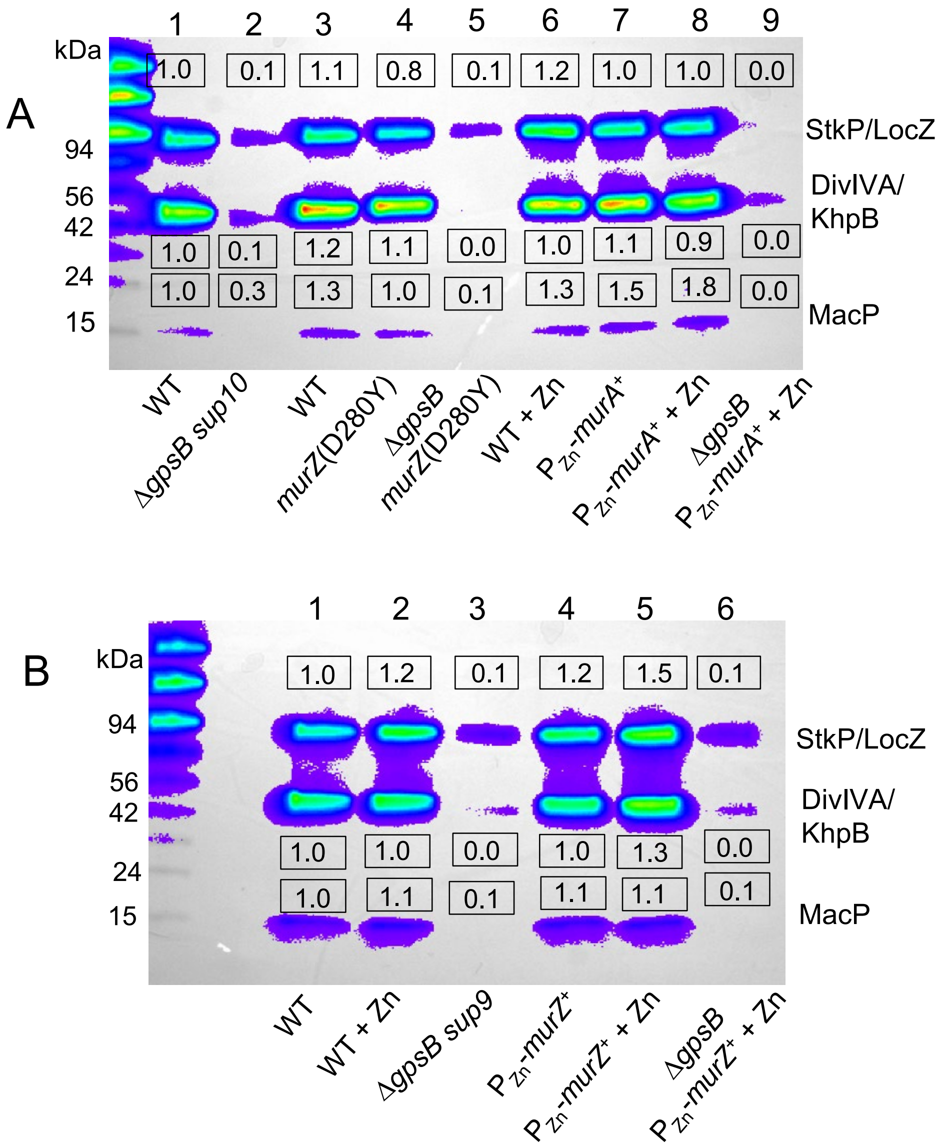

**Figure S7. Overexpression of MurZ or MurA or the presence of *murZ*(D280Y) suppresses Δ*gpsB* lethality by a protein-phosphorylation independent mechanism.** (A). Western blot with α-pThr antibody to detect protein phosphorylation on Thr resides for Δ*gpsB* strains containing *murZ*(D280Y) mutation, or *murA* overexpression. Mean relative values of band intensities (±SEM) compared to the wild-type (WT) strain indicated for the phosphorylated MapZ/StkP, DivIVA or MacP bands are shown in boxes above or below the blots. Strains 1 to 9 used for A are 1, IU1945 (WT D39 Δ*cps)*; 2, IU11918; 3, IU1824 (WT D39 Δ*cps rpsL1*); 4, IU13439; 5, IU13485; 6, IU1945 + 0.5 mM ($Zn^{2+}$/(1/10)$Mn^{2+}$) ; 7, IU11079; 8, IU11079 + 0.5 mM ($Zn^{2+}$/(1/10)$Mn^{2+}$); 9, IU13757 + 0.5 mM ($Zn^{2+}$/(1/10)$Mn^{2+}$). (B) Western blot for Δ*gpsB* suppressor strains overexpressing *murZ*. Strains are listed as follows: 1, IU1945; 2, IU1945 + 0.5 mM ($Zn^{2+}$/(1/10)$Mn^{2+}$); 3, IU11846; 4, IU11077; 5, IU11077 + 0.2 mM ($Zn^{2+}$/(1/10)$Mn^{2+}$); 6, IU13756 + 0.2 mM ($Zn^{2+}$/(1/10)$Mn^{2+}$).

Fig. S7

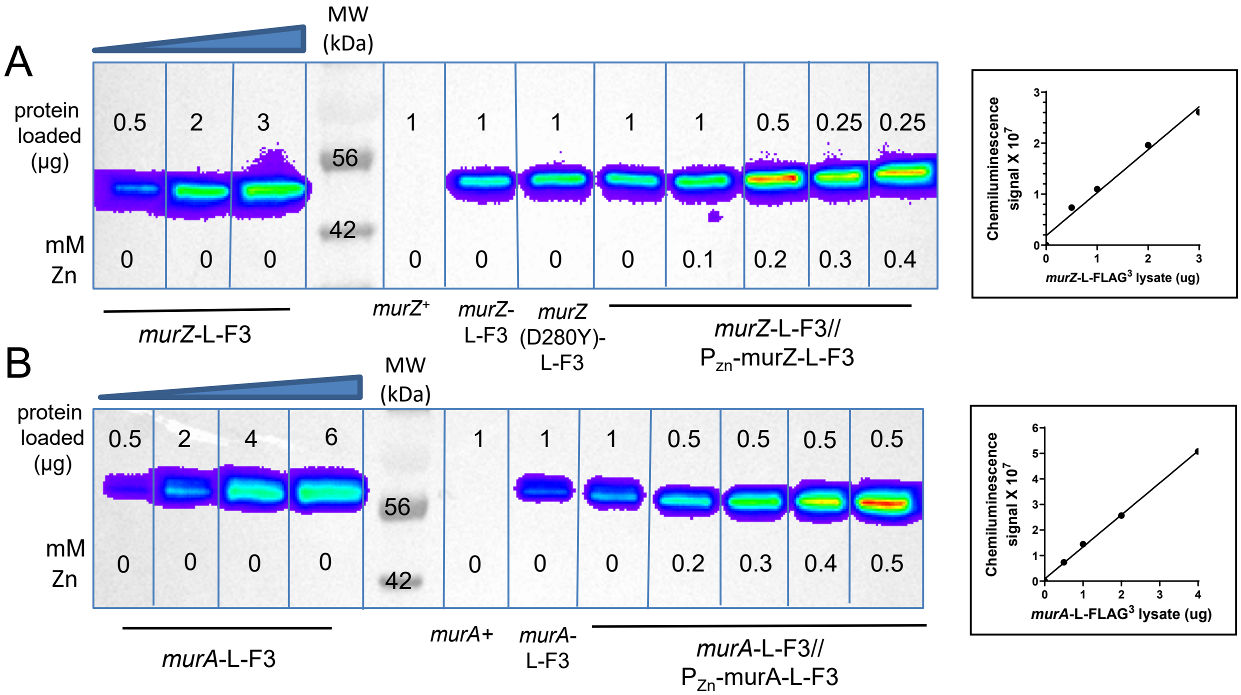

**Figure S8**. **Quantitation of relative MurZ-L-F3 (A) and MurA-L-F3 (B) cellular amounts by western blot using anti-FLAG antibody.** Strains and growth conditions are listed in legend to Fig. 5. The µg amounts of total protein loaded for each strain, and ($Zn^{2+}$/(1/10)$Mn^{2+}$) concentrations present in the BHI growth media are shown above and below the bands, respectively. The amounts of proteins loaded per lane for each sample were adjusted so that the intensity values are within the linear range obtained with the standard curve using various µg amounts of IU13502 or IU14028 samples. Plot of µg of lysate obtained from IU13502 or IU14028 loaded vs chemiluminescence signal intensities are shown to the right of the blots.

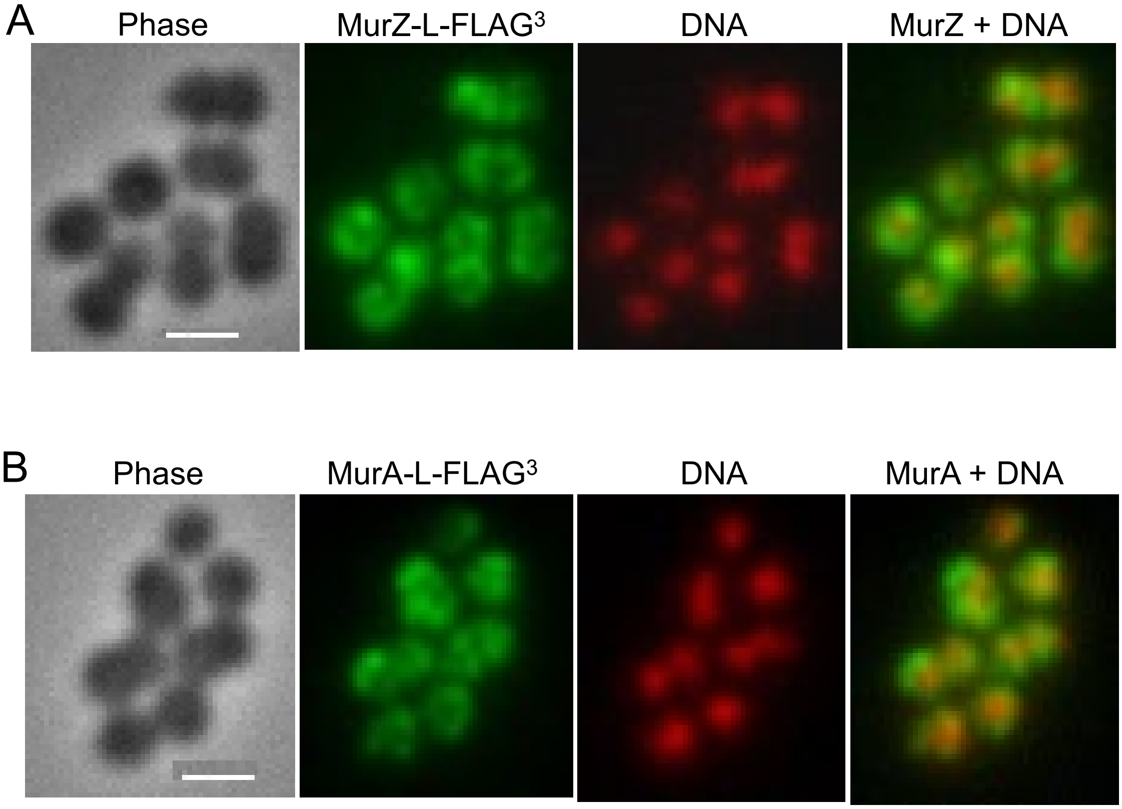

**Figure S9. MurZ and MurA show general cytoplasmic distribution.** Immunofluorescence microscopy was performed as described in *Experimental procedures* using (A) IU13502 (*murZ*-L-FLAG$^3$) or (B) IU14028 (*murA*-L-FLAG$^3$). Nucleoid DNA was labeled with a mounting media SlowFade gold antifade reagent containing DNA staining reagent DAPI. Scale bar: 1 μm.

Fig. S9

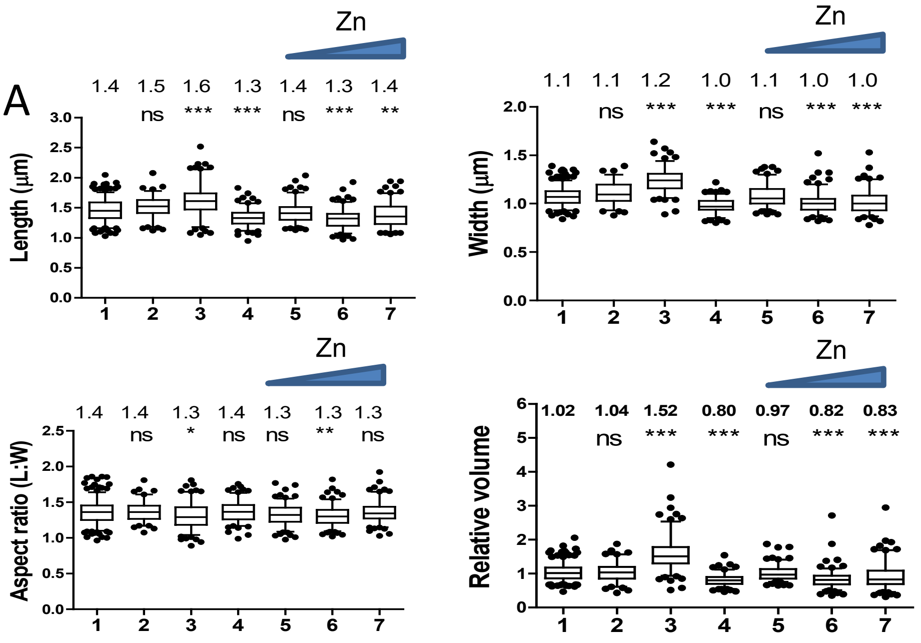

| | Genotype | Zn (mM) |
|---|---|---|
| 1 | WT | 0 |
| 2 | | 0.4 |
| 3 | Δ*murZ* | 0 |
| 4 | *murZ* (D280Y) | 0 |
| 5 | $P_{Zn}$-*murZ*⁺ | 0 |
| 6 | | 0.2 |
| 7 | | 0.4 |

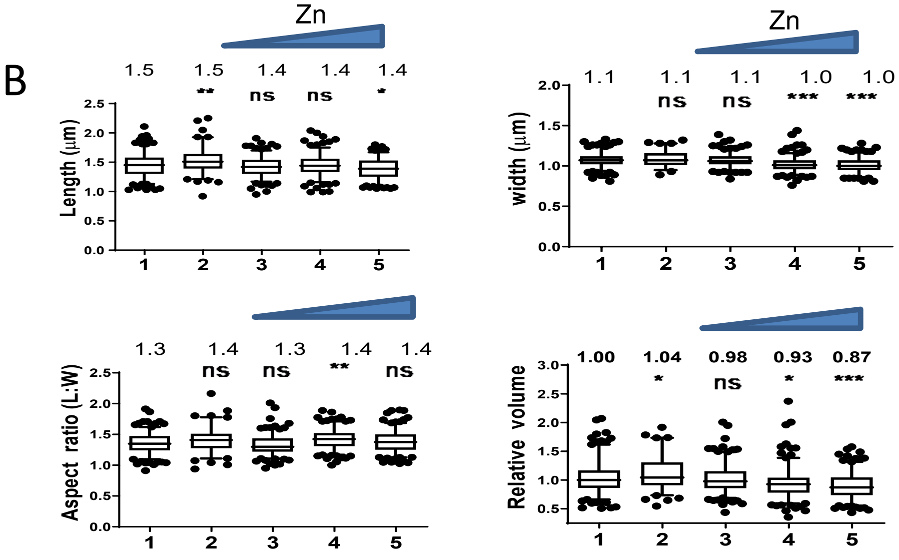

| | Genotype | Zn (mM) |
|---|---|---|
| 1 | WT | 0 |
| 2 | Δ*murA* | 0 |
| 3 | $P_{Zn}$-*murA*⁺ | 0 |
| 4 | | 0.3 |
| 5 | | 0.5 |

**Figure S10. Box-and-whisker plots of cell dimensions of *murZ*(D280Y) and overexpression strains of *murZ* and *murA*.** (A) Box-and-whisker plots (whiskers, 5 and 95 percentile) of cell lengths, widths, aspect ratios (cell length to width) and relative cell volumes of strains grown with or without ($Zn^{2+}$/(1/10)$Mn^{2+}$) of strains shown in Fig. 6. 1, WT (IU1824); 2, WT + 0.4 mM ($Zn^{2+}$/(1/10)$Mn^{2+}$); 3, Δ*murZ* (IU13536); 4, *murZ*(D280Y) (IU13438); 5, 6 and 7, *murZ*⁺//$P_{Zn}$-*murZ*⁺ (IU13393) grown in 0, 0.2, or 0.4 mM ($Zn^{2+}$/(1/10)$Mn^{2+}$), respectively. (B) 1, WT (IU1824); 2, Δ*murA* (IU13538); 3, 4 and 5, *murA*⁺//$P_{Zn}$-*murA*⁺ (IU13395) grown in 0, 0.3, or 0.5 mM ($Zn^{2+}$/(1/10)$Mn^{2+}$), respectively. P values were obtained by one-way ANOVA analysis (GraphPad Prism, Kruskal-Wallis test). *, **, *** and ns denote $p<0.05$, $p<0.01$, $p<0.001$, not significant, respectively when compared to WT.

Fig. S10

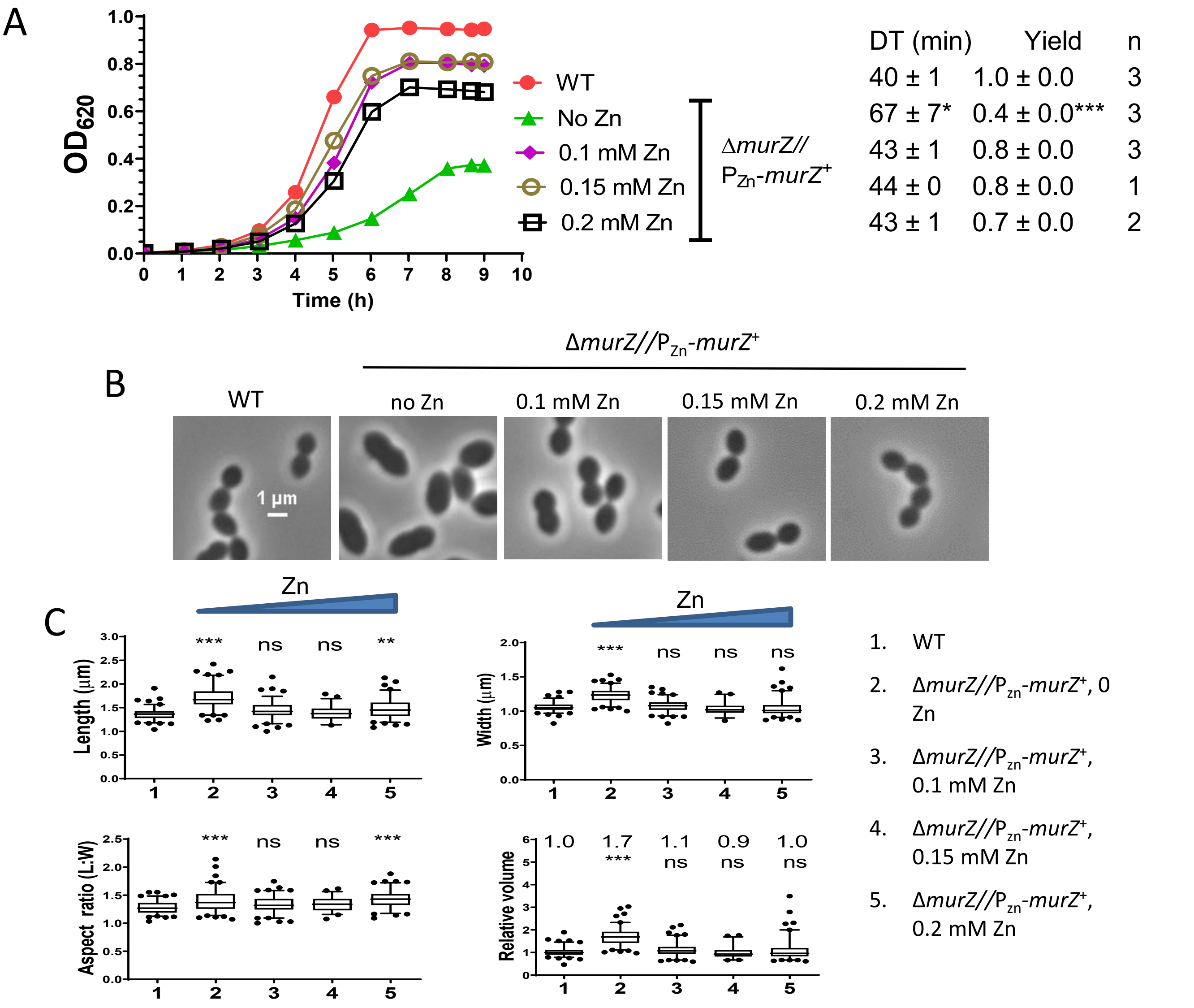

**Figure S11. Complementation of Δ*murZ* growth and morphological defects by ectopic overexpression of *murZ*.** Parent D39 Δ*cps rpsL1* strain (IU1824) was grown overnight in BHI broth with no additional ($Zn^{2+}$/(1/10)$Mn^{2}$+), and Δ*murZ*//$P_{Zn}$-*murZ*$^{+}$ (IU16259) strains was grown overnight in BHI supplemented with 0, 0.1, 0.15 or 0.2 mM ($Zn^{2+}$/(1/10)$Mn^{2+}$), and diluted with fresh BHI containing the same concentrations of $Zn^{2+}$/(1/10)$Mn^{2+}$ as the overnight cultures. (A) Representative growth curves, averages and SEMs of doubling times (DT) and maximal growth yields ($OD_{620}$) during 9 hours of growth. (B) Representative phase-contrast images taken between 3 and 4.5 h of growth for all strains and conditions. (C) Box-and-whisker plots of cell dimensions of strains grown with or without ($Zn^{2+}$/(1/10)$Mn^{2+}$). P values were obtained by one-way ANOVA analysis (GraphPad Prism, Kruskal-Wallis test). *, **, *** and ns denote p<0.05, p<0.01, p<0.001, not significant, respectively when compared to WT.

Fig. S11

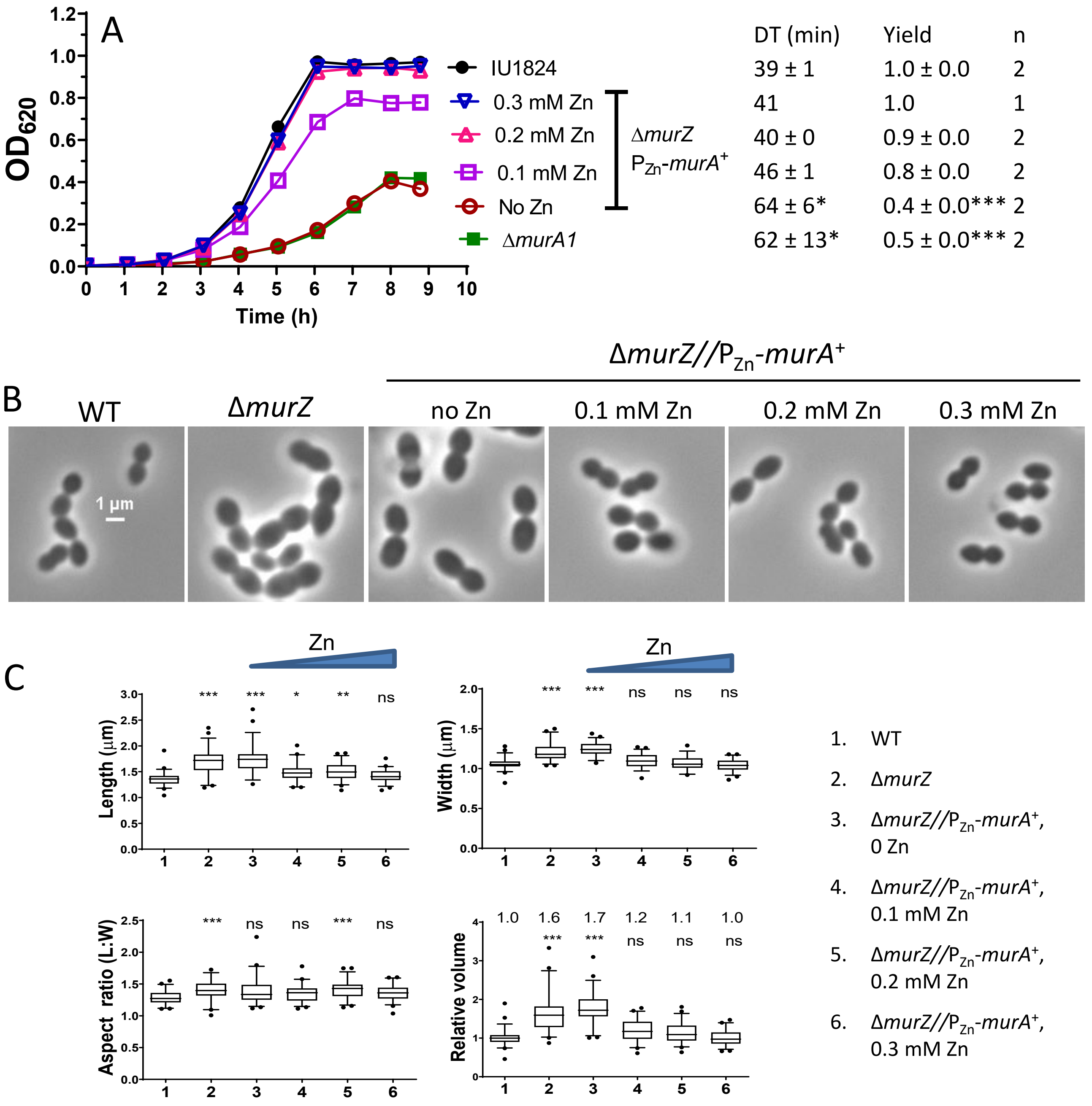

**Figure S12. Complementation of Δ*murZ* growth and morphological defects by ectopic overexpression of *murA*.** Parent D39 Δ*cps rpsL1* strain (IU1824), Δ*murZ* (IU13536), and Δ*murZ murA*//$P_{zn}$-*murA* (IU16262) strains were grown overnight and during the day in BHI broth with no additional or indicated concentrations of ($Zn^{2+}$/(1/10)$Mn^{2+}$) as described in legend to Fig. S11.

Fig. S12

D39 *cps*⁺ genetic background

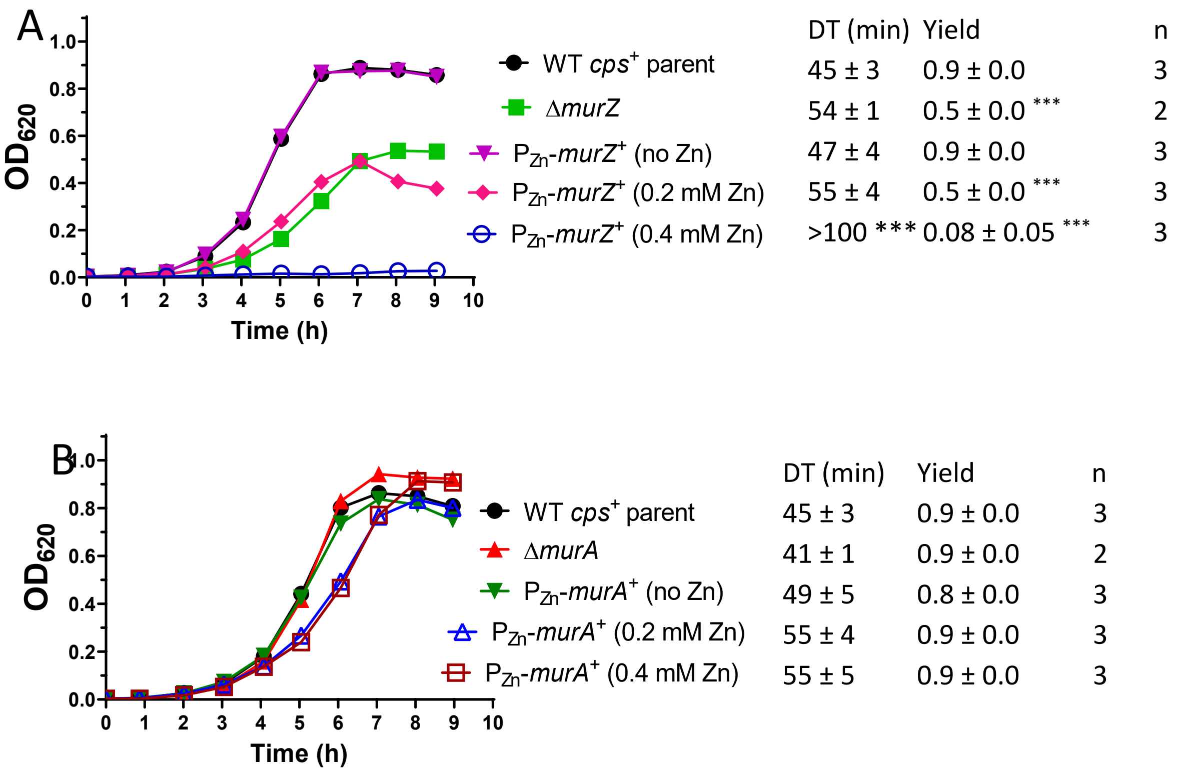

**Figure S13. Growth phenotypes of deletion and overexpression of *murZ* or *murA* in D39 encapsulated strains are similar to those in unencapsulated Δ*cps* D39 strains.** (A) Growth in BHI broth of wild-type D39 *cps*⁺ parent (IU1690), isogenic Δ*murZ*::$P_c$-*erm* (IU16176), and *murZ*//$P_{Zn}$-*murZ*⁺ (IU15879). (B) Deletion or overexpression of *murA* in an encapsulated derivative of strain D39 did not result in growth defects when cultured in BHI broth. Strains tested are D39 *cps*⁺ parent (IU1690), isogenic Δ*murA* (IU16178), and *murA*⁺//$P_{Zn}$-*murA*⁺ (IU15880).

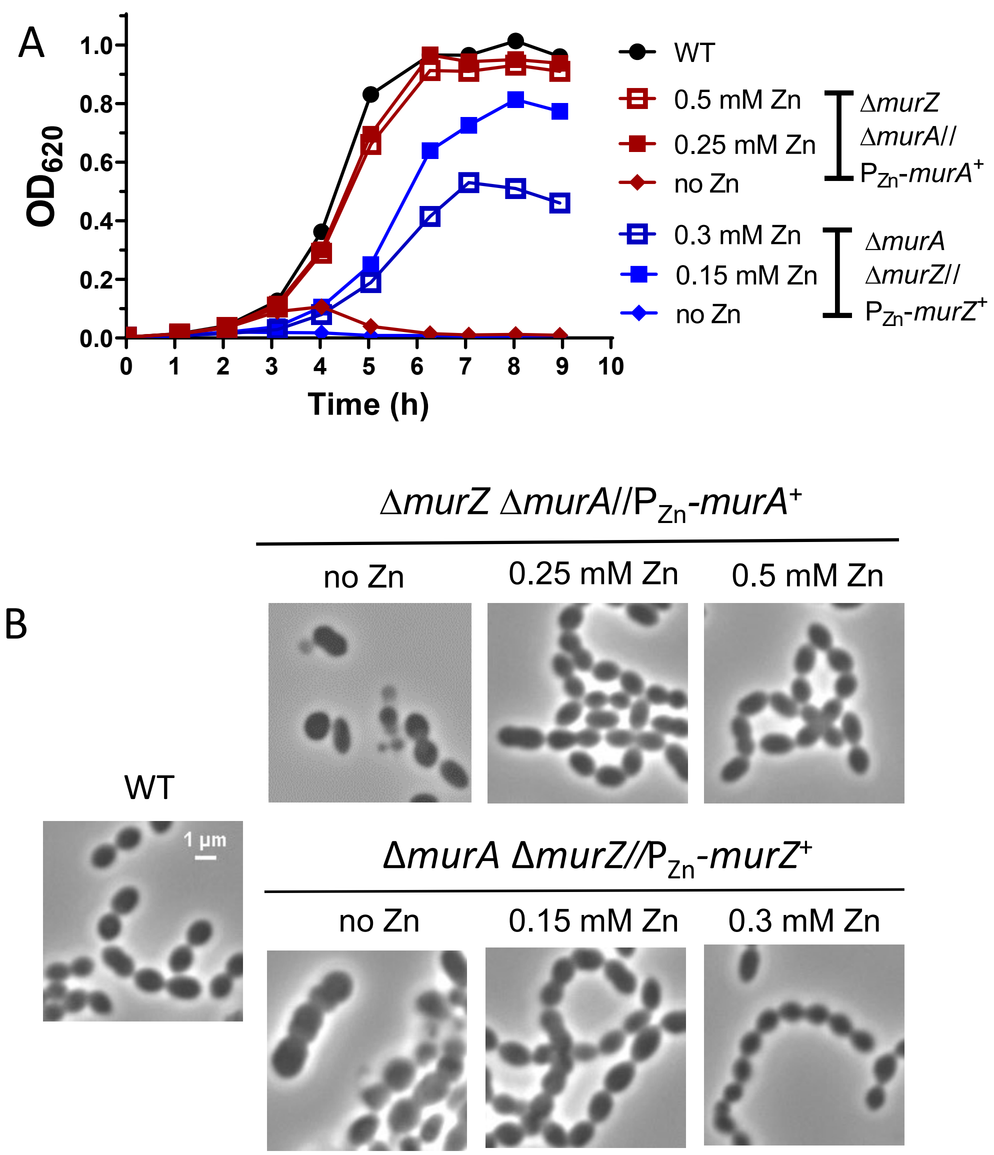

**Figure S14. Depletion of MurA in a Δ*murZ* strain, or depletion of MurZ in a Δ*murA* strain results in cell lysis, but not cell elongation.** Parent D39 Δ*cps rpsL1* strain (IU1824), Δ*murZ* Δ*murA*//$P_{Zn}$-*murA*$^{+}$ (IU16332), Δ*murA* Δ*murZ*//$P_{Zn}$-*murZ*$^{+}$ (IU16330) strains were grown overnight in BHI containing 0.15 or 0.25 mM $Zn^{2+}$ ($Zn^{2+}$/(1/10)$Mn^{2+}$) for IU16332 and IU16330, respectively, and diluted into BHI broth containing the indicated ($Zn^{2+}$/(1/10)$Mn^{2+}$) concentration. (A) Growth curve and (B) microscopic images taken between 3 to 4 h of growth.

Fig. S14

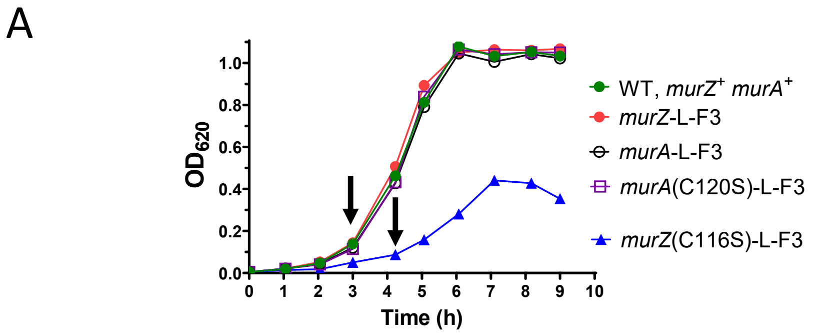

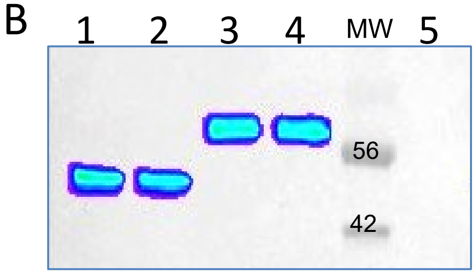

| Lane | Genotype |
|---|---|
| 1 | *murZ*-L-FLAG³ |
| 2 | *murZ*(C116S)-L-FLAG³ |
| 3 | *murA*-L-FLAG³ ) |
| 4 | *murA*(C120S)-L-FLAG³ |
| 5 | *murZ*⁺ *murA*⁺ WT |

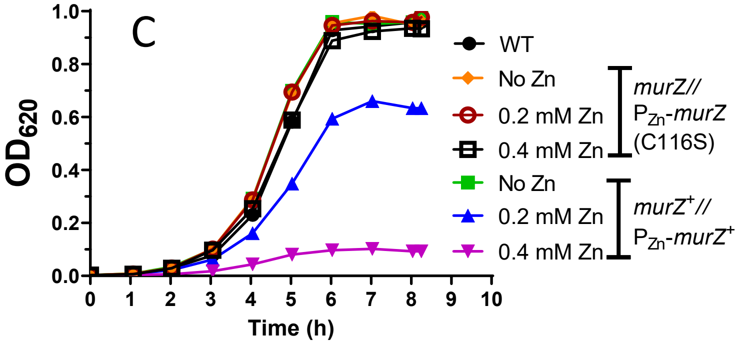

| | DT (min) | Yield | n |
|---|---|---|---|
| WT | 39 ± 0 | 1.0 ± 0.0 | 2 |
| *murZ*//$P_{Zn}$-*murZ*(C116S), No Zn | 37 ± 2 | 1.0 ± 0.0 | 2 |
| *murZ*//$P_{Zn}$-*murZ*(C116S), 0.2 mM Zn | 39 ± 0 | 1.0 ± 0.0 | 2 |
| *murZ*//$P_{Zn}$-*murZ*(C116S), 0.4 mM Zn | 41 ± 1 | 0.9 ± 0.0 | 2 |
| *murZ*⁺//$P_{Zn}$-*murZ*⁺, No Zn | 38 ± 1 | 1.0 ± 0.0 | 2 |
| *murZ*⁺//$P_{Zn}$-*murZ*⁺, 0.2 mM Zn | 42 ± 1 | 0.7 ± 0.0 | 2 |
| *murZ*⁺//$P_{Zn}$-*murZ*⁺, 0.4 mM Zn | 61 ± 6 | 0.1 ± 0.0 | 2 |

**Figure S15. Characterization of mutant strains with catalytic site mutations *murZ*(C116S) or *murA*(C120S).** (A) Growth curves in BHI broth of non-FLAG-tagged *murZ*⁺ WT (IU1824), *murZ*-L-FLAG³ (IU13502), *murZ*(C116S)-L-FLAG³ (IU15941), *murA*-L-FLAG³ (IU14028), and *murA*(C120S)-L-FLAG³ (IU15951). *murZ*(C116S)-L-FLAG³ strain showed a defective growth profile in BHI, similar to Δ*murZ* strains (see Fig. 6A). Arrows indicate when samples were withdrawn for protein preparation. (B) Western blot using an anti-FLAG antibody of protein samples prepared from above strains. C) Overexpression of MurZ in a *murZ*⁺//$P_{Zn}$-*murZ*⁺ strain (IU13393) resulted in growth inhibition in BHI broth, while overexpression of catalytically inactive *murZ*(C116S) in a *murZ*⁺//$P_{Zn}$-*murZ*(C116S) strain had no effect on growth.

Fig. S15

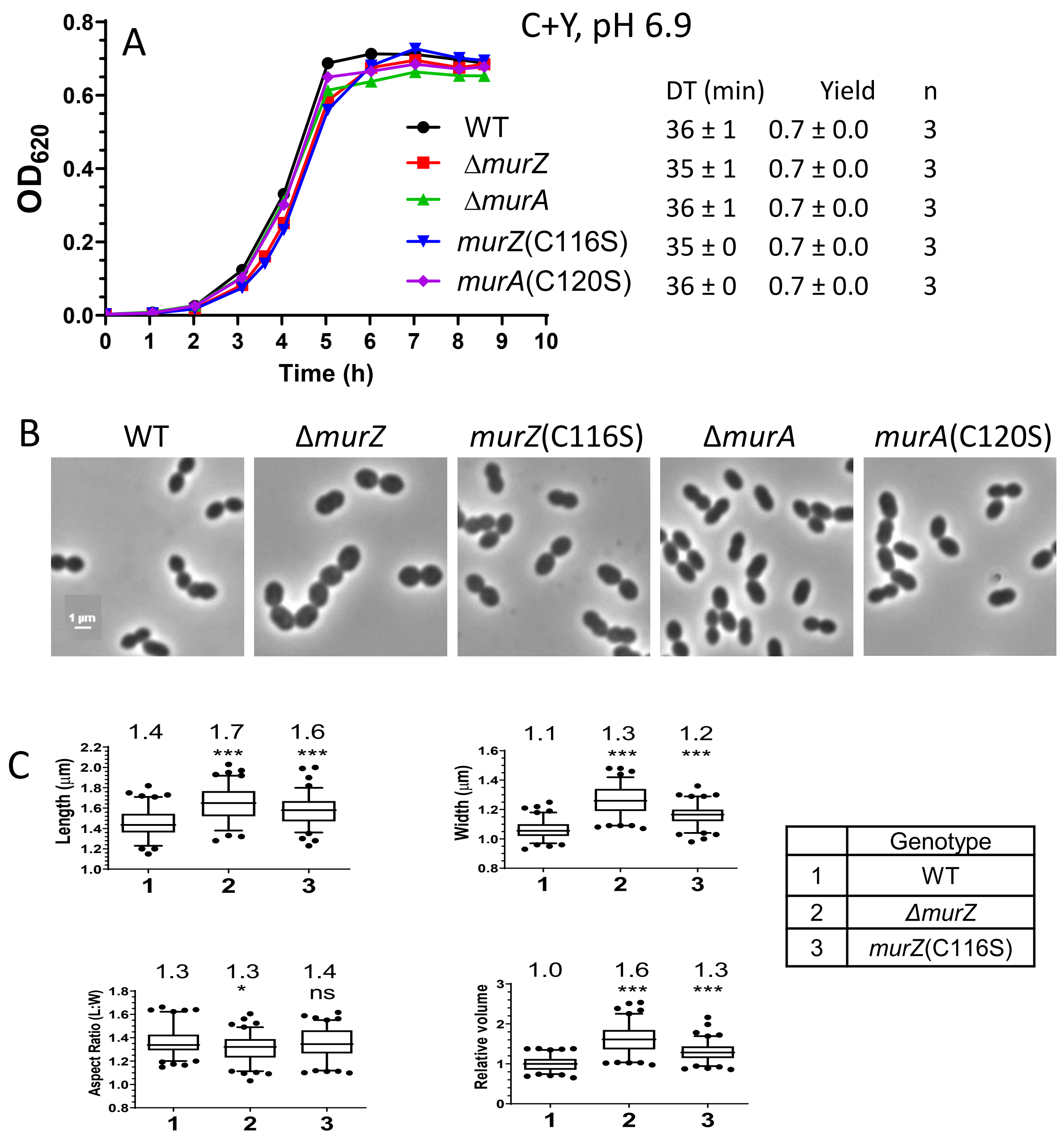

**Figure S16. Δ*murZ* or *murZ*(C116S) mutants cultured in C+Y media, pH 6.9 do not show growth defects, but form significantly enlarged cells compared to WT.** (A) Growth curves of WT (IU1824), Δ*murA* (IU13538), *murA*(C120S) (IU15949), Δ*murZ* (IU13536), and *murZ*(C116S) (IU15939). Strains were grown overnight in BHI broth, centrifuged to remove BHI, and resuspended in C+Y, pH 6.9 medium to $OD_{620}$ ≈ of 0.003 for growth curves. (B) Cells were imaged $OD_{620}$ ≈ of 0.1 to 0.15. Scale bar = 1 μm. (C) Box-and-whisker plots (whiskers, 5 and 95 percentile) of cell lengths, widths, aspect ratios (cell length to width) and relative cell volumes of 100 cells for each strain from two experiments. P values were obtained by one-way ANOVA analysis (GraphPad Prism, Kruskal-Wallis test). *, **, *** and ns denote p<0.05, p<0.01, p<0.001, not significant, respectively when compared to WT.

Fig. S16

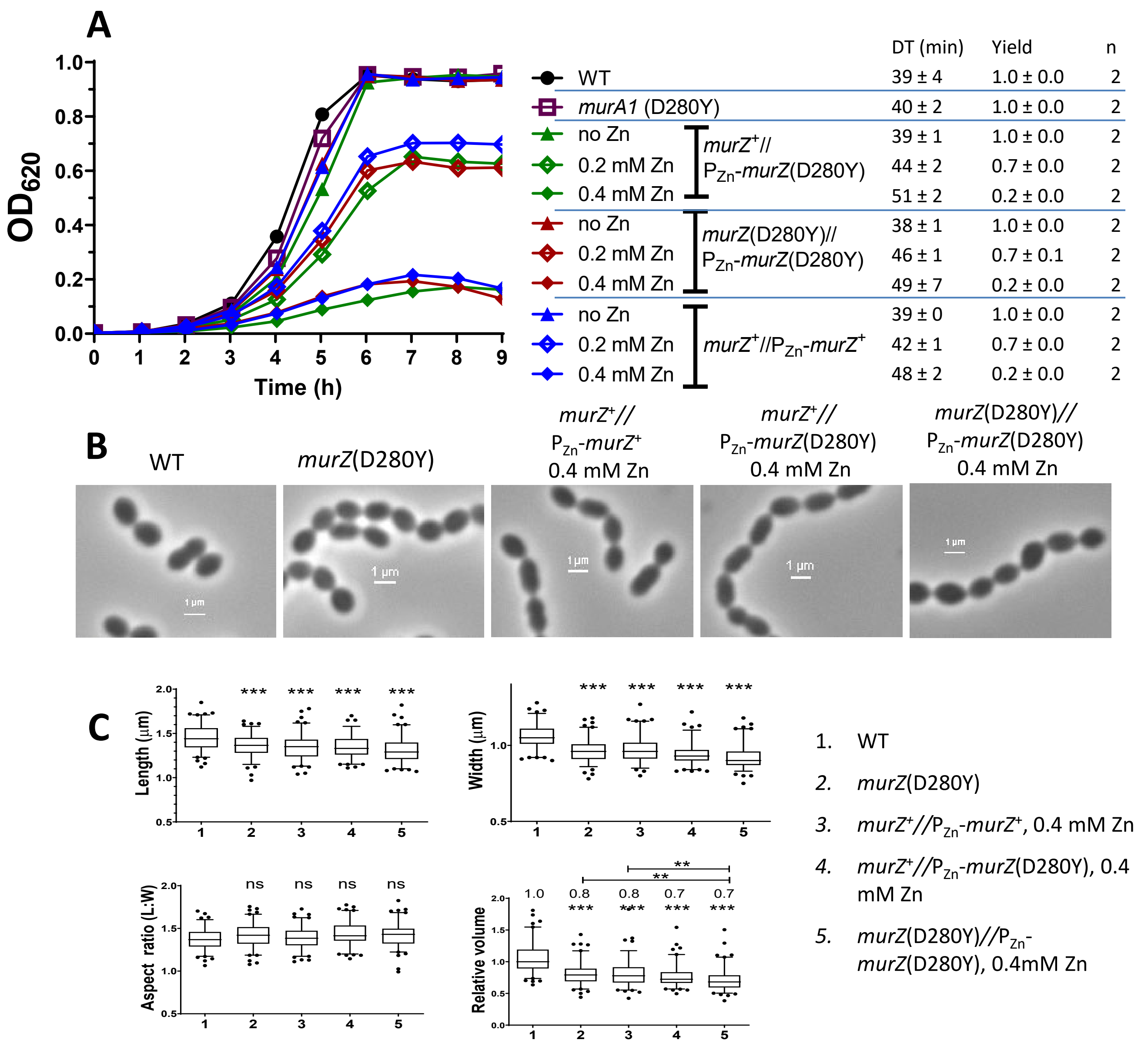

**Figure. S17. Ectopic overexpression of MurZ(D280Y) decreases cell size and inhibits growth similar to overexpression of MurZ.** Parent D39 Δ*cps rpsL1* strain (IU1824), *murZ*(D280Y) (IU13438), *murZ*+//P$_{Zn}$-*murZ*+ (IU13393), *murZ*+//P$_{Zn}$-*murZ*(D280Y) (IU16334), and *murZ*(D280Y)//P$_{Zn}$-*murZ*(D280Y) (IU16336) strains were grown overnight in BHI broth with no additional ($Zn^{2+}$/(1/10)$Mn^{2}$+), diluted to $OD_{620}$ ≈ 0.003 in the morning with no additional, 0.2 mM, or 0.4 mM ($Zn^{2+}$/(1/10)$Mn^{2+}$). (A) Representative growth curves, averages and SEMs of doubling times and maximal growth yields ($OD_{620}$) during 9 hours of growth. n denotes number of independent growths. (B) Representative phase-contrast images. (C) Box-and-whisker plots of cell dimension. P values were obtained by one-way ANOVA analysis (GraphPad Prism, Kruskal-Wallis test). *, **, *** and ns denote p<0.05, p<0.01, p<0.001, not significant, respectively when compared to WT.

Fig. S17

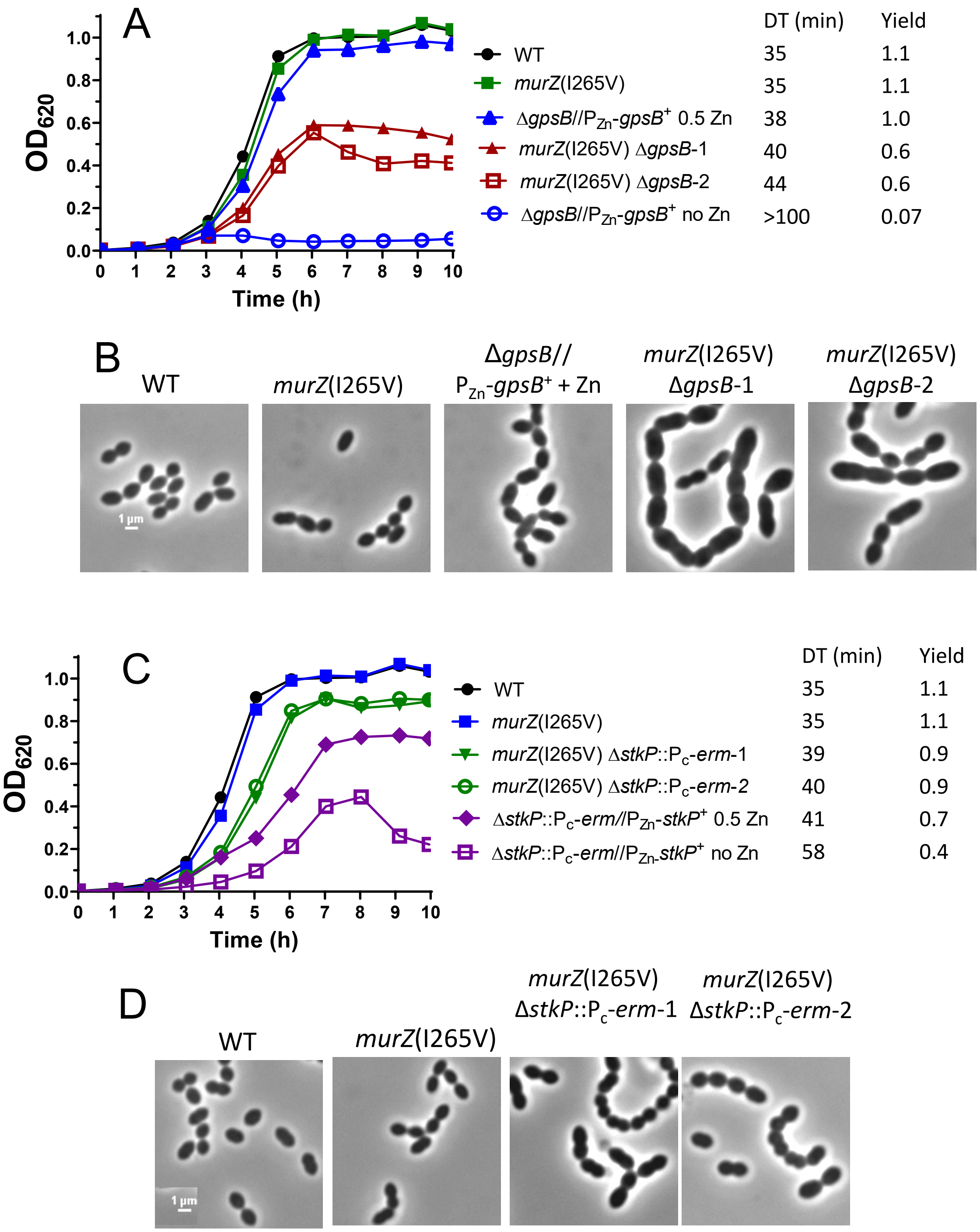

Fig. S18A to S18D

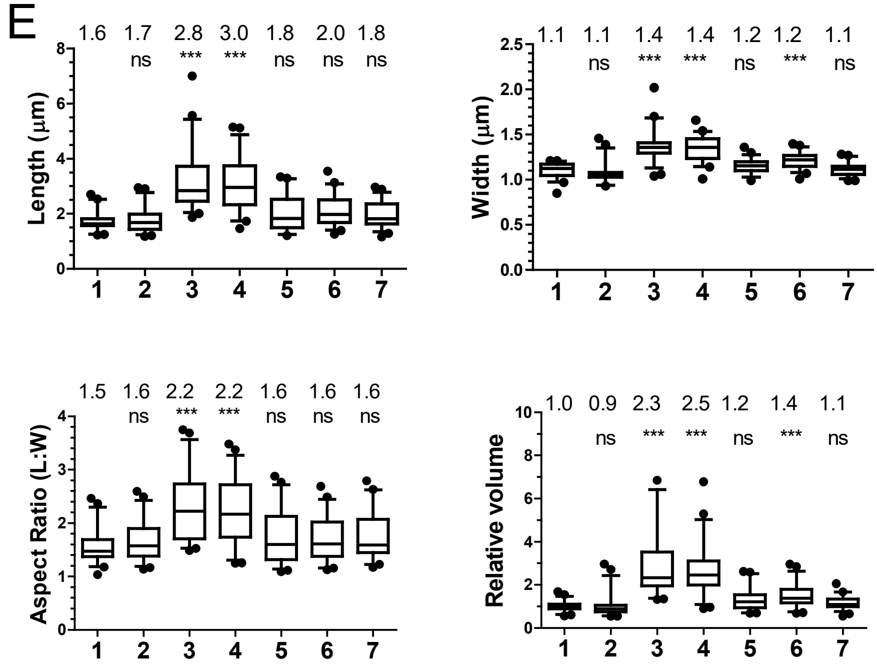

| | Genotype |
|---|---|
| 1 | WT |
| 2 | *murZ*(I265V) |
| 3 | *murZ*(I265V) Δ*gpsB-1* |
| 4 | *murZ*(I265V) Δ*gpsB-2* |
| 5 | *murZ*(I265V) Δ*stkP*::$P_c$-*erm*-1 |
| 6 | *murZ*(I265V) Δ*stkP*::$P_c$-*erm*-2 |
| 7 | Δ*gpsB//*$P_{Zn}$-*gpsB*$^+$*,* 0.5 mM Zn |

**Figure. S18. Suppression of Δ*gpsB* and Δ*stkP* lethality by *murZ*(I265V).** (A) Growth curves of WT (IU1824), *murZ*(I265V) (IU14210), two independent isolates of *murZ*(I265V) Δ*gpsB* (IU14234, IU15124), and Δ*gpsB*//$P_{Zn}$-*gpsB*$^+$ (IU16370) strains. Strains were grown overnight and diluted for growth during the day in BHI broth as described in *Experimental procedures*. IU16370 was grown overnight in BHI broth with 0.5 mM ($Zn^{2+}$/(1/10)$Mn^{2+}$), and diluted to $OD_{620}$ ≈ 0.003 in the morning with fresh BHI not supplemented with ($Zn^{2+}$/(1/10)$Mn^{2+}$) or containing 0.5 mM ($Zn^{2+}$/(1/10)$Mn^{2+}$). (B) Microscopic images of cells in (A) grown to $OD_{620}$ ≈0.15. (C) Growth curves of WT (IU1824), *murZ*(I265V) (IU14210), two independent isolates of *murZ*(I265V) Δ*stkP*::$P_c$-*erm* (IU17469, IU17475), and Δ*stkP*::$P_c$-*erm*//$P_{Zn}$-*stkP*$^+$ (IU16933) grown similar to IU16370. (D) Microscopic images of cells in (C) grown to OD ≈ 0.15. Scale bar = 1 µm. (E) Box-and-whisker plots (whiskers, 5 and 95 percentile) of cell lengths, widths, aspect ratios (cell length to width) and relative cell volumes of strains shown in (B) and (D). P values were obtained by one-way ANOVA analysis (GraphPad Prism, Kruskal-Wallis test). *, **, *** and ns denote $p<0.05$, $p<0.01$, $p<0.001$, not significant, respectively when compared to WT.

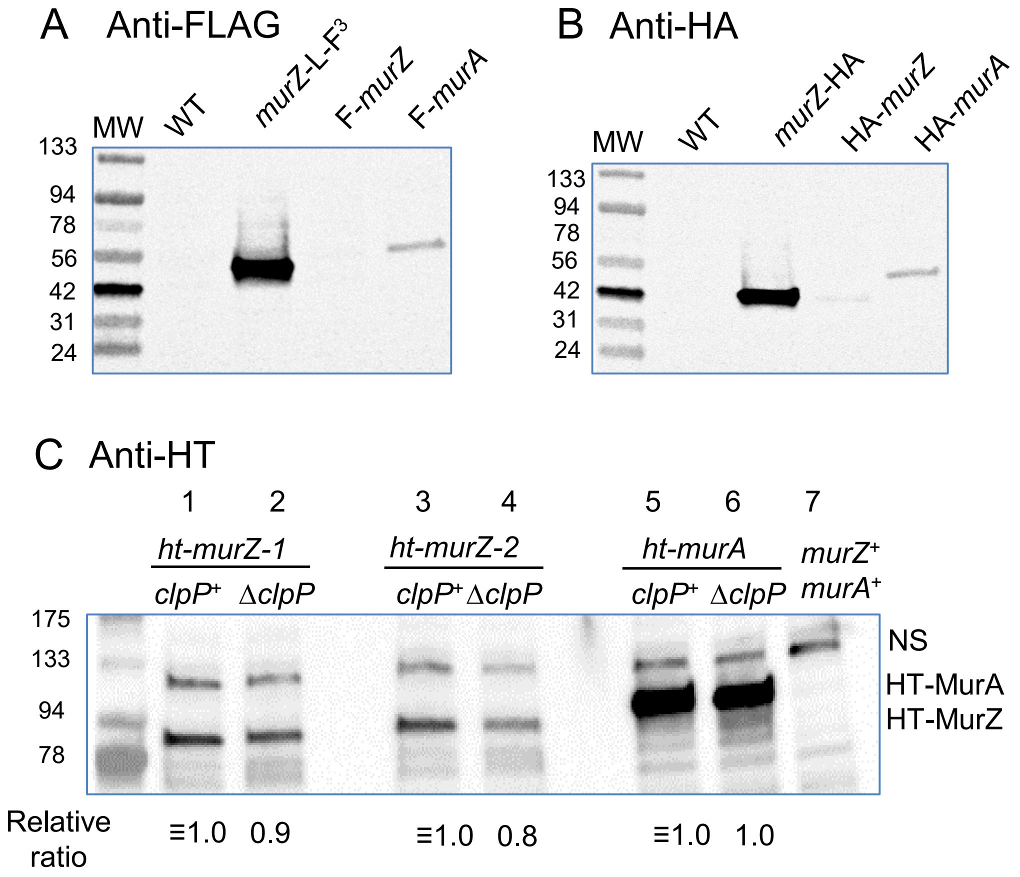

**Figure S19. Cellular amounts of N- or C-terminal-tagged MurZ and MurA fusion proteins are unchanged in Δ*clpP* mutants.** (A and B) Stable expression of C-terminal tagged MurZ (MurZ-L-$F^3$) and MurZ (MurZ-HA) compared to N-terminal tagged MurZ (F-MurZ, HA-MurZ) and MurA (F-MurA, HA-MurA). (A) Western blot results with lysates obtained from strains IU1824 (WT), IU13502 (*murZ*-L-$F^3$), IU17764 (F-*murZ*), and IU17768 (F-*murA*). (B) Western blot results of lysates obtained from strains IU1824 (WT), IU17170 (*murZ*-HA), IU17766 (HA-*murZ*), and IU17770 (HA-*murA*). (C) Western blot showing that HT-MurZ and HT-MurA cellular amounts are similar in *clpP*$^+$ and Δ*clpP* strains. Lysates were obtained from strains IU17838 (*ht-murZ* isolate 1, lane 1), IU17865 (*ht-murZ* Δ*clpP* isolate 1, lane 2), IU17840 (*ht-murZ* isolate 2, lane 3), IU17869 (*ht-murZ* Δ*clpP,* isolate 2 lane 4), IU17841 (*ht-murA,* lane 5), IU17869 (*ht-murZ* Δ*clpP,* lane 6) and WT (lane 7). Antibodies used for detection are described in *Experimental procedures* and signals were detected with an Azure Biosystem 600. 9 µg of crude lysate was loaded on each lane for A-C.

Fig. S19

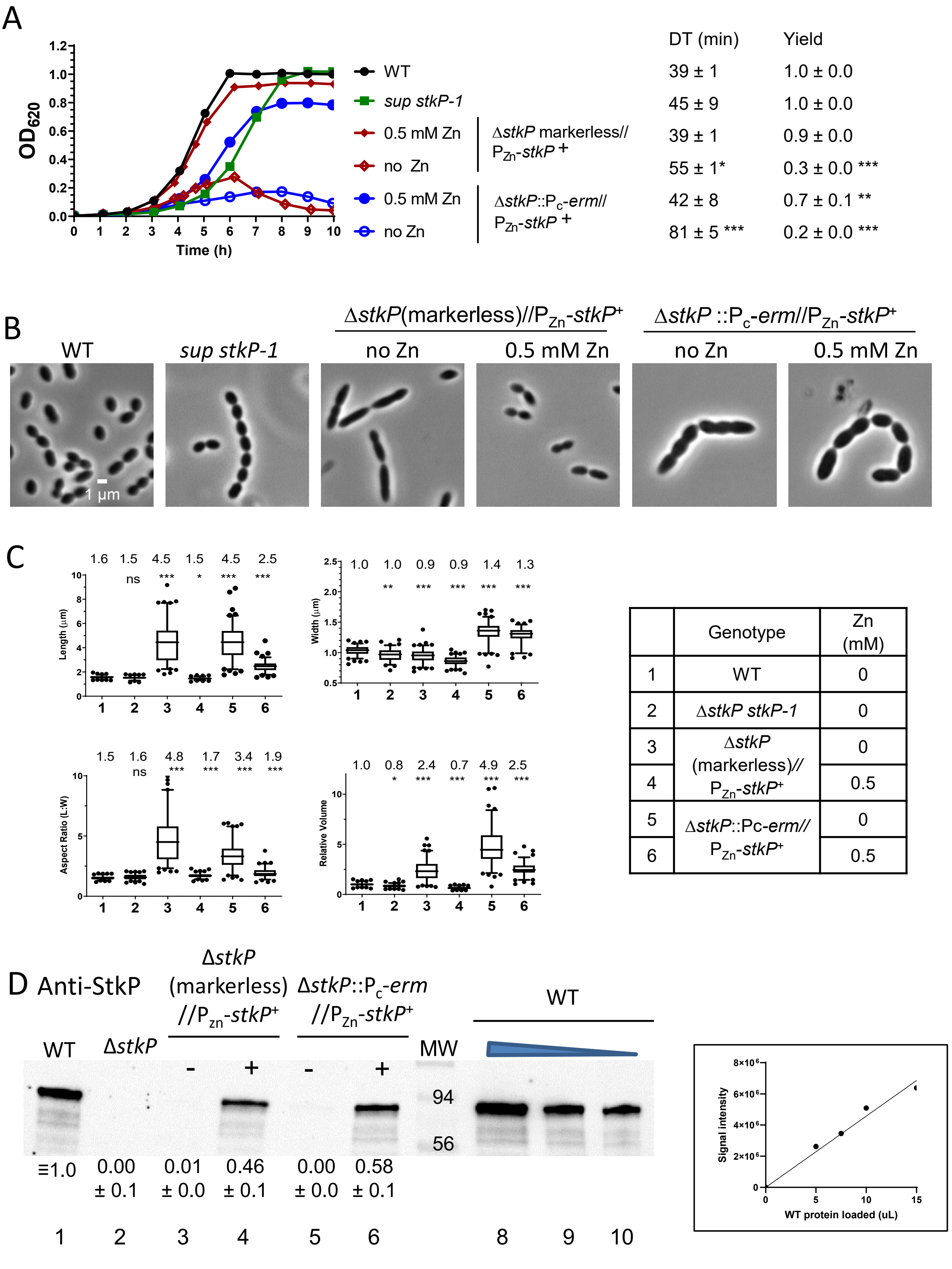
A
OD620
Time (h)
WT
sup stkP-1
0.5 mM Zn
no Zn
0.5 mM Zn
no Zn
ΔstkP markerless// PZn-stkP+
ΔstkP::Pc-erm// PZn-stkP+
DT (min)
Yield
39 ± 1
1.0 ± 0.0
45 ± 9
1.0 ± 0.0
39 ± 1
0.9 ± 0.0
55 ± 1*
0.3 ± 0.0 ***
42 ± 8
0.7 ± 0.1 **
81 ± 5 ***
0.2 ± 0.0 ***
B
WT
sup stkP-1
ΔstkP(markerless)//PZn-stkP+
ΔstkP ::Pc-erm//PZn-stkP+
no Zn
0.5 mM Zn
no Zn
0.5 mM Zn
1 μm
C
Length (μm)
1.6 1.5 4.5 1.5 4.5 2.5
ns *** * *** ***
Width (μm)
1.0 1.0 0.9 0.9 1.4 1.3
** *** *** *** ***
Aspect Ratio (L:W)
1.5 1.6 4.8 1.7 3.4 1.9
ns *** *** *** ***
Relative Volume
1.0 0.8 2.4 0.7 4.9 2.5
* *** *** *** ***
Genotype
Zn (mM)
1 WT 0
2 ΔstkP stkP-1 0
3 ΔstkP (markerless)// PZn-stkP+ 0
4 0.5
5 ΔstkP::Pc-erm// PZn-stkP+ 0
6 0.5
D Anti-StkP
WT
ΔstkP
ΔstkP (markerless) //Pzn-stkP+
ΔstkP::Pc-erm //PZn-stkP+
MW
WT
- + - +
94
56
≡1.0
0.00 ± 0.1
0.01 ± 0.0
0.46 ± 0.1
0.00 ± 0.0
0.58 ± 0.1
1 2 3 4 5 6 8 9 10
Signal intensity
WT protein loaded (uL)

Fig. S20

**Figure S20. Complementation of Δ*stkP* growth and morphological defects by ectopic overexpression of StkP*.*** Parent D39 Δ*cps rpsL1* strain (IU1824), *sup stkP-1* (IU16883), Δ*stkP*(*markerless)//*$P_{Zn}$-*stkP*$^+$ (IU18665) and Δ*stkP*::$P_c$-*erm//*$P_{Zn}$-*stkP*$^+$ (IU16933) strains were grown overnight in BHI broth with no additional ($Zn^{2+}$/(1/10)$Mn^{2+}$) (IU1824 and IU16883) or with 0.5 mM ($Zn^{2+}$/(1/10)$Mn^{2+}$) (IU18665 and IU16933) as described in *Experimental procedures*. Strains were diluted to $OD_{620}$ ≈ 0.003 in the morning with fresh BHI containing no ($Zn^{2+}$/(1/10)$Mn^{2+}$) or indicated concentrations of ($Zn^{2+}$/(1/10)$Mn^{2+}$). (A) Growth curves, doubling times, and maximal growth yields ($OD_{620}$) during 9 h of growth. (B) Representative phase-contrast images taken between 3.5 to 4 h of growth. Scale bar = 1 µm. Growth curves and microscopy were performed in two independent experiments. (C) Box-and-whisker plots (whiskers, 5 and 95 percentile) of cell lengths, widths, aspect ratios, and relative cell volumes. P values were obtained by one-way ANOVA analysis (GraphPad Prism, Kruskal-Wallis test). *, **, *** and ns denote p<0.05, p<0.01, p<0.001, not significant, respectively when compared to WT. (D) Quantitative Western blot with anti-StkP antibody showing relative StkP levels inducted by an ectopic Zn-controlled promoter. Lane 1, wild-type (WT, IU1824), 2, *sup stkP-1* (IU16883), 3-4, Δ*stkP*(markerless)//$P_{Zn}$-*stkP*$^+$ (IU18665) with 0 and 0.5 mM ($Zn^{2+}$/(1/10)$Mn^{2+}$), respectively, and 5-6 Δ*stkP*::$P_c$-*erm//*$P_{Zn}$-*stkP*$^+$ (IU16933) with 0 and 0.5 mM ($Zn^{2+}$/(1/10)$Mn^{2+}$), respectively. Samples were normalized based on culture $OD_{620}$ before addition of lysis buffer, and 10 µL (≈3 µg) of lysate were loaded in lanes 1-6. Lane 7, molecular weight standard. Lanes 8-10, 15, 7.5 and 5 µL of WT lysates, respectively, were used to generate the standard curve to the right. SDS-PAGE and western blotting were carried out as described in *Experimental procedures* using Licor IR Dye800 CW secondary antibody detected with Azure Biosystem 600. – or + indicates the absence of presence of ($Zn^{2+}$/(1/10)$Mn^{2+}$) in the BHI broth. Signals obtained with anti-StkP antibody were normalized with total protein stain in each lane using Totalstain Q-NC (Azure Scientific).

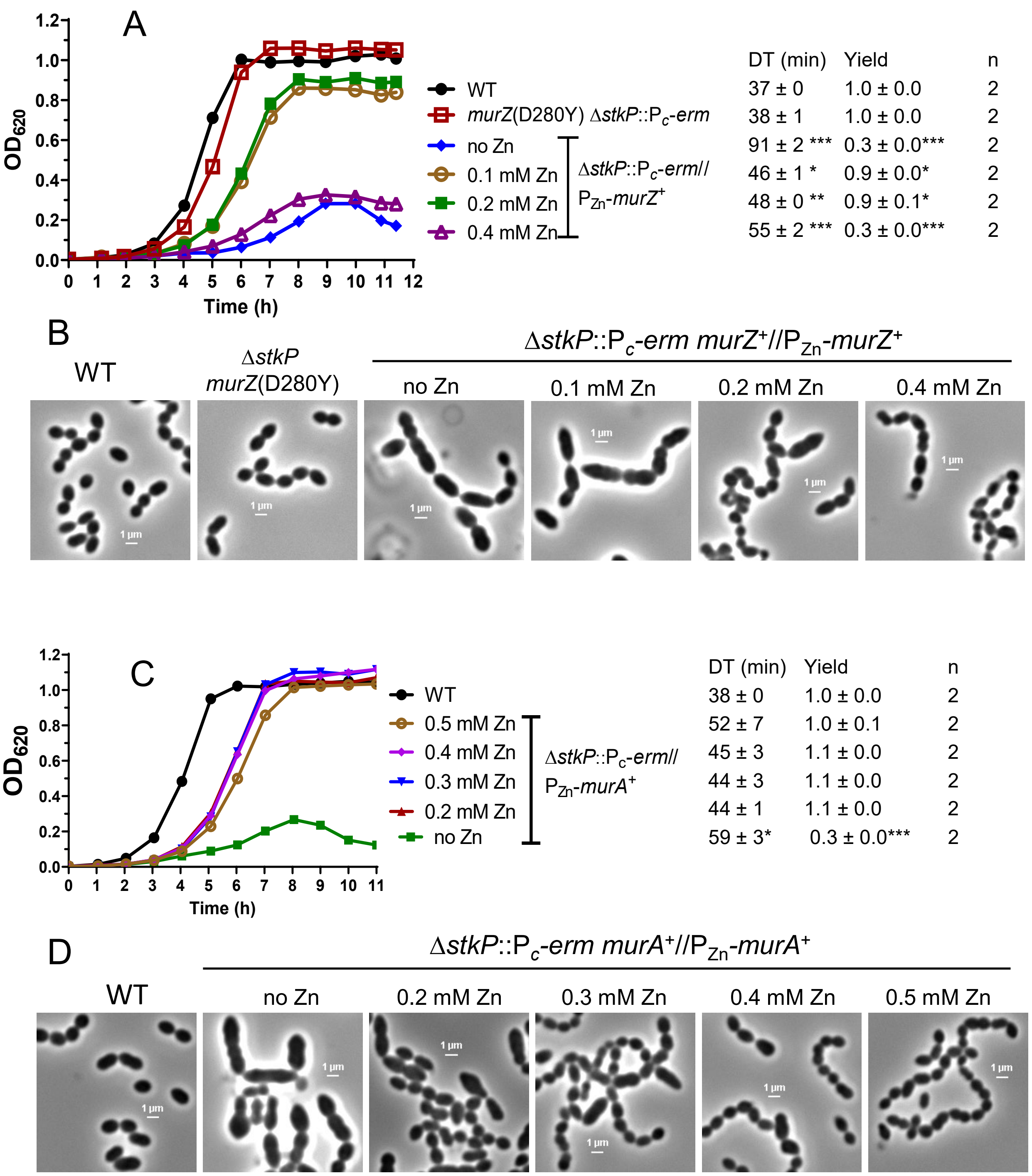

Fig. S21A to S21D

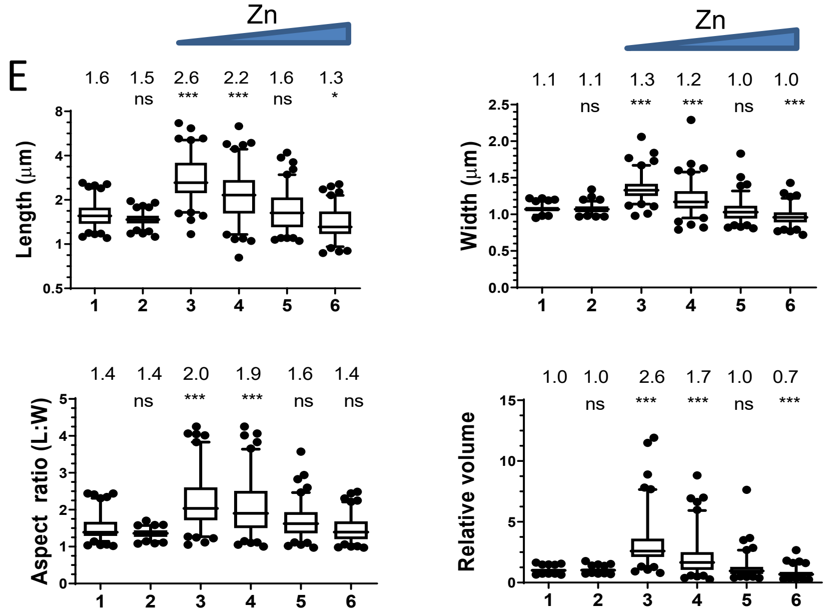

| | Genotype | Zn (mM) |
|---|---|---|
| 1 | WT | 0 |
| 2 | *murZ* (D280Y) Δ*stkP*::P$_c$-*erm* | 0 |
| 3 | Δ*stkP*::P$_c$-*erm*// P$_{Zn}$-*murZ*$^+$ | 0 |
| 4 | | 0.1 |
| 5 | | 0.2 |
| 6 | | 0.4 |

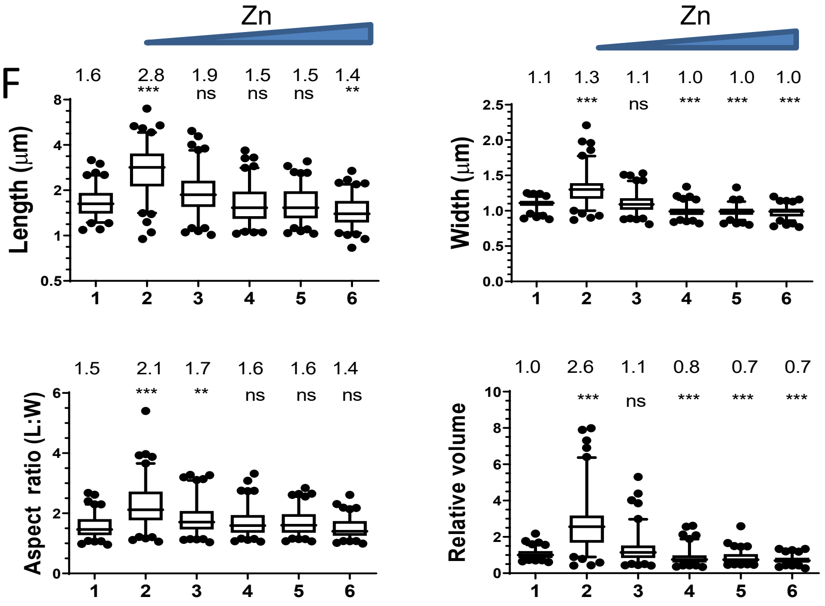

| | Genotype | Zn (mM) |
|---|---|---|
| 1 | WT | 0 |
| 2 | Δ*stkP*::P$_c$-*erm*// P$_{Zn}$-*murA*$^+$ | 0 |
| 3 | | 0.2 |
| 4 | | 0.3 |
| 5 | | 0.4 |
| 6 | | 0.5 |

Fig. S21E and S21F

**Figure S21. Suppression of growth and morphological Δ*stkP* phenotypes in BHI broth by *murZ*(D280Y) or overexpression of MurZ or MurA. (**A and B) Parent D39 Δ*cps rpsL1* strain (IU1824), *murZ*(D280Y) Δ*stkP*::Pc-*erm* (IU16885), and Δ*stkP*::Pc-*erm murZ*$^{+}$//$P_{Zn}$-*murZ*$^{+}$ (IU16897) were grown overnight in BHI broth without (IU1824 and IU16885) or with 0.2 mM ($Zn^{2+}$/(1/10)$Mn^{2+}$) . Overnight cultures were diluted in the morning in BHI for IU1824 and IU16885, and in BHI supplemented with 0 to 0.4 mM ($Zn^{2+}$/(1/10)$Mn^{2+}$) for IU16897. (A) Representative growth curves, averages, and SEMs of doubling times (DT) and maximal growth yields ($OD_{620}$). (B) Representative phase-contrast images taken between at 3.5 h for IU1824 and IU16885, and between 5 to 5.8 h for IU16897. Similar growth curves and morphology results were obtained with an independent Δ*stkP murZ*(D280Y) isolate, IU16895. (C) Parent D39 Δ*cps rpsL1* strain (IU1824) and a Δ*stkP*::$P_c$-*erm murA*$^{+}$//$P_{Zn}$-*murA*$^{+}$ (IU16915) strain were grown overnight in BHI broth with no or 0.4 mM ($Zn^{2+}$/(1/10)$Mn^{2+}$), respectively. Overnight cultures were diluted to $OD_{620} \approx 0.003$ in the morning in BHI for IU1824, and in BHI supplemented with 0 to 0.5 mM ($Zn^{2+}$/(1/10)$Mn^{2+}$) for IU16915. (D) Representative phase-contrast images taken at 3 h for IU1824, and between 4 to 5 h for IU16915. (E) and (F) Box-and-whisker plots (whiskers, 5 and 95 percentile) of cell lengths, widths, aspect ratios, and relative cell volumes of above strains grown with or without ($Zn^{2+}$/(1/10)$Mn^{2}$+). P values were obtained by one-way ANOVA analysis (GraphPad Prism, Kruskal-Wallis test). *, **, *** and ns denote p<0.05, p<0.01, p<0.001, not significant, respectively when compared to WT.
